# Towards high-resolution modeling of small molecule – ion channel interactions

**DOI:** 10.1101/2024.04.02.587818

**Authors:** Brandon J. Harris, Phuong T. Nguyen, Guangfeng Zhou, Heike Wulff, Frank DiMaio, Vladimir Yarov-Yarovoy

## Abstract

Ion channels are critical drug discovery targets for a wider range of pathologies, such as epilepsy, chronic pain, itch, autoimmunity, and cardiac arrhythmias. To develop effective and safe therapeutics, it is necessary to design small molecules with high potency and selectivity for specific ion channel subtypes. There has been increasing implementation of structure-guided drug design for the development of small molecules targeting ion channels. We evaluated the performance of two Rosetta ligand docking methods, RosettaLigand and GALigandDock, on structures of known ligand - cation channel complexes. Ligands were docked to voltage-gated sodium (Na_V_), voltage-gated calcium (Ca_V_), and transient receptor potential vanilloid (TRPV) channel families. For each test case, RosettaLigand and GALigandDock methods were able to frequently sample a ligand binding pose within 1-2 Å root mean square deviation (RMSD) relative to the native ligand – channel structure. However, RosettaLigand and GALigandDock scoring functions cannot consistently identify native drug binding poses as top-scoring models. Our study reveals that the proper scoring criteria for RosettaLigand and GALigandDock modeling of ligand - ion channel complexes should be assessed on a case-by-case basis using sufficient ligand and receptor interface sampling, knowledge about state specific interactions of the ion channel and inherent receptor site flexibility that could influence ligand binding.

## Introduction

The voltage-gated cation channel families consist of pore-forming transmembrane proteins that selectively conduct ions across lipid bilayers, and mediate physiological processes such as signal transduction, gene expression, synaptic transmission, and the activation and proliferation of cells in the immune system (Catterall, 1995; Hille, 2001; Catterall, 2011; Feske, Wulff, and Skolnik, 2015; Nanou and Catterall, 2018). Cation channels function in a finely regulated manner across spatial and temporal domains to complete these cellular functions. Current drug discovery efforts aim to modulate channel activity by targeting specific channel domains. For instance, a voltage-gated sodium channel’s therapeutically relevant structural domains are the selectivity filter (otherwise known as the outer pore vestibule), the central pore cavity (otherwise known as the inner pore vestibule), and the voltage-sensing domain (Nguyen and Yarov-Yarovoy, 2022). There have been considerable academic and industry efforts to identify therapeutically relevant small molecules that selectively target ion channels (Bagal et al., 2013; Zhang et al., 2020). However, developing effective and safe therapeutics targeting ion channels has been challenging (Wulff et al., 2019).

To address these challenges, drug discovery pipelines are trending towards incorporation of virtual drug screening and computer-aided drug design processes, for their ability to minimize drug development time and cost (Maia et al, 2020). Among these processes, molecular docking has demonstrated its usefulness for structure-based drug discovery. Molecular docking involves predicting conformations of the small molecule’s orientation with the protein (known as the pose) and scoring the poses to rank the likely protein-ligand interaction (Meng et al, 2011).

Among the numerous molecular docking software packages, Rosetta is a protein modeling software and design suite with two established small molecule docking methods: RosettaLigand (Meiler and Baker, 2006; Davis and Baker, 2009; Smith and Meiler, 2020) and GALigandDock (Park et al, 2021). RosettaLigand uses a Monte Carlo minimization procedure using the Rosetta energy function (Alford et al, 2017) to dock a pre-generated set of ligand conformers, while allowing side chain flexibility within a protein-receptor site. GALigandDock utilizes a different approach with two distinct features. First, the scoring function, RosetteGenFF, is a new generalized energy function tailored for small molecules. RosettaGenFF was trained from the Cambridge Structural Database (Groom et al, 2016), which at the time contained 1,386 small molecule crystal lattice arrangements to create a balanced force field that discriminates true lattice packing arrangements of the ligand from decoy (alternative lattice packing and conformational) arrangements. During docking, an orientation-dependent water-bridging energy term is incorporated within RosettaGenFF to further discriminate the protein-ligand orientation (Pavlovicz, Park, and DiMaio, 2020). Second, GALigandDock samples conformational space using a genetic algorithm. The ligand rigid body degrees of freedom and rotatable torsions are encoded as ‘genes’ to generate new ligand inputs for successive docking iterations. This allows efficient sampling of the protein-ligand energetic landscape when paired with the RosettaGenFF score function, canonical Monte Carlo optimization, and quasi-Newtonian minimization procedures within the Rosetta framework (Park et al, 2021).

While Rosetta protein-ligand docking methods are able to perform well with soluble protein – ligand benchmarks, the application of these methods to membrane-embedded ion channels has not been explored. Since there is a need to better assess and screen small molecules targeting different ion channel domains, we selected a diverse set of ten known cation channel-ligand structures for evaluation. From this set of high-resolution ion channel – small-molecule complexes, we assessed the accuracy of the RosettaLigand and GALigandDock methods in sampling ligand poses near the native structural coordinates and predicting the closest matching pose by energy ranking.

Our case studies include four voltage-gated sodium (Na_V_) channel structures, five voltage-gated calcium (Ca_V_) channel structures, and one transient receptor potential vanilloid (TRPV) channel structure. The ion channel ligand binding sites include the voltage-sensing domain, the selectivity filter, and the central pore cavity. Our results demonstrate that RosettaLigand and GALigandDock methods can frequently sample ligand binding poses within 1-2 Å root-mean-square deviation (RMSD) from the native channel – ligand structure. However, the ability to identify a near-native pose from energy ranking remains a challenge. When considering factors like the targeted ion channel domain, the ligand library features, and the sampling of ligand and receptor site conformations, our work demonstrates that high-resolution structures paired with RosettaLigand or GALigandDock can support drug discovery pipelines on a case-by-case basis.

## Material and Methods

### Ligand generation

Ligands were extracted as Structure Data Files (.sdf) from PubChem (Kim et al, 2023). Using the Avogadro software (Hanwell et al, 2012), each ligand structure underwent bond-correction, protonation at pH 7.4, and energy minimization using the Merck Molecular Force Field (Halgren, 1996a; b; c; Halgren and Nachbar, 1999; Halgren, 1999a; b). The resulting models were saved as Tripos Mol2 (.mol2) files. The protonation and bond order of saxitoxin and tetrodotoxin was matched to experimentally reported work (Hinman and DuBois, 2003; Thomas-Tran and Du Bois, 2016). Both experimentally resolved structures of verapamil docked to rabbit Ca_V_1.1 were tested (Zhao et al, 2019).

Next, using Amber Tools’ Antechamber protocol, the partial atomic charge, atom, and bond type assignments for each ligand were AM1-BCC corrected (Case et al, 2021; Salomon-Ferrer, Case, and Walker, 2012; **Appendix S1**). AM1-BCC correction is commonly used in Rosetta-based ligand docking protocols (Smith and Meiler, 2020; Park et al, 2021) and has demonstrated a similar performance correlation with other RosettaLigand input preparation protocols (Smith and Meiler, 2020).

The AM1-BCC corrected ligands were used for subsequent steps specific to each method. For RosettaLigand, an in-house script (**Appendix S2**) using the OpenEye Omega toolkit (Hawkins et al, 2010) was used to generate the conformer library, followed by Rosetta to generate the associated ligand parameters file. For GALigandDock, the input conformer was generated using the RosettaGenFF crystal structure prediction protocol (Park et al, 2021; **Appendix S3**), taking the lowest energy packing arrangement as input.

### Ion channel preparation

Ion channel structures were downloaded from the Protein Data Bank (Berman et al, 2000). Prior to RosettaLigand docking, structures were relaxed with backbone constraints using the RosettaRelax protocol (Nivón, Moretti, and Baker, 2013). This protocol allows the repacking of protein sidechains and minimization of the structure into the Rosetta score function for comparison between poses. The lowest energy pose from 100 relaxed poses was used for docking.

### RosettaLigand docking

RosettaLigand docking was performed using previously described RosettaScripts protocols (Davis and Baker, 2009; DeLuca, Khar, and Meiler, 2015; **Appendix S4-S7**). Briefly, the initial placement of all ligands into their respective ion channels was performed by superimposing the initial ligand poses onto the experimental ligand coordinates. RosettaLigand uses grid-based sampling to score the ion channel – ligand interface, whereby all ligand atoms and ion channel atoms a set distance from the ligand are scored with the defined scoring function. The scoring grid width was calculated uniquely for each ligand to ensure all ligand atoms were bound to the scoring grid. As used previously, the scoring grid width was calculated as the maximum conformer atom-atom distance plus twice the box size value used in the `Transform` mover (Moretti, Bender, Allison, and Meiler, 2016; **Appendix S4**).

RosettaLigand has a low-resolution and high-resolution sampling phase. For an unbiased sampling of the domain, an initial transformation during the low-resolution sampling phase is performed on the initial ligand position using Rosetta’s `Transform` mover with a box size of 7-8 Å. The low-resolution `Transform` step is a grid-based Monte Carlo simulation, where the ligand is translated up to 0.2 Å and rotated up to 20 degrees per iteration, for a total of 500 iterations. The pose with the lowest score is then used as the starting pose for high-resolution docking. For high-resolution docking using Rosetta’s `HighResDocker` mover, six docking cycles of rotamer sampling are performed with repacking of sidechains every third iteration. Lastly, the ion channel – ligand complex is minimized using Rosetta’s `FinalMinimizer` mover, and the interface scores are reported using the `InterfaceScoreCalculator`.

For all RosettaLigand docking runs, the `ligand_soft_rep` and `hard_rep` scoring functions were reweighted based on previous work assessing Rosetta score functions with the Comparative Assessment of Score Function 2016 (CASF-2016) dataset (Su et al., 2019; Smith and Meiler, 2020). Specifically, reweights of Coulombic electrostatic potential (`fa_elec`), Lennard-Jones repulsive energy between atoms in the same residue (`fa_intra_rep`), sidechain-backbone hydrogen bond energy (`hbond_bb_sc`), sidechain-sidechain hydrogen bond energy (`hbond_sc`), and Ramachandran preferences (`rama`) were applied for the soft-repulsive and hard-repulsive docking phases (**Appendix S6**).

For each docking run, either 20,000 poses or 100,000 poses were generated to assess if there are statistically significant differences in nearest native pose RMSD. In RosettaLigand, a ligand interface is defined either by a representative ligand atom (a “neighbor atom”, defined as the geometric center of mass by default), or all ligand atoms, relative to all ion channel C_β_ atoms within a specified radius from the ligand (commonly default to 6 or 7 Å). For RosettaLigand, two mutually exclusive ligand area interface modes are available for scoring the pose interface: the ligand neighbor atom cutoffs mode (`add_nbr_radius=”True”`), and the all ligand atom cutoffs mode (`all_atom_mode=”True”`). In previous work, both modes were used (Moretti, Bender, Allison, and Meiler, 2016; Smith and Meiler, 2020), thus, both modes were evaluated for any differences in performance. Four individual RosettaLigand docking sets were performed for each PDB structure, by combining different pose totals and ligand area interface modes, for performance comparison.

### GALigandDock docking

As described in the original study, docking was performed using the RosettaScripts’ `GALigandDock` mover in DockFlex mode (Park et al, 2021; **Appendix S8-S10**). Replicate runs of GALigandDock were performed in parallel for each structure evaluated. Each run consisted of 20 generations with a pool of 100 poses, where each generation updates the pool by total energy. By default, GALigandDock outputs the top 20 structures from the final generation, however, for this study, the entire pool of poses were used. For each docking run, 20,000 poses (1,000 runs) or 100,000 poses (5,000 runs) were output with a padding value of 2 Å, 4 Å, or 7 Å to test for statistical differences in the lowest RMSD pose (RMSD_Min_). In total, six individual GALigandDock docking sets were performed for each PDB structure, by combining different pose totals and padding sizes, for performance comparison.

### P_Near_ calculation

P_Near_ is a quantitative metric to evaluate an energy funnel’s quality from a population of poses, with the lowest-scoring pose as the converged minima, or native state. P_Near_ was calculated as previously defined (Bhardwaj, Mulligan, Bahl, et al., 2016), and applied as previously described for ligand docking (Smith and Meiler, 2020), with the native state being the ligand structure coordinates. P_Near_ ranges from 0 (protein will not converge to native state) to 1 (protein will always converge to the native state).

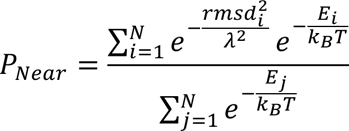

The numerator is the Boltzmann probability of an individual pose being near the native state, governed by the “native-ness” parameter (λ), and the thermal energy, the product of the Boltzmann constant and absolute temperature (k_B_T). The denominator is the partition function of a canonical ensemble. For small-molecule docking, the root mean square deviation (RMSD) of the pose’s ligand coordinates relative to the structure’s ligand coordinates when the protein pose is superimposed to the entire protein structure, or the receptor site is used. The energy scoring metric (E_i_) is the interface energy: the energy solely composed of protein-ligand interactions.

A previous Rosetta small-molecule docking study (Park et al., 2021) categorized native-like models at 1 Å or 2 Å RMSD without calculating P_Near_, while another study evaluating RosettaLigand performance calculated P_Near_ using λ =1.5 Å, and k_B_T =0.62 (Meiler and Smith, 2020). Therefore, we calculated P_Near_ with native-like models defined by λ=2.0 Å and k_B_T =0.62 Rosetta energy units (REU), however we calculated P_Near_ using all previously reported values for the parameters λ and k_B_T (**Tables S4.1-4.10**). A specific P_Near_ cutoff indicative for drug discovery pipelines has not been established, hence we will refer to P_Near_ ≥ 0.5 as a ‘first pass’ cutoff for this study when evaluating energy funnel convergence to the ligand structure coordinates.

### Statistical tests

Tests for normality, heteroscedasticity, and Pearson’s correlation between covariates were performed in Python using NumPy (Harris et al., 2020), SciPy (Virtanen et al., 2020), and Pingouin (Vallat, 2018) with a significance level (α) of 0.05. Population data consisted of the RMSD_Min_ for each docking set. Not all RMSD_Min_ data for each docking set fit a normal distribution. Since RMSD_Min_ data should skew towards zero Å, a lognormal base 10 transformation was applied to all RMSD_Min_ population data when comparing across docking sets. Shapiro Wilk tests for normality and Levene heteroscedasticity tests were performed to ensure transformed data were normal and of equal variance, respectively.

Covariates for both methods were the number of rotatable bonds of the ligand, the number of ligand heavy atoms, and the resolution of the structure. Statistical tests for RosettaLigand also included the number of conformers generated, transformed with a lognormal base 10, as a covariate. The ligand molecular weight was initially included as a covariate but was discarded after identifying strong Pearson’s correlation with the number of ligand heavy atoms (**Table S3, Figure S1.**).

To assess RMSD_Min_ across all docking sets for a given method, a repeated measures factorial analysis of variance (ANOVA) with covariates were performed using IBM SPSS (version 29). For RosettaLigand, the two factors were sample size and ligand area interface mode, while for GALigandDock the two factors were sample size and padding value. Mauchly’s test for sphericity was performed for levels of padding (p = 0.63).

## Results

In this study, we evaluated ligands that are suggested to modulate ion channel activity for a given clinical effect, such as antiarrhythmic, anticonvulsant, and antihypertensive (Table 1). Our study also includes canonical ion channel blockers, such as tetrodotoxin and saxitoxin, small molecules whose structures have been resolved for drug discovery campaigns, such as GX-936, Z944, and Bay K 8544, and drugs currently approved for therapy, such as nifedipine, diltiazem, and flecainide (**Figure 1).**

**Figure 1.**
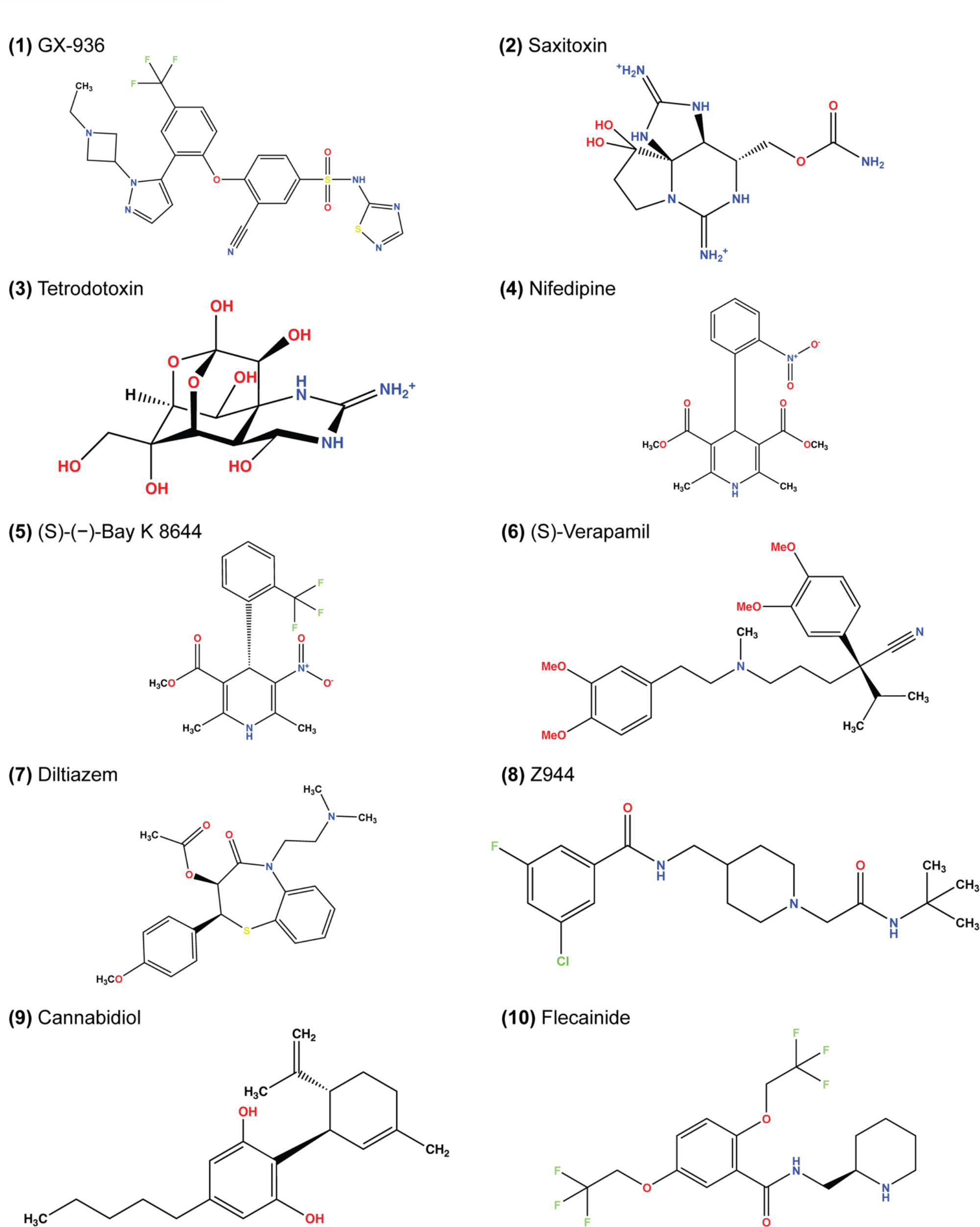
Two-dimensional structures of ligands docked in this study. The depicted stereochemistry reflects that resolved from structures. The protonation and bond order of saxitoxin and tetrodotoxin from prior reported work was used (Hinman and DuBois, 2003; Thomas-Tran and Du Bois, 2016).

**Table 1.** Summary of ligand – ion channel structures docked, domain, and covariates included in the study.

| PDB ID | Description | Ion channel modulation / therapeutic class | Domain | Total ligand rotatable bonds | Total ligand heavy atoms | Ligand molecular weight (Da) | Structure resolution (Å) | Total RosettaLigand Conformers |
| --- | --- | --- | --- | --- | --- | --- | --- | --- |
| 5EK0 | human Nav1.7-VSD4-NavAb / GX-936 | Inhibitor / NA | Voltage-sensing domain | 8 | 39 | 575.6 | 3.53 | 2179 |
| 6J8G | human Nav1.7 / Saxitoxin | Blocker / NA | Selectivity filter | 3 | 21 | 299.3 | 3.20 | 28 |
| 6J8I | human Nav1.7 / Tetrodotoxin | Blocker / NA | Selectivity filter | 1 | 22 | 319.3 | 3.20 | 2 |
| 6JP5 | rabbit Cav1.1 / Nifedipine | Blocker / vasodilator, antihypertensive, antianginal | CPC† | 5 | 25 | 346.3 | 2.90 | 22 |
| 6JP8 | rabbit Cav1.1 / (S)-(-)-Bay K 8644 | Activator / NA | CPC | 3 | 25 | 356.3 | 2.70 | 3 |
| 6JPA | rabbit Cav1.1 / (S)-Verapamil | Blocker / antiarrhythmic, vasodilator | CPC | 13 | 33 | 454.6 | 2.60 | 785 |
| 6JPB | rabbit Cav1.1 / Diltiazem | Blocker / antihypertensive, vasodilator | CPC | 7 | 29 | 414.5 | 2.90 | 45 |
| 6KZP | human Cav3.1 / Z944 | Blocker / NA | CPC | 6 | 26 | 383.9 | 3.10 | 1992 |
| 6U88 | rat TRPV2 / Cannabidiol | Activator / anticonvulsants | CPC | 6 | 23 | 314.5 | 3.20 | 71 |
| 6UZ0 | rat Nav1.5 / Flecainide | Blocker / antiarrhythmic class 1c | CPC | 7 | 28 | 414.3 | 3.24 | 1169 |

† CPC = Central pore cavity.

### Performance criteria for Rosetta ligand docking

A previous study has emphasized that two criteria must be satisfied for accurate modeling of protein-ligand interactions (Kaufmann and Meiler, 2012). First, the method must produce native-like poses through sufficient sampling. For small molecule docking, a native-like ligand pose has a root mean square deviation (RMSD) below a context-dependent predetermined value. Previously reported RMSD ranges when assessing a native-like pose are 1.0 Å or 2.0 Å, with 2.0 Å being a common cutoff for drug discovery pipelines (Maia et al., 2020; Park et al., 2021). When calculating RMSD, the protein pose is superimposed to the entire protein structure or the receptor site, and then the ligand coordinates of the pose are evaluated relative to the ligand coordinates of the structure.

Second, the pose population must produce an energy funnel converging to the lowest-energy, native-like pose (Meiler and Baker, 2006). Rather than the lowest-energy pose, we use empirical structural evidence in this study to define the native pose as the ligand’s structural coordinates. An energy funnel is qualitatively evaluated using the interface energy between ligand and protein with respect to ligand RMSD (Meiler and Baker, 2006, Smith and Meiler, 2020). A quantitative metric of funnel convergence, P_Near_, has been utilized in Rosetta protein design (Guarav, Mulligan, Bahl, et al., 2016; Mulligan et al., 2021), and has been adopted for small molecule docking (Smith and Meiler, 2020). Due to the large number of poses, funnel quality is often assessed with a population subset consisting of the best-scoring poses (Combs et al., 2010; Lemmon, Kaufmann, and Meiler, 2012; Shim et al., 2019).

With these criteria in mind, we evaluated the performance of RosettaLigand and GALigandDock for the following ion channel receptor sites: the voltage-sensing domain, the pore-forming domain, and the central pore cavity (**Table 1**). We evaluated RosettaLigand using combinations of sample size (20,000 poses vs 100,000 poses) and ligand area interface mode (ligand neighbor atom vs all ligand atom). We evaluated GALigandDock using combinations of sample size (20,000 poses vs 100,000 poses) and padding of the sampling grid (2 Å, 4 Å, and 7 Å). Increasing the padding of the sampling grid enables additional rotamer sampling around the receptor and increases translational sampling of the ligand around the receptor site.

### Covariates potentially influencing RMSD

We controlled for the following covariates that are dependent upon the ligand or structure used for docking. We speculated that the number of ligand rotatable bonds and the number of heavy atoms could influence RMSD_Min_ by increasing the amount of internal sampling required during the docking run. We also speculated that a poorer, higher reported structural resolution could result in an increased overall RMSD_Min_. The total number of ligand heavy atoms was also used as a covariate for RMSD_Min_, which is strongly correlated to the molecular weight of each ligand (**Figure S1**). Lastly, for RosettaLigand, the total number of conformers provided as input could affect the RMSD_Min_ since more conformers could inherently require more sampling.

### RosettaLigand and GALigandDock generate native-like ion channel-ligand poses

This study evaluated docking sets using population data consisting of the lowest RMSD pose (RMSD_Min_) from each channel – ligand test case. We provide four docking sets for RosettaLigand and six docking sets for GALigandDock, using combinations of the sample size with either the RosettaLigand-specific ligand area interface calculation, or GALigandDock-specific padding value. For brevity, we will discuss the RosettaLigand docking set using a sample size of 100,000 poses and the all-ligand atoms interface mode, and the GALigandDock docking set using a sample size of 100,000 poses and padding value of 7 Å. These specific docking sets provide the greatest breadth of receptor sampling and scoring among our docking sets by appropriately encompassing the entirety of each ion channel domain tested (**Table 2**). The results for other docking sets are provided in the Supplemental Material (**Tables S1.1-1.2**).

**Table 2.** The minimum RMSD (Å) pose for each ligand-channel docked that is detailed in the primary text.

| PDB ID | Description | RosettaLigand, all ligand atom interface cutoff, 100,000 poses | GALigandDock, padding 7 Å, 100,000 poses |
| --- | --- | --- | --- |
| **5EK0** | human Nav<sub>v</sub>1.7-VSD4-NavAb / GX-936 | 0.91 | 0.65 |
| **6J8G** | human Nav<sub>v</sub>1.7 / Saxitoxin | 0.70 | 0.76 |
| **6J8I** | human Nav<sub>v</sub>1.7 / Tetrodotoxin | 0.54 | 0.81 |
| **6JP5** | rabbit Cav<sub>v</sub>1.1 / Nifedipine | 0.77 | 1.3 |
| **6JP8** | rabbit Cav<sub>v</sub>1.1 / (S)-(-)-Bay K 8644 | 0.93 | 0.97 |
| **6JPA-1<sup>†</sup>** | rabbit Cav<sub>v</sub>1.1 / (S)-Verapamil | 1.4 | 2.0 |
| **6JPA-2** | rabbit Cav<sub>v</sub>1.1 / (S)-Verapamil | 2.2 | 2.1 |
| **6JPB** | rabbit Cav<sub>v</sub>1.1 / Diltiazem | 2.5 | 1.1 |
| **6KZP** | human Cav<sub>v</sub>3.1 / Z944 | 0.73 | 0.84 |
| **6U88** | rat TRPV2 / Cannabidiol | 1.0 | 0.90 |
| **6UZ0** | rat Nav<sub>v</sub>1.5 / Flecainide | 1.2 | 0.94 |
| **Average** |  | 1.2 ± 0.6 | 1.1 ± 0.5 |
| **Median** |  | 0.93 | 0.94 |

<sup>†</sup> Verapamil was resolved with two binding orientations to the rabbit Cav<sub>v</sub>1.1 central pore cavity in 6JPA and are named 6JPA-1 and 6JPA-2 in this study respectively.

For RosettaLigand, the average RMSD_Min_ across the docking set was 1.2 ± 0.6 Å (**Table 2**). When comparing RosettaLigand docking sets and using repeated measures ANOVA with covariates, an interaction from both factors (sample size and ligand area interface mode) did not result in rejection of the null hypothesis of equivalent means from common logarithm transformed RMSD_Min_ data (p=0.71). Likewise, individual interactions from sample size (p=0.38) and ligand area interface mode (p=0.72) did not result in rejection of the null hypothesis of equivalent means. Each RosettaLigand docking set produced comparable RMSD_Min_ averages and standard deviations within a few sub-Angstroms (**Table S1.1**). Therefore, our results suggest that using either ligand area interface mode with 20,000 or 100,000 total poses could generate similar near-native models.

For GALigandDock, the average RMSD_Min_ across the docking set was 1.1 ± 0.5 Å (**Table 2)**, while all docking sets produced similar RMSD_Min_ averages and standard deviations within a few sub-Angstroms (**Table S1.2**). Again, using repeated measures ANOVA with covariates, an interaction from both factors (sample size and padding size) failed to result in a rejection of the null hypothesis of equivalent means (p=0.91). Likewise, individual interactions from sample size (p=0.59) and padding size (p=0.34) did not result in rejection of the null hypothesis of equivalent means. While there was no statistical advantage to using a specific padding value to achieve a lowered average RMSD_Min_, we note that ion channel structures within the central pore cavity had greater sampling with a padding value of 7 Å, since a padding value of 5 Å did not encompass the entire pore. Therefore, the appropriate padding size is context dependent and should be adjusted when using GALigandDock to provide sufficient sampling grid space.

### RosettaLigand and GALigandDock energy funnels for ion channel-ligand docking

We evaluated whether the entire population of generated poses and the top 10% of the lowest total energy-scoring poses would achieve P_Near_ values indicative of an energy funnel converging onto the ligand structural coordinates (**Tables S4.1-4.10**). Since interface energy describes how a ligand is interacting with the ion channel, P_Near_ was calculated with the interface energy rather than the total energy. A specific P_Near_ cutoff indicative for drug discovery pipelines has not been established. Hence, we will refer to P_Near_ ≥ 0.5 as a ‘first pass’ cutoff for this study when evaluating energy funnel convergence to the ligand structure coordinates. For brevity, we will discuss P_Near_ with a “native-ness” parameter (λ) of 2.0, and a thermal energy parameter (k_B_T) of 0.62. Other P_Near_ values matching parameter values reported in other work are provided in the Supplemental Material (Bhardwaj, Mulligan, Bahl, et al., 2016; Smith and Meiler, 2020; **Tables S4.1-4.10**).

The P_Near_ of the full population and the top 10% of total energy-scoring poses were similar within methods, with the bulk of P_Near_ values being in the thousandths or lower (**Table 3**). However, RosettaLigand and GALigandDock were able to identify energy funnels for unique cases. For RosettaLigand, the human Ca_V_3.1-Z944 complex (PDB: 6KZP) achieved P_Near_ ≥ 0.5, while for GALigandDock, human Na_V_1.7-VSD4-Na_V_Ab - GX-936 (PDB: 5EK0) achieved P_Near_ ≥ 0.5. For both methods, rabbit Ca_V_1.1-Bay K8644 (PDB: 6JP8) achieved P_Near_ ≥ 0.5.

**Table 3.** P_Near_ of RMSD_Min_ and interface energy for each ligand-channel docked that is detailed in the primary text, calculated with k_B_T=0.62 and λ=2.0.

| PDB ID | Description | RosettaLigand, all ligand atom interface cutoff, 100,000 poses |  | GALigandDock, padding 7 Å, 100,000 poses |  |
| --- | --- | --- | --- | --- | --- |
|  |  | Full population | Top 10% total score | Full population | Top 10% total score |
| 5EK0 | human Nav1.7-VSD4-NavAb / GX-936 | 0.22 | 0.26 | 0.83 | 0.83 |
| 6J8G | human Nav1.7 / Saxitoxin | $1.2 \cdot 10^{-2}$ | $9.3 \cdot 10^{-3}$ | $1.6 \cdot 10^{-2}$ | $1.6 \cdot 10^{-2}$ |
| 6J8I | human Nav1.7 / Tetrodotoxin | 0.12 | $8.8 \cdot 10^{-2}$ | 0.27 | 0.27 |
| 6JP5 | rabbit Cav1.1 / Nifedipine | 0.21 | 0.28 | 0.24 | 0.29 |
| 6JP8 | rabbit Cav1.1 / (S)-(-)-Bay K 8644 | 0.62 | 0.66 | 0.60 | 0.61 |
| 6JPA-1 | rabbit Cav1.1 / (S)-Verapamil | $4.7 \cdot 10^{-3}$ | $4.3 \cdot 10^{-3}$ | $2.4 \cdot 10^{-3}$ | $2.8 \cdot 10^{-3}$ |
| 6JPA-2 | rabbit Cav1.1 / (S)-Verapamil | $5.0 \cdot 10^{-4}$ | $1.2 \cdot 10^{-4}$ | $1.8 \cdot 10^{-5}$ | $1.9 \cdot 10^{-5}$ |
| 6JPB | rabbit Cav1.1 / Diltiazem | $6.8 \cdot 10^{-4}$ | $6.4 \cdot 10^{-4}$ | $2.8 \cdot 10^{-6}$ | $7.8 \cdot 10^{-6}$ |
| 6KZP | human Cav3.1 / Z944 | 0.50 | 0.62 | $1.2 \cdot 10^{-4}$ | $1.1 \cdot 10^{-4}$ |
| 6U88 | rat TRPV2 / Cannabidiol | $8.3 \cdot 10^{-2}$ | $4.3 \cdot 10^{-2}$ | $5.0 \cdot 10^{-3}$ | $3.8 \cdot 10^{-3}$ |
| 6UZ0 | rat Nav1.5 / Flecainide | 0.14 | 0.12 | $5.9 \cdot 10^{-6}$ | $2.9 \cdot 10^{-5}$ |

Following the standard of reporting a percentage of poses (Combs et al., 2010; Lemmon, Kaufmann, and Meiler, 2012; Shim et al., 2019), specific P_Near_ values reported herein refer to the top 10% of total energy-scoring poses (**Table 3**).

Notably, there are visual distinctions in interface energy funnel plots between RosettaLigand and GALigandDock (**Figures S2-S12**). Generally, RosettaLigand can sample more poses up to 2 Å RMSD compared to GALigandDock, but with a less distinguishable energy funnel since some poses score greater than zero, indicating an unfavorable energy score. Since GALigandDock uses the lowest total energy poses as input for future conformer generations, poses with interface energy greater than zero are infrequent.

Lastly, when using the lowest interface energy pose from the full docking population as ranking criteria, the RMSD_Min_ increases to a range between 1.2 Å and 10.8 Å (mean 4.5 ± 2.8 Å) with RosettaLigand (**Table S2.1**), and a range between 0.83 Å and 14.5 Å (mean 6.4 ± 5.0 Å) with GALigandDock (**Table S2.5**). This suggests that extracting the lowest energy pose is not a reliable indicator of a near-native pose and does not necessarily reflect the native binding mode for ion channel-ligand docking.

### Ligand Docking into the voltage-sensing domain

The only ion channel-ligand structure evaluated for ligand docking into the voltage-sensing domain (VSD) was GX-936 in complex with the VSD4 of the hNa_V_1.7-Na_V_Ab chimera (PDB: 5EK0). The hNa_V_1.7 channel has been validated as a drug target for pain signaling, and aryl sulfonamides have been reported as selective inhibitors. Specifically, GX-936 exhibits selectivity for hNa_V_1.7 compared to other hNa_V_ isoforms (McCormack et al., 2013; Ahuja et al., 2015; Nguyen and Yarov-Yarovoy, 2022).

After docking GX-936 to the hNa_V_1.7 VSD4, the RMSD_Min_ poses were 0.91 Å using RosettaLigand and 0.65 Å using GALigandDock (**Figure 2**, **Table 2).** The largest deviations from the native structure were observed at the peripheral pyrazole ring and the ethyl azetidine (**Figure 2).** Notably, GX-936 has eight rotatable bonds, making it the second most flexible ligand used in this study **(Table 1**; **Figure 1)**. However, the sampled ligand poses were consistently below 2 Å RMSD for each docking run (**Table S1.1-1.2).** RosettaLigand was unable to identify the RMSD_Min_ pose as the lowest interaction energy pose (**Table S2.1**), scoring the lowest energy pose 2.7 Å from the structure coordinates. This pose has the same binding orientation, but the pyrazole ring clashes with the structurally resolved lipid, while the ethyl azetidine withing the VSD is orientated up towards the extracellular space, rather than down. Indeed, the lowest ten interface energy poses all possess these features (**Figure S13.1**). GALigandDock scored the lowest energy pose 0.83 Å from the structure coordinates (**Table S2.5**). Compared to RosettaLigand, the lowest ten interface energy poses converge well with the RMSD_Min_ pose, with only one of the ten poses clashing with the lipid (**Figure S13.2)**. Further, GALigandDock was able to discriminate with good confidence an energy funnel (P_Near_ = 0.83; **Table 3**), whereas RosettaLigand was unable to distinguish a clear energy funnel (P_Near_ = 0.26; **Table 3**).

**Figure 2.**
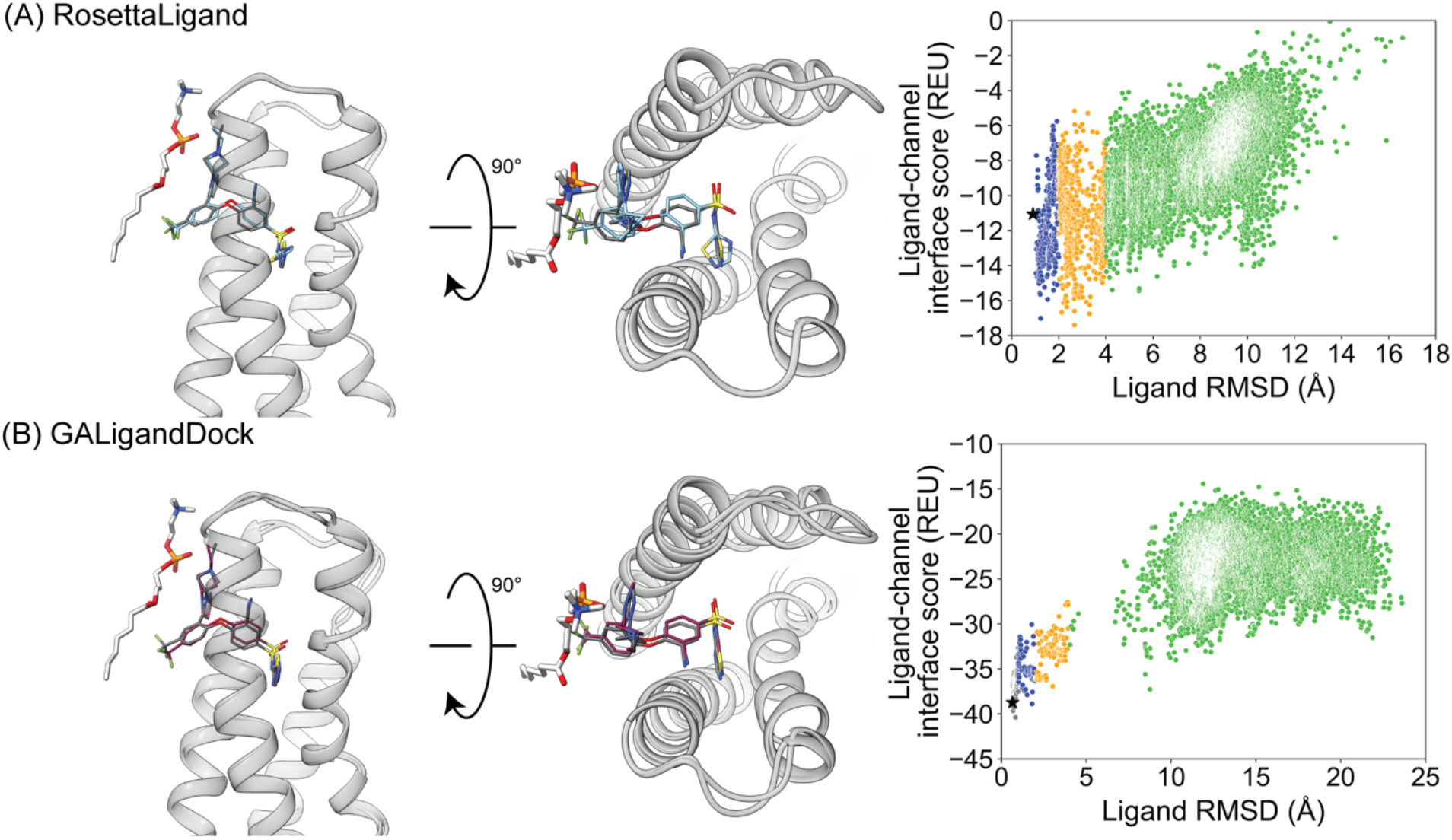
GX-936 docking to hNa_V_1.7-Na_V_Ab voltage sensing domain 4 (PDB 5EK0). **(A)** RosettaLigand (RMSD_Min_ = 0.91 Å, P_Near_ = 0.26) and **(B)** GALigandDock (RMSD_Min_ = 0.65 Å, P_Near_ = 0.83). Left: transmembrane view. Middle: extracellular view. Right: ligand RMSD vs interface energy distribution of the top 10% of total energy poses. Grey dots: < 1 Å, blue dots: 1-2 Å, yellow dots: 2-4 Å, green dots: > 4 Å. Carbon atoms of RosettaLigand molecule are shown in cyan, GALigandDock in dark pink, native structure coordinates in dark grey and phospholipid head group resolved from the structure in light grey. Non-carbon atoms match the Corey-Pauling-Koltun coloring scheme. Black star indicates RMSD_Min_ pose from the entire population.

### Ligand Docking into the Pore-Forming Domain

Tetrodotoxin (TTX) and saxitoxin (STX) are both guanidinium-based small molecules derived from puffer fish and shellfish, and act as selective blockers of sodium channels by binding to the pore-forming domain (Hille, 2018; Shen et al., 2019). Being potent pore blockers for only some Na_V_ channel isoforms, hNa_V_ channel isoforms are classified in physiology as TTX-resistant (hNa_V_1.5, hNa_V_1.8-1.9) and TTX-sensitive (hNa_V_1.1-1.4, hNa_V_1.6-1.7) (Stevens, Peigneur, and Tytgat, 2011). Like aryl sulfonamides for VSDs, the discovery of TTX and STX binding to the pore-forming domain of Na_V_ channels has spurred the design of similar blockers with greater selectivity for a specific hNa_V_ isoform, usually hNa_V_1.7 for pain therapy (Hagen et al., 2017; Pajouhesh et al., 2020).

We chose STX and TTX as test cases to evaluate ligand docking into the pore-forming domain of hNa_V_1.7 (PDB: 6J8G and 6J8I, respectively; Shen et al., 2019). The protonation and bond order of STX and TTX from a previously reported work was used (Thomas-Tran and Du Bois, 2016). Based on their cage-like structures, STX has only 3 rotatable bonds and TTX has only 1 rotatable bond: making them the most rigid ligands in this study (**Table 1**; **Figure 1**). In both cases, RosettaLigand docking resulted in RMSD_Min_ values of 0.70 Å for STX and 0.54 Å for TTX, while GALigandDock reported RMSD_Min_ values of 0.76 Å for STX and 0.81 Å for TTX **(Table 2**; **Figures 3 and 4)**. The P_Near_ values suggested little to no energy funnel convergence with STX (RosettaLigand P_Near_ = 9.3•10^-3^ and GALigandDock P_Near_ = 1.6•10^-2^), and TTX (RosettaLigand P_Near_ = 7.8•10^-2^ and GALigandDock P_Near_ = 0.27) (**Table 3**). This lack of convergence is due to other energy minima occurring within 3-6 Å RMSD (**Figures 3 and 4**).

**Figure 3.**
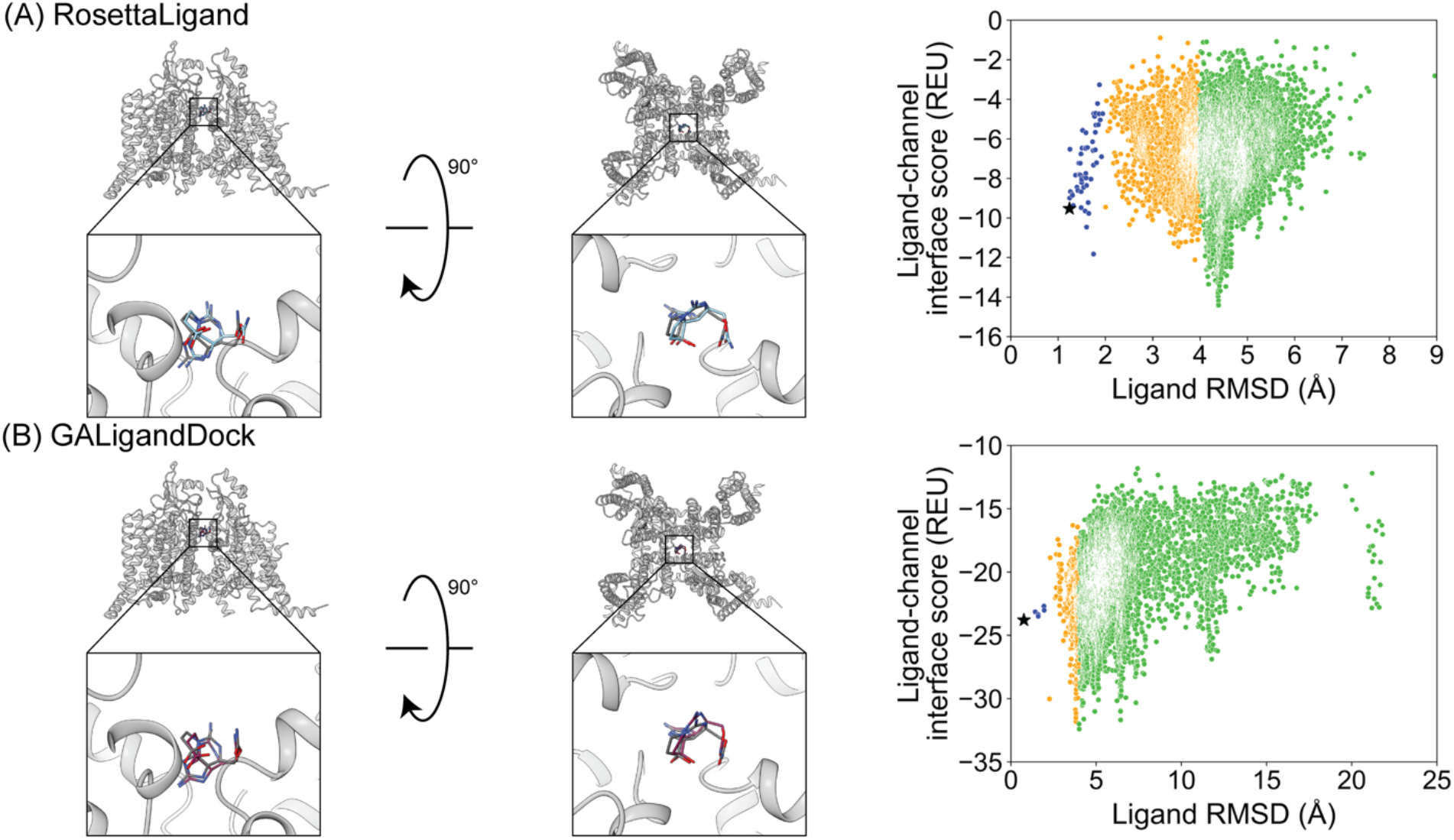
Saxitoxin docking to hNa_V_1.7 selectivity filter (PDB 6J8G). **(A)** RosettaLigand (RMSD_Min_ = 0.70 Å, P_Near_ = 9.3•10^-3^) and **(B)** GALigandDock (RMSD_Min_ = 0.76 Å, P_Near_ = 1.6•10^-2^). Left: transmembrane view. Middle: extracellular view. Right: ligand RMSD vs interface energy distribution of the top 10% of total energy poses. Grey dots: < 1 Å, blue dots: 1-2 Å, yellow dots: 2-4 Å, green dots: > 4 Å. Carbon atoms of RosettaLigand molecule are shown in cyan, GALigandDock in dark pink, and native structure coordinates in dark grey. Non-carbon atoms match the Corey-Pauling-Koltun coloring scheme. Black star indicates RMSD_Min_ pose from the entire population.

**Figure 4.**
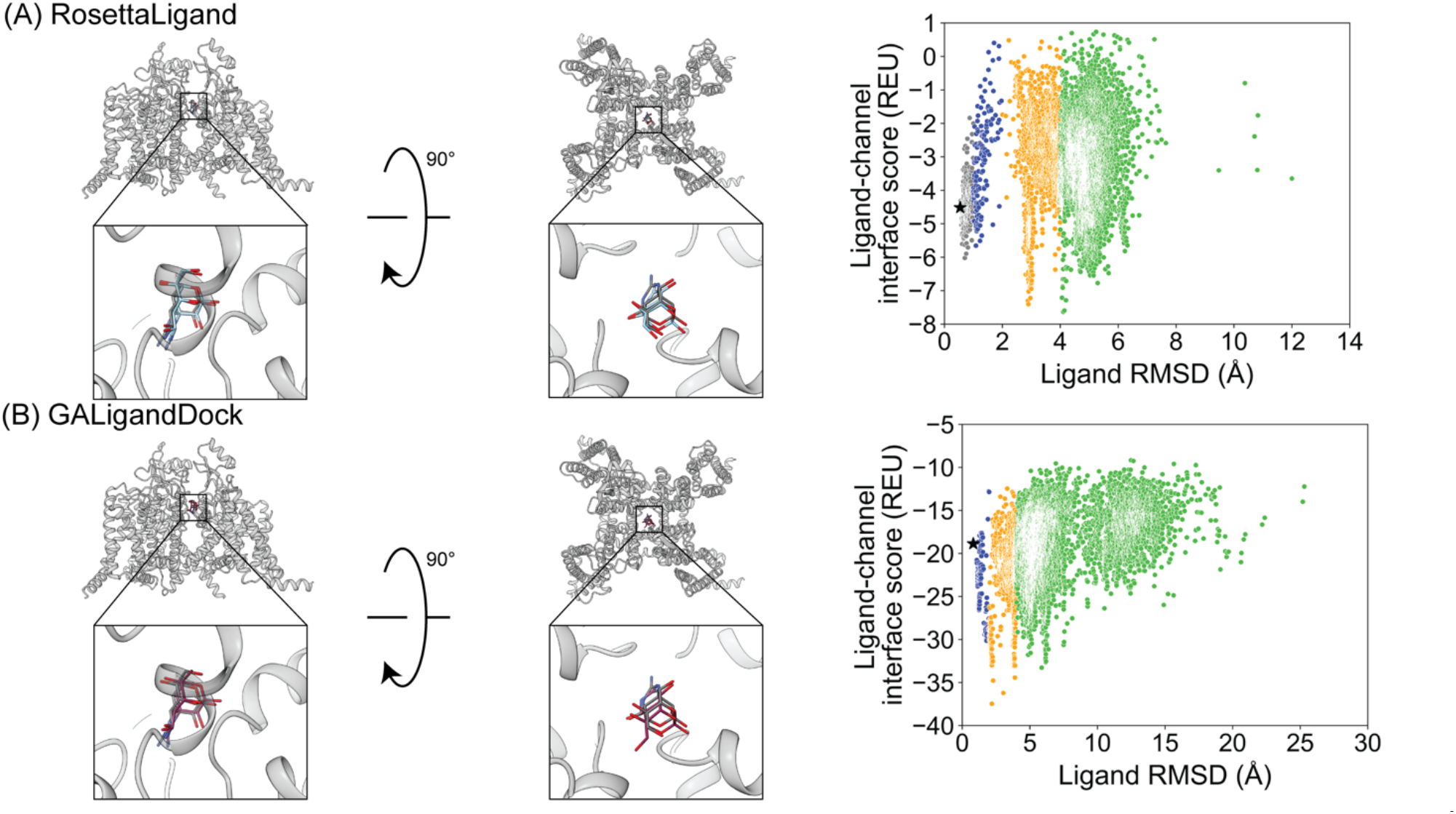
Tetrodotoxin docking to hNa_V_1.7 selectivity filter (PDB 6J8I). **(A)** (RMSD_Min_ = 0.54 Å, P_Near_ = 7.8•10^-2^) and **(B)** GALigandDock (RMSD_Min_ = 0.81 Å, P_Near_ = 0.27). Left: transmembrane view. Middle: extracellular view. Right: ligand RMSD vs interface energy distribution of the top 10% of total energy poses. Grey dots: < 1 Å, blue dots: 1-2 Å, yellow dots: 2-4 Å, green dots: > 4 Å. Carbon atoms of RosettaLigand molecule are shown in cyan, GALigandDock in dark pink, and native structure coordinates in dark grey. Non-carbon atoms match the Corey-Pauling-Koltun coloring scheme. Black star indicates RMSD_Min_ pose from the entire population.

For STX, the lowest ten interface energy poses with RosettaLigand converged at 4.4 Å RMSD, with the carbamoyl group positioned deeper into the selectivity filter, while GALigandDock rendered a 3.8-6.4 Å RMSD, with the center of STX further away from the selectivity filter in various rotations (**Figure S14.1-2)**. For TTX, the lowest ten interface energy poses with RosettaLigand contained poses between 2.9-4.2 Å RMSD; five of ten poses converged to the structural coordinates with the amino group pointing away from the selectivity filter, while the other five poses converged to the structural coordinates with the amino group pointing perpendicular to the selectivity filter path, (**Figure S15.1**). With GALigandDock the poses were between 2.2-5.9 Å RMSD, with five of ten poses having the TTX amino group pointing towards the pore and one set of P1/P2 helices, three of ten poses with the TTX amino group pointing toward the selectivity filter path, and one pose with the TTX amino group pointing toward the extracellular space. (**Figure S15.2**).

### Ligand docking into the central pore cavity

Seven test cases were used to evaluate ligand docking into the central pore cavity involving the following channels: four cases from rabbit Ca_V_1.1, one from human Ca_V_3.1, one from rat Na_V_1.5, and one from TRPV2. The central pore cavity is a broad target, with small molecules reported to traverse through the gate or fenestration sites to reach their binding site and modulate channel activity via pore blockade or allosteric mechanism (Hille, 1977; Hockerman et al., 1997; Jiang et al., 2020). Drugs targeting this region can act as vasodilators (dihydropyridines), antiarrhythmics (benzothiazepines, phenylalkylamines, flecainide), antiepileptics (Z944, cannabidiol), or local anesthetics (flecainide) (Pumroy et al., 2019; Zhao et al., 2019a; Zhao et al., 2019b; Jiang et al., 2020).

We docked nifedipine (dihydropyridine channel blocker), Bay K 8644 (dihydropyridine channel activator), diltiazem (benzothiazepine channel blocker), and two conformations of verapamil (phenylalkylamine channel blocker) into rabbit Ca_V_1.1. The number of rotatable bonds was five, three, seven, and thirteen, respectively (**Table 1**; **Figure 1**).

Nifedipine docking resulted in a RosettaLigand RMSD_Min_ of 0.77 Å and a GALigandDock RMSD_Min_ of 1.3 Å (**Figure 5**, **Table 2**). The calculated P_Near_ suggested little energy funnel convergence (RosettaLigand P_Near_ = 0.28 and GALigandDock P_Near_ = 0.29) (**Table 3**). This lack of energy convergence is exemplified by the interaction energy distributions containing multiple low energy minima 1-2 Å and 4-6 Å RMSD away from the ligand’s structural coordinates (**Figure 5**). The lowest ten interface energy poses with RosettaLigand contained poses between 1.1-4.4 Å RMSD, with most poses being 1.7 Å or less; eight of ten poses converged to the structural coordinates with slight variation in rotamers, while two of the ten poses flipped the position of the dihydropyridine and carbon ring relative to the structural coordinates (**Figure S16.1**). With GALigandDock, the lowest ten interface energy poses were at 1.3 Å RMSD, in a similar position and orientation to the low RMSD conformations from RosettaLigand (**Figure S16.2)**.

**Figure 5.**
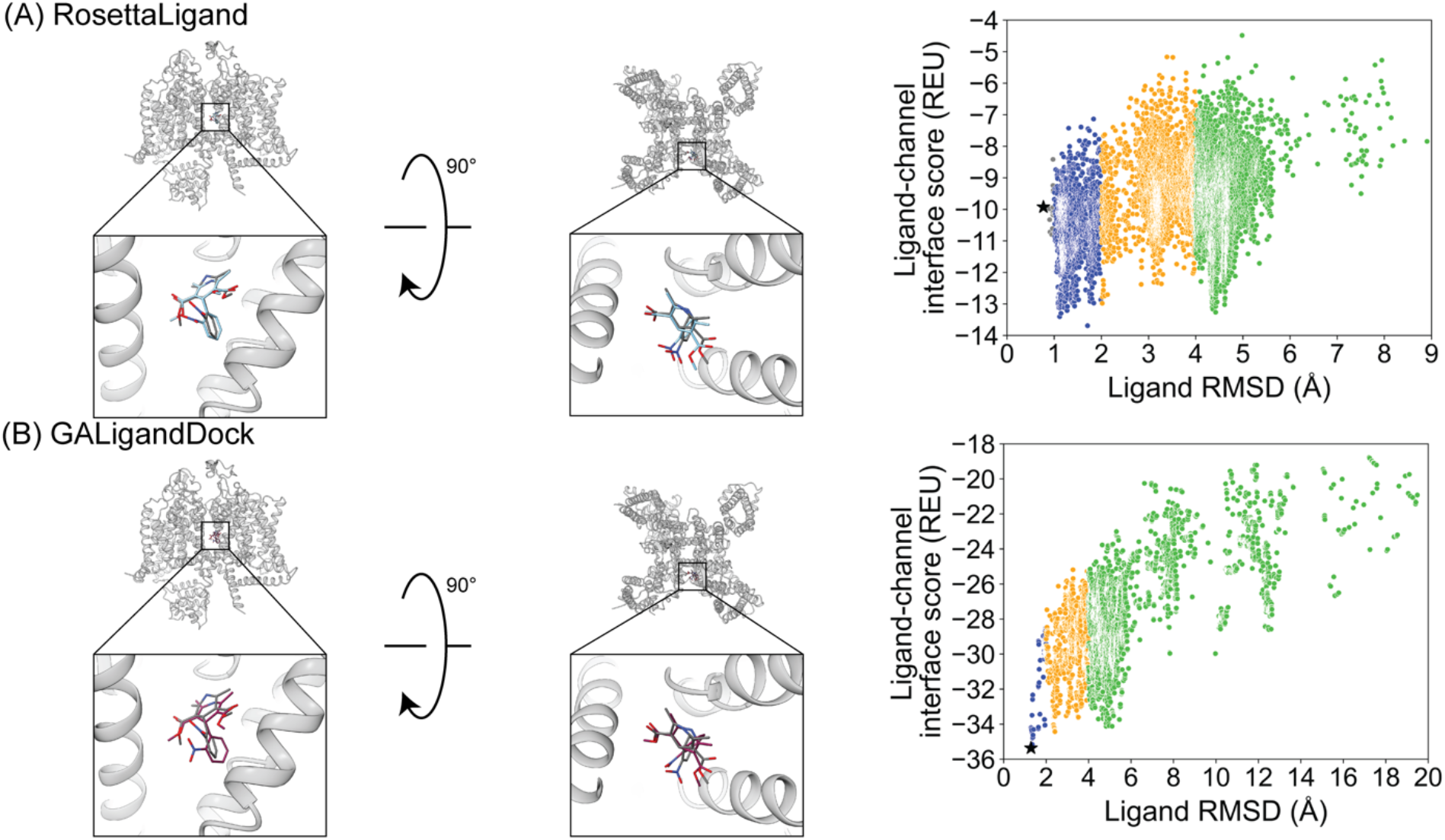
Nifedipine docking to rabbit Ca_V_1.1 central pore cavity (PDB 6JP5). **(A)** RosettaLigand (RMSD_Min_ = 0.77 Å, P_Near_ = 0.28) and **(B)** GALigandDock (RMSD_Min_ = 1.3 Å, P_Near_ = 0.29). Left: transmembrane view. Middle: extracellular view. Right: ligand RMSD vs Interface energy distribution of the top 10% of total energy poses. Grey dots: < 1 Å, blue dots: 1-2 Å, yellow dots: 2-4 Å, green dots: > 4 Å. Carbon atoms of RosettaLigand molecule are shown in cyan, GALigandDock in dark pink, and native structure coordinates in dark grey. Non-carbon atoms match the Corey-Pauling-Koltun coloring scheme. Black star indicates RMSD_Min_ pose from the entire population.

Bay K 8644 docking resulted in a RosettaLigand RMSD_Min_ of 0.93 Å and GALigandDock RMSD_Min_ of 0.97 Å (**Figure 6**, **Table 2**). The calculated RosettaLigand P_Near_ was 0.66 and the GALigandDock P_Near_ was 0.61 (**Table 3**). The P_Near_ values, paired with the interaction energy distribution data, indicate well-converged energy funnels. Further, the lowest interaction energy poses with RosettaLigand was within 0.3 Å of the RMSD_Min_ pose (**Table S2.1**) and for GALigandDock was within 0.2 Å of the RMSD_Min_ pose (**Table S2.5**). With RosettaLigand, the lowest ten interface energy poses were between 1.2-1.3 Å RMSD, with all ten poses converged to the structural coordinates with slight variation in rotamers (**Figure S17.1**). With GALigandDock, the lowest ten interface energy poses were at 1.1 Å or 4.6 Å RMSD. Eight of ten poses had 1.1 Å RMSD with slight deviation in position to structural coordinates, while two of ten poses had 4.6 Å RMSD with the dihydropyridine in the correct position, but the carbon ring flipped 180 degrees such that the trifluoromethyl group was pointed in the opposite direction (**Figure S17.2)**.

**Figure 6.**
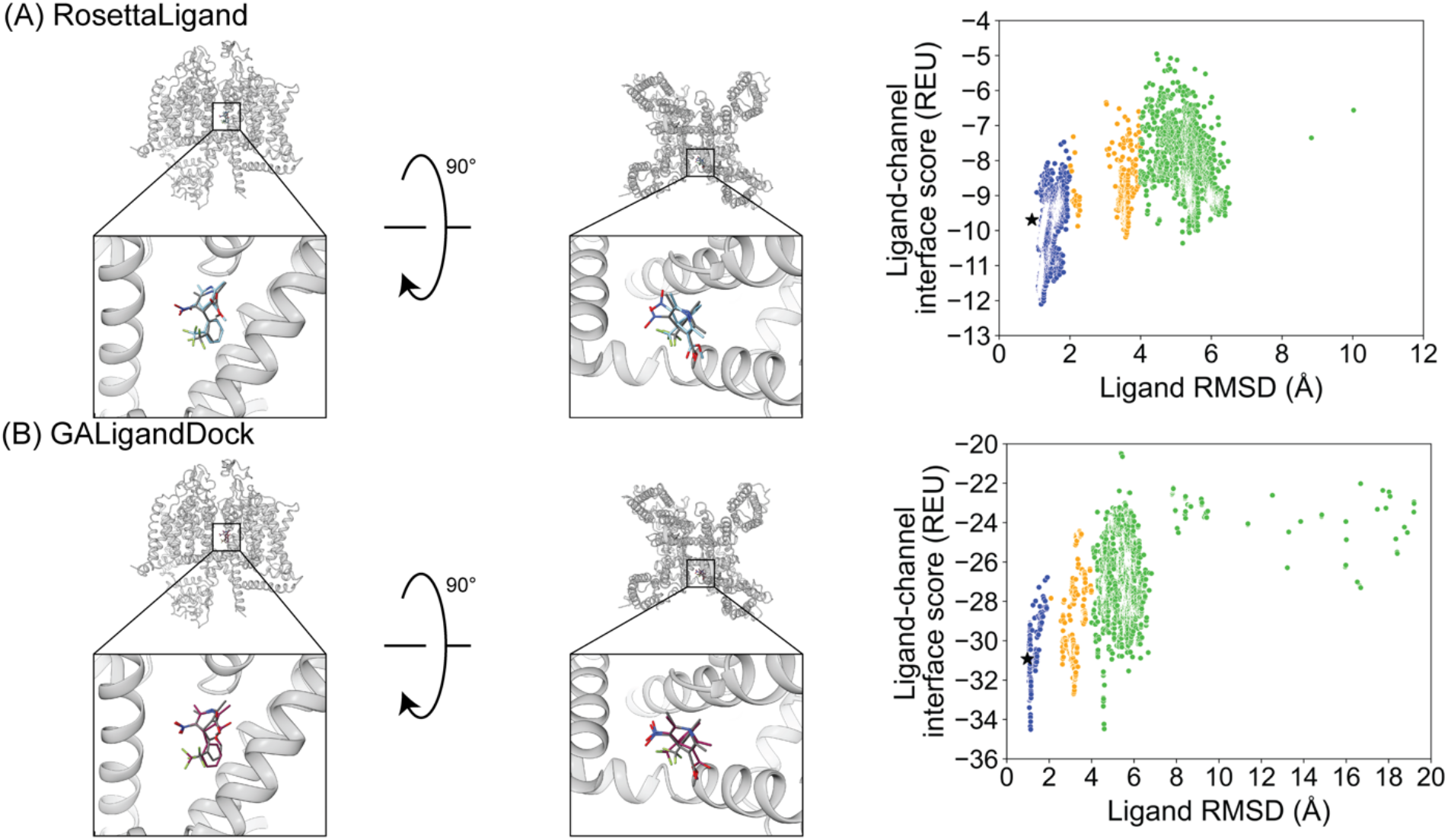
Bay K 8644 docking to rabbit Ca_V_1.1 central pore cavity (PDB 6JP8). **(A)** RosettaLigand (RMSD_Min_ = 0.93 Å, P_Near_ = 0.66) and **(B)** GALigandDock (RMSD_Min_ = 0.97 Å, P_Near_ = 0.61). Left: transmembrane view. Middle: extracellular view. Right: ligand RMSD vs interface energy distribution of the top 10% of total energy poses. Grey dots: < 1 Å, blue dots: 1-2 Å, yellow dots: 2-4 Å, green dots: > 4 Å. Carbon atoms of RosettaLigand molecule are shown in cyan, GALigandDock in dark pink, and native structure coordinates in dark grey. Non-carbon atoms match the Corey-Pauling-Koltun coloring scheme. Black star indicates RMSD_Min_ pose from the entire population.

Diltiazem docking resulted in a RosettaLigand RMSD_Min_ of 2.5 Å and a GALigandDock RMSD_Min_ of 1.1 Å (**Figure 7**, **Table 2**). The calculated P_Near_ suggested no energy funnel convergence for either method (RosettaLigand P_Near_ = 6.4•10^-4^ and GALigandDock P_Near_= 7.8•10^-6^) (**Table 3**). This lack of energy funnel convergence is exemplified by the interaction energy distributions containing local minima 5-14 Å RMSD away from the ligand’s structural coordinates (**Figure 7**). With RosettaLigand, the lowest ten interface energy poses were between 5.2-7.6 Å RMSD, with all ten poses at a similar channel depth at the pore center, but with no convergence in local position or rotamer placement. (**Figure S20.1**). With GALigandDock, the lowest ten interface energy poses were between 10.5-14.5 Å RMSD. Rather than being positioned in the pore center, all ten poses were in a similar channel depth to the structural coordinates but positioned in the fenestration with no convergence in local position or rotamer placement (**Figure S20.2**).

**Figure 7.**
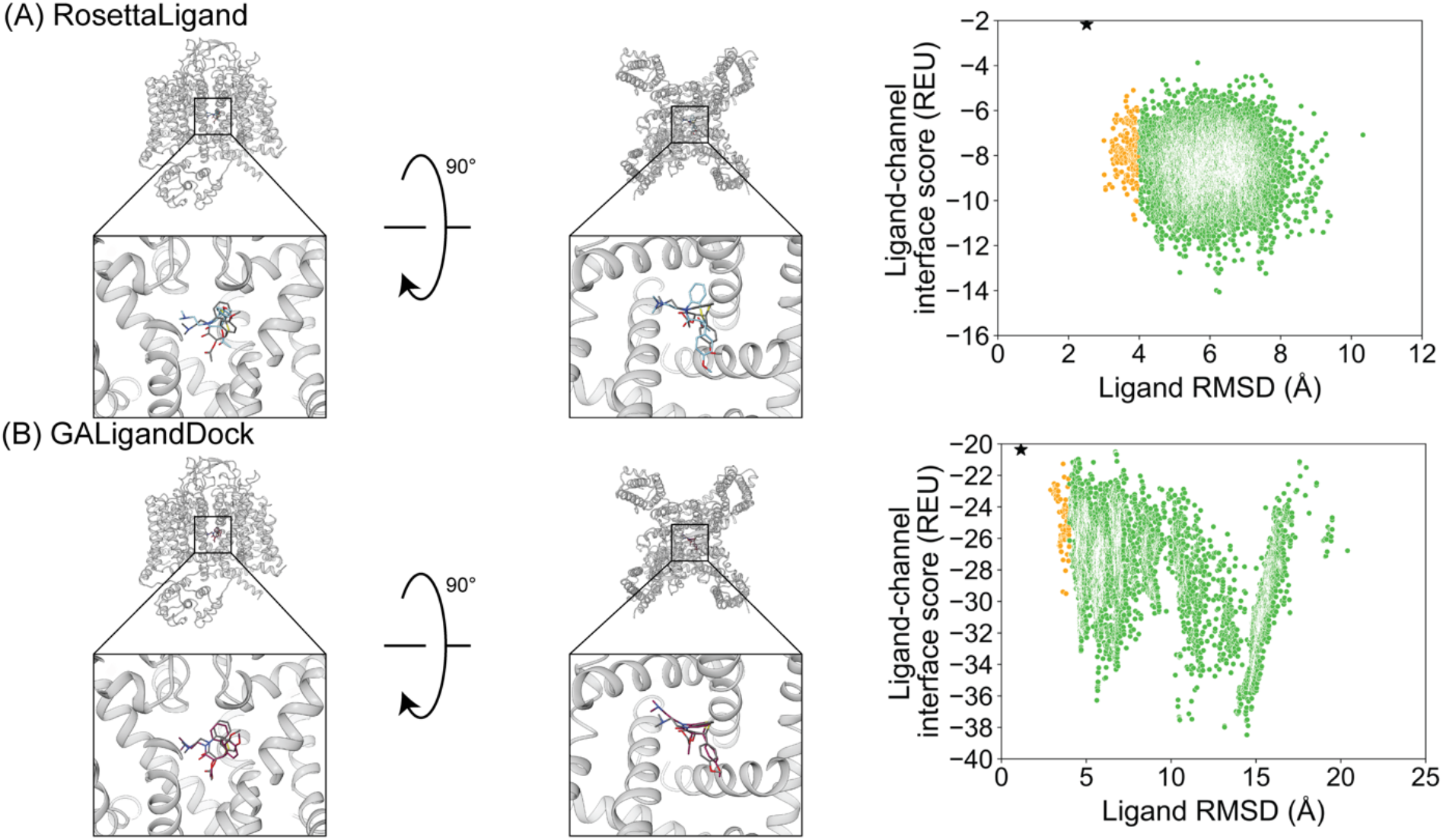
Diltiazem docking to rabbit Ca_V_1.1 central pore cavity (PDB 6JPB). **(A)** RosettaLigand (RMSD_Min_ = 2.5 Å, P_Near_ = 6.4•10^-4^) and **(B)** GALigandDock (RMSD_Min_ = 1.1 Å, P_Near_ = 7.8•10^-6^). Left: transmembrane view. Middle: extracellular view. Right: ligand RMSD vs interface energy distribution of the top 10% of total energy poses. Yellow dots: 2-4 Å, green dots: > 4 Å. Carbon atoms of RosettaLigand molecule are shown in cyan, GALigandDock in dark pink, and native structure coordinates in dark grey. Non-carbon atoms match the Corey-Pauling-Koltun coloring scheme. Black star indicates RMSD_Min_ pose from the entire population.

Verapamil was previously resolved in a complex with rabbit Ca_V_1.1 in two binding modes with flipped orientations (Zhao et al., 2019a). We evaluated if RosettaLigand and GALigandDock sampling favored a particular orientation. For the first orientation, docking resulted in a RosettaLigand RMSD_Min_ of 1.4 Å and a GALigandDock RMSD_Min_ of 2.0 Å (**Figure 8**, **Table 2**). The calculated P_Near_ suggested no energy funnel convergence (RosettaLigand P_Near_ = 4.3•10^-3^, GALigandDock P_Near_ = 2.8•10^-3^) (**Table 3**). This nonexistent energy funnel is evident by local interaction energy minima beyond 4 Å RMSD from the ligand structure coordinates (**Figure 8)**. With RosettaLigand, the lowest ten interface energy poses were between 4.1-9.7 Å RMSD, with all ten poses at a similar channel depth, and some poses converging in local position and rotamer placement at 5.7 Å and 9.2 Å RMSD (**Figure S18.1**). With GALigandDock, the lowest ten interface energy poses were between 3.8-11.9 Å RMSD. All ten poses were at different channel depths, positioned in the pore center or at the pore center – fenestration interface, and did not converge in local position or rotamer placement (**Figure S18.2**).

**Figure 8.**
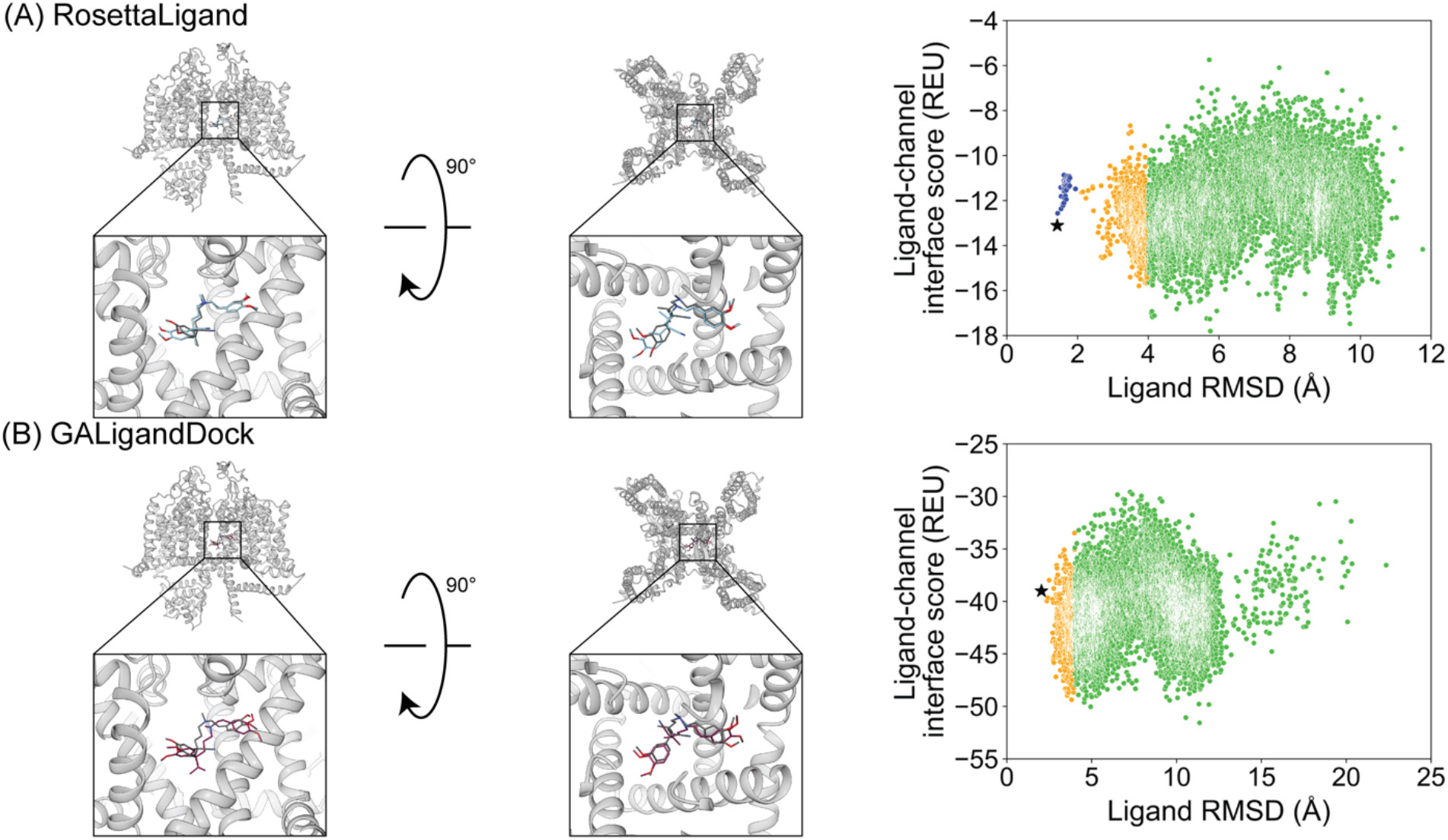
Verapamil docking in its first orientation to rabbit Ca_V_1.1 central pore cavity (PDB 6JPA). **(A)** RosettaLigand (RMSD_Min_ = 1.4 Å, P_Near_ = 4.3•10^-3^) and **(B)** GALigandDock (RMSD_Min_ = 2.0 Å, P_Near_ = 2.8•10^-3^). Left: transmembrane view. Middle: extracellular view. Right: ligand RMSD vs interface energy distribution of the top 10% of total energy poses. Blue dots: 1-2 Å, yellow dots: 2-4 Å, green dots: > 4 Å. Carbon atoms of RosettaLigand molecule are shown in cyan, GALigandDock in dark pink, and native structure coordinates in dark grey. Non-carbon atoms match the Corey-Pauling-Koltun coloring scheme. Black star indicates RMSD_Min_ pose from the entire population.

For the second orientation, docking resulted in a RosettaLigand RMSD_Min_ of 2.2 Å and a GALigandDock RMSD_Min_ of 2.1 Å (**Figure 9**, **Table 2**). The calculated P_Near_ values suggest no energy funnel convergence (RosettaLigand P_Near_ = 1.2•10^-4^, GALigandDock P_Near_ = 1.9•10^-5^) (**Table 3**). Similar to the first orientation, the interaction energy distribution for the second orientation yields local energy minima greater than 4 Å RMSD from the ligand structure coordinates (**Figure 9**). With RosettaLigand, the lowest ten interface energy poses were between 8.5-10.8 Å RMSD, with all ten poses at a similar channel depth, and some poses converging in local position, similar to the docking set for the first structural orientation of verapamil (**Figure S19.1**). With GALigandDock, the lowest ten interface energy poses were between 6.5-10.8 Å RMSD. All ten poses were at different channel depths, positioned in the pore center or at the pore center – fenestration interface, and did not converge in local position or rotamer placement, similar to the docking set for the first structural orientation of verapamil (**Figure S19.2**).

**Figure 9.**
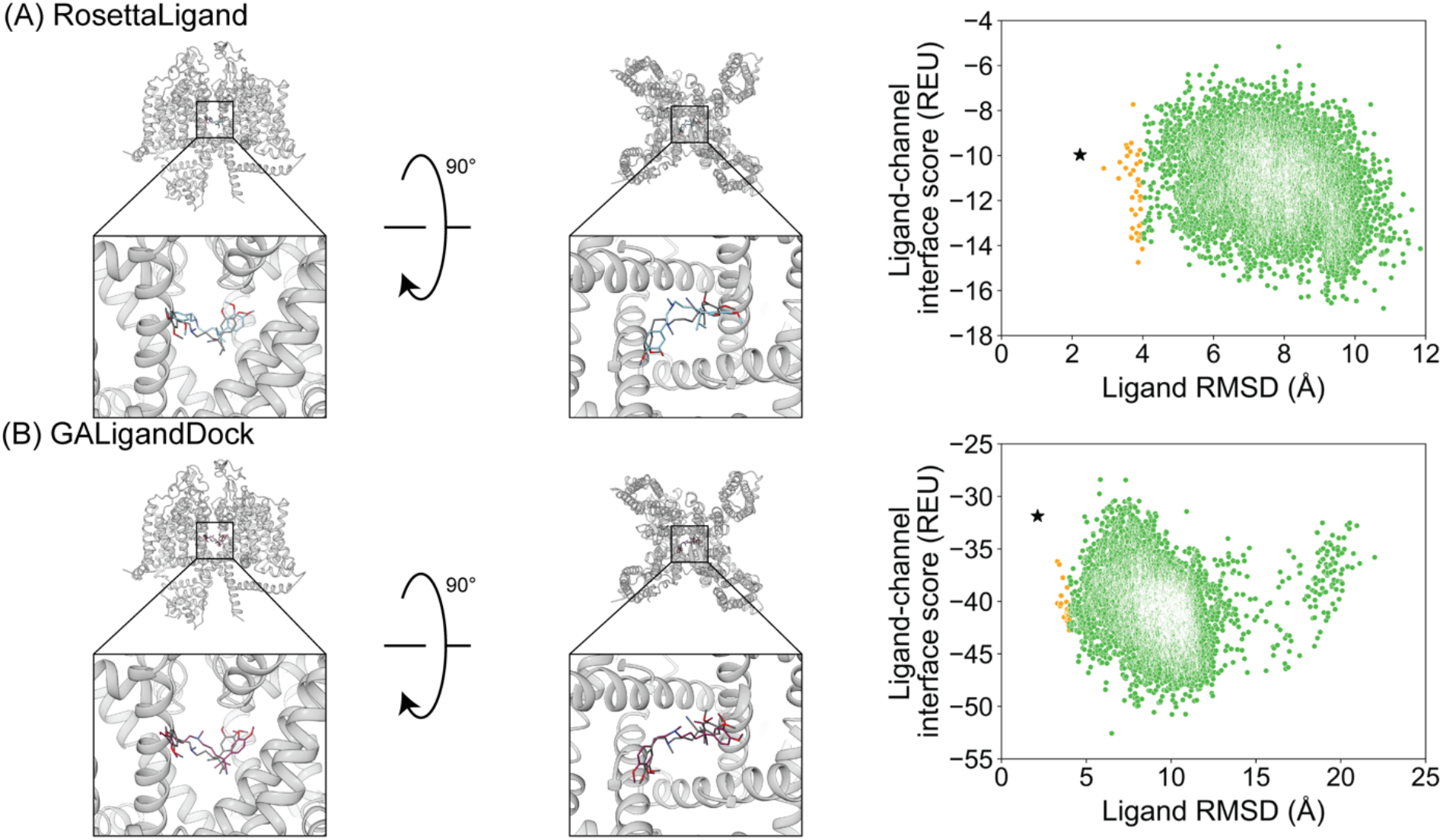
Verapamil docking in its second orientation to rabbit Ca_V_1.1 central pore cavity (PDB 6JPA). **(A)** RosettaLigand (RMSD_Min_ = 2.2 Å, P_Near_ = 1.2•10^-4^) and **(B)** GALigandDock (RMSD_Min_ = 2.1 Å, P_Near_ = 1.9•10^-5^). Left: transmembrane view. Middle: extracellular view. Right: ligand RMSD vs interface energy distribution of the top 10% of total energy poses. Yellow dots: 2-4 Å, green dots: > 4 Å. Carbon atoms of RosettaLigand molecule are shown in cyan, GALigandDock in dark pink, and native structure coordinates in dark grey. Non-carbon atoms match the Corey-Pauling-Koltun coloring scheme. Black star indicates RMSD_Min_ pose from the entire population.

The small molecule Z944 (channel blocker) has 6 rotatable bonds and is resolved bound to human Ca_V_3.1. Docking with RosettaLigand and GALigandDock both resulted in a RMSD_Min_ of 0.73 Å and 0.84 Å, respectively (**Figure 10**, **Table 2**). Interestingly, the calculated P_Near_ suggests good energy funnel convergence for RosettaLigand (P_Near_ = 0.62), but no energy funnel convergence for GALigandDock (P_Near_ = 1.1•10^-4^) (**Table 3**). For RosettaLigand, the interaction energy produced an energy funnel between 1-2 Å, while for GALigandDock produced an energy funnel near 10-15 Å (**Figure 10**). With RosettaLigand, the lowest ten interface energy poses were between 0.90-1.7 Å RMSD, with all ten poses converged to the structural coordinates with slight variation in rotamers. (**Figure S21.1**). With GALigandDock, the lowest ten interface energy poses were between 10.6-12.4 Å RMSD, with four sets of unique poses identified. The first set, containing three of the ten poses, is in a slightly lower channel depth and in a similar orientation to the structural coordinates, but rotated approximately 180 degrees about the pore center. The second set is one pose and is in a similar location as the first set but is rotated such that the phenyl ring and tertiary butylamine positions are flipped. The third group, containing four poses, is similar to the second set but rotated approximately 180 degrees about the pore center. The last group, containing two poses, is in a similar channel depth and location to the structural coordinates, but the phenyl ring and the tertiary butylamine positions are flipped (**Figure S21.2**).

**Figure 10.**
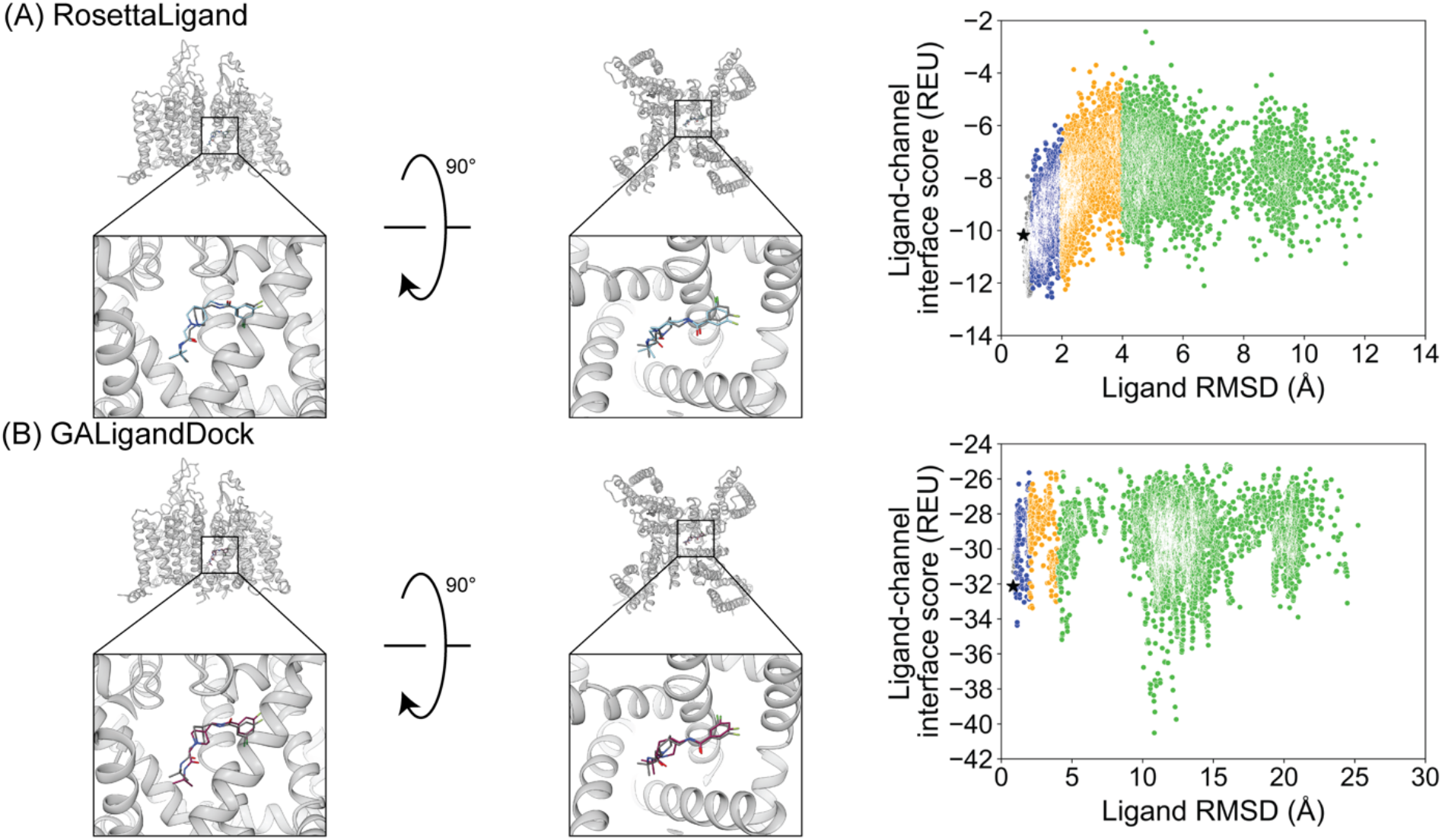
Z944 docking to human Ca_V_3.1 central pore cavity (6KZP). **(A)** RosettaLigand (RMSD_Min_ = 0.73 Å, P_Near_ = 0.62) and **(B)** GALigandDock (RMSD_Min_ = 0.84 Å, P_Near_ = 1.1•10^-4^). Left: transmembrane view. Middle: extracellular view. Right: ligand RMSD vs interface energy distribution of the top 10% of total energy poses. Grey dots: < 1 Å, blue dots: 1-2 Å, yellow dots: 2-4 Å, green dots: > 4 Å. Carbon atoms of RosettaLigand molecule are shown in cyan, GALigandDock in dark pink, and native structure coordinates in dark grey. Non-carbon atoms match the Corey-Pauling-Koltun coloring scheme. Black star indicates RMSD_Min_ pose from the entire population.

Flecainide (channel blocker) has 7 rotatable bonds and is resolved bound to rat Na_V_1.5. RosettaLigand docking resulted in a RMSD_Min_ of 1.2 Å, while GALigandDock resulted in a RMSD_Min_ of 0.94 Å (**Figure 11**, **Table 2**). The calculated P_Near_ values suggest no energy funnel convergence with either method (RosettaLigand P_Near_ = 0.12, GALigandDock P_Near_ = 2.9•10^-5^) (**Table 3**). The interaction energy distribution from RosettaLigand produced an energy minimum between 2-4 Å and 7-10 Å, while from GALigandDock produced energy minima near 10-12 Å (**Figure 11**). With RosettaLigand, the lowest ten interface energy poses were between 2.1-4.0 Å RMSD, with all ten poses in the same channel depth as the structural coordinates, but with a tilt such that the trifluoromethyl groups are level in channel depth, rather than the trifluoromethyl group facing the pore being lower in depth. In one pose, the piperidin-2-yl-methylamine was positioned lower into the channel than the rest of the ligand and the structural coordinates, while the rest of the ligand pose was in a similar position to other poses (**Figure S23.1**). With GALigandDock, the lowest ten interface energy poses were between 9.7-11.9 Å RMSD, with all ten poses positioned at the fenestration in the same channel depth as the structural coordinates, but oriented outside of the pore with one of the trifluoromethyl groups pointing towards the pore center but positioned at the exterior of the channel fenestration (**Figure S23.2**).

**Figure 11.**
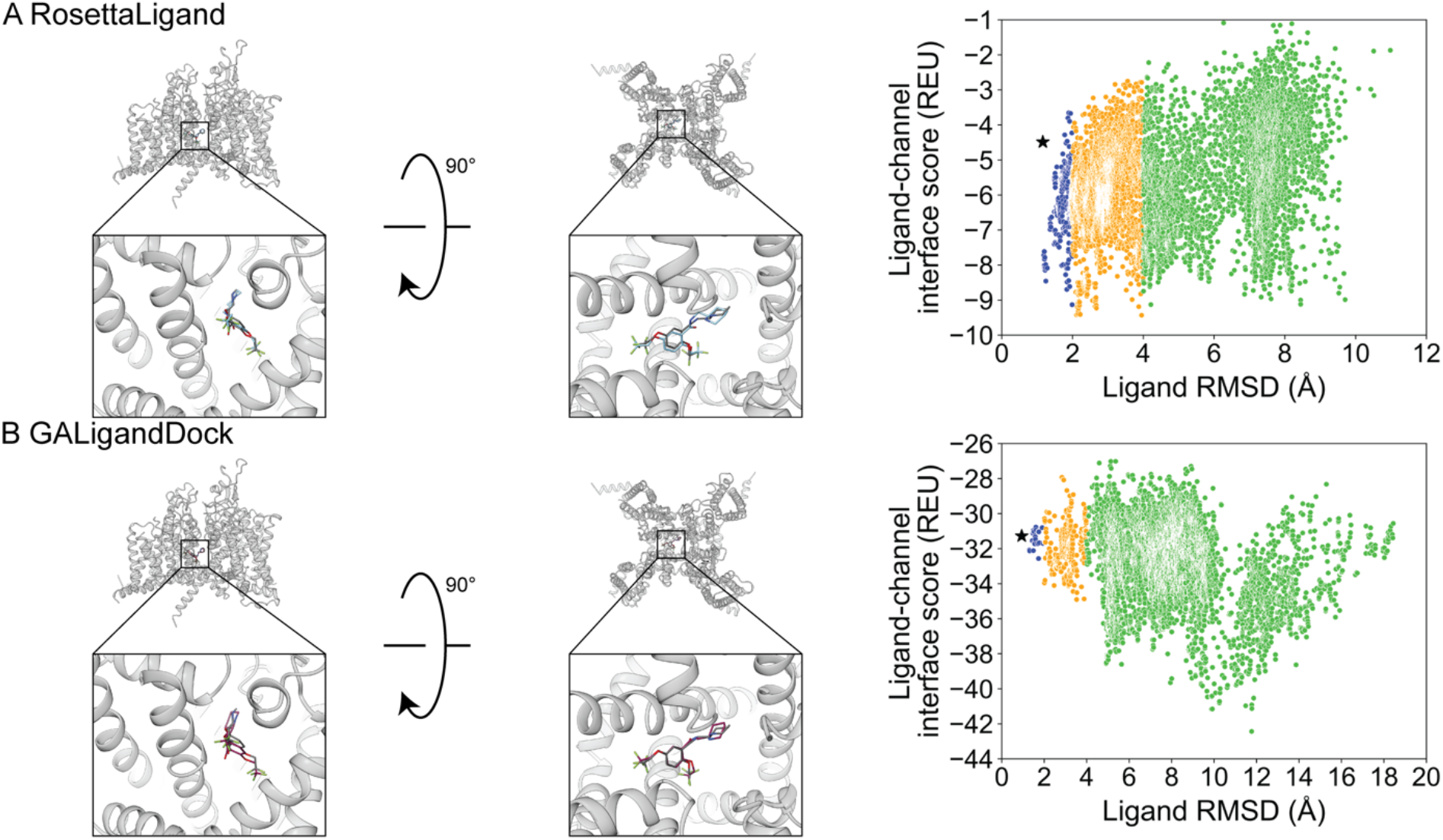
Flecainide docking to rat Na_V_1.5 central pore cavity (PDB 6UZ0). **(A)** RosettaLigand (RMSD_Min_ = 1.2 Å, P_Near_ = 0.12) and **(B)** GALigandDock (RMSD_Min_ = 0.94 Å, P_Near_ = 2.9•10^-5^). Left: transmembrane view. Middle: extracellular view. Right: ligand RMSD vs interface energy distribution of the top 10% of total energy poses. Blue dots: 1-2 Å, yellow dots: 2-4 Å, green dots: > 4 Å. Carbon atoms of RosettaLigand molecule is indicated in cyan, GALigandDock in dark pink, and native structure coordinates in dark grey. Non-carbon atoms match the Corey-Pauling-Koltun coloring scheme. Black star indicates RMSD_Min_ pose from the entire population.

The transient receptor potential vanilloid (TRPV) subfamily of cation channels broadly play roles in thermo-sensation and thermoregulation in response to heat (>53 °C), notably in noxious heat sensing (Nilius, 2007; Gees et al., 2012). TRPV2 channels are widely expressed, with identification in dorsal root ganglion neurons, as well as brain, heart, and smooth muscle tissue, among others (Gees et al., 2012). The TRPV2 channel has been implicated in thermal pain sensing, muscular dystrophy, and cardiomyopathy, among other diseases (Nilius, 2007; Gees et al., 2012). Structurally, TRPV2 channels contain six transmembrane spanning domains and commonly assemble as a homomer with four identical subunits (Zheng and Trudeau, 2015). They resemble canonical voltage-gated ion channels with a pore domain and a weak voltage sensor domain, while their gating is regulated by heat and a diverse set of agonists such as 2-aminoethoxydiphenyl borate and cannabidiol (Gees et al., 2012; Zheng and Trudeau, 2015; Pumroy et al., 2019).

Cannabidiol (channel activator) has 6 rotatable bonds and is resolved bound to rat TRPV2. Docking with RosettaLigand resulted in a RMSD_Min_ of 1.0 Å, while GALigandDock resulted in a RMSD_Min_ of 0.90 Å (**Figure 12**, **Table 2**). The calculated P_Near_ values suggest no energy funnel convergence for both methods (RosettaLigand P_Near_ = 4.3•10^-2^, GALigandDock P_Near_ = 3.8•10^-3^) (**Table 3**). For RosettaLigand, the interaction energy distribution did not demonstrate a clear minimum, with minima seen between 2-8 Å (**Figure 12**). For GALigandDock, there is a clear minimum between 6-7 Å (**Figure 12**). With RosettaLigand, the lowest ten interface energy poses were between 2.8-7.0 Å RMSD, with two groups of conformations in the same channel depth and interfaced with the S5 and S6 helices of adjacent TRPV2 monomers like the structural coordinates. In one group, cannabidiol is positioned parallel to the S6 helical segment, with the cyclohexene at the lowest channel depth. In the second group, the poses are positioned in the same orientation as the structural coordinates, with the pentyl chain facing away from the pore center, but the overall position of cannabidiol is slightly elevated relative to the structural coordinates (**Figure S22.1**). With GALigandDock, the lowest ten interface energy poses were either 6.4 Å or 6.7 Å RMSD, with all ten poses positioned at the fenestration in the same channel depth and interfacing with the S5 and S6 helices of adjacent TRPV2 monomers like the structural coordinates, but with the ligand rotated around the cyclohexene plane such that the pentyl chain is oriented parallel to the S6 helix, rather than pointing outward (**Figure S22.2).**

**Figure 12.**
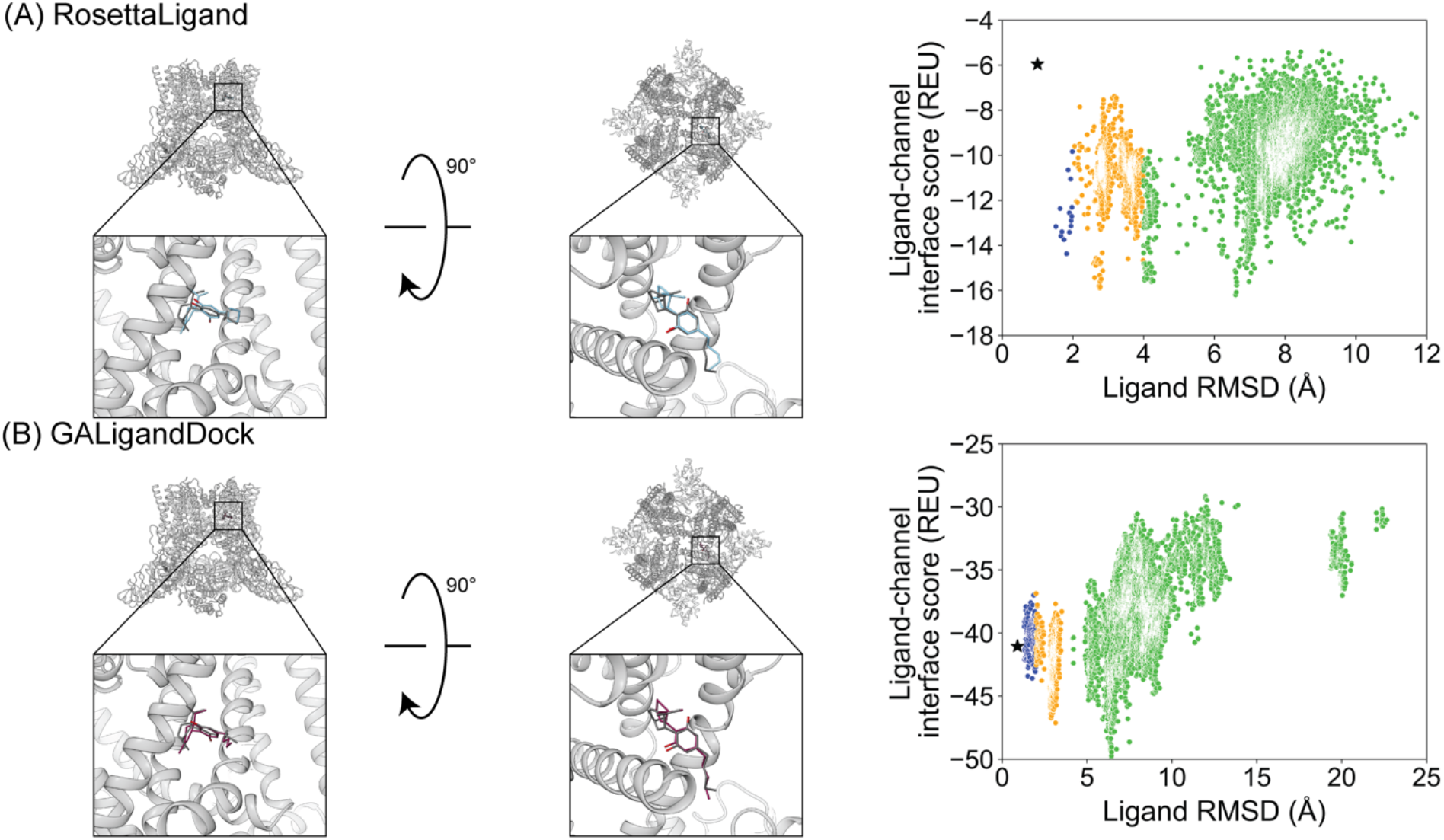
Cannabidiol docking to rat TRPV2 central pore cavity (PDB 6U88). **(A)** RosettaLigand (RMSD_Min_ = 1.0 Å, P_Near_ = 4.3•10^-2^) and **(B)** GALigandDock (RMSD_Min_ = 0.90 Å, P_Near_ = 3.8•10^-3^). Left: transmembrane view. Middle: extracellular view. Right: ligand RMSD vs interface energy distribution of the top 10% of total energy poses. Blue dots: 1-2 Å, yellow dots: 2-4 Å, green dots: > 4 Å. Carbon atoms of RosettaLigand molecule are shown in cyan, GALigandDock in dark pink, and native structure coordinates in dark grey. Non-carbon atoms match the Corey-Pauling-Koltun coloring scheme. Black star indicates RMSD_Min_ pose from the entire population.

## Discussion

Previous studies have underscored the importance of ligand docking methods for generating ion channel structure-based hypotheses (Yang et al., 2013; Shim et al., 2019). Furthermore, ligand docking methods when combined with high-resolution structures can aid in rational drug design (Wang et al., 2007; Liu et al., 2018; Wulff et al., 2019; Maia et al., 2020). Notably, RosettaLigand has been extensively used to predict the molecular mechanisms of ligand – ion channel interactions (Yang et al., 2013; Yang et al., 2015; Nguyen et al., 2017; Nguyen et al., 2019; Shim et al., 2019; Craig 2^nd^ et al., 2022; Vu et al., 2020; Maly et al., 2022; Pumroy et al., 2022). While GALigandDock has not yet been tested on ion channel structure-function relationships, it utilizes a new generalized energy function tailored for small molecules while sampling ligand conformations with a genetic algorithm, making it an attractive complement to RosettaLigand.

Indeed, this study demonstrates that high-resolution structures paired with RosettaLigand and GALigandDock can be useful tools for formulating structural hypotheses and predicting binding modes for drug discovery. Notably, both Rosetta methods can produce ligand poses near the native ligand structure coordinates. Using a standard 2 Å RMSD_Min_ cutoff (Maia et al., 2020; Park et al., 2021), the RosettaLigand docking set yielded an average RMSD_Min_ of 1.2 ± 0.6 Å, and the GALigandDock docking set yielded an average RMSD_Min_ of 1.1 ± 0.5 Å (**Table 2**). However, the ability to discriminate the RMSD_Min_ pose remains challenging, and highlights important practical considerations when applying Rosetta ligand docking methods to a chosen ion channel target.

When performing Rosetta ligand docking, the features of the ligand conformer library, the size of the receptor site, and prior knowledge of critical functional residues, need to be considered to determine the appropriate amount of pose sampling. For our study, we generated either 20,000 poses or 100,000 poses per docking run. While we did notice a statistically significant difference in RMSD_Min_ between 20,000 and 100,000 total poses, the difference was within a few sub-Angstroms. Furthermore, another RosettaLigand benchmarking study of the CASF-2016 dataset generated 1,000 total poses and was able to sufficiently sample a RMSD_Min_ below 2 Å (Su et al., 2019; Smith and Meiler, 2020). Thus, for ion channel docking, we suggest sampling in the ones to tens of thousands, especially when docking in large receptor sites like the pore-forming domain.

Overall, our docking data was able to achieve a P_Near_ greater than 0.5 for two test cases for each docking method when using the top 10% of total-scoring poses. Both methods achieved P_Near_ greater than 0.5 for Bay K8644 to the rabbit Ca_V_1.1 central pore cavity, while RosettaLigand achieved P_Near_ greater than 0.5 for Z944 to the human Ca_V_3.1 central pore cavity, and GALigandDock achieved P_Near_ greater than 0.5 for GX-936 docking to hNa_V_1.7-Na_V_Ab voltage sensing domain four (**Table 3**). The reasons for a low P_Near_ value in most test cases are possibly due to 1) improper scoring by Rosetta score functions to discriminate native-like poses by energy, 2) multiple favorable ligand binding states in the receptor site that have not been structurally resolved, and/or 3) insufficient pose sampling.

Currently, we suggest that Rosetta score functions are unable to sufficiently score near-native poses accurately in ion channel docking. For each docking run, comparing the RMSD_Min_ pose and the lowest interface energy pose indicates that the RMSD_Min_ pose is under-scored (**Tables S2.1-2.5**). Conversely, a previous RosettaLigand study using the CASF-2016 dataset (containing 285 crystal structures of protein – ligand complexes with an overall resolution < 2.5 Å and an R-factor < 0.25) and 1,000 total poses per docking run was able to frequently achieve P_Near_ values between 0.8 and 1.0 (Su et al., 2019; Smith and Meiler, 2020), suggesting that the Rosetta score functions can be utilized for other docking studies, but should be verified for accuracy with test cases. Furthermore, the CASF-2016 dataset does not contain ion channels, while the sample size per docking run is sufficiently lower in the RosettaLigand docking study using CASF-2016 than those in our study, suggesting that sampling is not the predominant issue, but rather favorable scoring of near-native poses is the primary issue.

For the voltage-sensing domains, GX-936 in complex with the VSD4 of hNa_V_1.7-Na_V_Ab chimera (PDB: 5EK0) was the only small-molecule docked yet that was consistently near or below 1 Å RMSD_Min_ for each docking set regardless of the method (**Tables S1.1-1.2**). This may be due to the VSD4 receptor site being the narrowest binding pocket tested, thereby limiting the number of binding configurations. This suggests that VSDs are generally well-suited for Rosetta ligand docking since the receptor is constrained to allow shape-complementary between ligand and protein while reducing the required sampling compared to pore-forming receptor sites. Notably, the Rosetta ligand docking methods employed did not use any implicit membrane parameters, while GX-936 in a biologically realistic context is partially exposed to a lipid head group (Ahuja et al., 2015). Thus, in this case, docking without an implicit membrane energy function was still able to achieve the experimental structurally resolved position, however, artifactual, low interface energy poses where GX-936 would be overlapping the phospholipid space are present (**Figure S13.1-13.2**). Further, GALigandDock was consistently able to achieve a P_Near_ > 0.5 when using a padding of 2.0 Å, 4.0 Å, or 7.0 Å, suggesting that ligands docked to VSDs could be screened by energy funnel convergence (**Tables S4.5-4.10**). More small molecules structurally resolved and docked to the VSD are needed to validate this observation.

For the pore-forming domain, TTX and STX were docked to hNa_V_1.7, as they are canonical small-molecule channel blockers used when studying the Na_V_ family. TTX and STX had the fewest rotatable bonds in this study, suggesting that little conformer sampling would be needed to achieve a near-native pose. Both TTX and STX docking achieved sub-Angstrom RMSD_Min_, except STX docking with GALigandDock at a padding of 7 Å (RMSD_Min_ =2.3 Å, padding 7 Å/100,000 poses vs RMSD_Min_ = 1.0 Å, padding 4 Å/100,000 poses; **Table S1.2**). However, all docking methods employed were unable to achieve a P_Near_ > 0.5, suggesting that 1) STX and TTX could have alternative binding conformations, 2) the docking methods employed have wrongfully scored alternative low-energy conformations, or 3) that inherent loop flexibility in the pore-forming domain is a requirement for docking an energy-optimized, induced-fit conformation. It should be noted that the hNa_V_1.7 selectivity filter is lined with polar residues that could contribute to a vast hydrogen bonding network, in addition to water hydrating the selectivity filter and the filter opening. Thus, it is unclear if the lowest energy poses, with potential salt bridge interactions, are possible alternate binding modes. It is thus possible that the sum of hydrogen bonding interactions and water-bridging effects could bias the energetic potential to a certain state, such as the one structurally resolved. Thus, further experimental characterization is needed to test these structural hypotheses.

For the central pore cavity, while there should be little to no lipid interactions (excluding the fenestration regions), docking of the central pore cavity produced the most pose variability, likely due to it being the widest receptor site compared to the pore-forming domain and voltage sensing domain. For example, the second orientation of verapamil positioned primarily in the central cavity of rabbit Ca_V_1.1 achieved a RMSD_Min_ of 1.4 Å for RosettaLigand and 2.0 Å for GALigandDock (**Table 2**). However, Z944 bound to hCa_V_3.1, with the wide aromatic end of Z944 in the narrow fenestration and the narrow amide end in the wide cavity achieved a RMSD_Min_ of 0.73 Å for RosettaLigand and 0.84 Å for GALigandDock (**Table 2**). Further, RosettaLigand was able to achieve a P_Near_ value greater than 0.5 in some docking conditions for Z944 (**Table 3; Table S4.1-4.4**), suggesting that the Rosetta docking methods could prove useful when docking similar ligands that bridge between the fenestration and central pore, and target the narrow fenestration with an aromatic moiety.

Small molecules docked in the central cavity were bound to the central pore (6JPA, orientation 2, 6JPB), the channel fenestration region (6JP5, 6JP8, 6U88), or both regions (6JPA orienation1, 6KZP, 6UZ0). It appears that those bound in the channel fenestration produced a lower RMSD_Min_, those primarily bound in the pore center produced the largest RMSD_Min_, while those bound to both regions produced intermediate RMSD_Min_ values, with Z944 docked to human Ca_V_3.1 central pore cavity (6KZP) being an exception. The same trend is roughly observed for P_Near_, where fenestration-only cases have P_Near_ values orders of magnitude greater than pore center cases (**Table S5**). Due to the limited number of cases for each sub-classification, further studies will need to be performed to assess a correlation.

It appears that molecules with predominantly planar aromatic rings and space-filling, “bulky” structures targeting the fenestration can be scored and ranked effectively; both Bay K 8644 (6JP8) and Z944 (6KZP) possess these features in contrast to other small molecules, that possess aromatic or aliphatic rings but are generally more linear and “floppy” targeting the fenestration, like flecainide (6UZ0) and the first orientation of verapamil (6JPA). (**Table S5**). This general trend in increased RMSD_Min_ for non-aromatic containing small molecules when the receptor site is solely the central pore could be due to 1) greater ligand flexibility within a larger area, which requires increased sampling, 2) a bulkier ligand having fewer conformations and orientations relative to the channel, 3) the absence of explicit water molecules and ions that would crowd the pore or directly bind to the ligand (Tikhonov and Zhorov, 2008). Furthermore, if the ligand is expected to bind to a central cavity motif present on multiple subunits, then post-hoc tools implementing symmetry to discriminate unique binding modes will be necessary to calculate an appropriate adjusted RMSD_Min_.

## Conclusions

In this study, we aimed to assess if 1) the RosettaLigand and GALigandDock molecular docking methodologies can recapitulate structurally resolved ion channel – small-molecule binding orientations with known pharmacological significance, 2) if their scoring functions can be used to accurately rank small molecule binding orientation for the purpose of blindly screening small molecules, and 3) if there are practical considerations when docking to specific domains of ion channels. With 2.0 Å RMSD_Min_ as a performance cutoff (Maia et al., 2020; Park et al., 2021), our results demonstrate that both RosettaLigand and GALigandDock can frequently sample the experimentally resolved ligand binding mode with less than 2.0 Å RMSD. However, when using an estimate of the Boltzmann probability for energy “funnel-likeness” (P_Near_) as a scoring function assessment, we currently perceive Rosetta score functions as unable to sufficiently score near-native poses accurately in ion channel docking; from this study, small molecules targeting voltage-sensing domains and bulky small molecules primarily composed of aromatic rings targeting fenestration regions appear to be most suited for score-based ranking. Thus, when performing an ion channel virtual drug discovery campaign, special consideration should be given to sufficient pose sampling to account for multiple rotameric and conformational states, to identify the size of the sampling required for sufficient interface scoring of the receptor site, to identify the appropriate state of the ion channel, to identify inherent channel flexibility that could influence ligand binding, and to potentially identify specific chemical functional groups known experimentally to influence binding to the target when selecting candidate conformations.

## Acknowledgements

We thank members of the VY-Y, FD, and HW laboratories for helpful discussions. We also thank Igor Vorobyov for helpful discussions and Jerome Manera for help with generating 2d structures of ligands. The work in VY-Y lab was supported by NIH grants R61NS127285, R01HL128537, R01HL159304, R01GM132110, R56NS9706, and R01NS128180. BJH was supported by NIH F31 Predoctoral Fellowship F31NS124337 and NIH T32 Predoctoral Fellowship GM007377. We are grateful to OpenEye Scientific for OpenEye Academic License.

