## Supplement File for "Towards high-resolution modeling of small molecule – ion channel interactions"

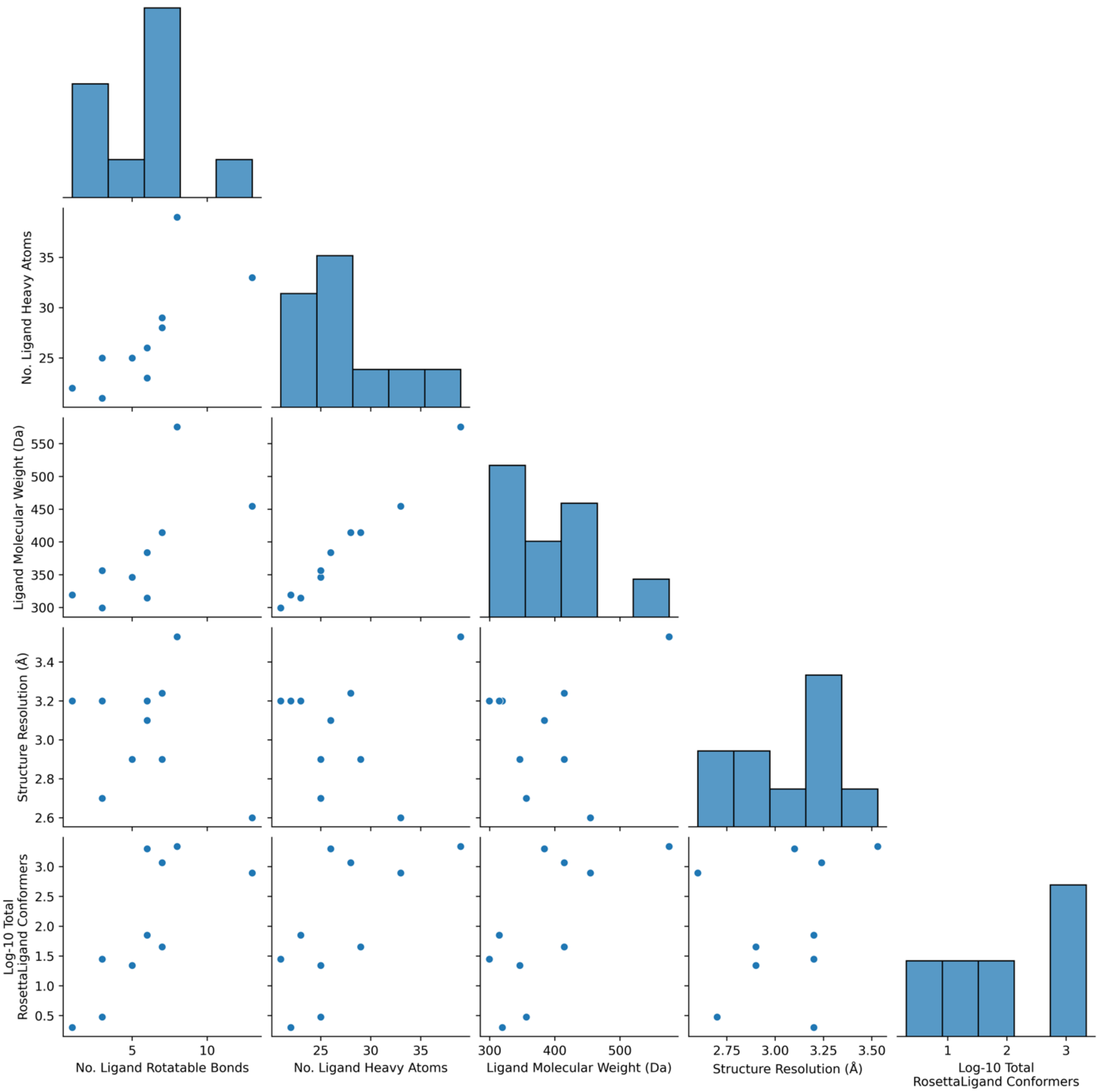


**Figure S1.** Pair plots and histograms of covariates. The logarithm base-10 of the total number of conformers is used in RosettaLigand only.

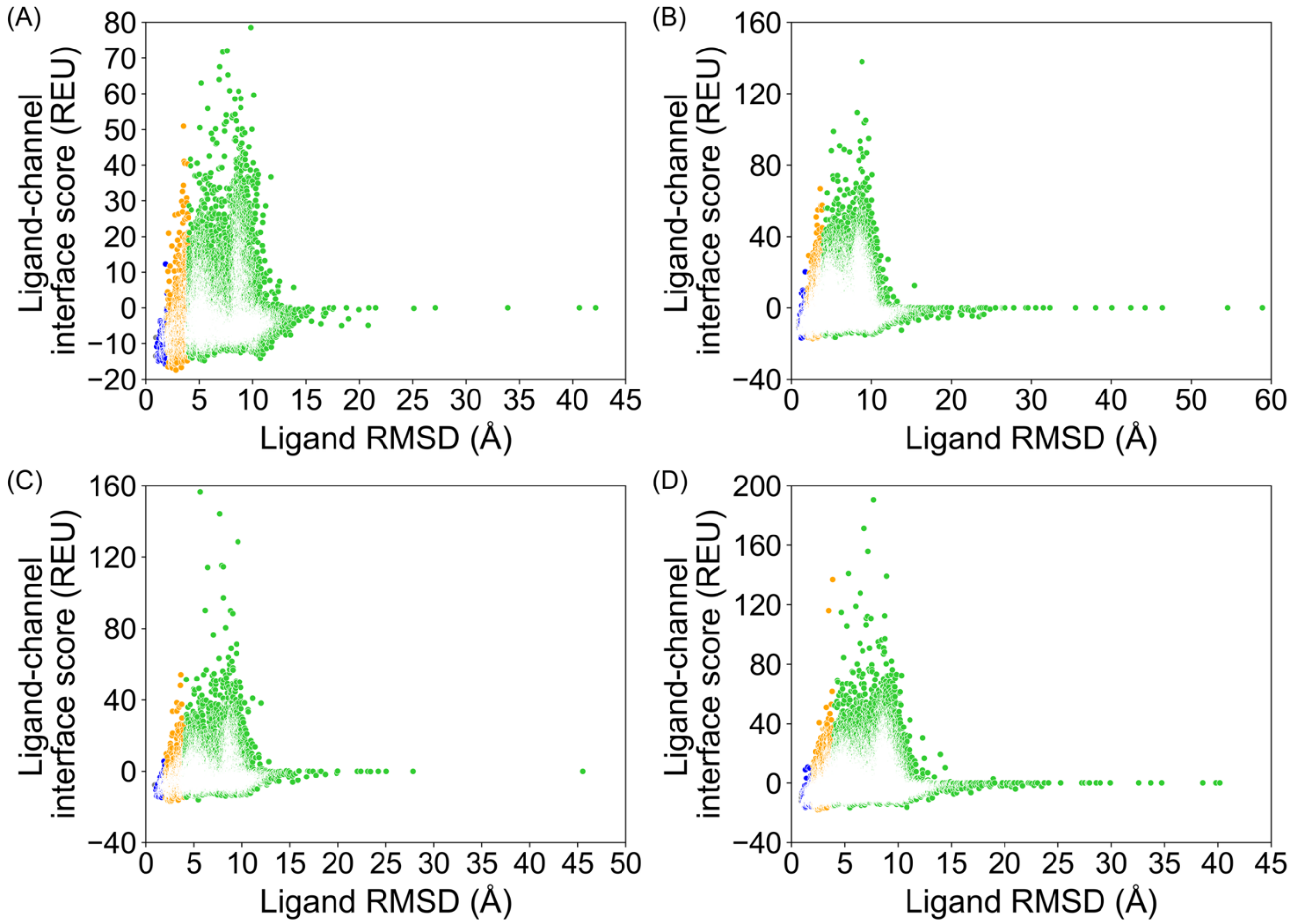


**Figure S2.1** PDB 5EK0, RosettaLigand, all poses from docking. **(A)** 20,000 total poses, all ligand atom interface, **(B)** 100,000 total poses, all ligand atom interface, **(C)** 20,000 total poses, ligand neighbor atom interface, **(D)** 100,000 total poses, ligand neighbor atom interface.

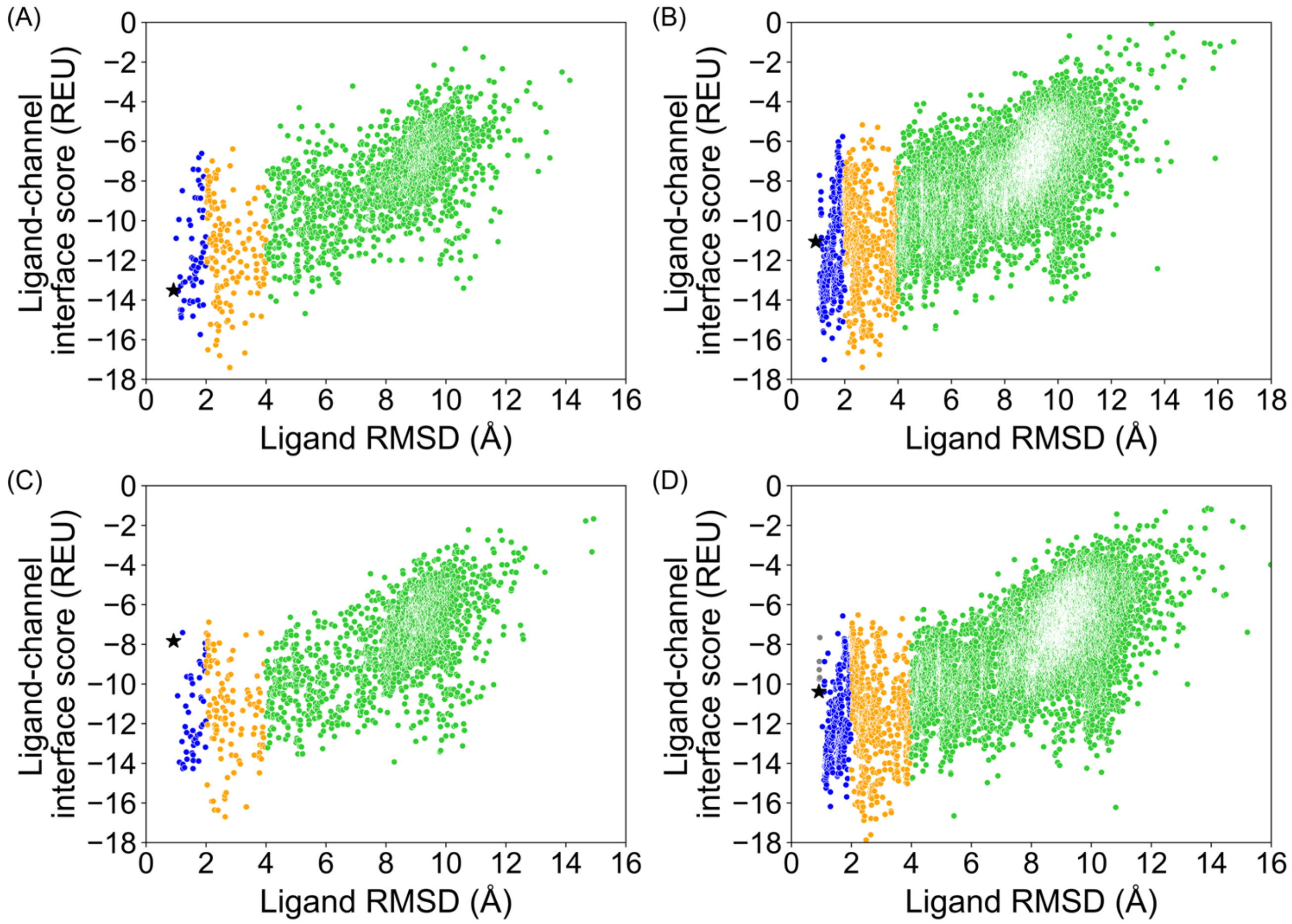


**Figure S2.2** PDB 5EK0, RosettaLigand, top 10 % of total poses by total energy (`total_score`). **(A)** 20,000 total poses, all ligand atom interface, **(B)** 100,000 total poses, all ligand atom interface, **(C)** 20,000 total poses, ligand neighbor atom interface, **(D)** 100,000 total poses, ligand neighbor atom interface.

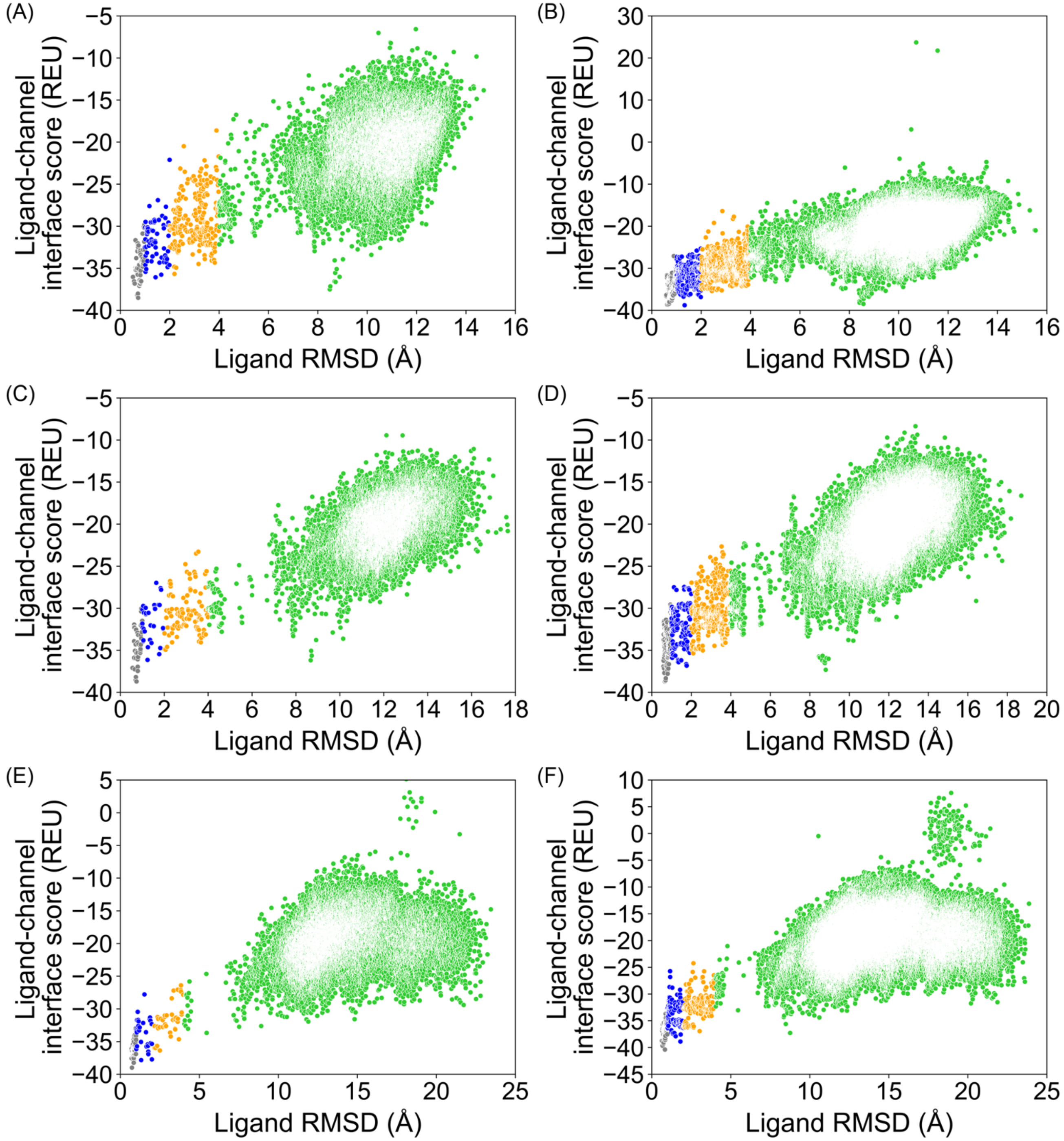


**Figure S2.3** PDB 5EK0, GALigandDock, all poses from docking. **(A)** 20,000 total poses, padding 2 Å, **(B)** 100,000 total poses, padding 2 Å, **(C)** 20,000 total poses, padding 4 Å, **(D)** 100,000 total poses, padding 4 Å, **(E)** 20,000 total poses, padding 7 Å, **(F)** 100,000 total poses, padding 7 Å.

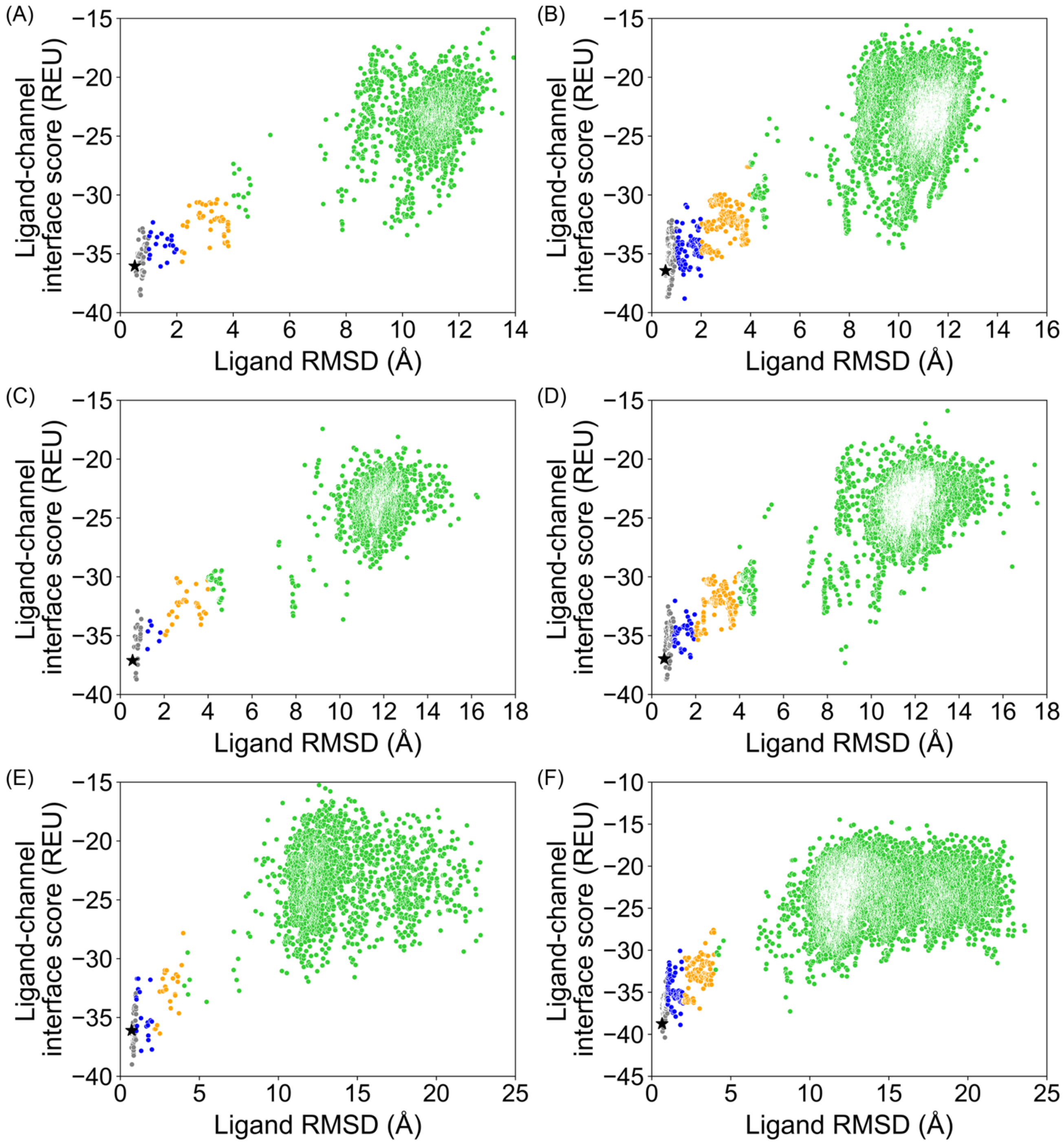


**Figure S2.4** PDB 5EK0, GALigandDock, top 10 % of total poses by total energy (`total_score`). **(A)** 20,000 total poses, padding 2 Å, **(B)** 100,000 total poses, padding 2 Å, **(C)** 20,000 total poses, padding 4 Å, **(D)** 100,000 total poses, padding 4 Å, **(E)** 20,000 total poses, padding 7 Å, **(F)** 100,000 total poses, padding 7 Å.

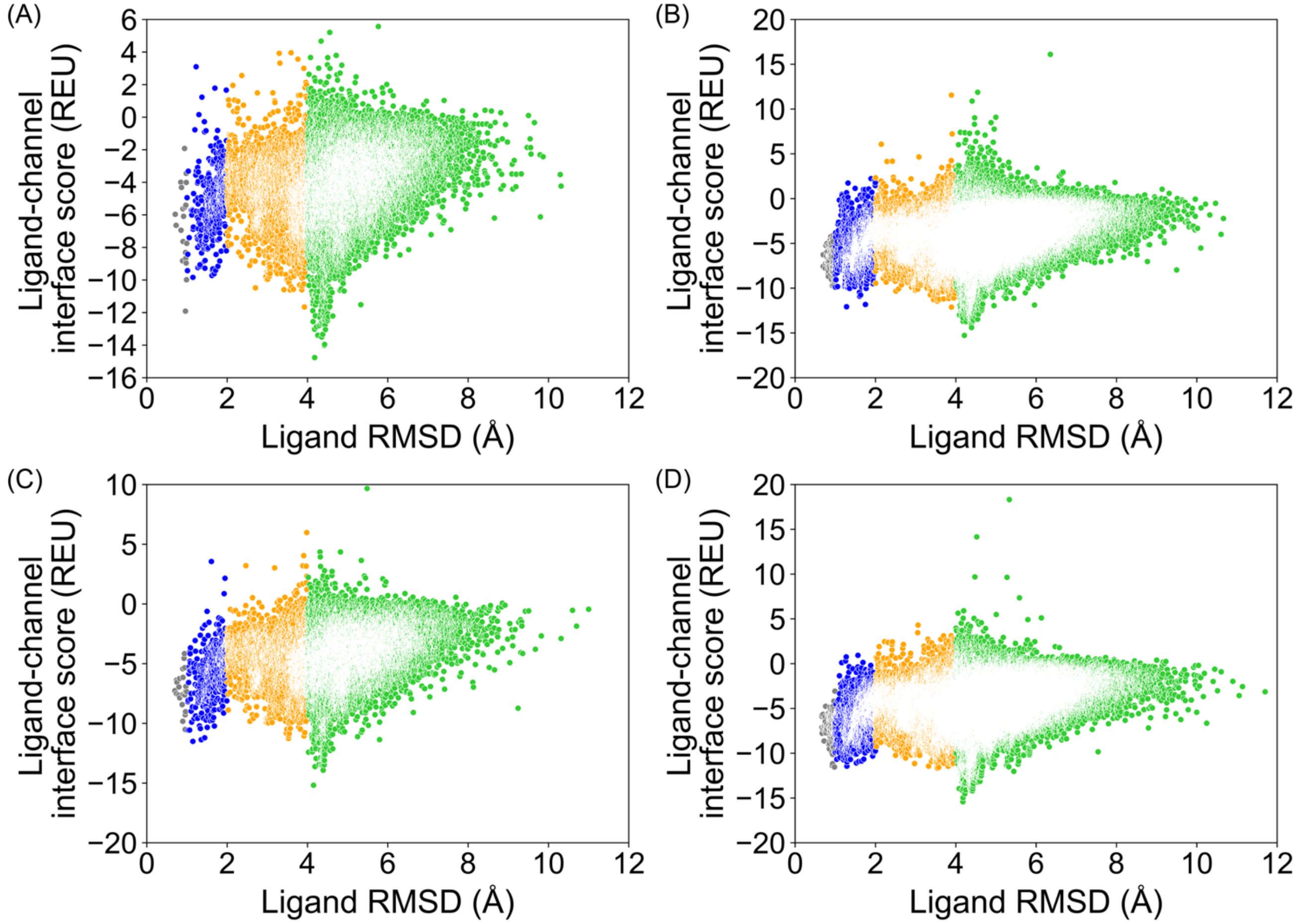


**Figure S3.1** PDB 6J8G, RosettaLigand, all poses from docking. **(A)** 20,000 total poses, all ligand atom interface, **(B)** 100,000 total poses, all ligand atom interface, **(C)** 20,000 total poses, ligand neighbor atom interface, **(D)** 100,000 total poses, ligand neighbor atom interface.

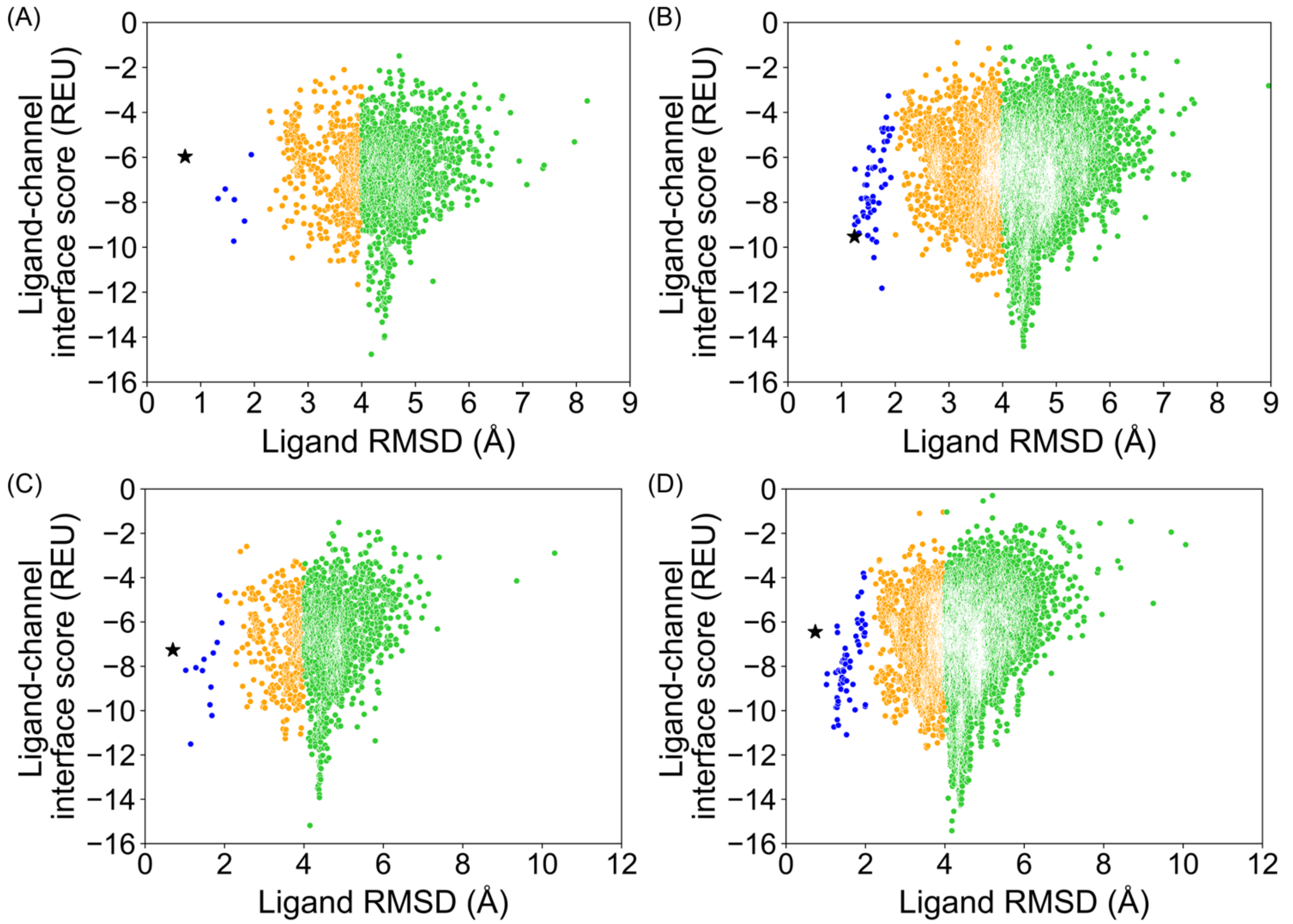


**Figure S3.2** PDB 6J8G, RosettaLigand, top 10 % of total poses by total energy (`total_score`). **(A)** 20,000 total poses, all ligand atom interface, **(B)** 100,000 total poses, all ligand atom interface, **(C)** 20,000 total poses, ligand neighbor atom interface, **(D)** 100,000 total poses, ligand neighbor atom interface.

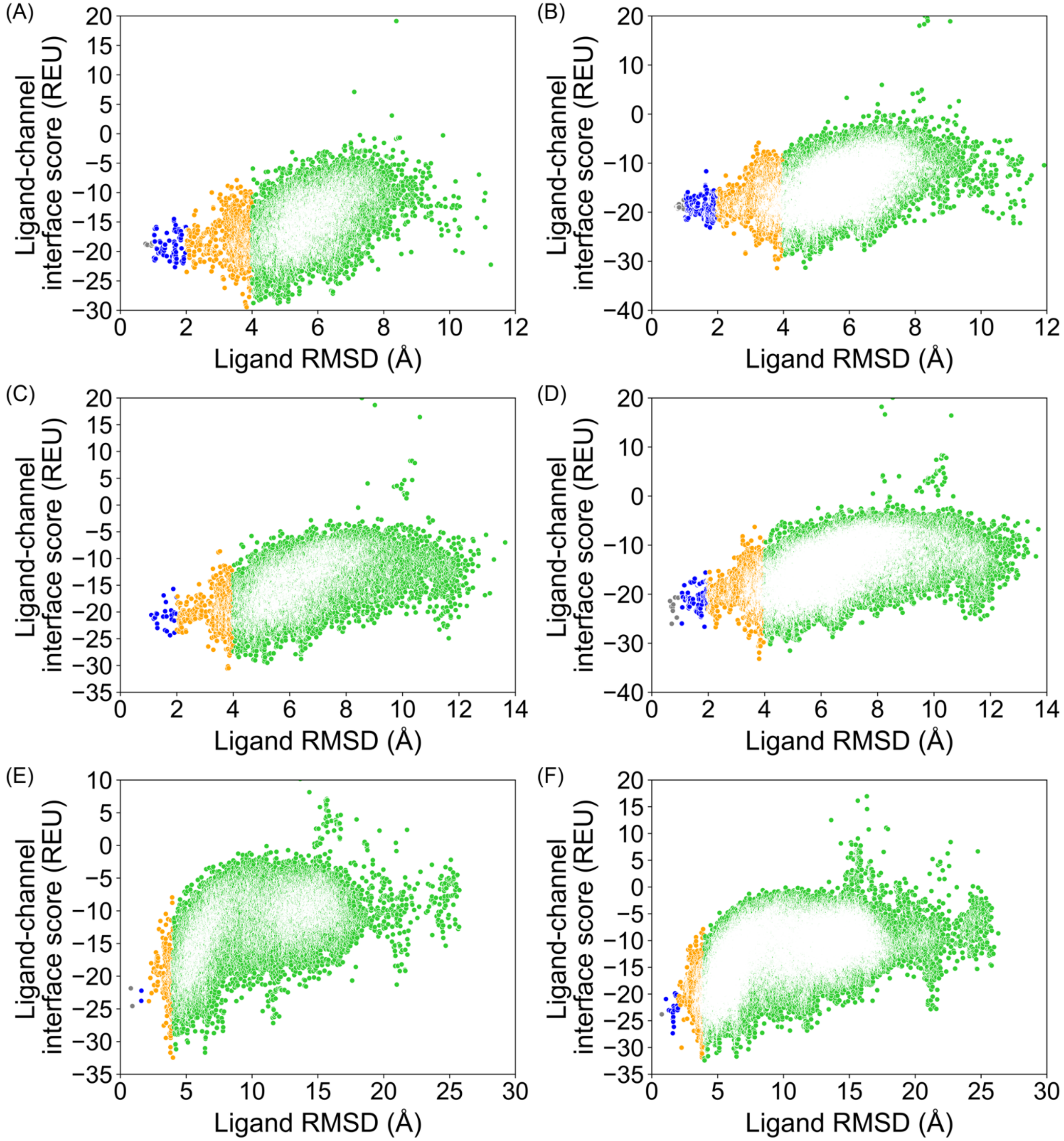


**Figure S3.3** PDB 6J8G, GALigandDock, all poses from docking. **(A)** 20,000 total poses, padding 2 Å, **(B)** 100,000 total poses, padding 2 Å, **(C)** 20,000 total poses, padding 4 Å, **(D)** 100,000 total poses, padding 4 Å, **(E)** 20,000 total poses, padding 7 Å, **(F)** 100,000 total poses, padding 7 Å.

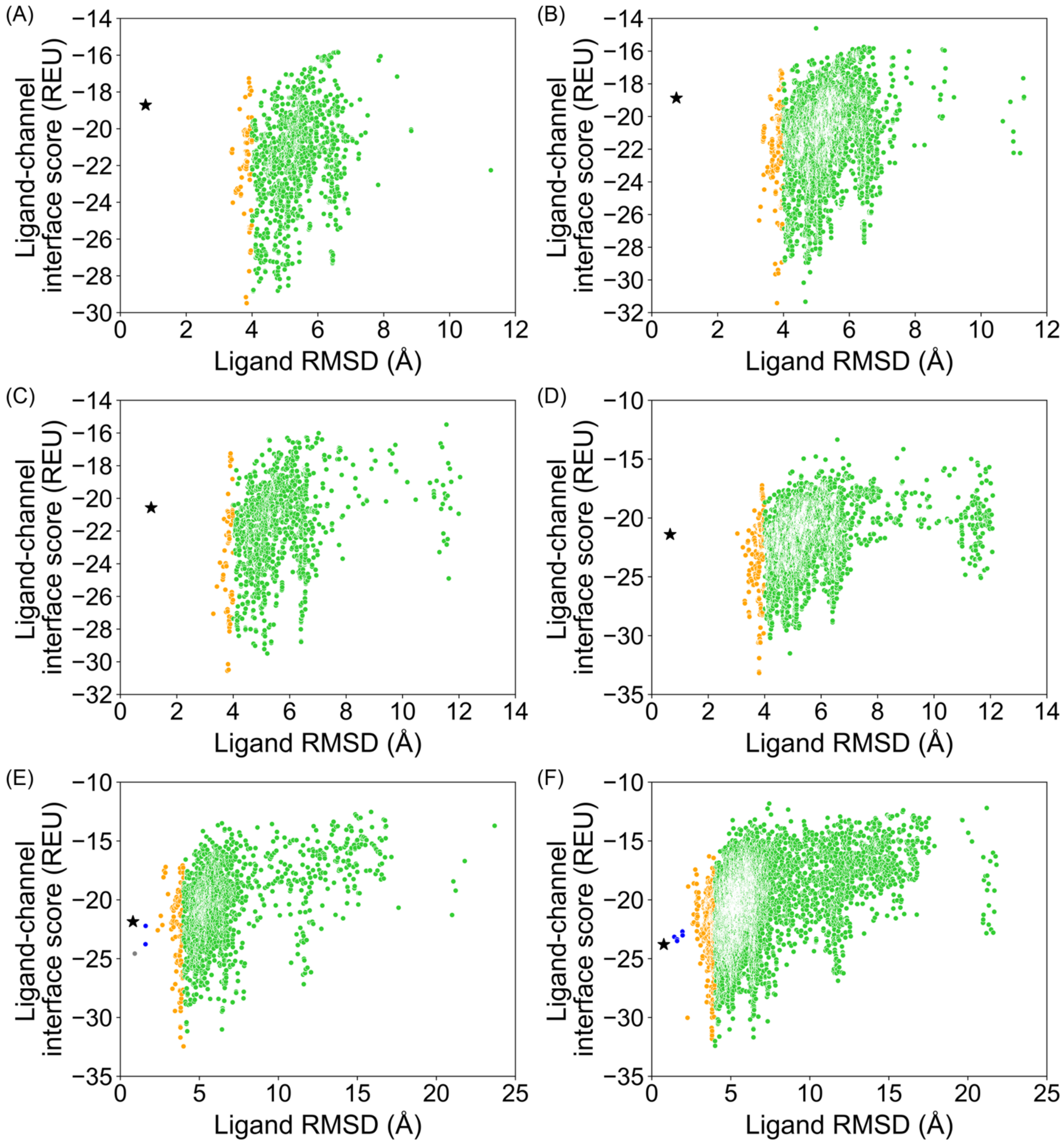


**Figure S3.4** PDB 6J8G, GALigandDock, top 10 % of total poses by total energy (`total_score`). **(A)** 20,000 total poses, padding 2 Å, **(B)** 100,000 total poses, padding 2 Å, **(C)** 20,000 total poses, padding 4 Å, **(D)** 100,000 total poses, padding 4 Å, **(E)** 20,000 total poses, padding 7 Å, **(F)** 100,000 total poses, padding 7 Å.

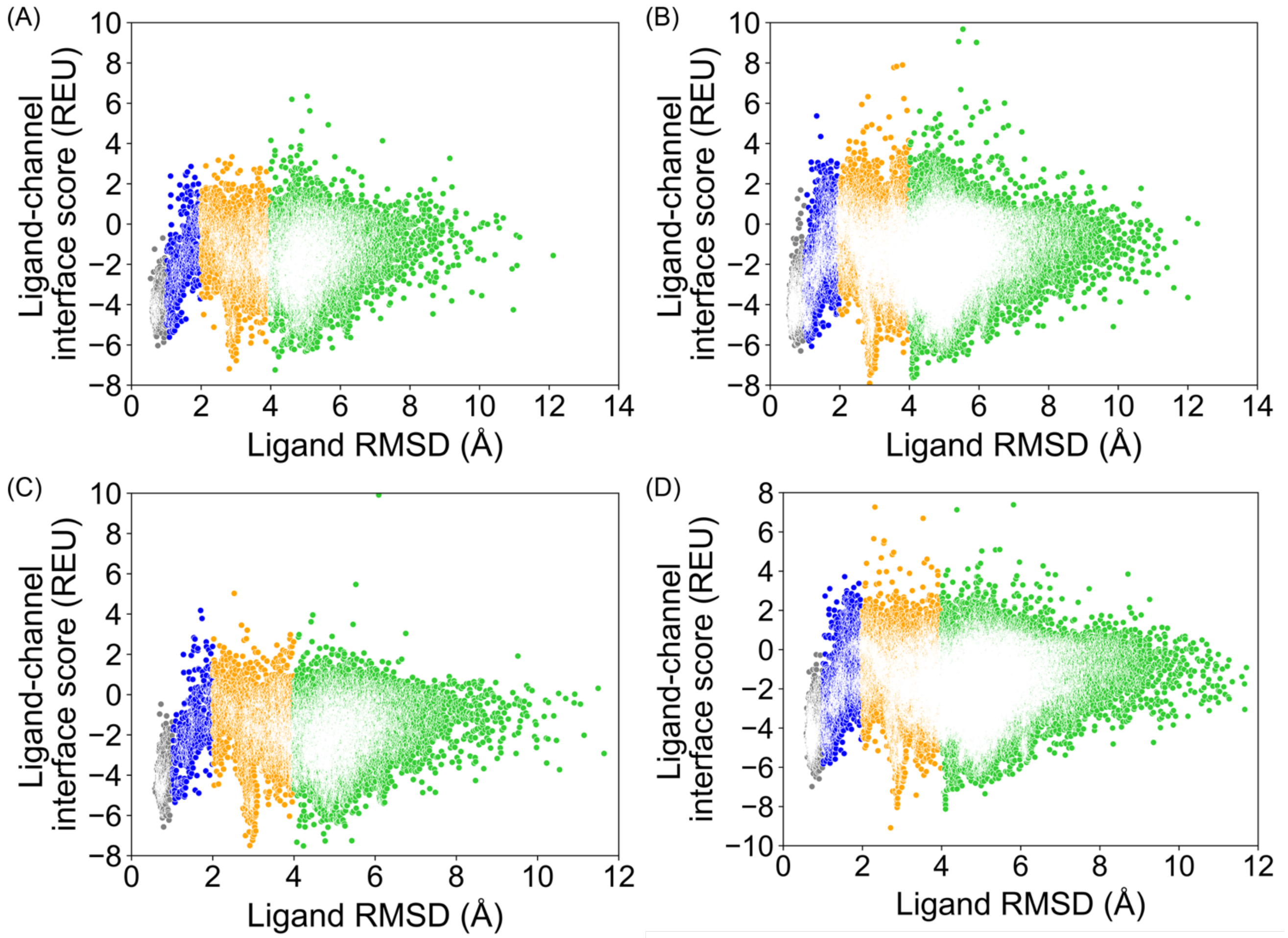


**Figure S4.1** PDB 6J8I, RosettaLigand, all poses from docking. **(A)** 20,000 total poses, all ligand atom interface, **(B)** 100,000 total poses, all ligand atom interface, **(C)** 20,000 total poses, ligand neighbor atom interface, **(D)** 100,000 total poses, ligand neighbor atom interface.

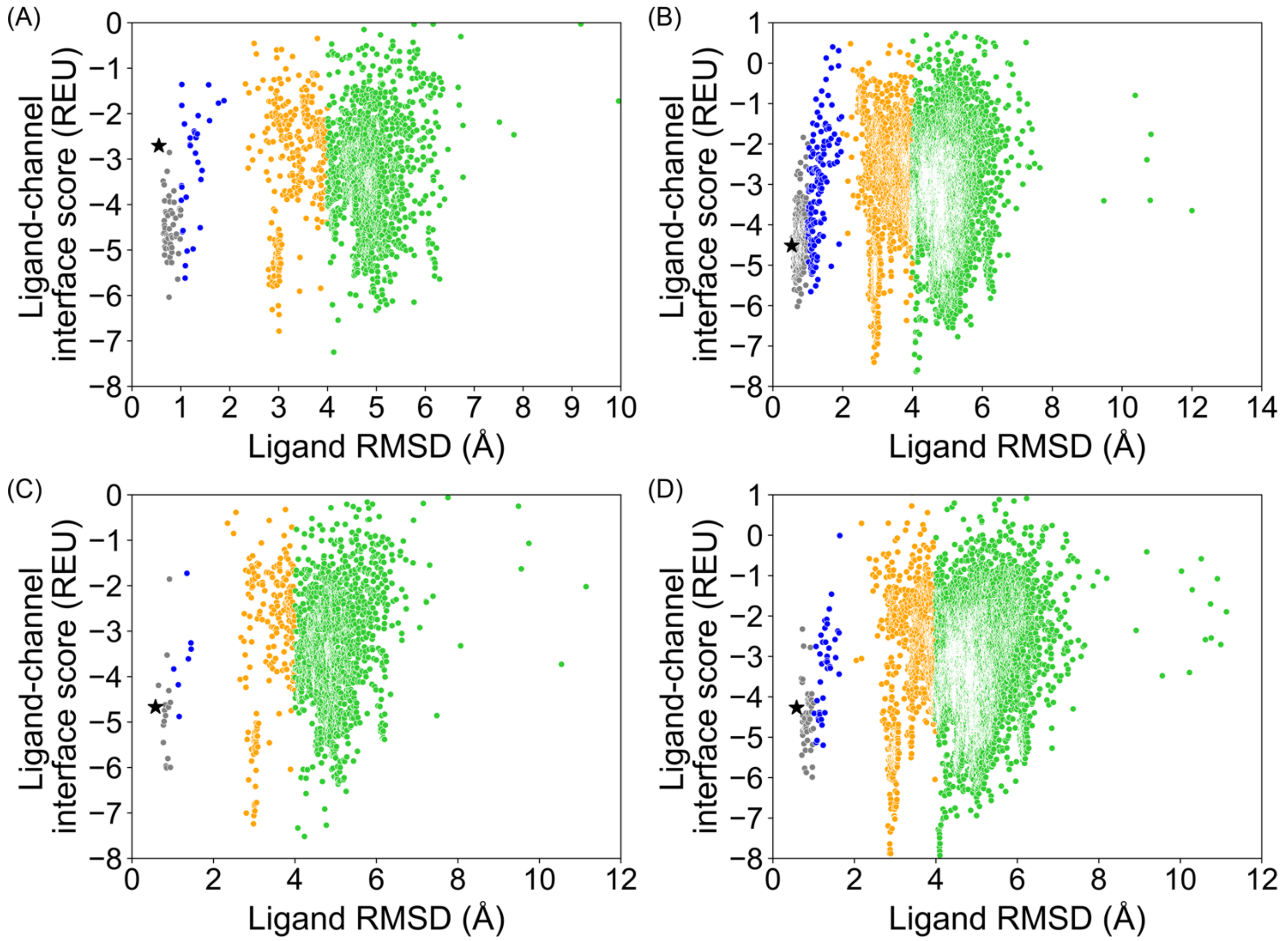


**Figure S4.2** PDB 6J8I, RosettaLigand, top 10 % of total poses by total energy (`total_score`). **(A)** 20,000 total poses, all ligand atom interface, **(B)** 100,000 total poses, all ligand atom interface, **(C)** 20,000 total poses, ligand neighbor atom interface, **(D)** 100,000 total poses, ligand neighbor atom interface.

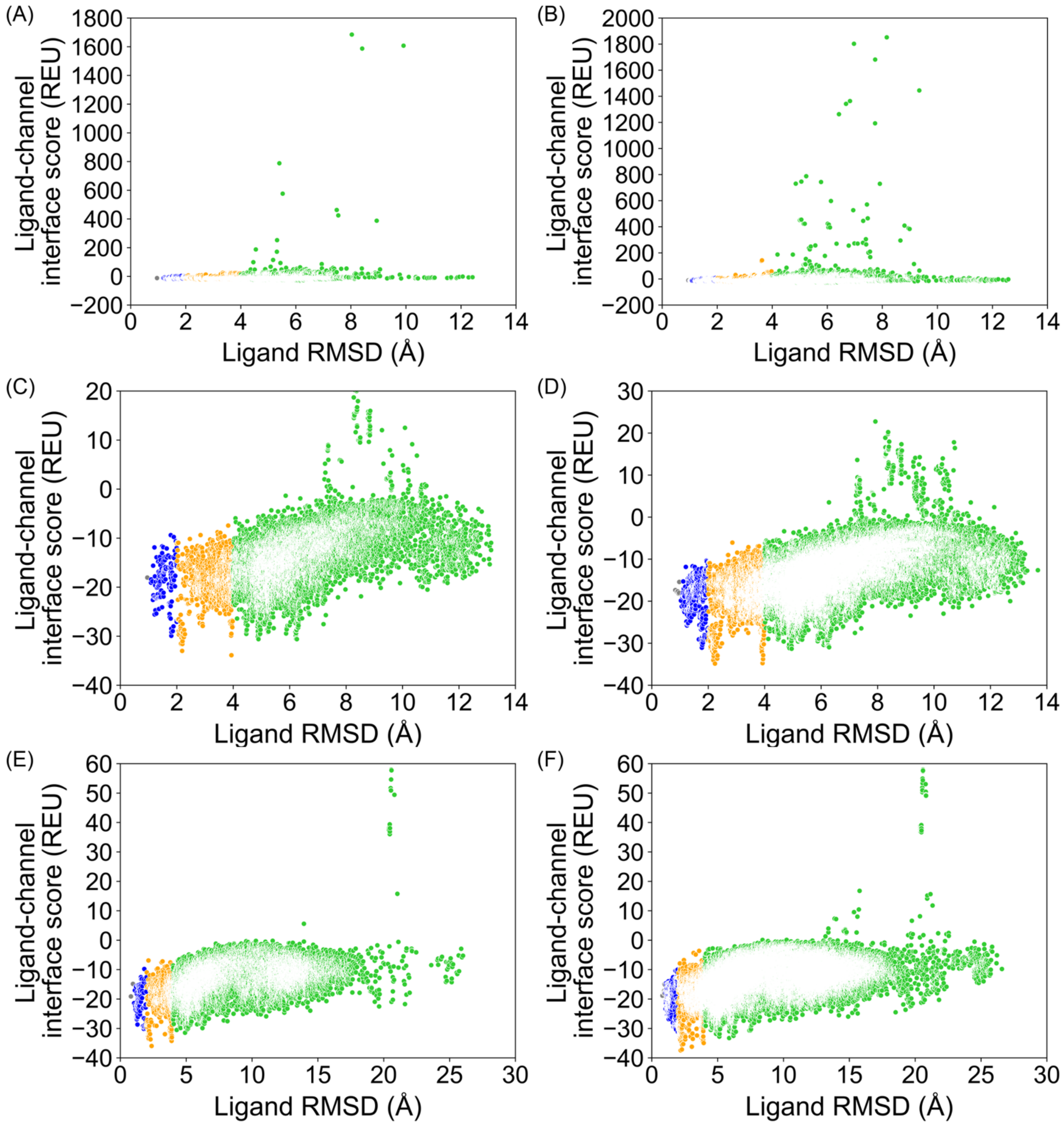


**Figure S4.3** PDB 6J8I, GALigandDock, all poses from docking. **(A)** 20,000 total poses, padding 2 Å, **(B)** 100,000 total poses, padding 2 Å, **(C)** 20,000 total poses, padding 4 Å, **(D)** 100,000 total poses, padding 4 Å, **(E)** 20,000 total poses, padding 7 Å, **(F)** 100,000 total poses, padding 7 Å.

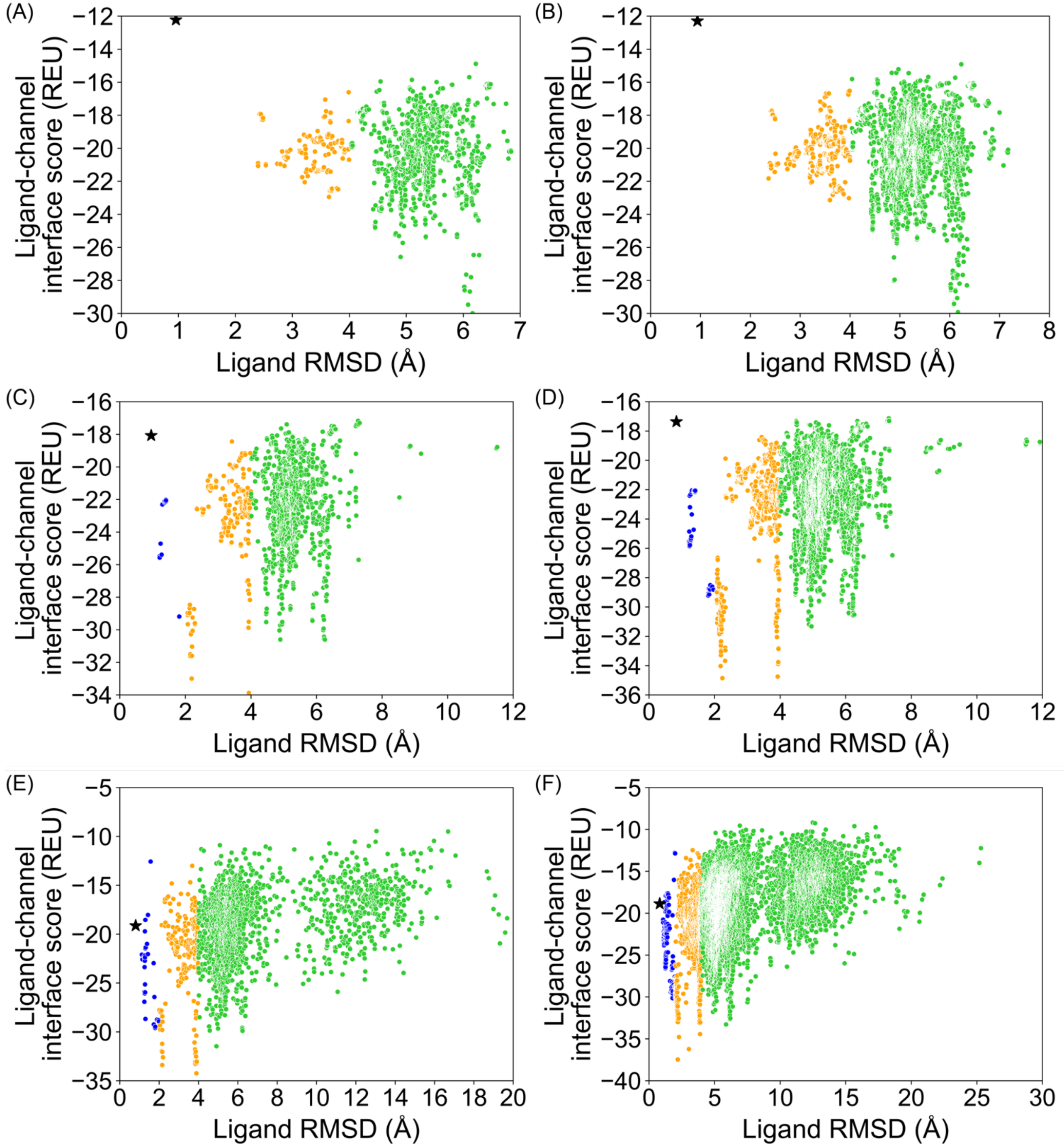


**Figure S4.4** PDB 6J8I, GALigandDock, top 10 % of total poses by total energy (`total_score`). **(A)** 20,000 total poses, padding 2 Å, **(B)** 100,000 total poses, padding 2 Å, **(C)** 20,000 total poses, padding 4 Å, **(D)** 100,000 total poses, padding 4 Å, **(E)** 20,000 total poses, padding 7 Å, **(F)** 100,000 total poses, padding 7 Å.

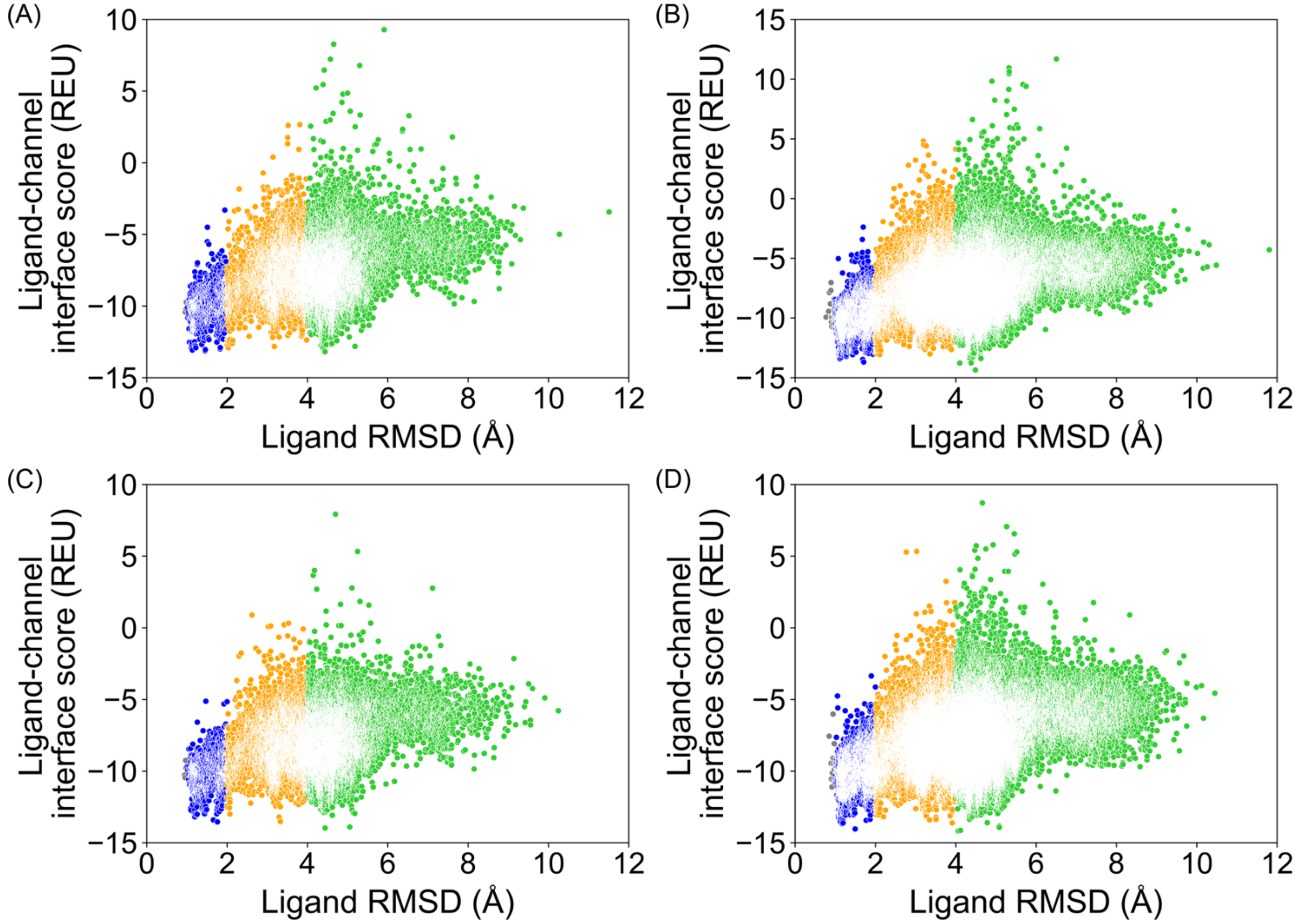


**Figure S5.1** PDB 6JP5, RosettaLigand, all poses from docking. **(A)** 20,000 total poses, all ligand atom interface, **(B)** 100,000 total poses, all ligand atom interface, **(C)** 20,000 total poses, ligand neighbor atom interface, **(D)** 100,000 total poses, ligand neighbor atom interface.

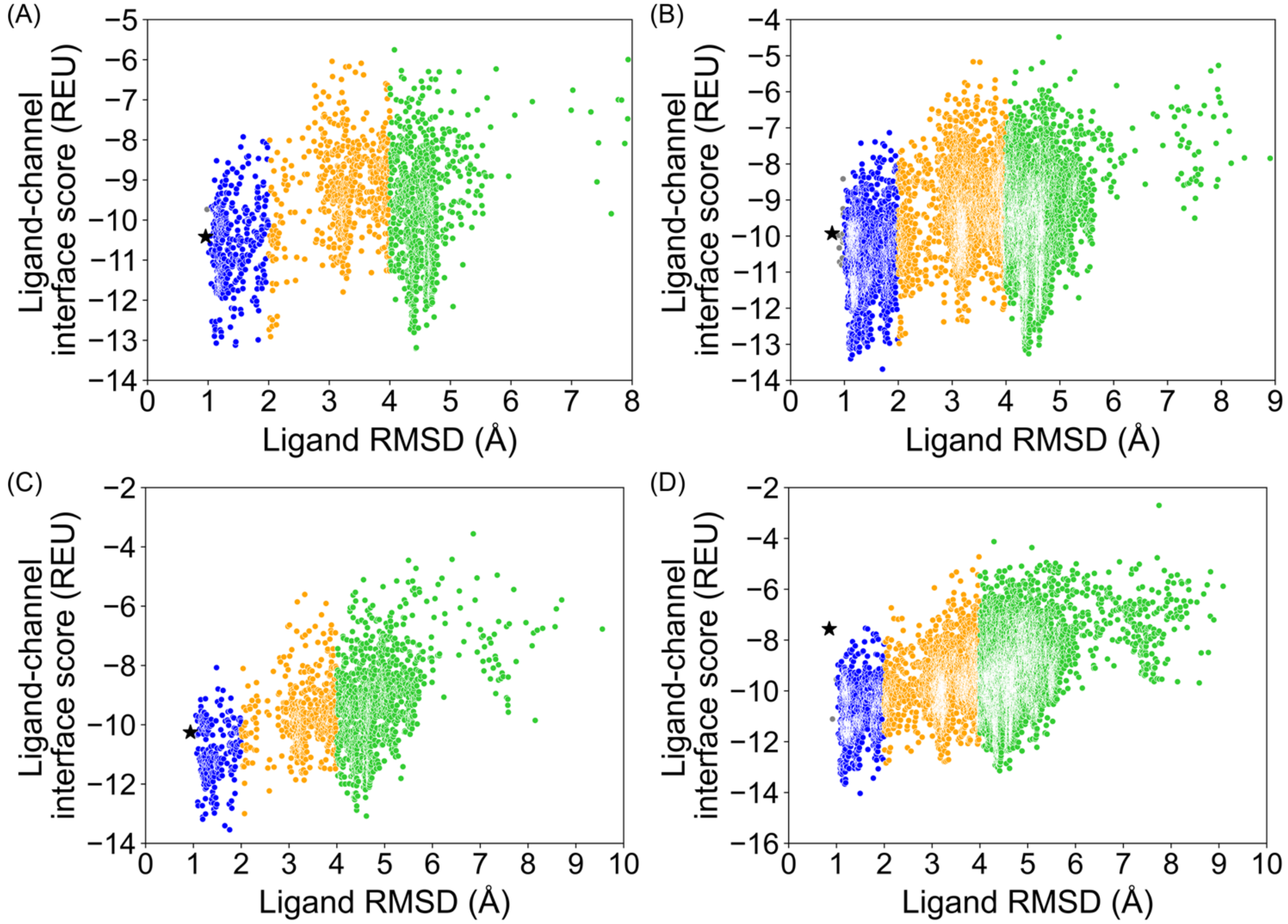


**Figure S5.2** PDB 6JP5, RosettaLigand, top 10 % of total poses by total energy (`total_score`). **(A)** 20,000 total poses, all ligand atom interface, **(B)** 100,000 total poses, all ligand atom interface, **(C)** 20,000 total poses, ligand neighbor atom interface, **(D)** 100,000 total poses, ligand neighbor atom interface.

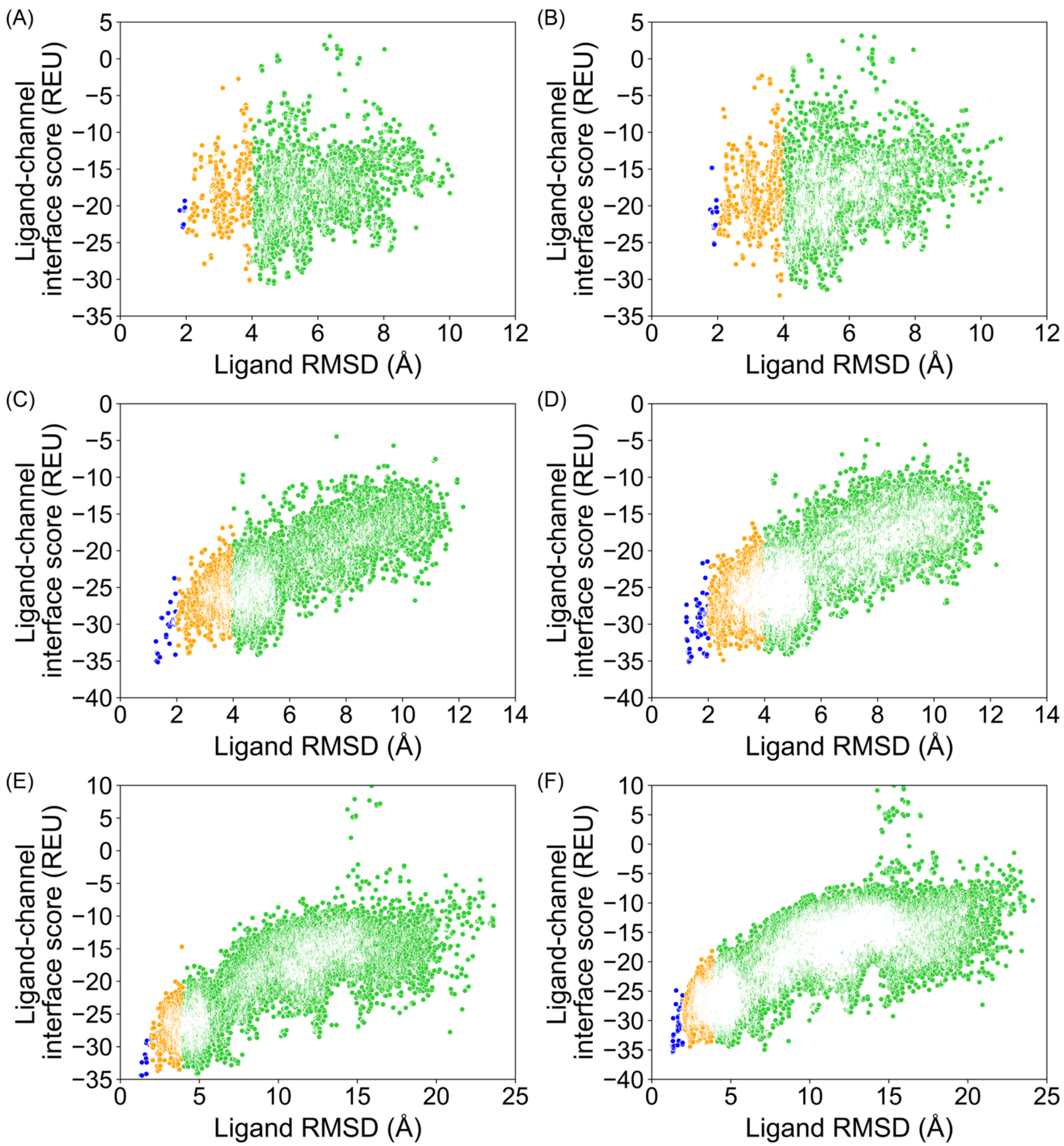


**Figure S5.3** PDB 6JP5, GALigandDock, all poses from docking. **(A)** 20,000 total poses, padding 2 Å, **(B)** 100,000 total poses, padding 2 Å, **(C)** 20,000 total poses, padding 4 Å, **(D)** 100,000 total poses, padding 4 Å, **(E)** 20,000 total poses, padding 7 Å, **(F)** 100,000 total poses, padding 7 Å.

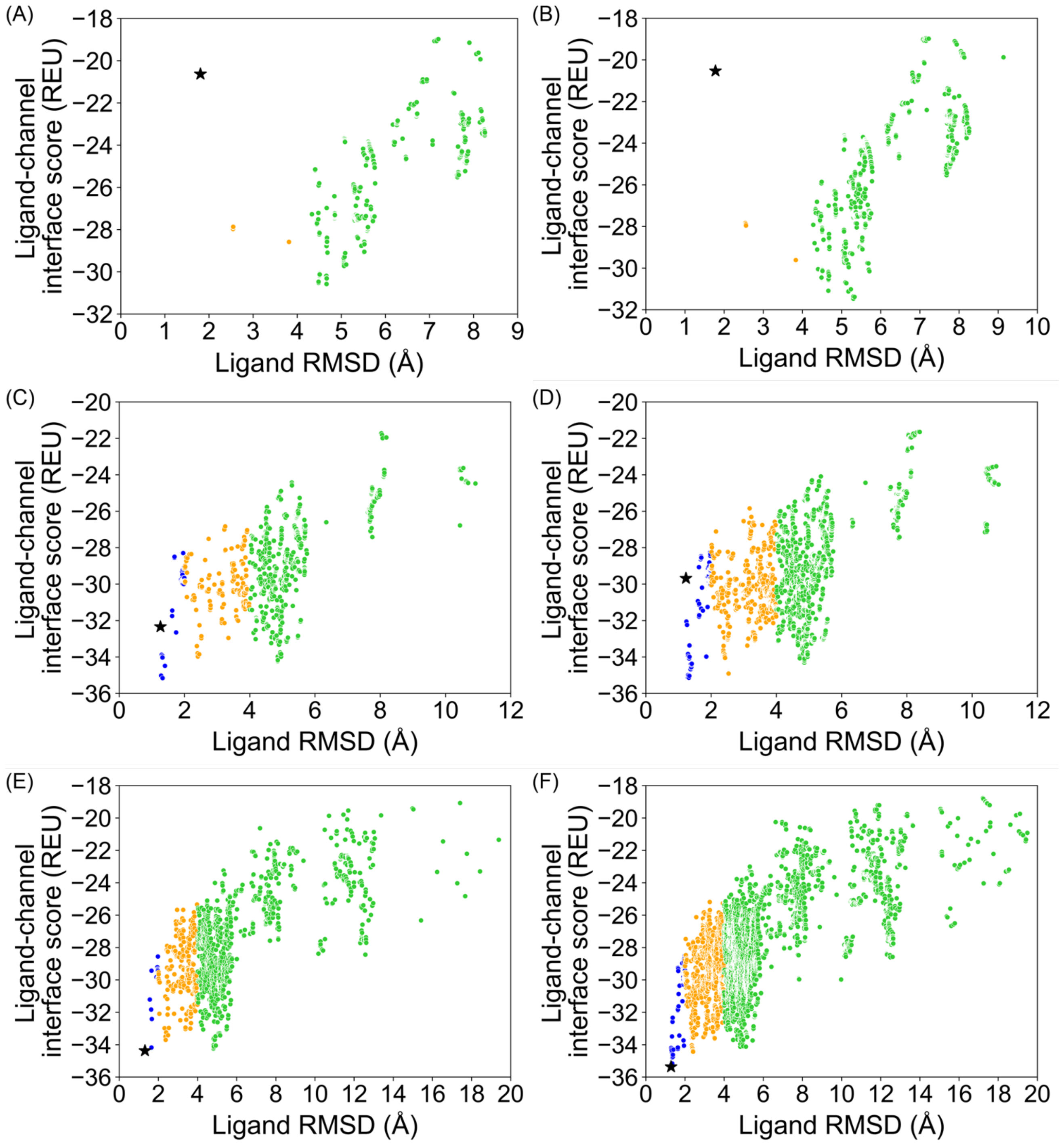


**Figure S5.4** PDB 6JP5, GALigandDock, top 10 % of total poses by total energy (`total_score`). **(A)** 20,000 total poses, padding 2 Å, **(B)** 100,000 total poses, padding 2 Å, **(C)** 20,000 total poses, padding 4 Å, **(D)** 100,000 total poses, padding 4 Å, **(E)** 20,000 total poses, padding 7 Å, **(F)** 100,000 total poses, padding 7 Å.

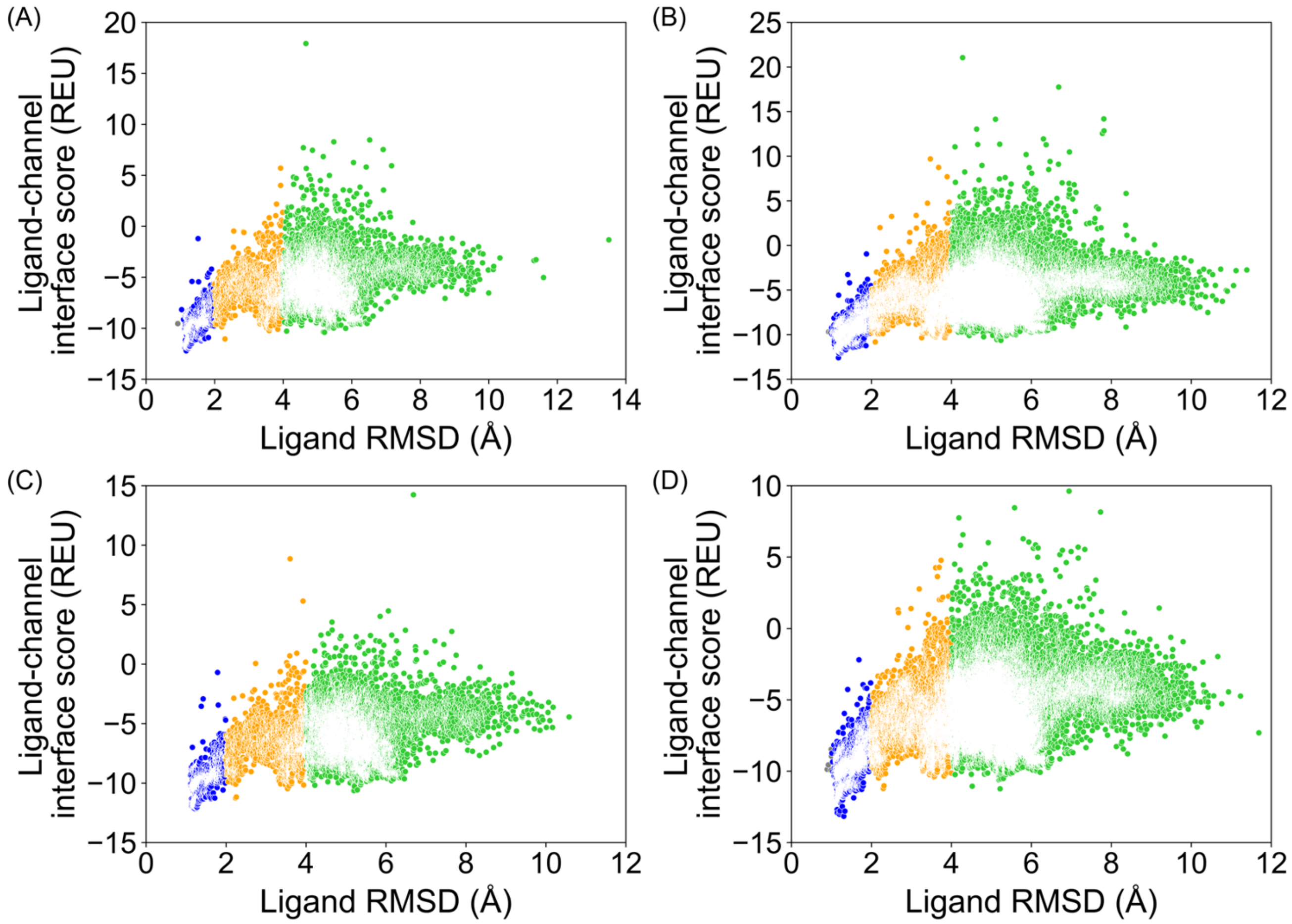


**Figure S6.1** PDB 6JP8, RosettaLigand, all poses from docking. **(A)** 20,000 total poses, all ligand atom interface, **(B)** 100,000 total poses, all ligand atom interface, **(C)** 20,000 total poses, ligand neighbor atom interface, **(D)** 100,000 total poses, ligand neighbor atom interface.

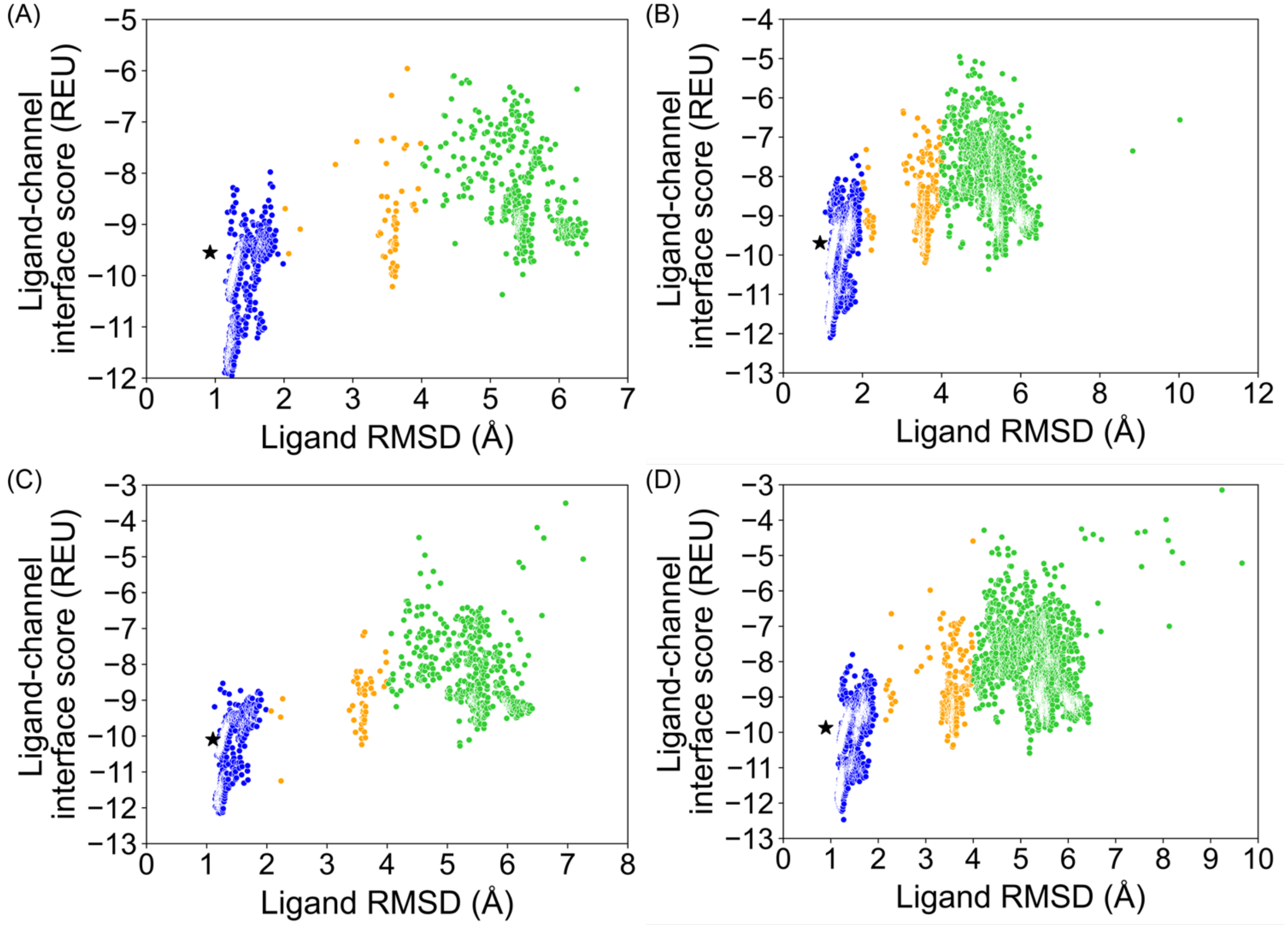


**Figure S6.2** PDB 6JP8, RosettaLigand, top 10 % of total poses by total energy (`total_score`). **(A)** 20,000 total poses, all ligand atom interface, **(B)** 100,000 total poses, all ligand atom interface, **(C)** 20,000 total poses, ligand neighbor atom interface, **(D)** 100,000 total poses, ligand neighbor atom interface.

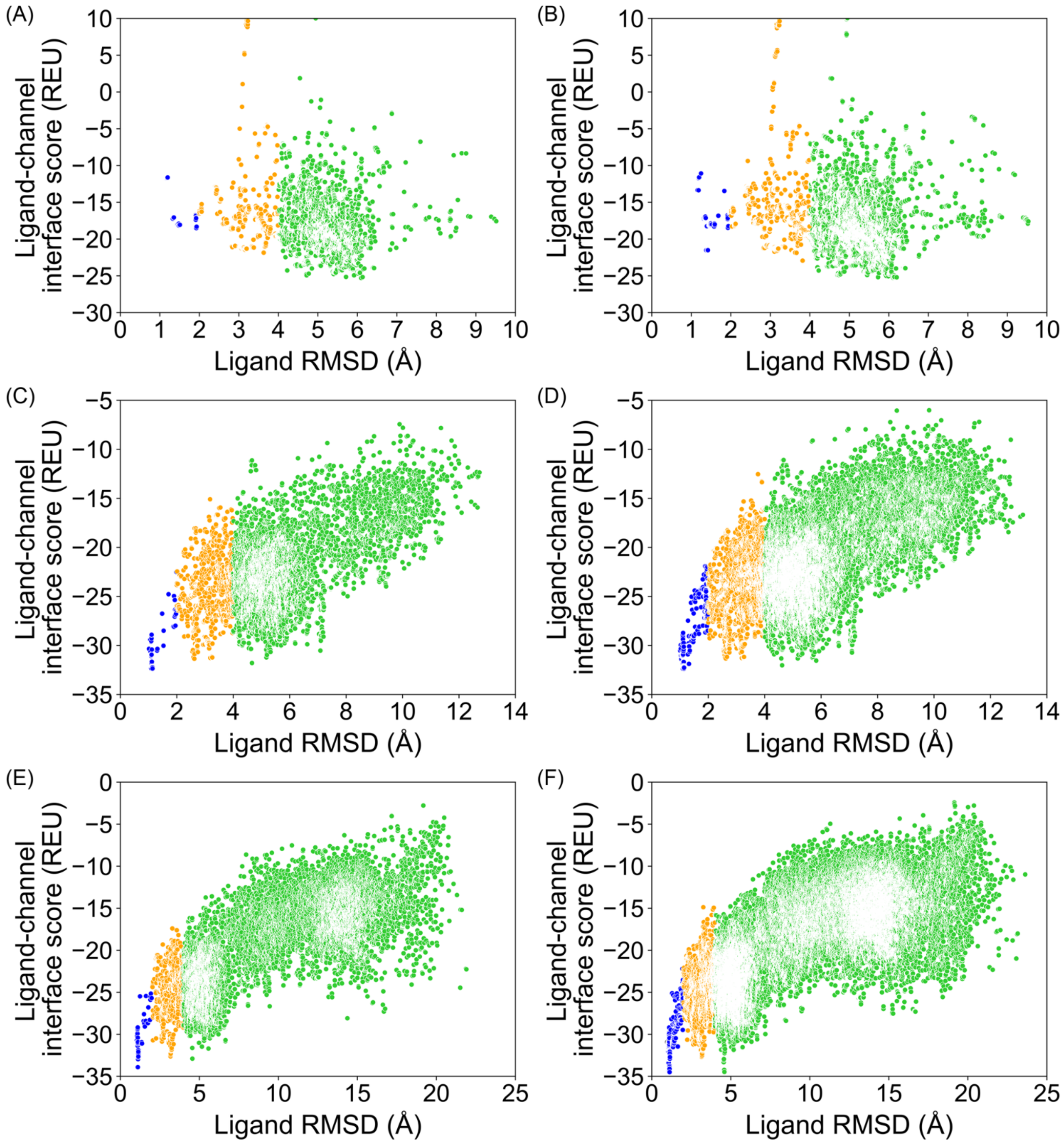


**Figure S6.3** PDB 6JP8, GALigandDock, all poses from docking. **(A)** 20,000 total poses, padding 2 Å, **(B)** 100,000 total poses, padding 2 Å, **(C)** 20,000 total poses, padding 4 Å, **(D)** 100,000 total poses, padding 4 Å, **(E)** 20,000 total poses, padding 7 Å, **(F)** 100,000 total poses, padding 7 Å.

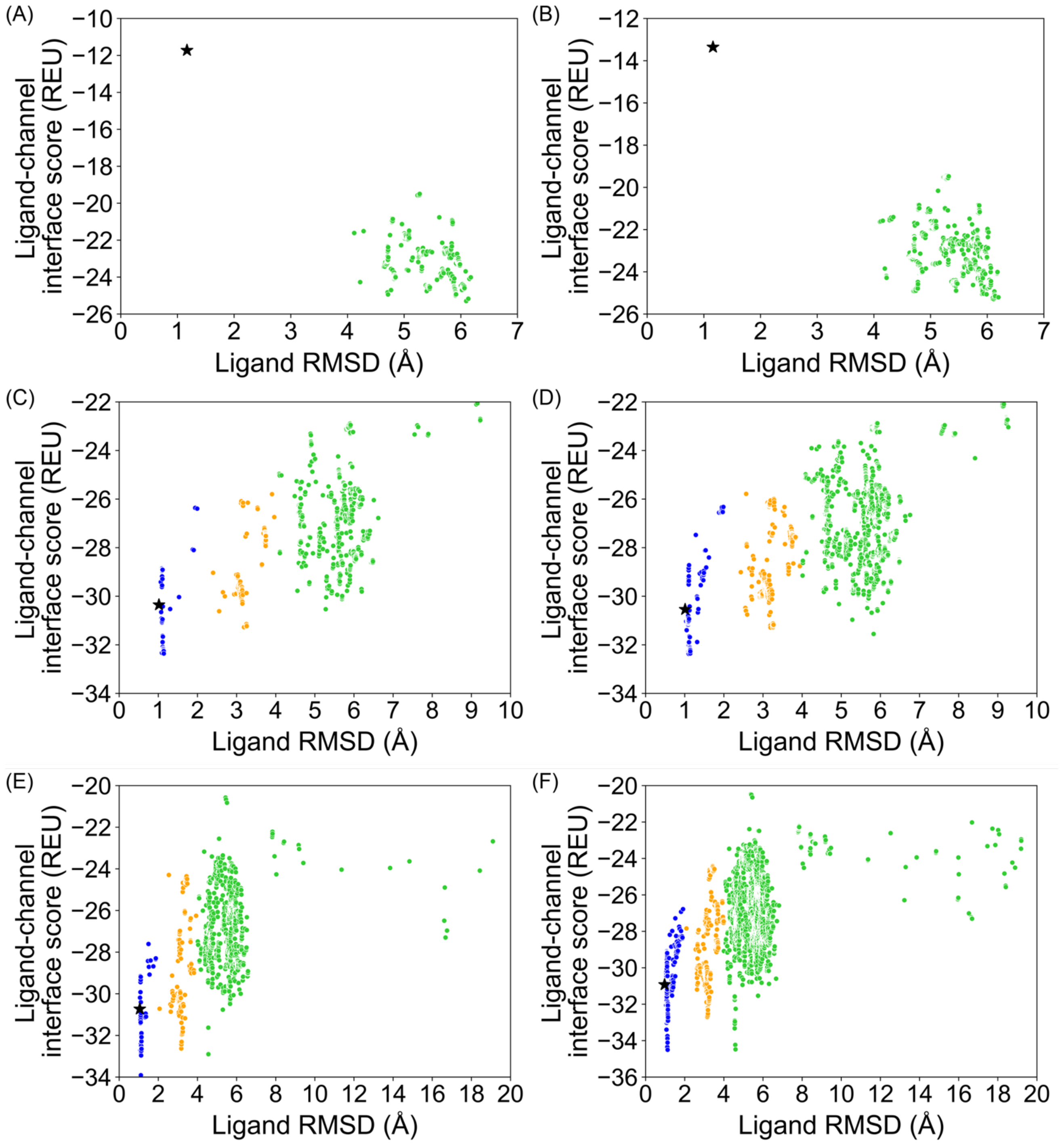


**Figure S6.4** PDB 6JP8, GALigandDock, top 10 % of total poses by total energy (`total_score`). **(A)** 20,000 total poses, padding 2 Å, **(B)** 100,000 total poses, padding 2 Å, **(C)** 20,000 total poses, padding 4 Å, **(D)** 100,000 total poses, padding 4 Å, **(E)** 20,000 total poses, padding 7 Å, **(F)** 100,000 total poses, padding 7 Å.

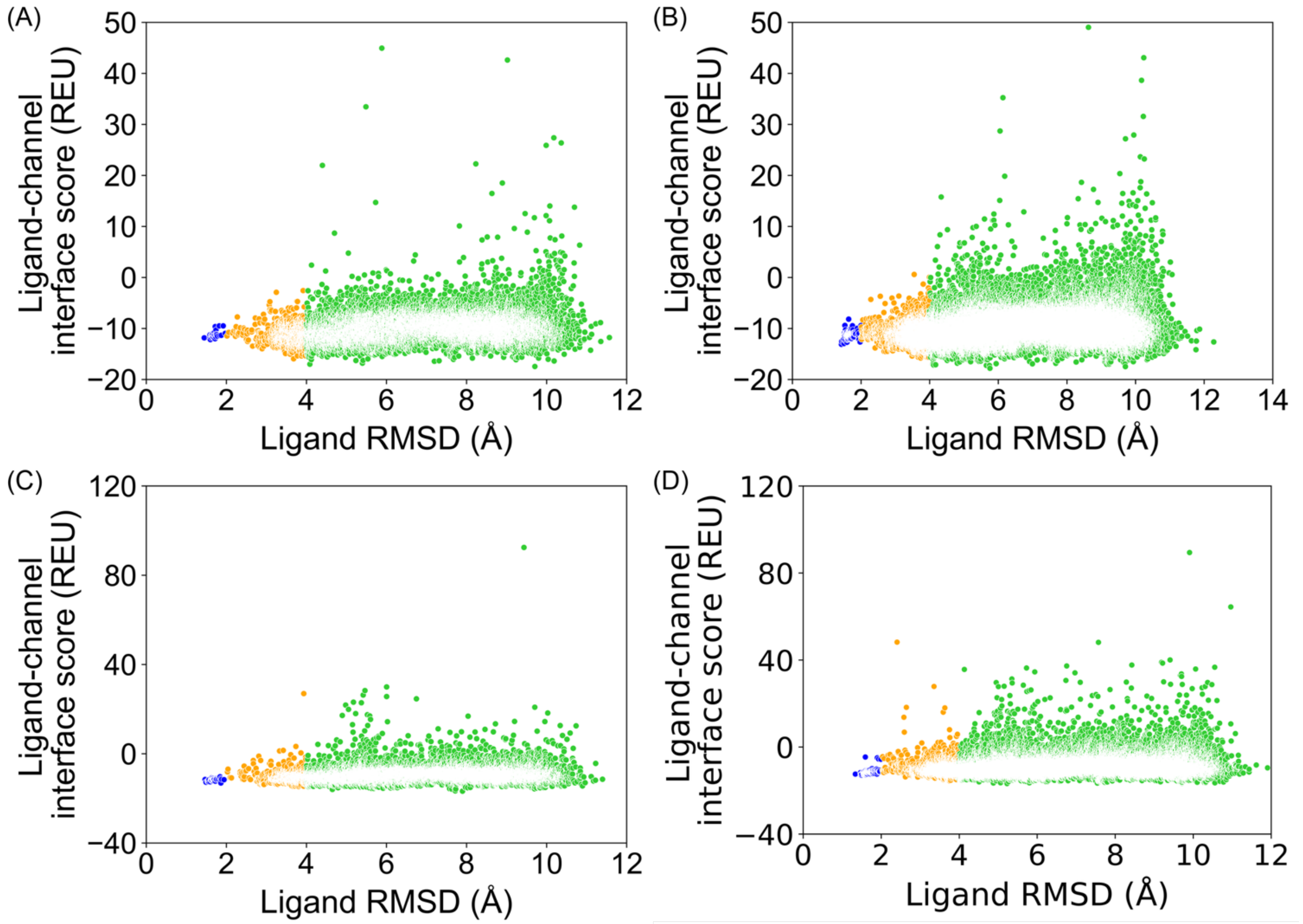


**Figure S7.1** PDB 6JPAV1, RosettaLigand, all poses from docking. **(A)** 20,000 total poses, all ligand atom interface, **(B)** 100,000 total poses, all ligand atom interface, **(C)** 20,000 total poses, ligand neighbor atom interface, **(D)** 100,000 total poses, ligand neighbor atom interface.

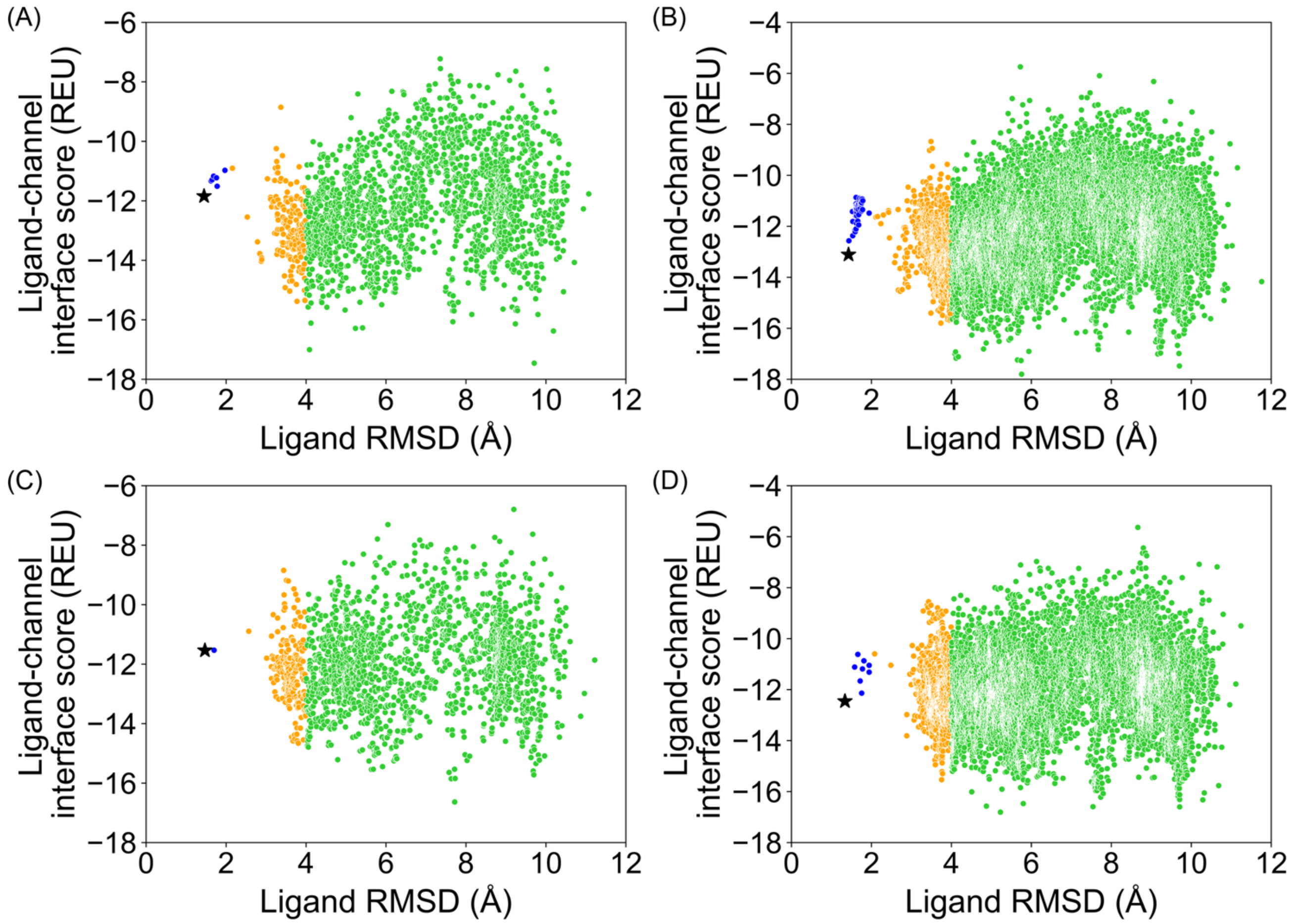


**Figure S7.2** PDB 6JPAV1, RosettaLigand, top 10 % of total poses by total energy (`total_score`). **(A)** 20,000 total poses, all ligand atom interface, **(B)** 100,000 total poses, all ligand atom interface, **(C)** 20,000 total poses, ligand neighbor atom interface, **(D)** 100,000 total poses, ligand neighbor atom interface.

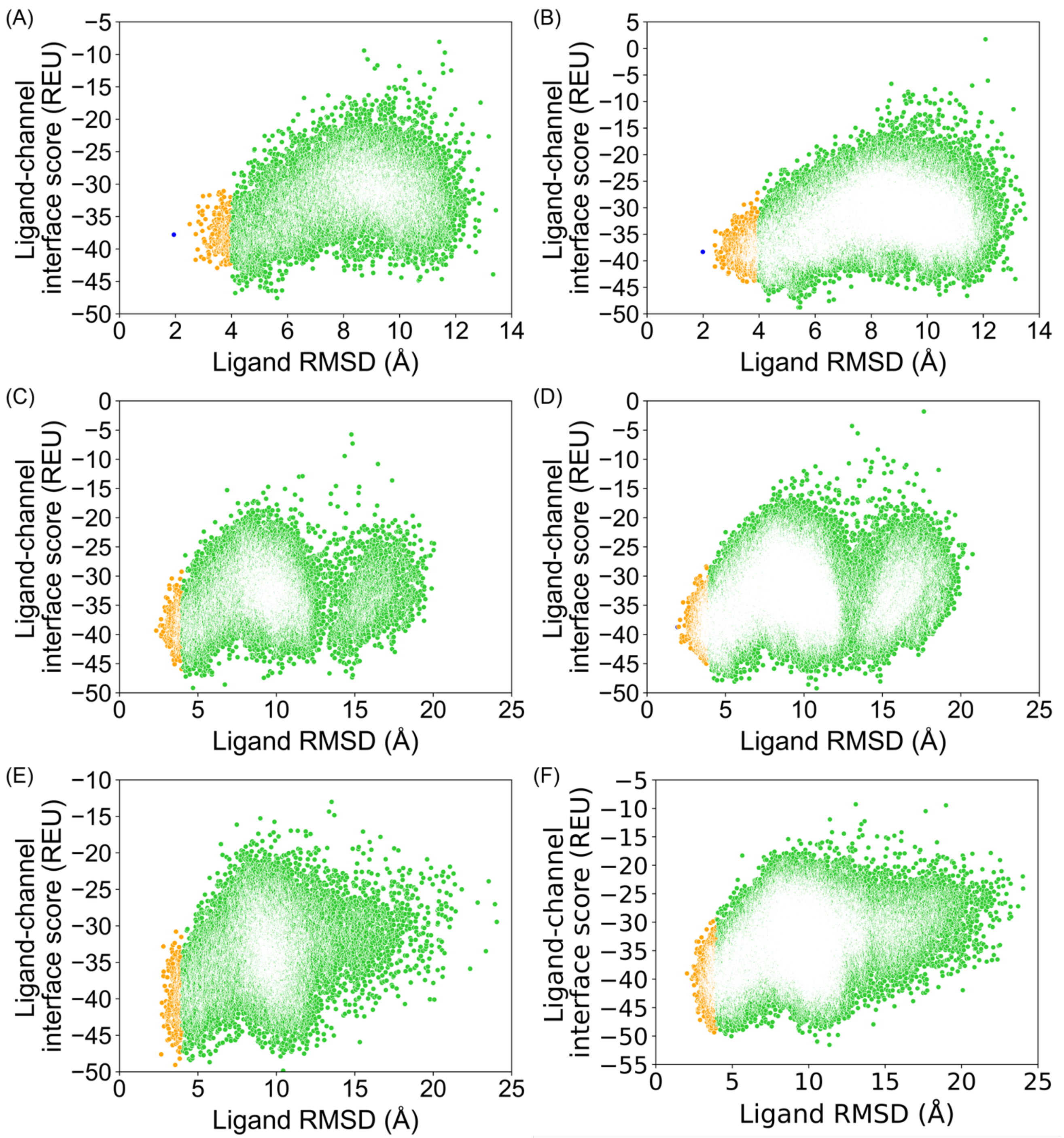


**Figure S7.3** PDB 6JPAV1, GALigandDock, all poses from docking. **(A)** 20,000 total poses, padding 2 Å, **(B)** 100,000 total poses, padding 2 Å, **(C)** 20,000 total poses, padding 4 Å, **(D)** 100,000 total poses, padding 4 Å, **(E)** 20,000 total poses, padding 7 Å, **(F)** 100,000 total poses, padding 7 Å.

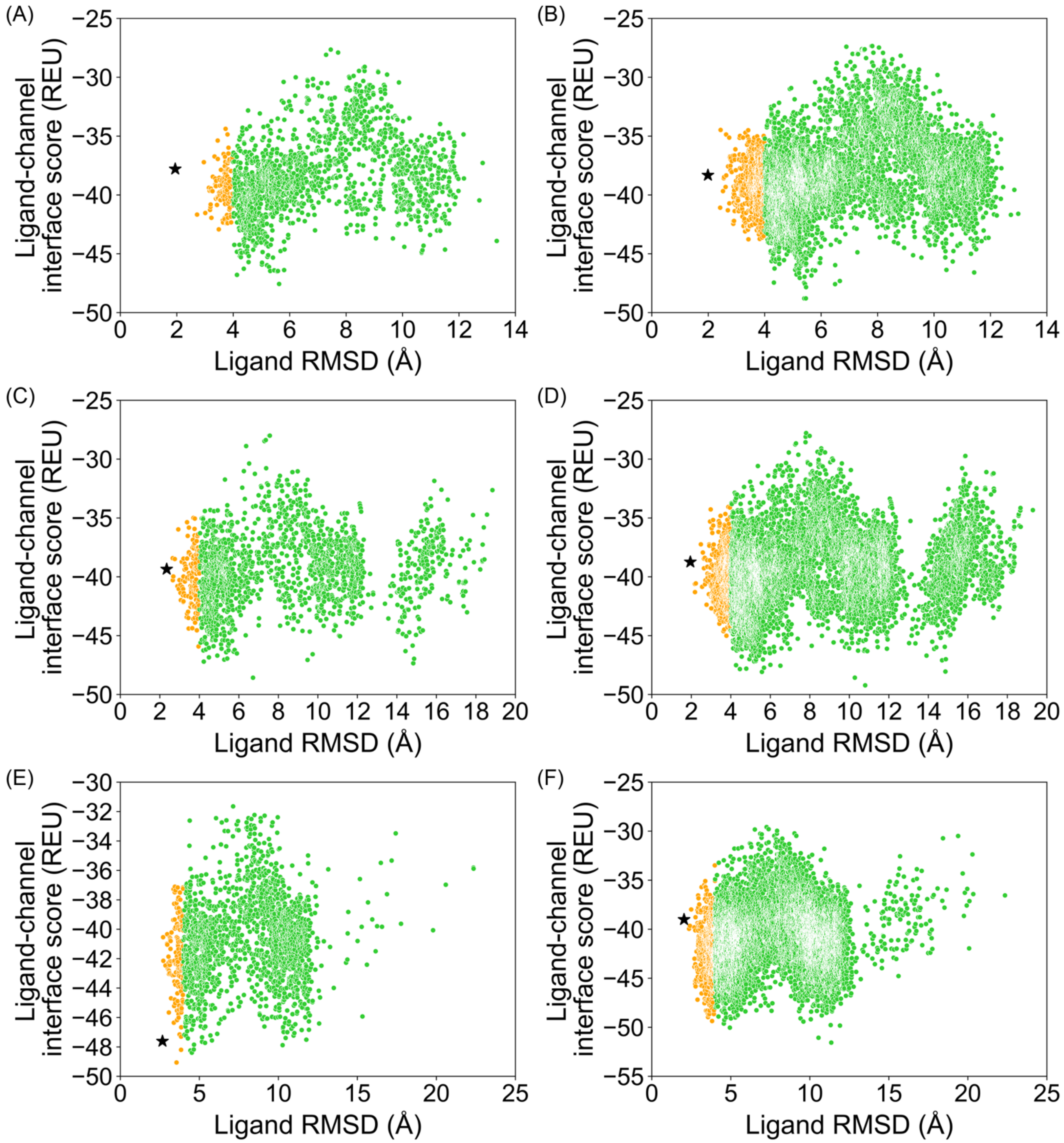


**Figure S7.4** PDB 6JPAV1, GALigandDock, top 10 % of total poses by total energy (`total_score`). **(A)** 20,000 total poses, padding 2 Å, **(B)** 100,000 total poses, padding 2 Å, **(C)** 20,000 total poses, padding 4 Å, **(D)** 100,000 total poses, padding 4 Å, **(E)** 20,000 total poses, padding 7 Å, **(F)** 100,000 total poses, padding 7 Å.

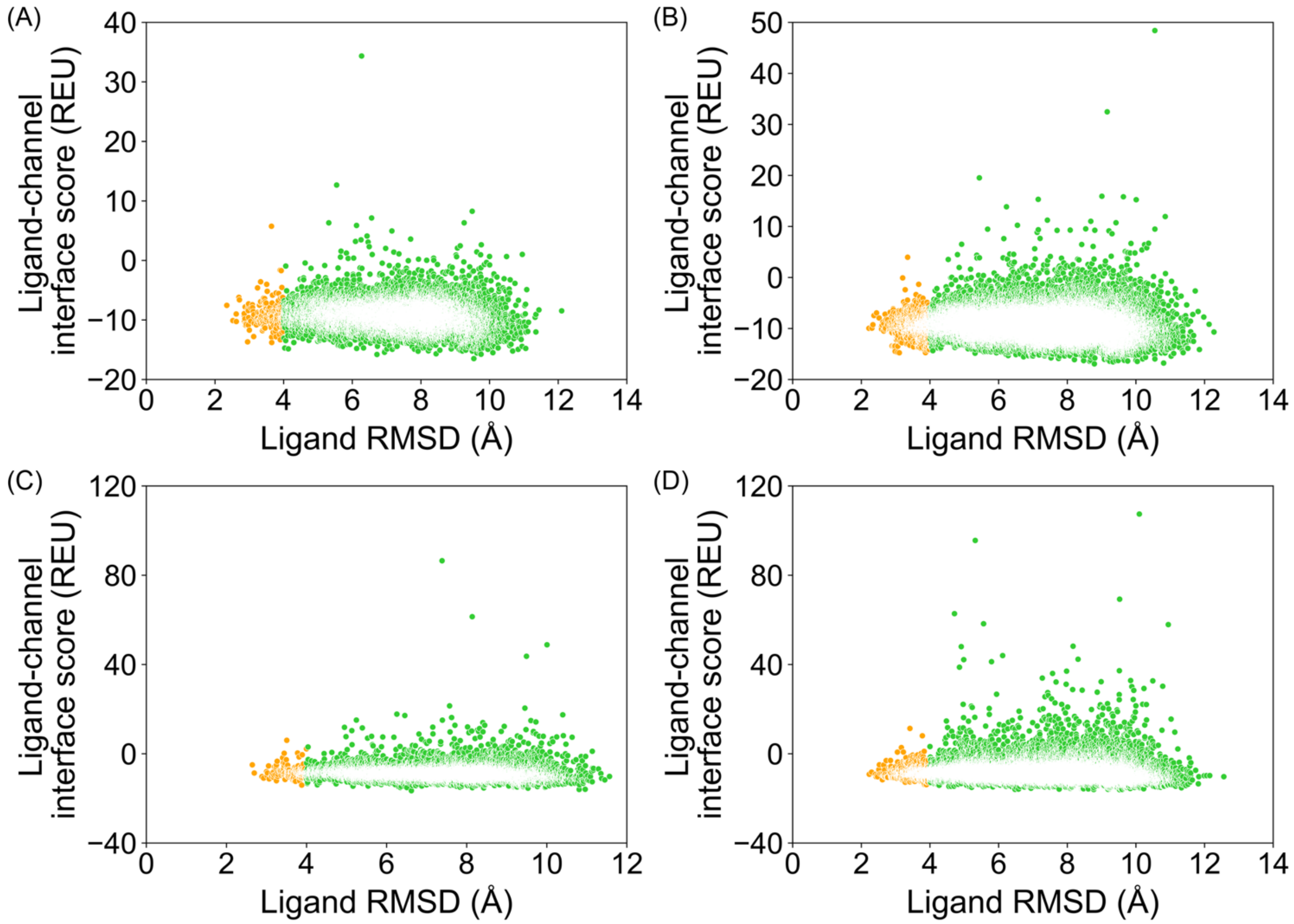


**Figure S8.1** PDB 6JPAV2, RosettaLigand, all poses from docking. **(A)** 20,000 total poses, all ligand atom interface, **(B)** 100,000 total poses, all ligand atom interface, **(C)** 20,000 total poses, ligand neighbor atom interface, **(D)** 100,000 total poses, ligand neighbor atom interface.

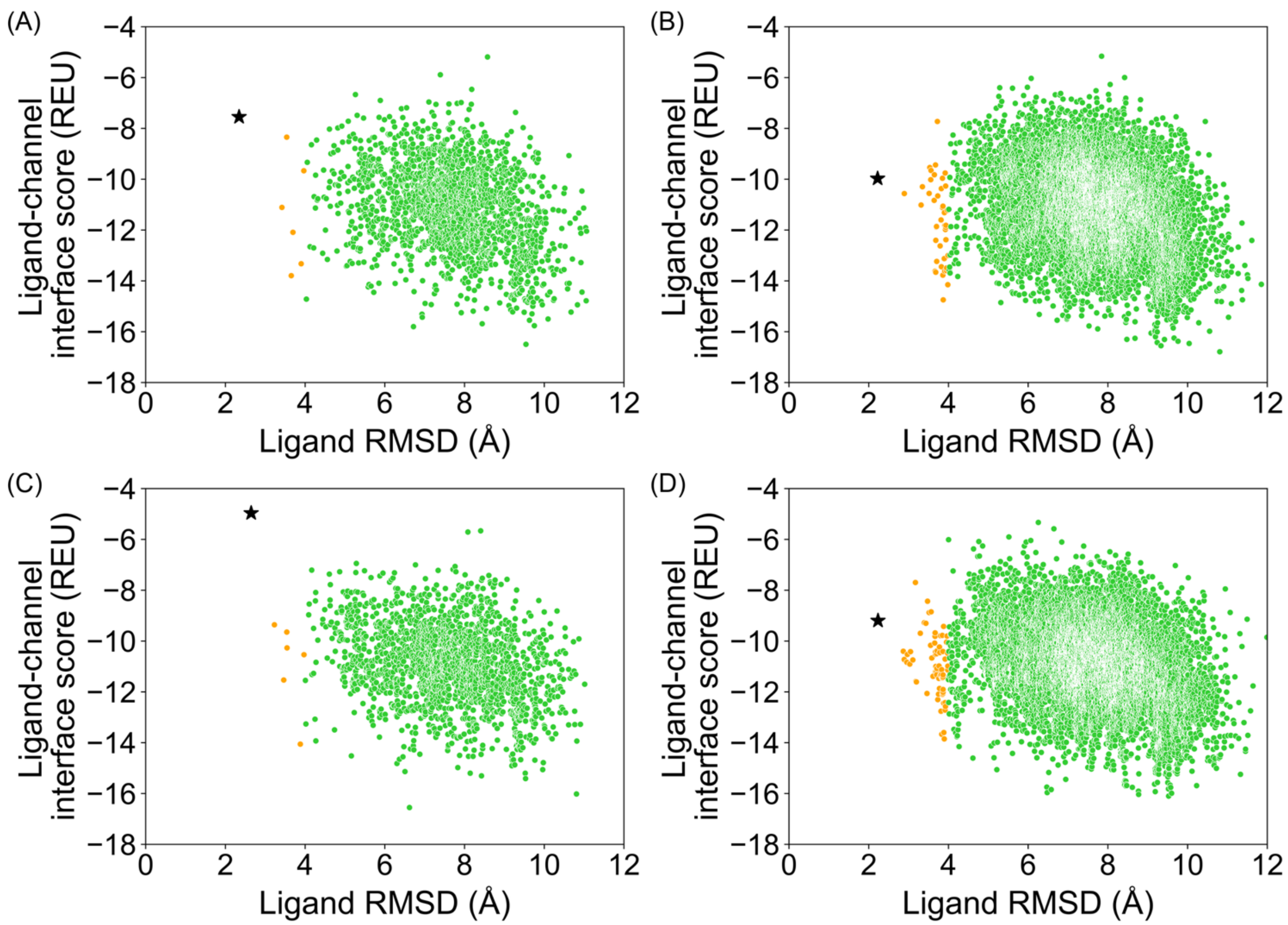


**Figure S8.2** PDB 6JPAV2, RosettaLigand, top 10 % of total poses by total energy (`total_score`). **(A)** 20,000 total poses, all ligand atom interface, **(B)** 100,000 total poses, all ligand atom interface, **(C)** 20,000 total poses, ligand neighbor atom interface, **(D)** 100,000 total poses, ligand neighbor atom interface.

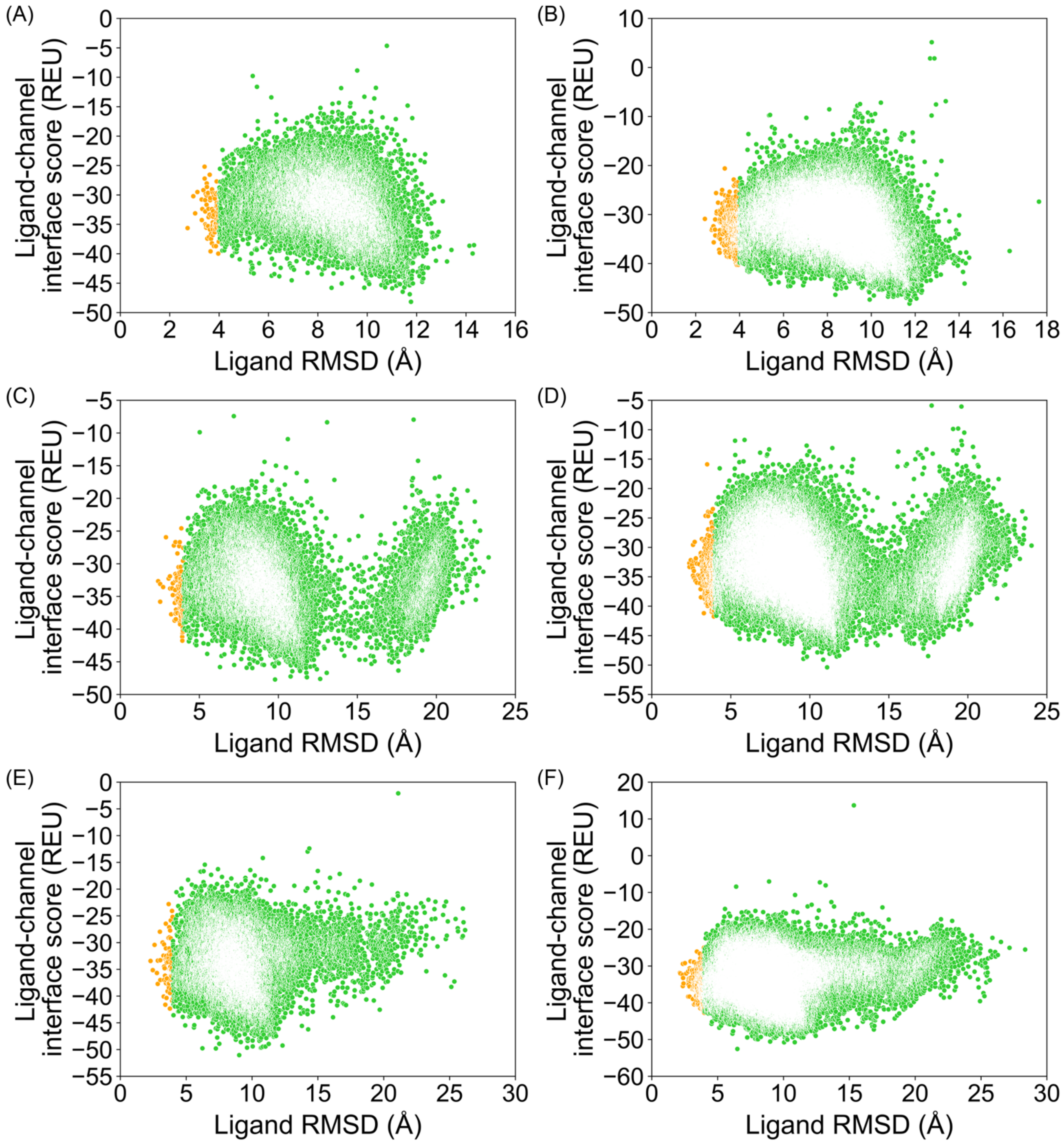


**Figure S8.3** PDB 6JPAV2, GALigandDock, all poses from docking. **(A)** 20,000 total poses, padding 2 Å, **(B)** 100,000 total poses, padding 2 Å, **(C)** 20,000 total poses, padding 4 Å, **(D)** 100,000 total poses, padding 4 Å, **(E)** 20,000 total poses, padding 7 Å, **(F)** 100,000 total poses, padding 7 Å.

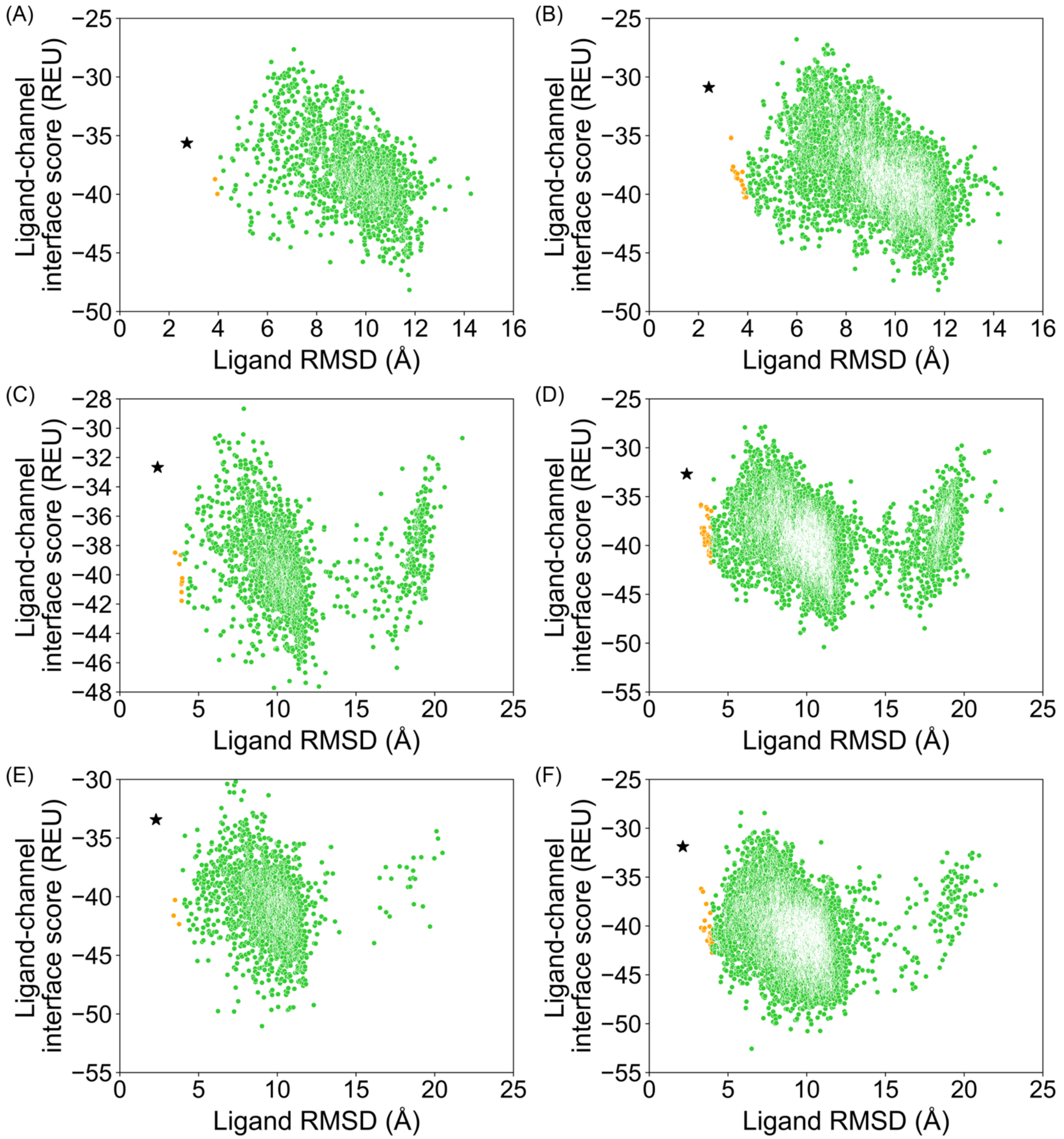


**Figure S8.4** PDB 6JPAV2, GALigandDock, top 10 % of total poses by total energy (`total_score`). **(A)** 20,000 total poses, padding 2 Å, **(B)** 100,000 total poses, padding 2 Å, **(C)** 20,000 total poses, padding 4 Å, **(D)** 100,000 total poses, padding 4 Å, **(E)** 20,000 total poses, padding 7 Å, **(F)** 100,000 total poses, padding 7 Å.

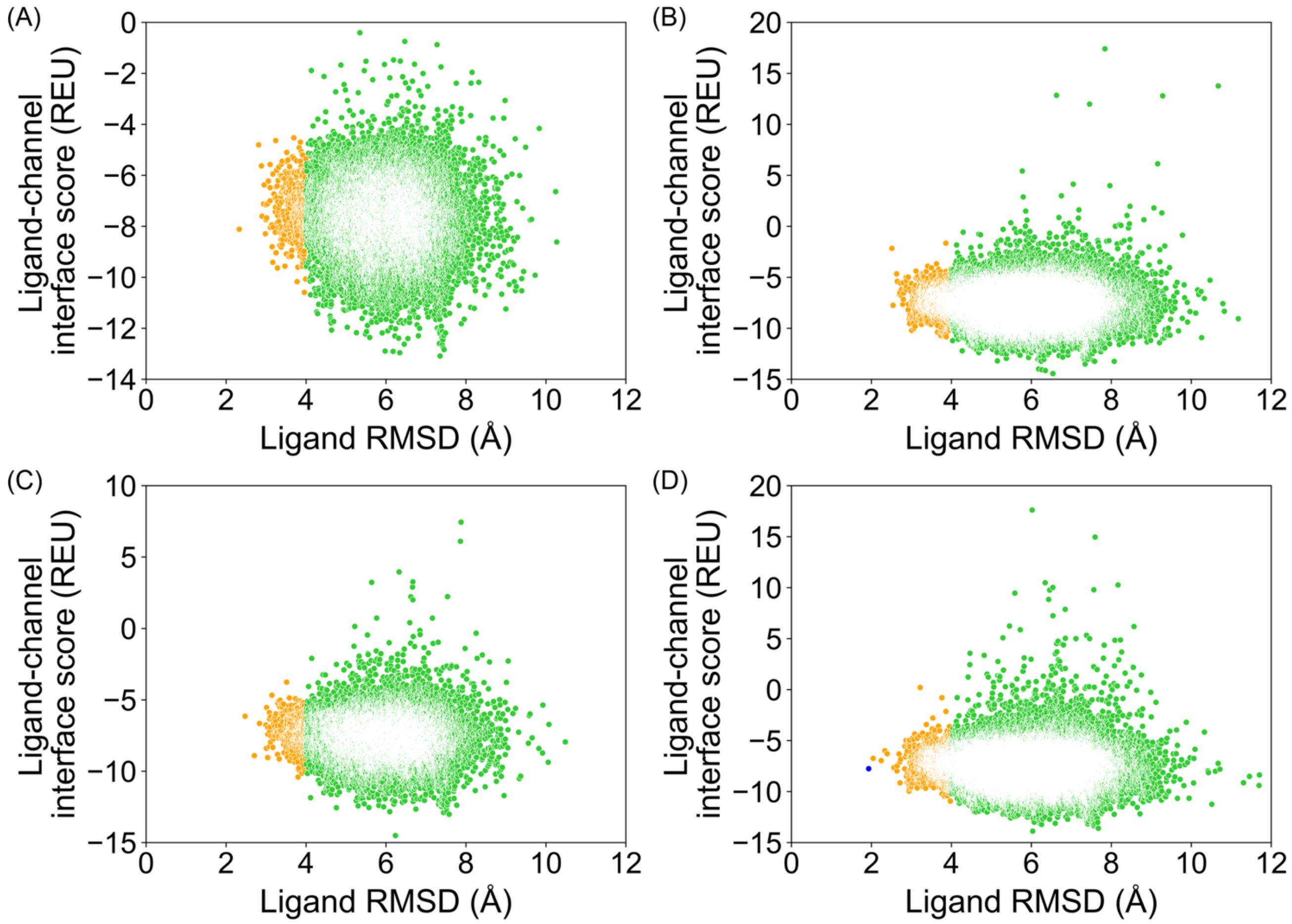


**Figure S9.1** PDB 6JPB, RosettaLigand, all poses from docking. **(A)** 20,000 total poses, all ligand atom interface, **(B)** 100,000 total poses, all ligand atom interface, **(C)** 20,000 total poses, ligand neighbor atom interface, **(D)** 100,000 total poses, ligand neighbor atom interface.

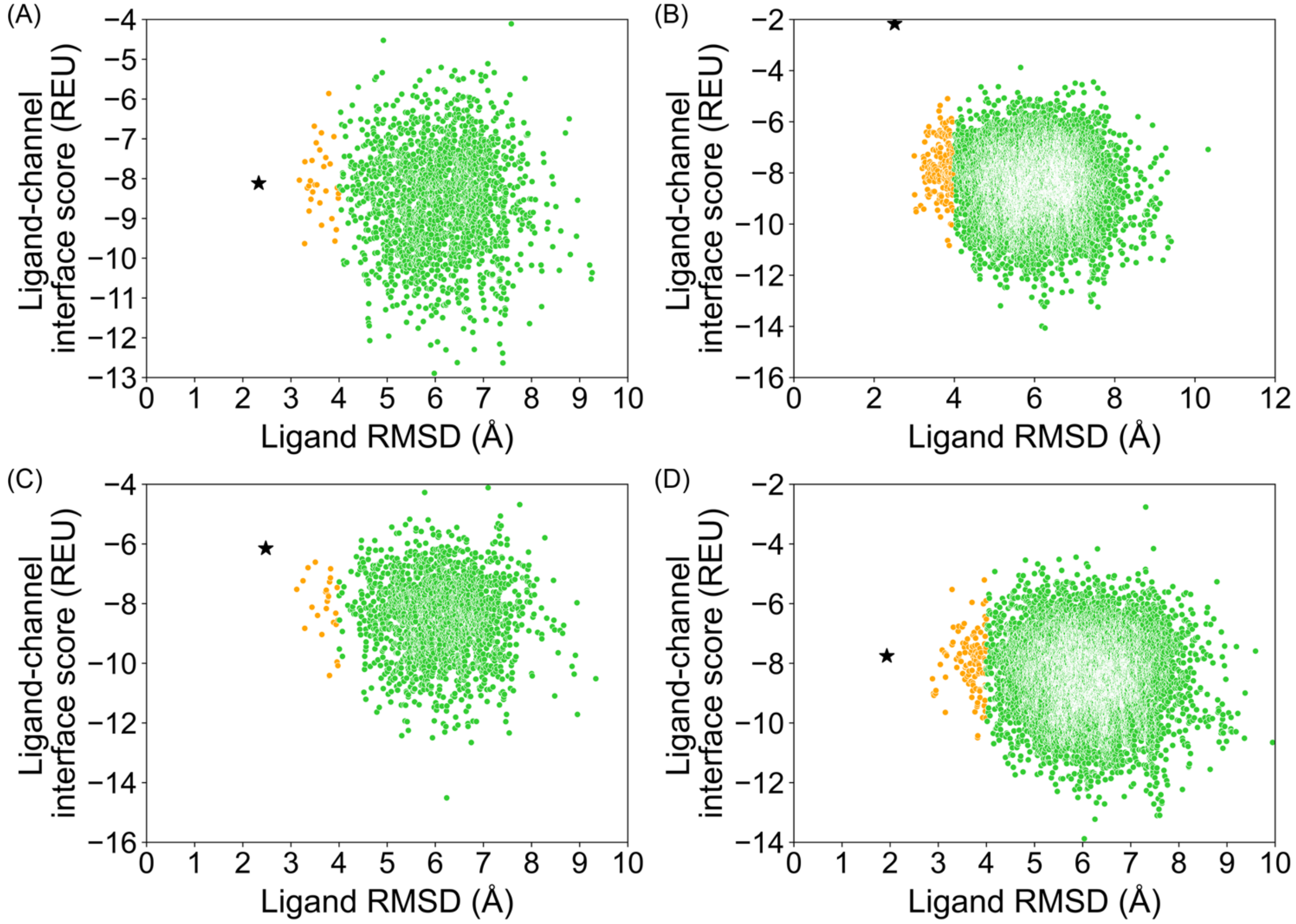


**Figure S9.2** PDB 6JPB, RosettaLigand, top 10 % of total poses by total energy (`total_score`). **(A)** 20,000 total poses, all ligand atom interface, **(B)** 100,000 total poses, all ligand atom interface, **(C)** 20,000 total poses, ligand neighbor atom interface, **(D)** 100,000 total poses, ligand neighbor atom interface.

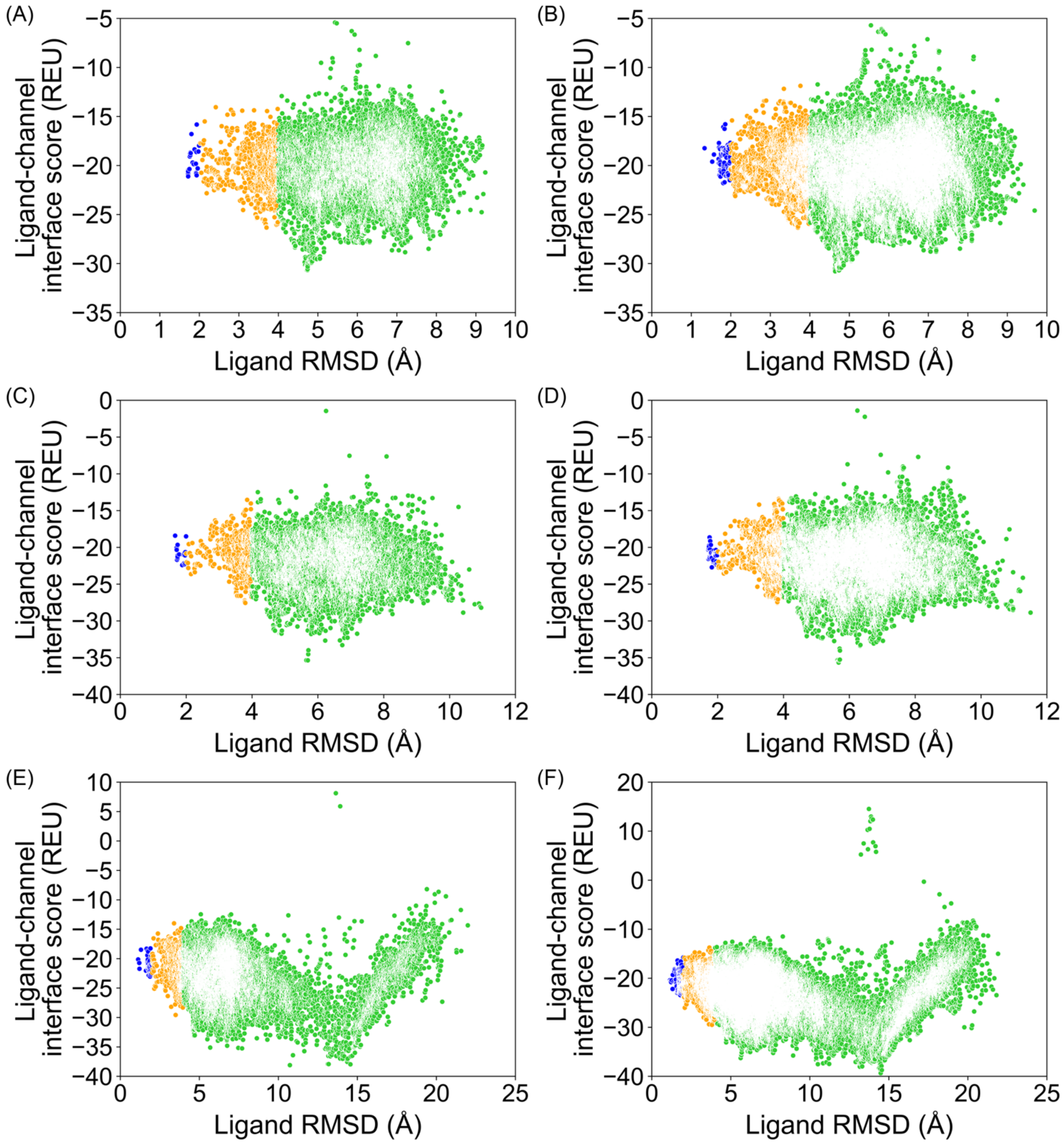


**Figure S9.3** PDB 6JPB, GALigandDock, all poses from docking. **(A)** 20,000 total poses, padding 2 Å, **(B)** 100,000 total poses, padding 2 Å, **(C)** 20,000 total poses, padding 4 Å, **(D)** 100,000 total poses, padding 4 Å, **(E)** 20,000 total poses, padding 7 Å, **(F)** 100,000 total poses, padding 7 Å.

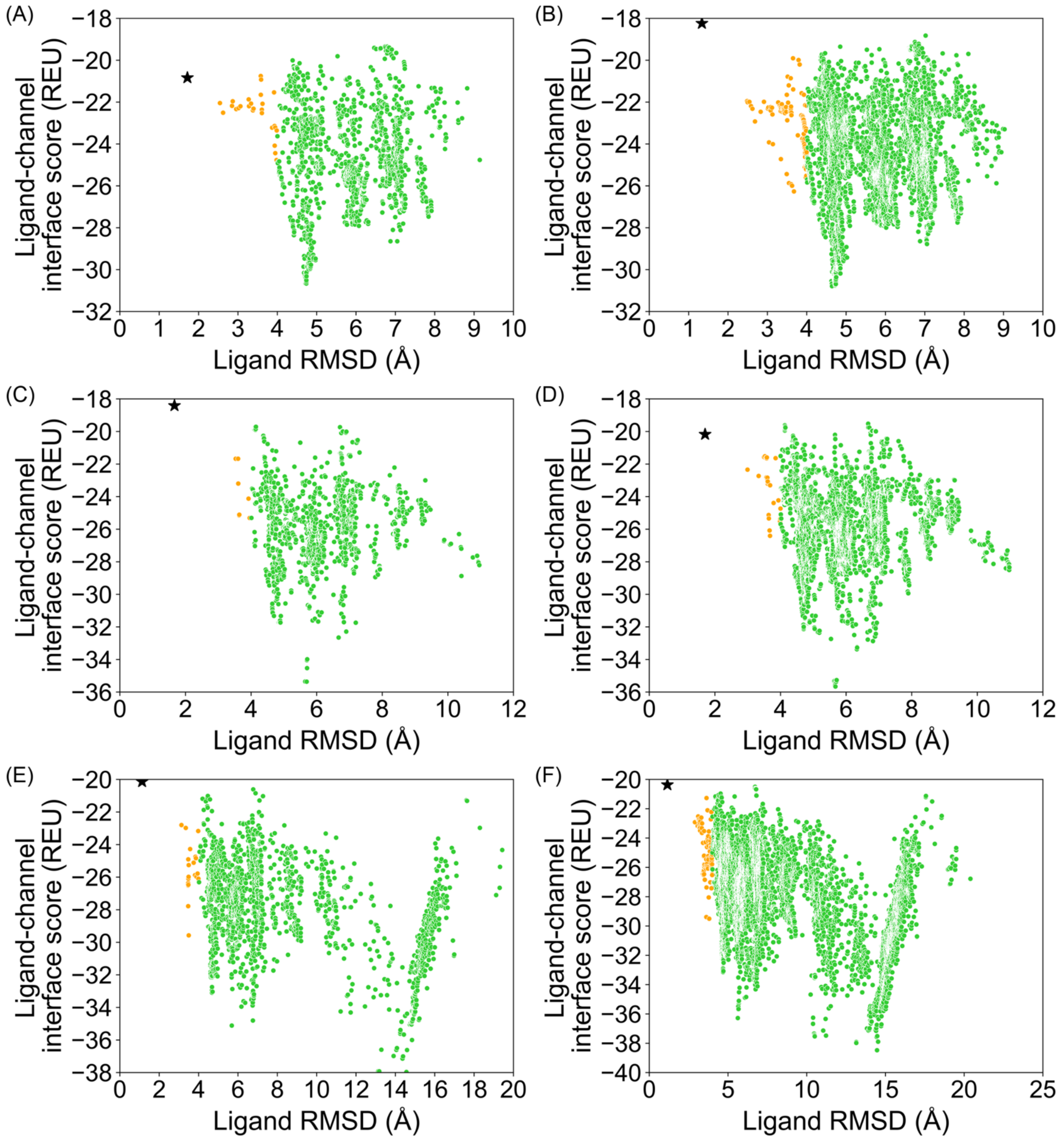


**Figure S9.4** PDB 6JPB, GALigandDock, top 10 % of total poses by total energy (`total_score`). **(A)** 20,000 total poses, padding 2 Å, **(B)** 100,000 total poses, padding 2 Å, **(C)** 20,000 total poses, padding 4 Å, **(D)** 100,000 total poses, padding 4 Å, **(E)** 20,000 total poses, padding 7 Å, **(F)** 100,000 total poses, padding 7 Å.

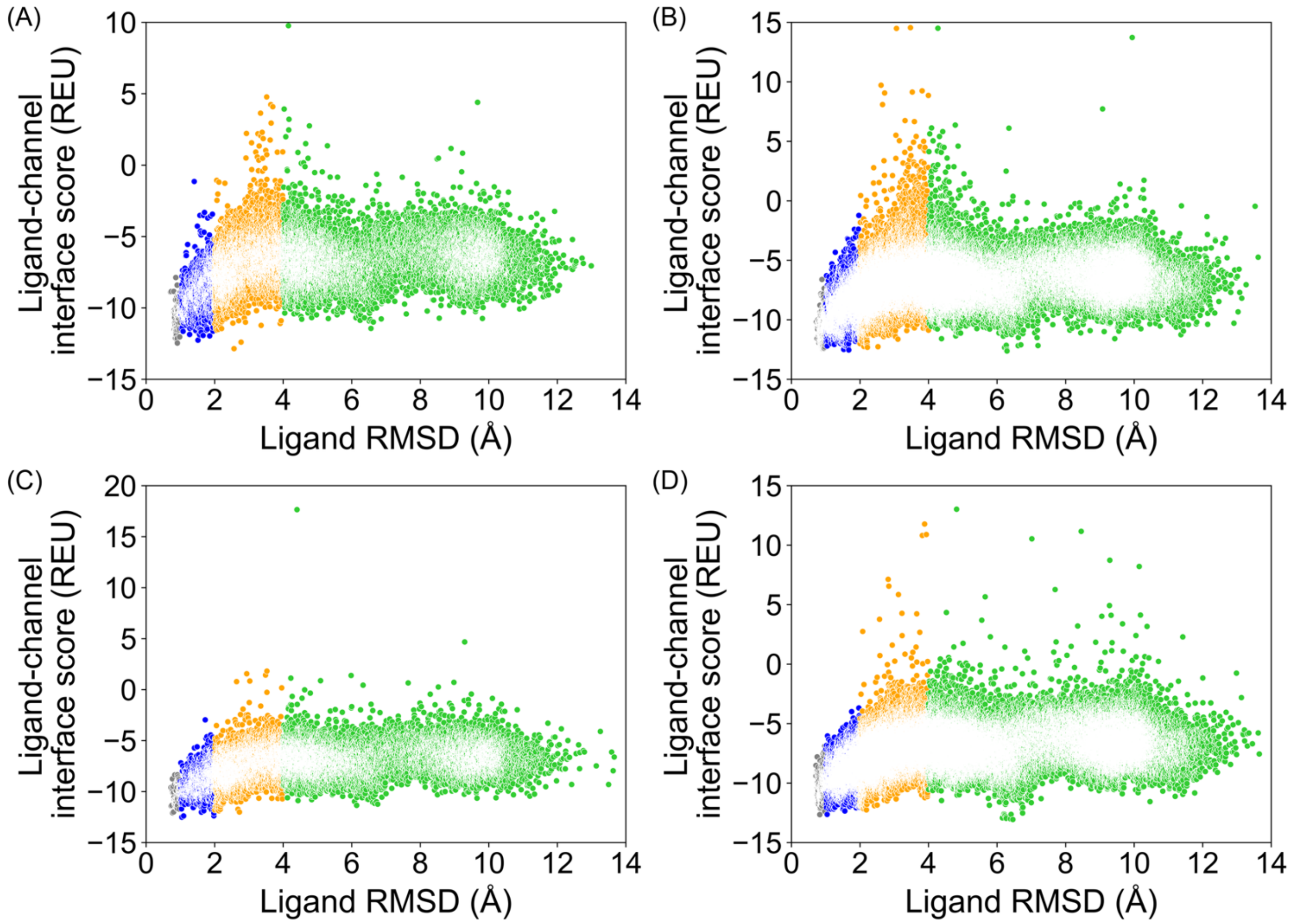


**Figure S10.1** PDB 6KZP, RosettaLigand, all poses from docking. **(A)** 20,000 total poses, all ligand atom interface, **(B)** 100,000 total poses, all ligand atom interface, **(C)** 20,000 total poses, ligand neighbor atom interface, **(D)** 100,000 total poses, ligand neighbor atom interface.

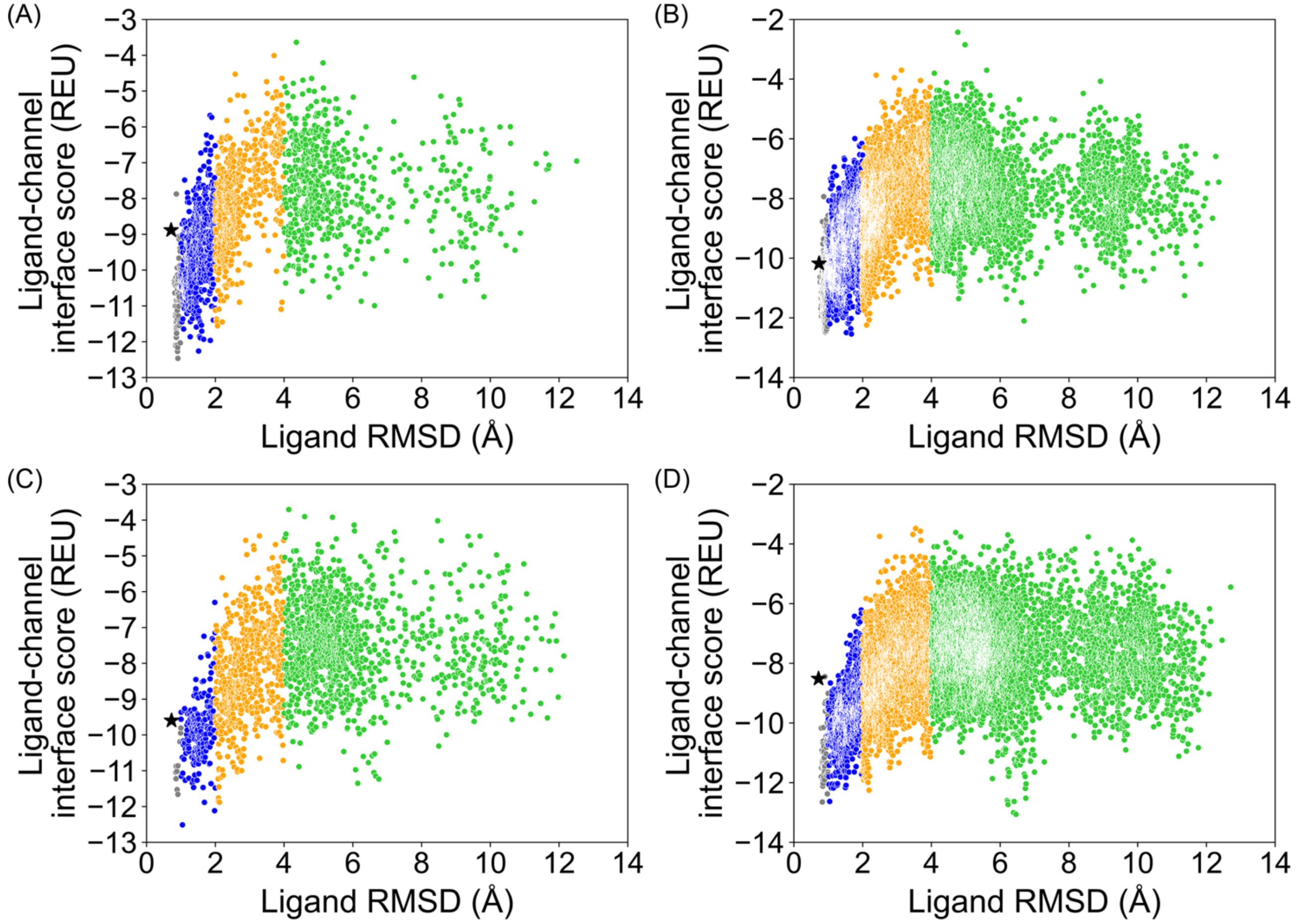


**Figure S10.2** PDB 6KZP, RosettaLigand, top 10 % of total poses by total energy (`total_score`). **(A)** 20,000 total poses, all ligand atom interface, **(B)** 100,000 total poses, all ligand atom interface, **(C)** 20,000 total poses, ligand neighbor atom interface, **(D)** 100,000 total poses, ligand neighbor atom interface.

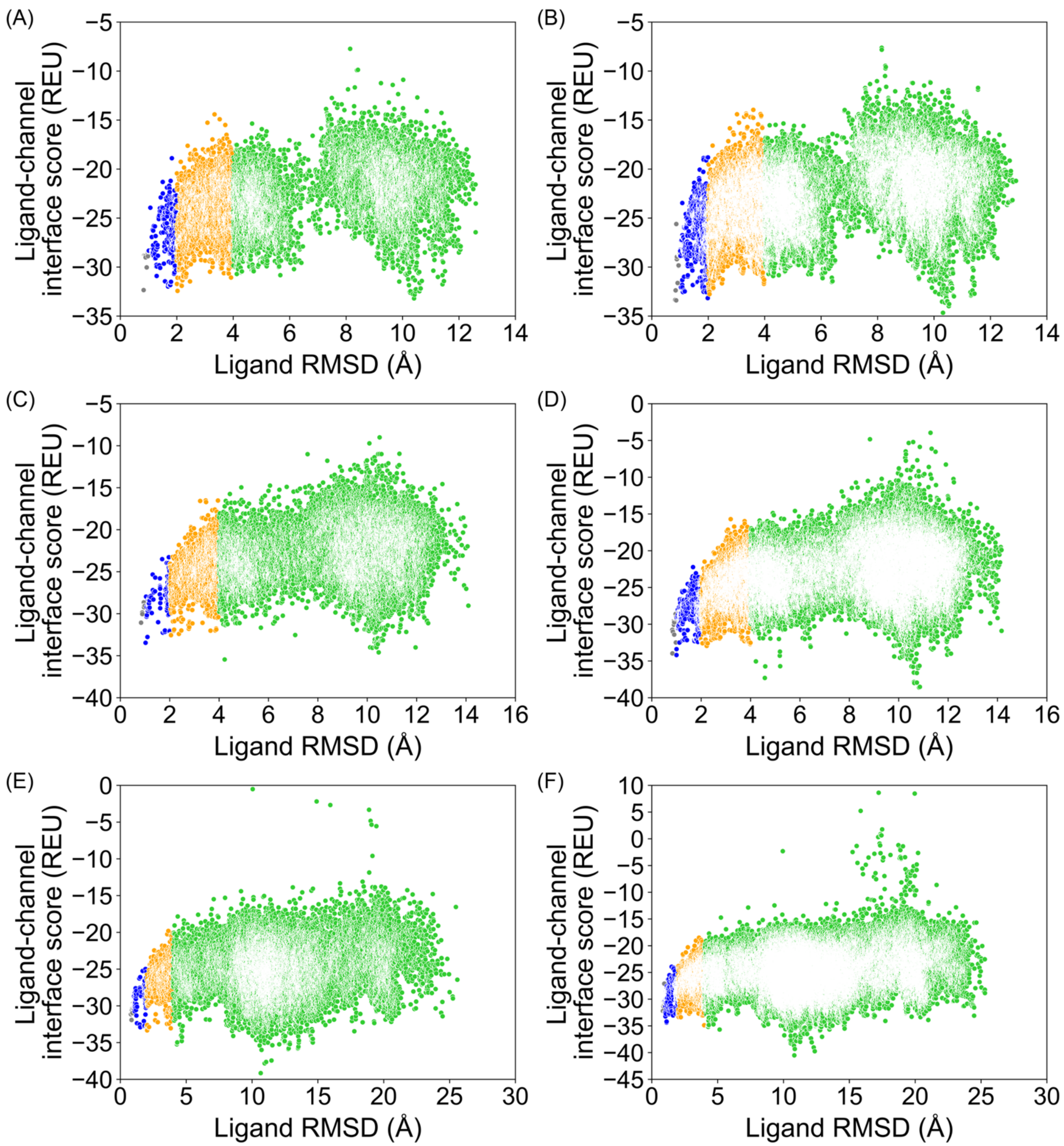


**Figure S10.3** PDB 6KZP, GALigandDock, all poses from docking. **(A)** 20,000 total poses, padding 2 Å, **(B)** 100,000 total poses, padding 2 Å, **(C)** 20,000 total poses, padding 4 Å, **(D)** 100,000 total poses, padding 4 Å, **(E)** 20,000 total poses, padding 7 Å, **(F)** 100,000 total poses, padding 7 Å.

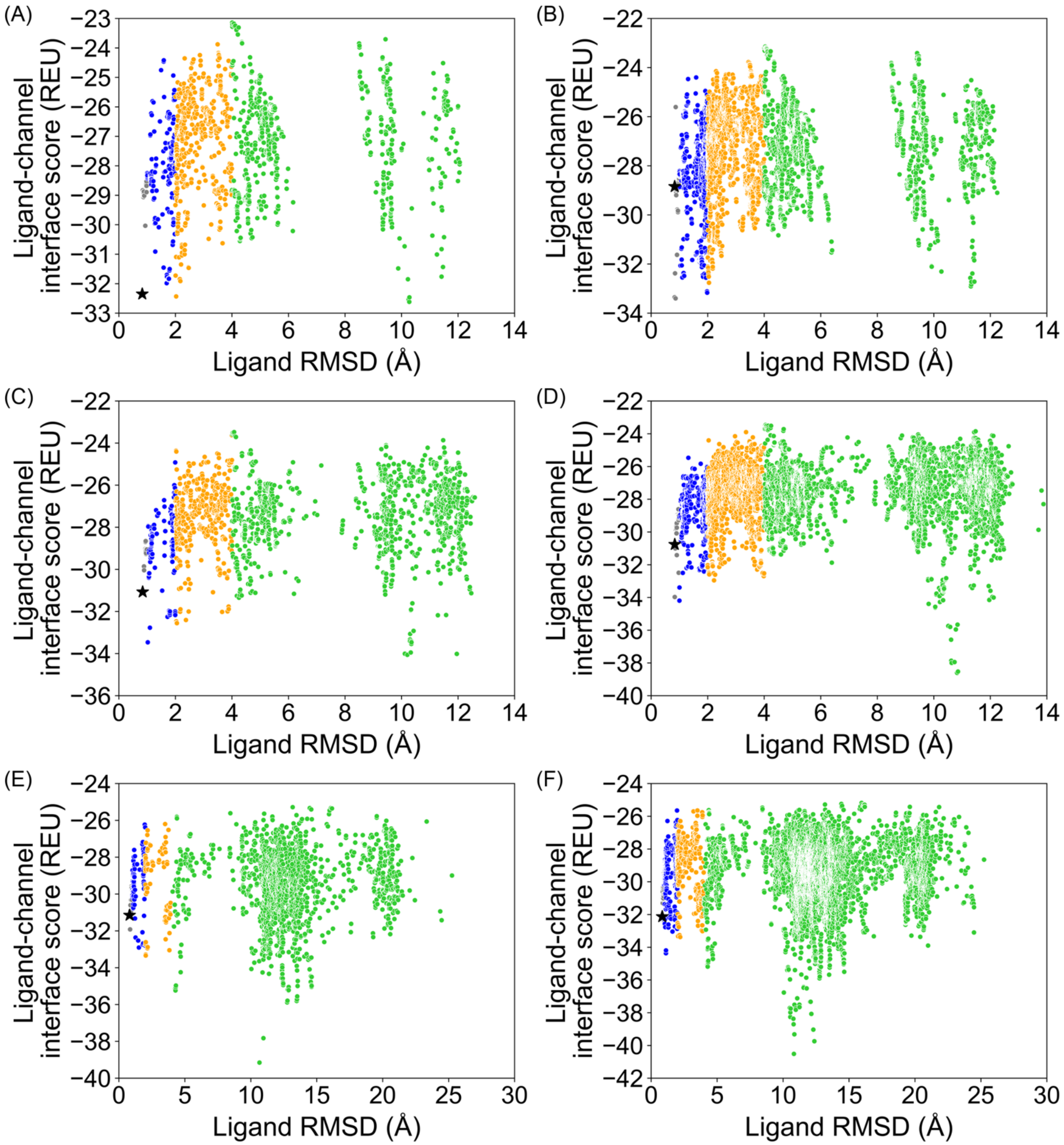


**Figure S10.4** PDB 6KZP, GALigandDock, top 10 % of total poses by total energy (`total_score`). **(A)** 20,000 total poses, padding 2 Å, **(B)** 100,000 total poses, padding 2 Å, **(C)** 20,000 total poses, padding 4 Å, **(D)** 100,000 total poses, padding 4 Å, **(E)** 20,000 total poses, padding 7 Å, **(F)** 100,000 total poses, padding 7 Å.

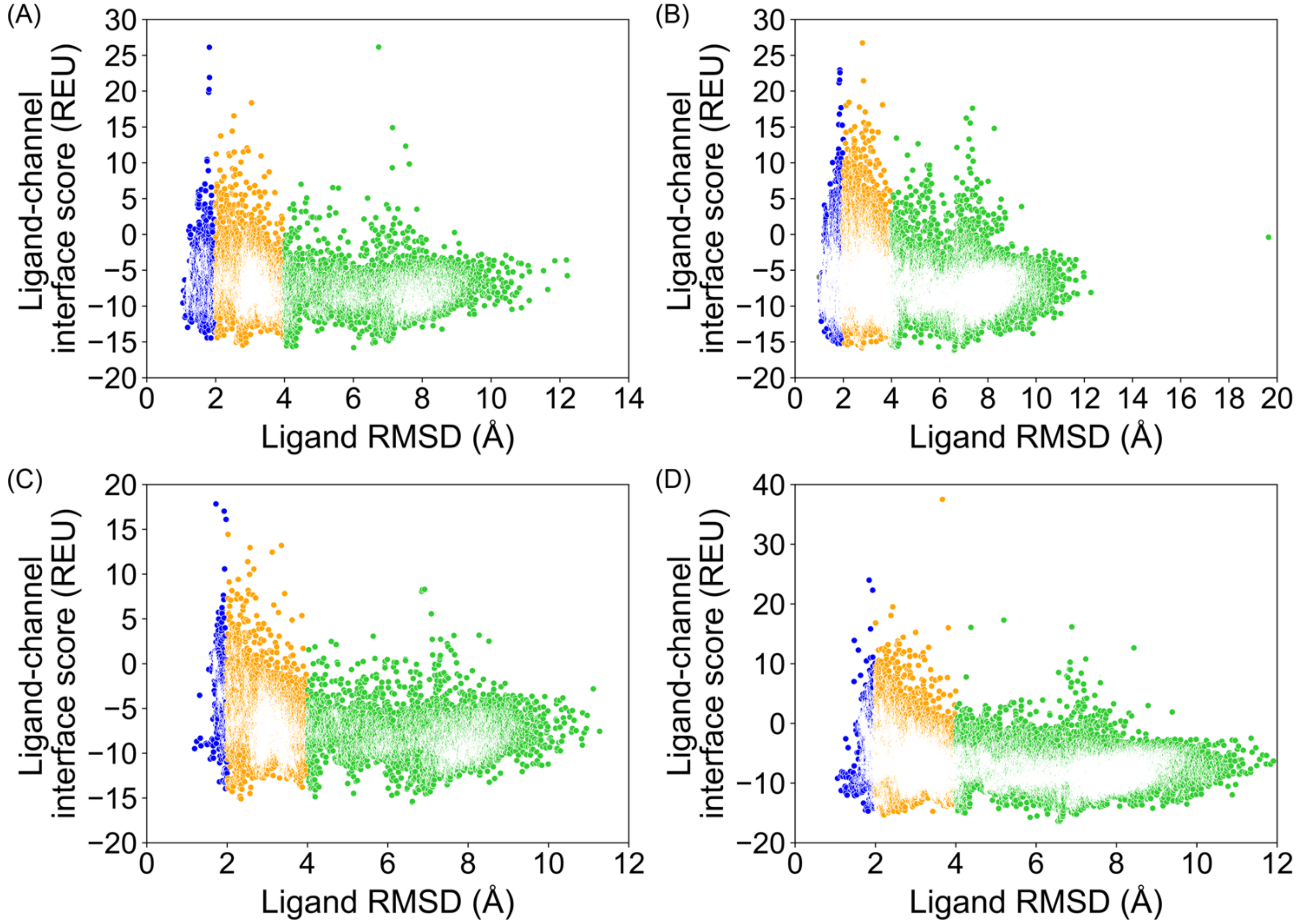


**Figure S11.1** PDB 6U88, RosettaLigand, all poses from docking. **(A)** 20,000 total poses, all ligand atom interface, **(B)** 100,000 total poses, all ligand atom interface, **(C)** 20,000 total poses, ligand neighbor atom interface, **(D)** 100,000 total poses, ligand neighbor atom interface.

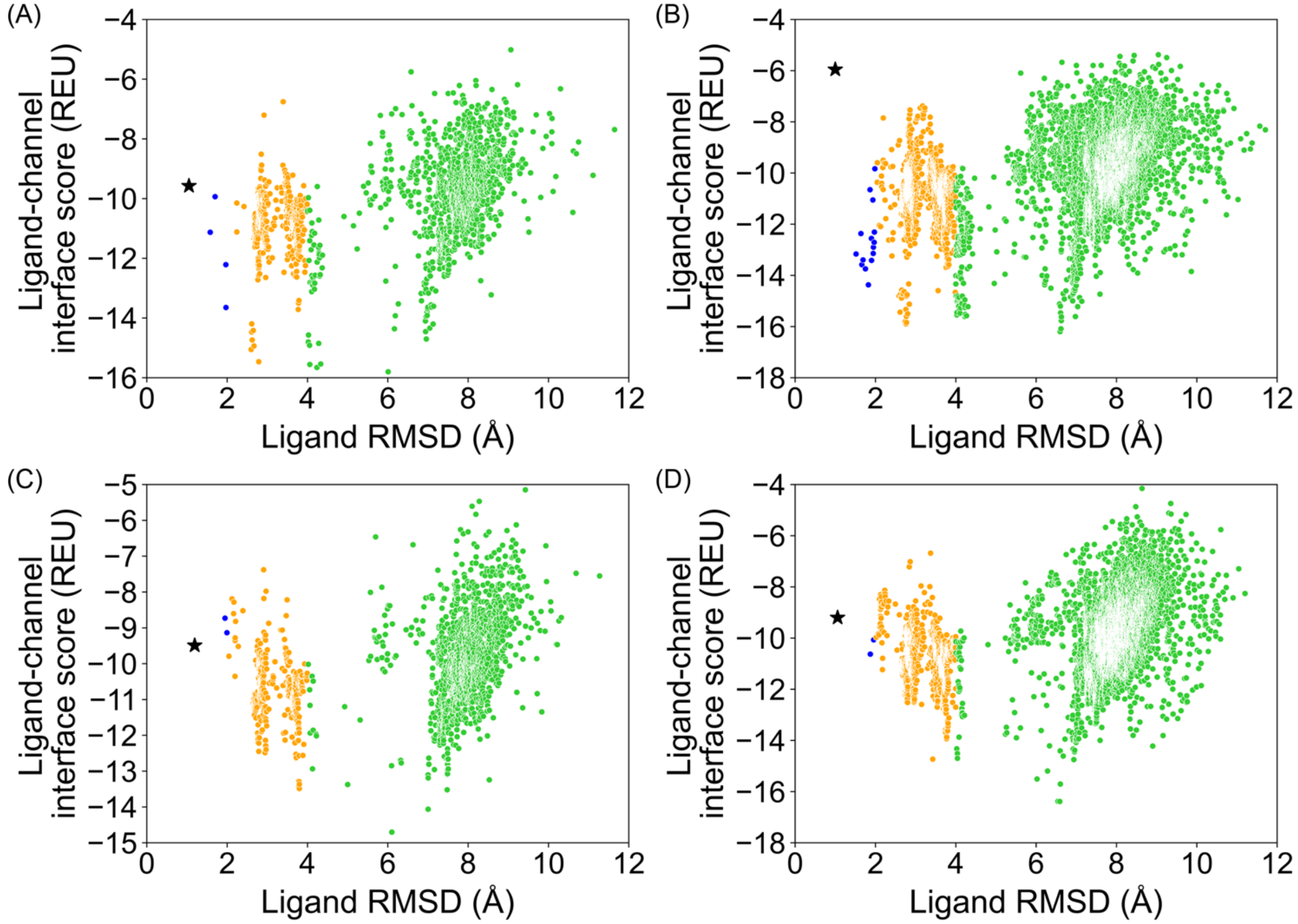


**Figure S11.2** PDB 6U88, RosettaLigand, top 10 % of total poses by total energy (`total_score`). **(A)** 20,000 total poses, all ligand atom interface, **(B)** 100,000 total poses, all ligand atom interface, **(C)** 20,000 total poses, ligand neighbor atom interface, **(D)** 100,000 total poses, ligand neighbor atom interface.

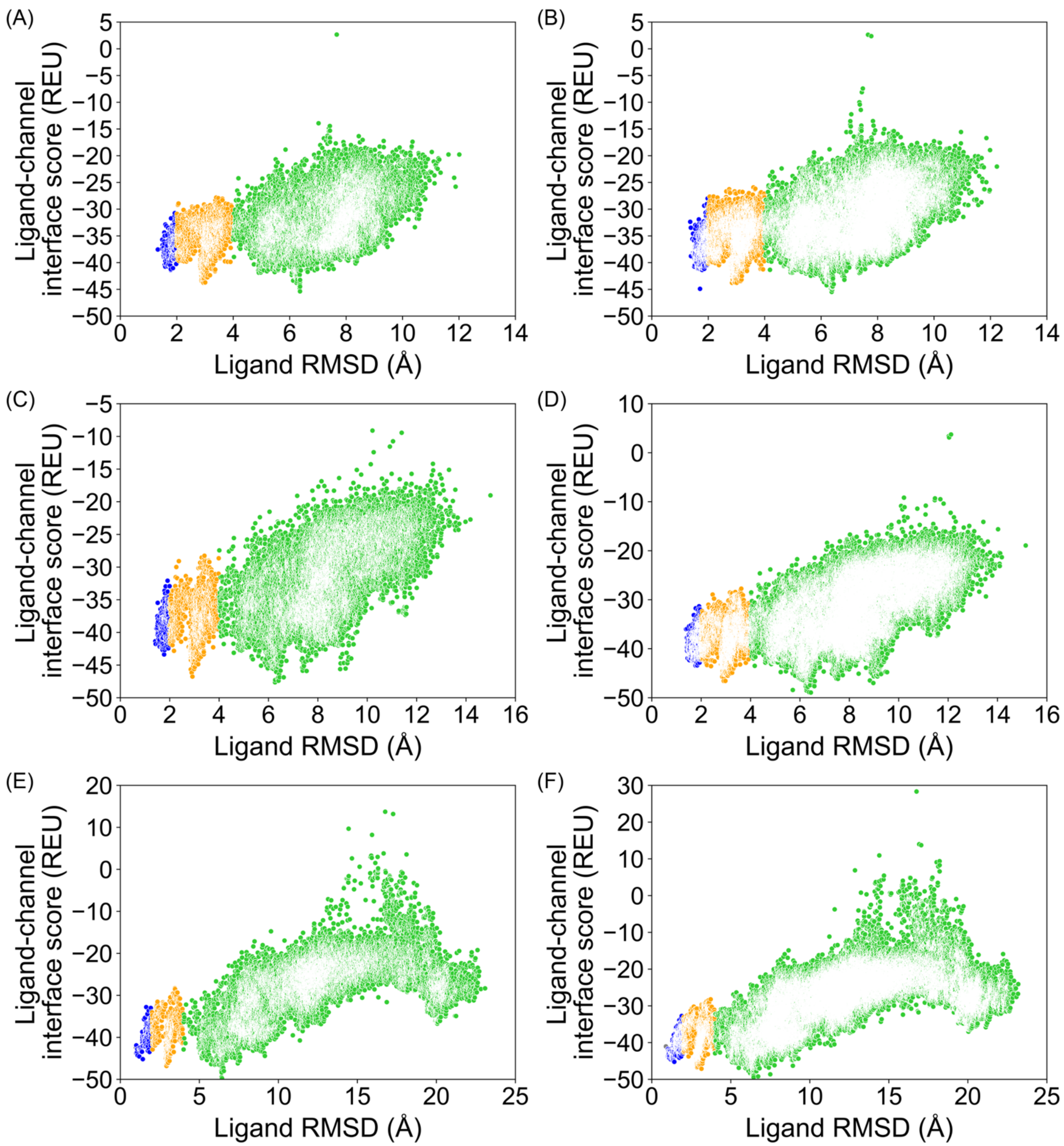


**Figure S11.3** PDB 6U88, GALigandDock, all poses from docking. **(A)** 20,000 total poses, padding 2 Å, **(B)** 100,000 total poses, padding 2 Å, **(C)** 20,000 total poses, padding 4 Å, **(D)** 100,000 total poses, padding 4 Å, **(E)** 20,000 total poses, padding 7 Å, **(F)** 100,000 total poses, padding 7 Å.

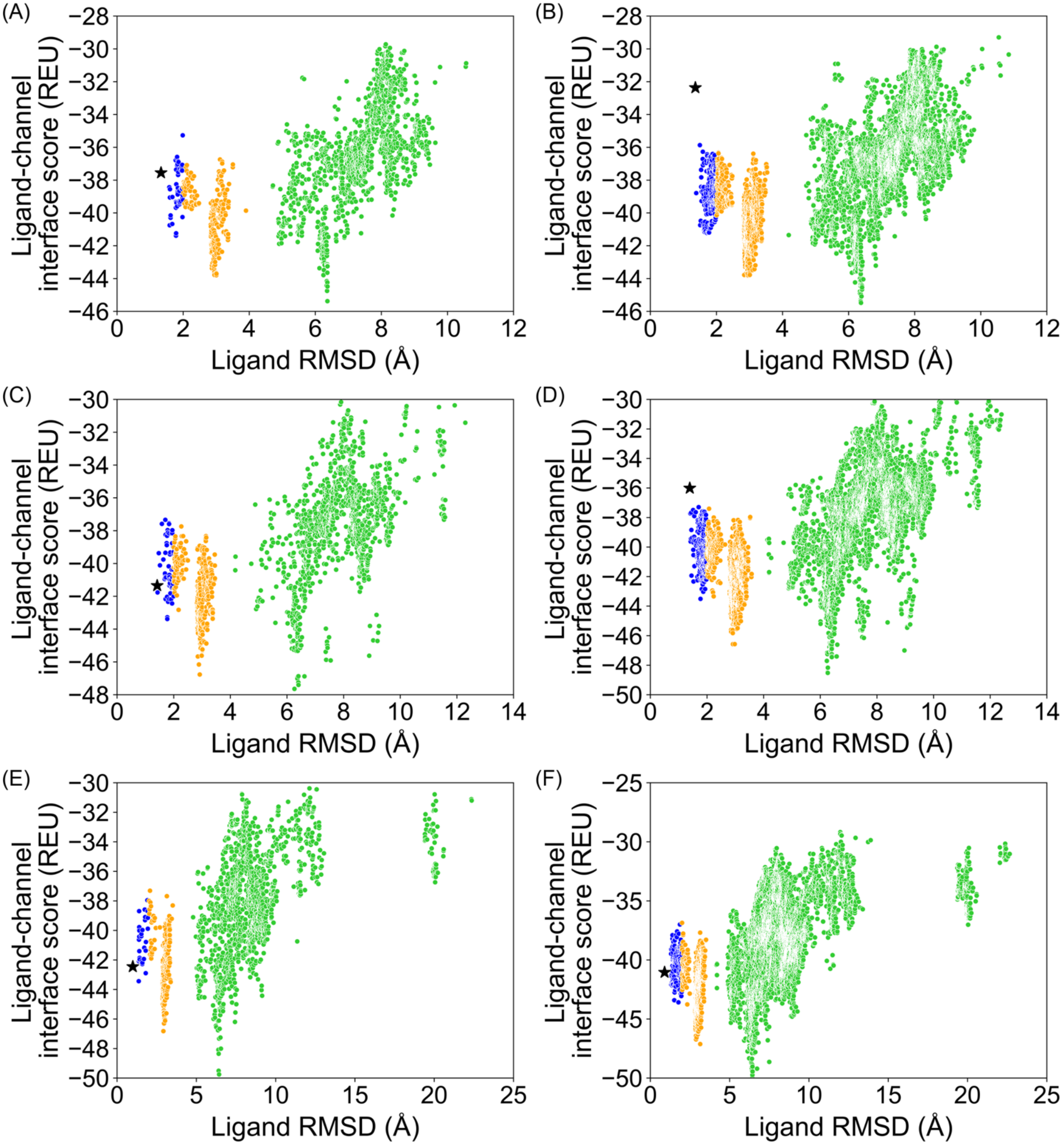


**Figure S11.4** PDB 6U88, GALigandDock, top 10 % of total poses by total energy (`total_score`). **(A)** 20,000 total poses, padding 2 Å, **(B)** 100,000 total poses, padding 2 Å, **(C)** 20,000 total poses, padding 4 Å, **(D)** 100,000 total poses, padding 4 Å, **(E)** 20,000 total poses, padding 7 Å, **(F)** 100,000 total poses, padding 7 Å.

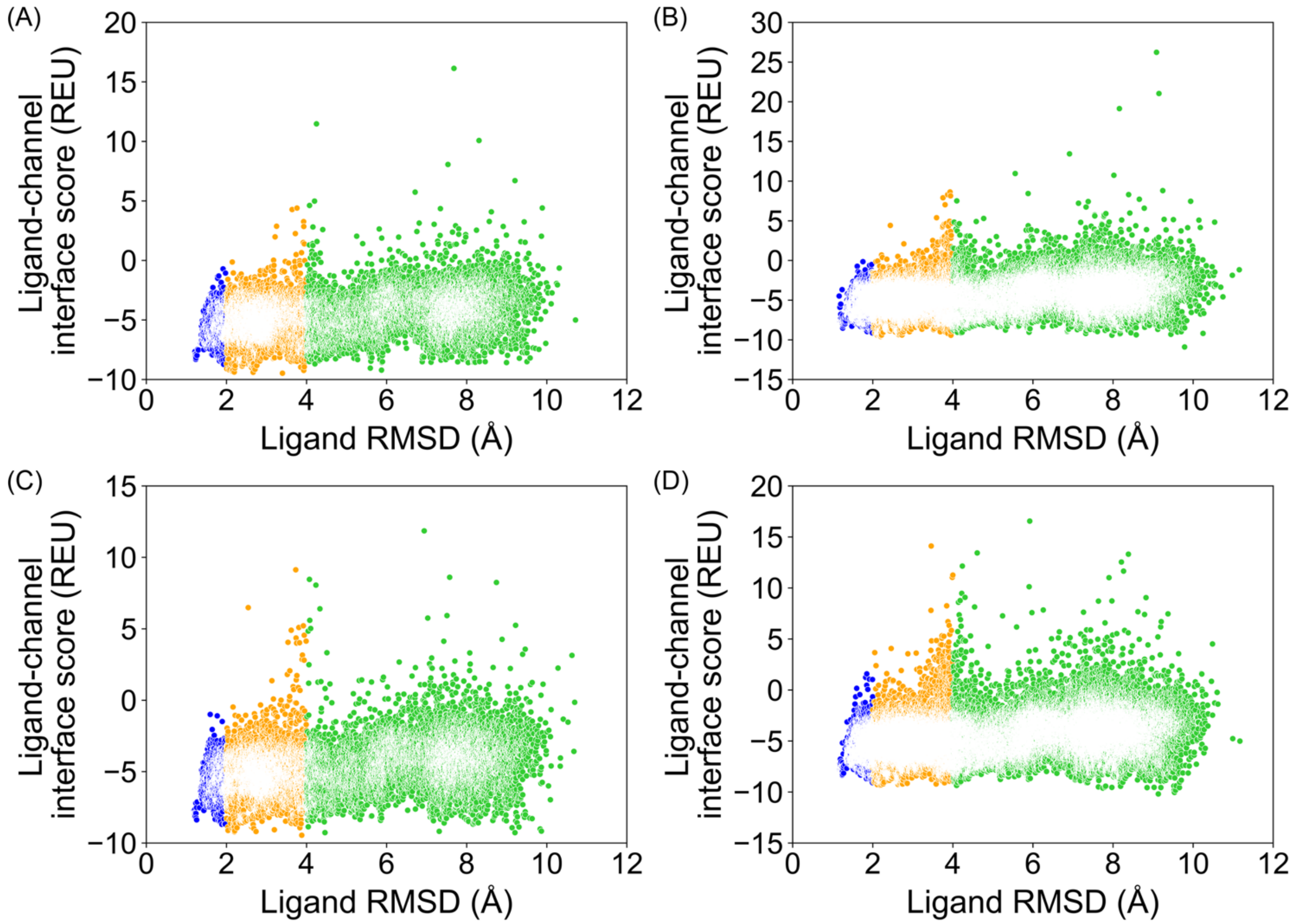


**Figure S12.1** PDB 6UZ0, RosettaLigand, all poses from docking. **(A)** 20,000 total poses, all ligand atom interface, **(B)** 100,000 total poses, all ligand atom interface, **(C)** 20,000 total poses, ligand neighbor atom interface, **(D)** 100,000 total poses, ligand neighbor atom interface.

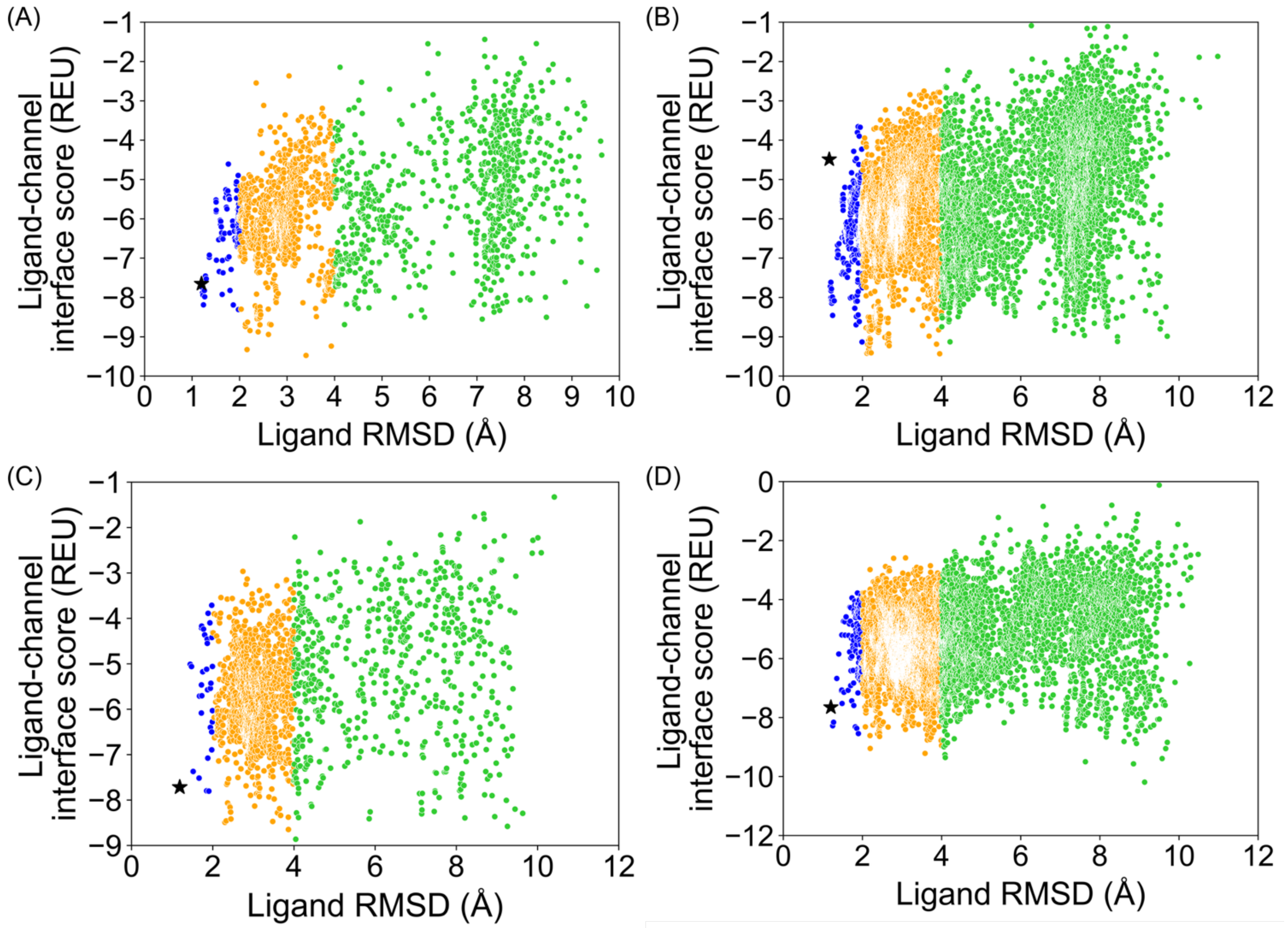


**Figure S12.2** PDB 6UZ0, RosettaLigand, top 10 % of total poses by total energy (`total_score`). **(A)** 20,000 total poses, all ligand atom interface, **(B)** 100,000 total poses, all ligand atom interface, **(C)** 20,000 total poses, ligand neighbor atom interface, **(D)** 100,000 total poses, ligand neighbor atom interface.

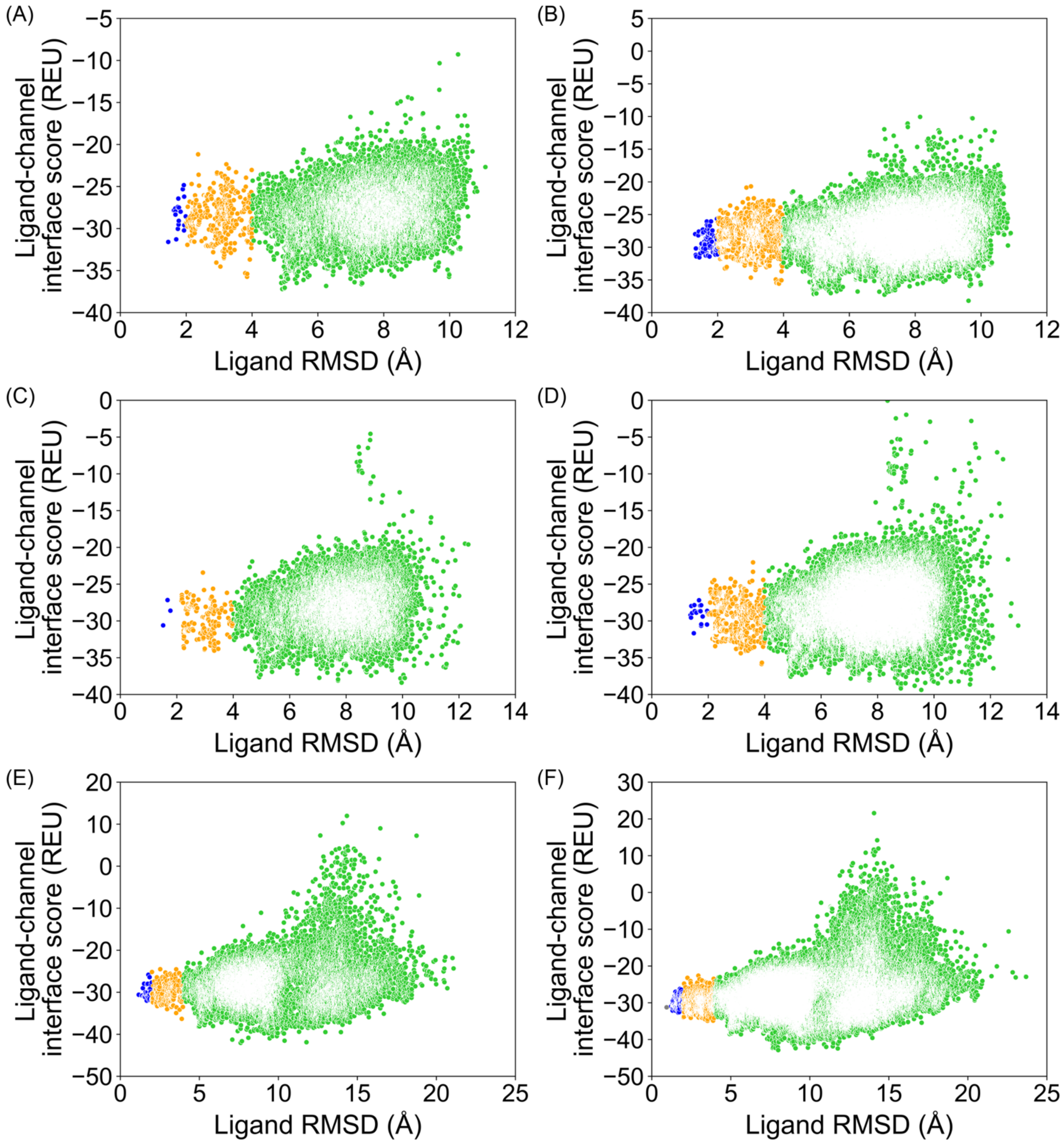


**Figure S12.3** PDB 6UZ0, GALigandDock, all poses from docking. **(A)** 20,000 total poses, padding 2 Å, **(B)** 100,000 total poses, padding 2 Å, **(C)** 20,000 total poses, padding 4 Å, **(D)** 100,000 total poses, padding 4 Å, **(E)** 20,000 total poses, padding 7 Å, **(F)** 100,000 total poses, padding 7 Å.

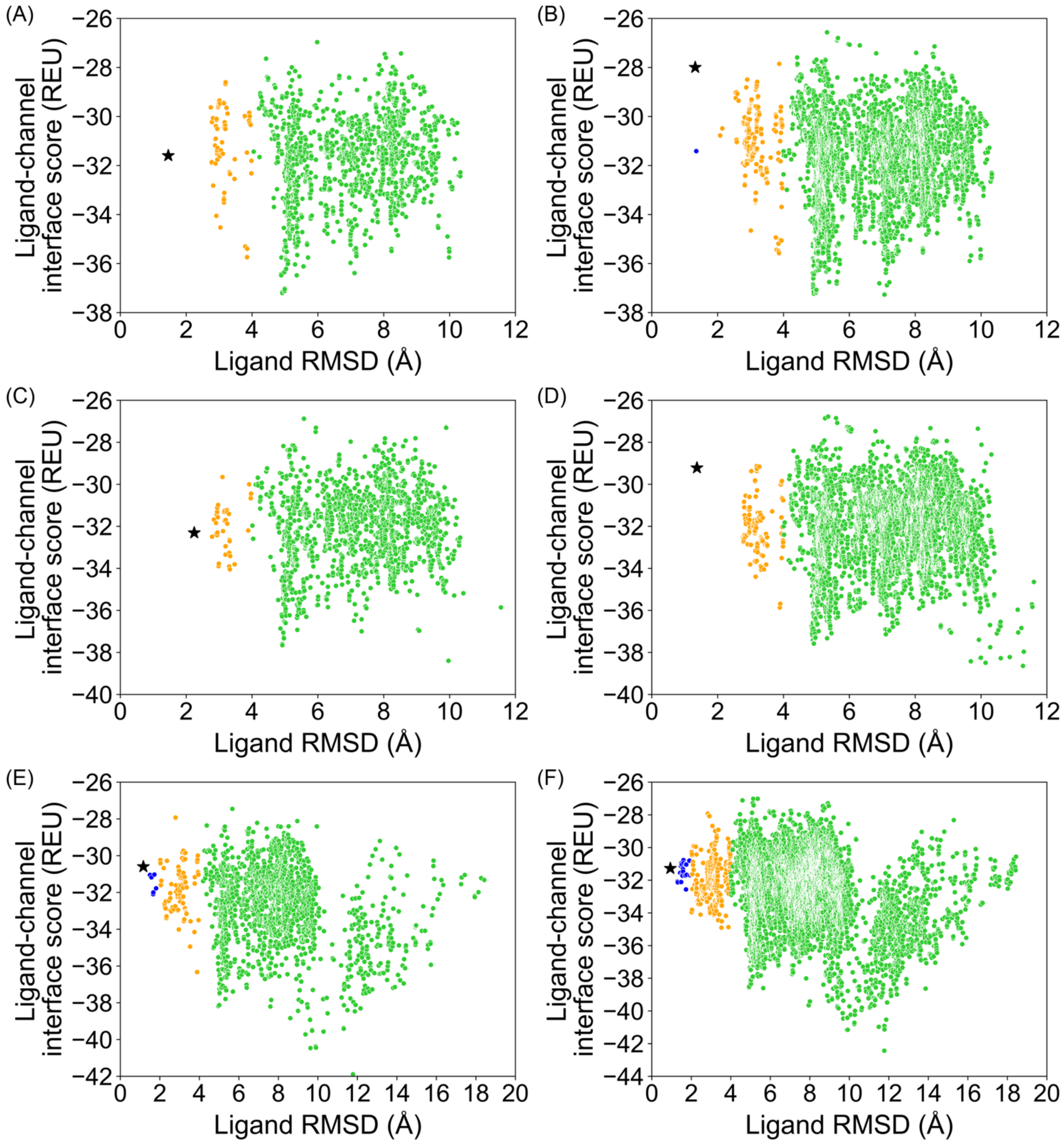


**Figure S12.4** PDB 6UZ0, GALigandDock, top 10 % of total poses by total energy (`total_score`). **(A)** 20,000 total poses, padding 2 Å, **(B)** 100,000 total poses, padding 2 Å, **(C)** 20,000 total poses, padding 4 Å, **(D)** 100,000 total poses, padding 4 Å, **(E)** 20,000 total poses, padding 7 Å, **(F)** 100,000 total poses, padding 7 Å.

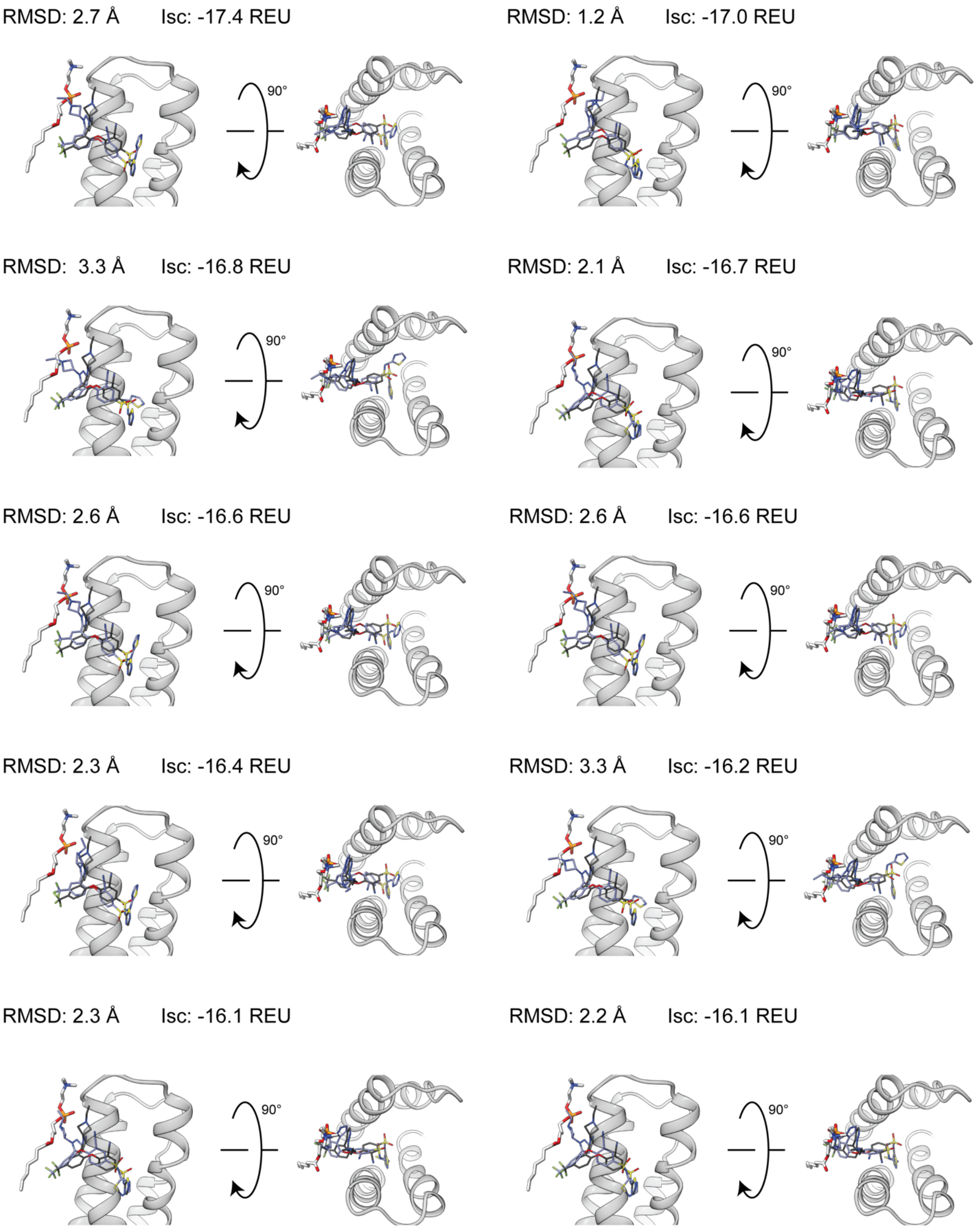


**Figure S13.1.** PDB 5EK0. Top 10 interface-scoring (`interface_delta_X`) poses from RosettaLigand, 100,000 total poses, all atom ligand interface.

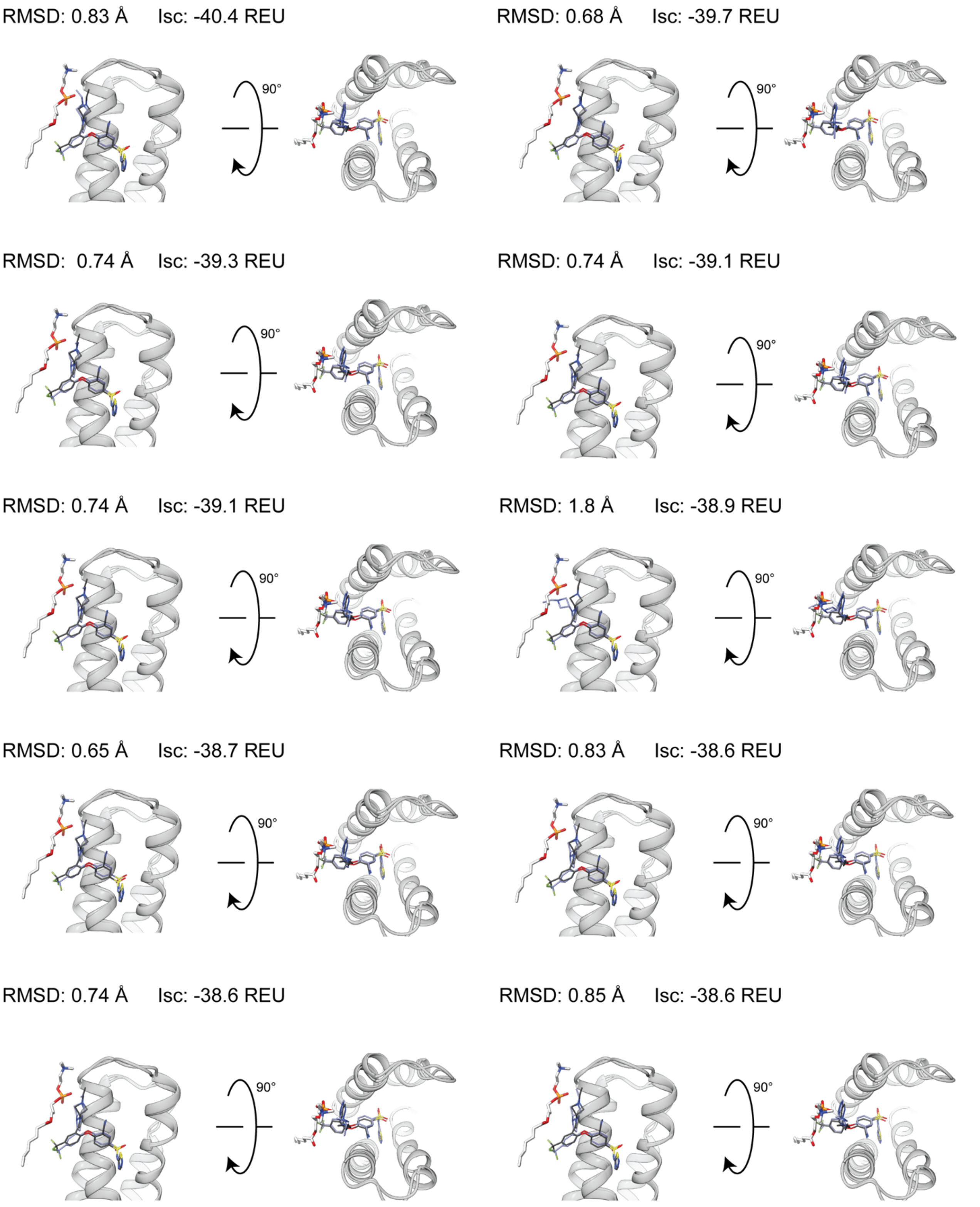


**Figure S13.2.** PDB 5EK0. Top 10 interface-scoring (`LigInterface`) poses from GALigandDock, 100,000 total poses, padding 7 Å.

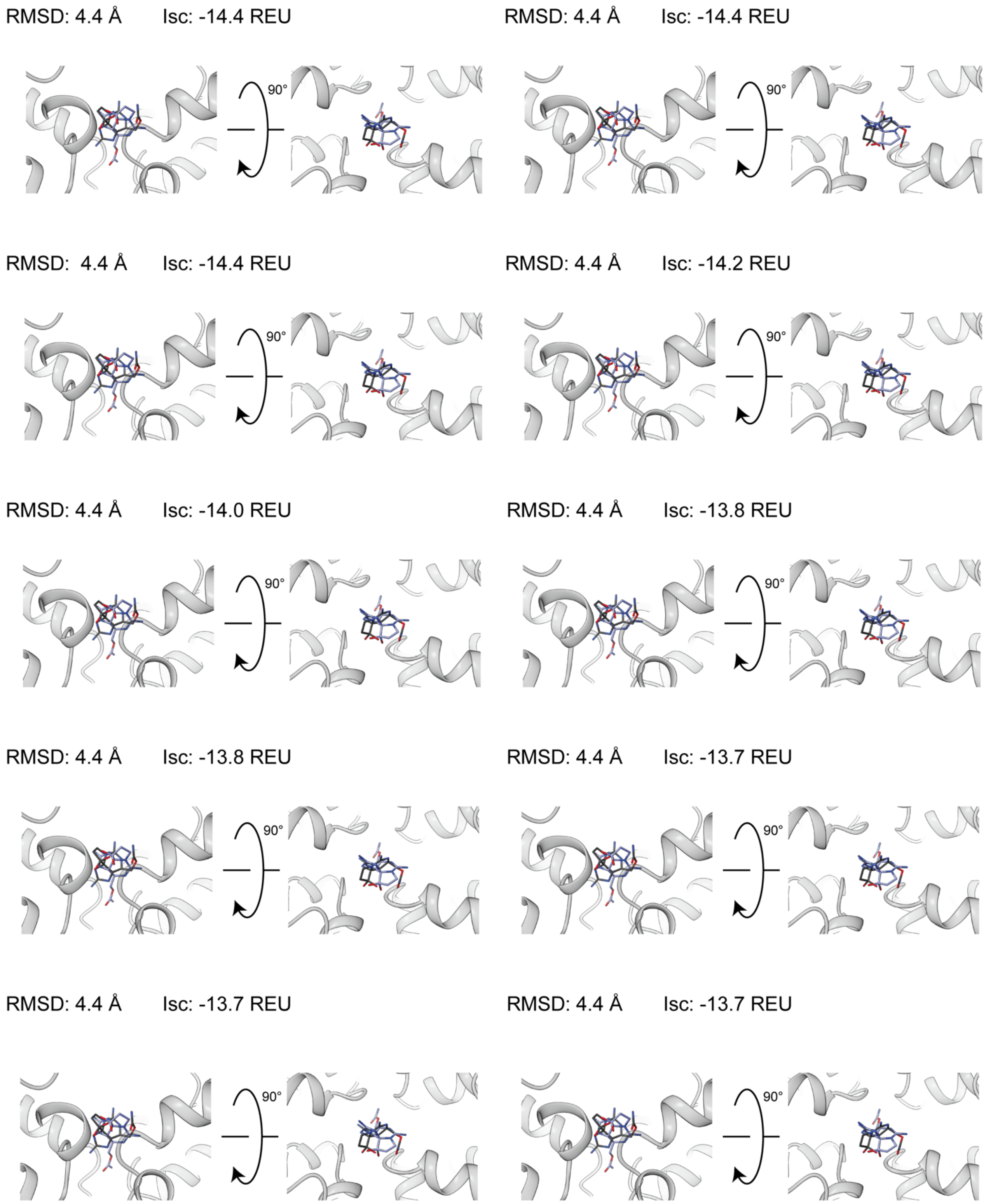


**Figure S14.1.** PDB 6J8G. Top 10 interface-scoring (`interface_delta_X`) poses from RosettaLigand, 100,000 total poses, all atom ligand interface.

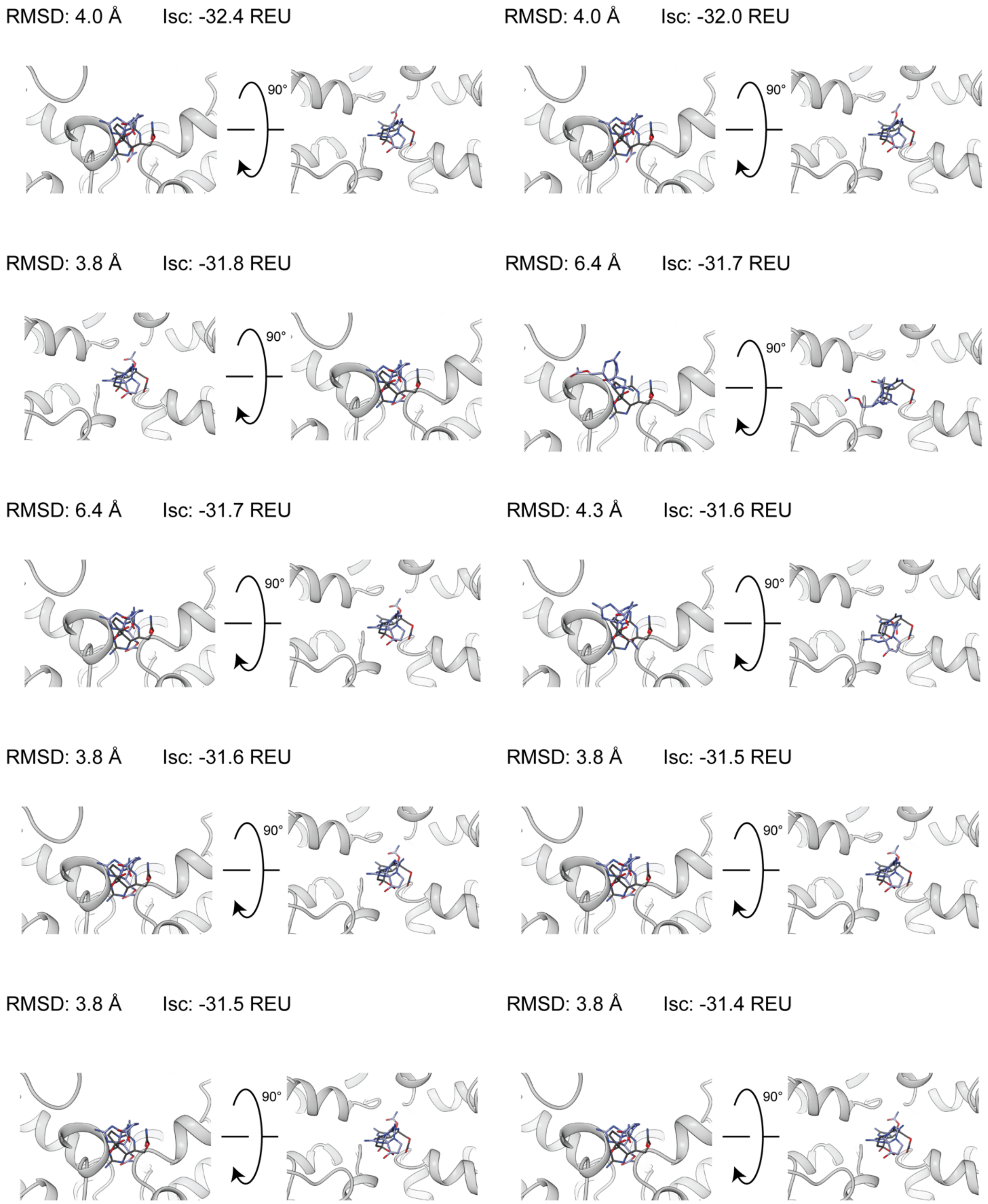


**Figure S14.2.** PDB 6J8G. Top 10 interface-scoring (`LigInterface`) poses from GALigandDock, 100,000 total poses, padding 7 Å.

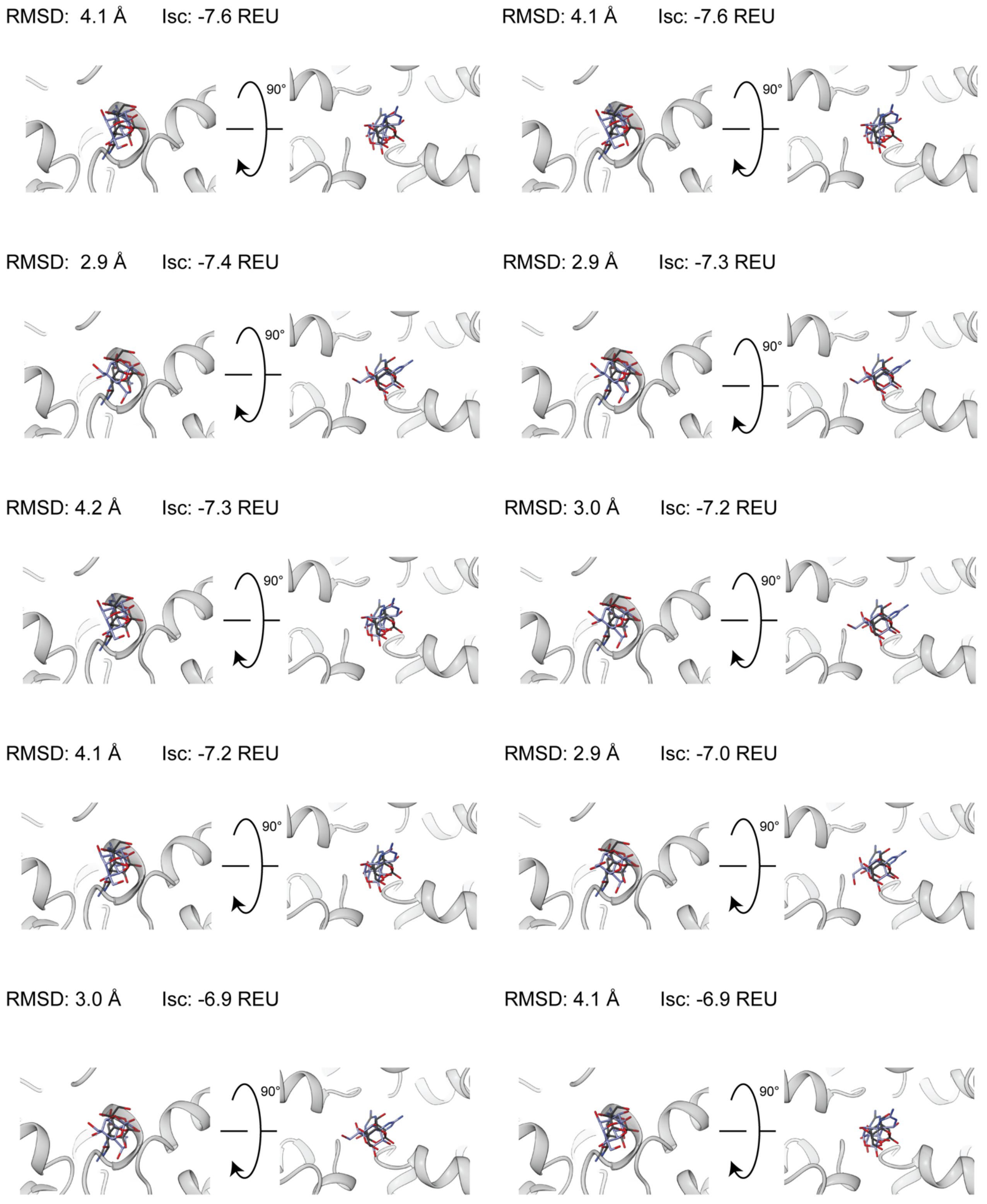


**Figure S15.1.** PDB 6J8I. Top 10 interface-scoring (`interface_delta_X`) poses from RosettaLigand, 100,000 total poses, all atom ligand interface.

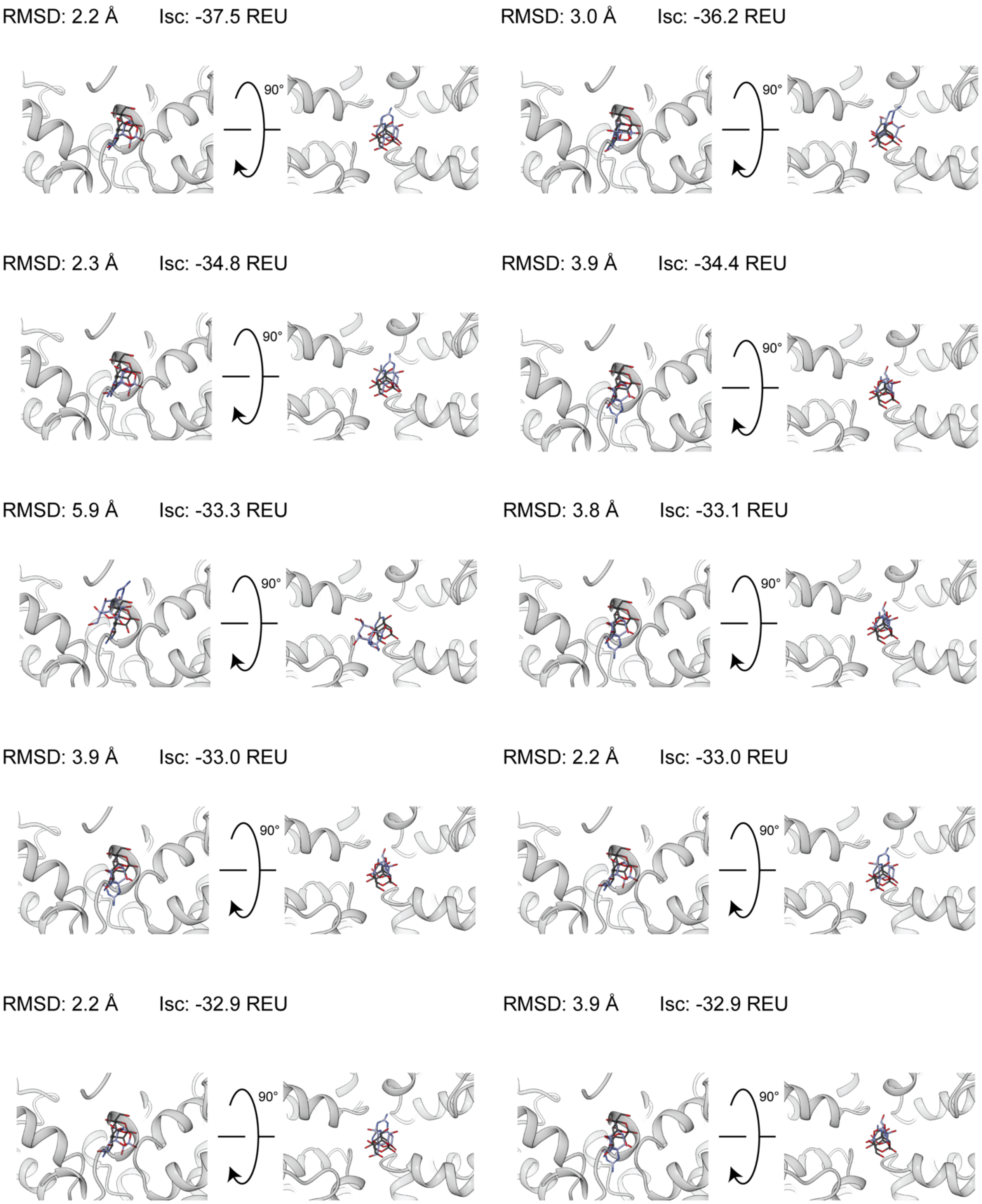


**Figure S15.2.** PDB 6J8I. Top 10 interface-scoring (`LigInterface`) poses from GALigandDock, 100,000 total poses, padding 7 Å.

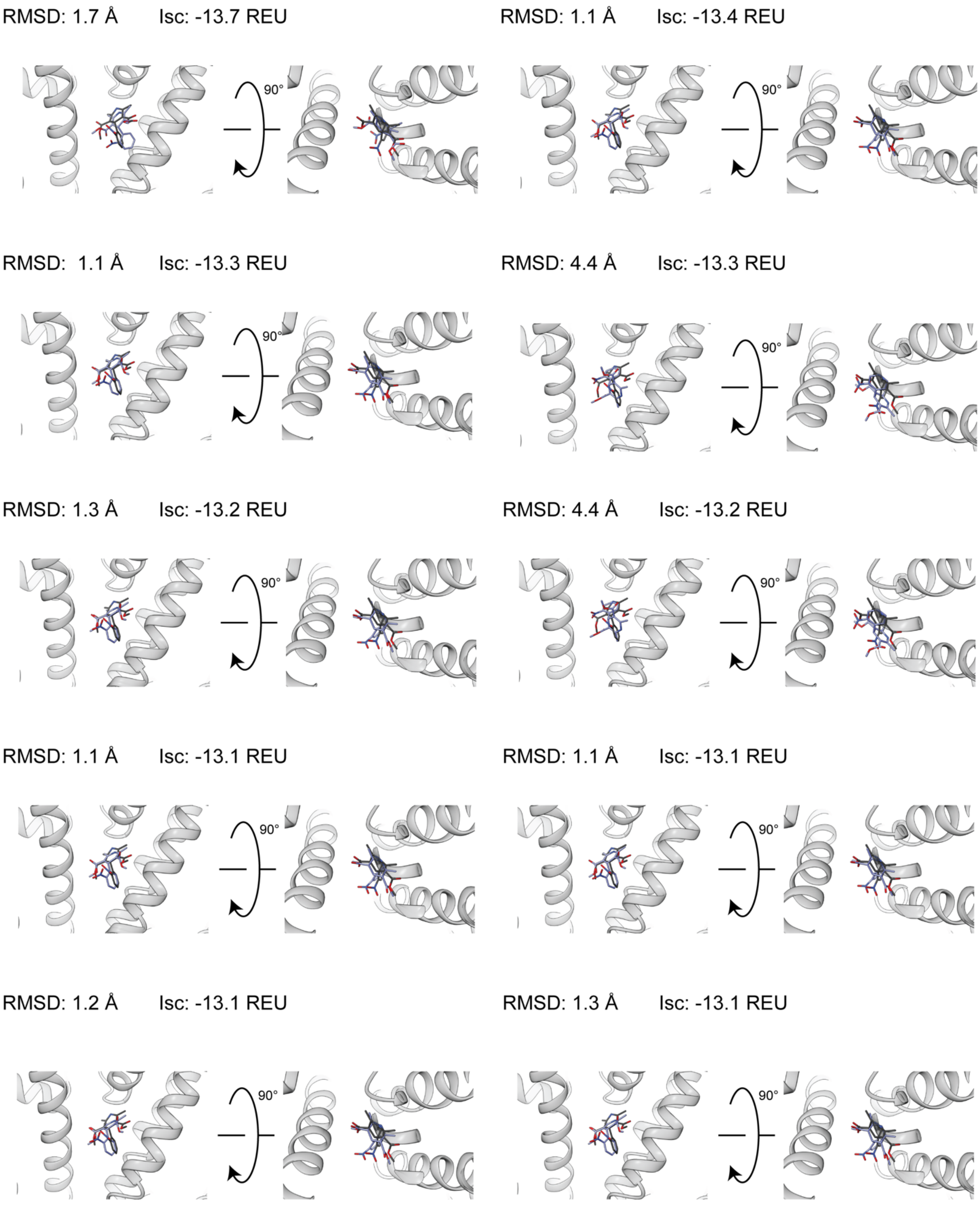


**Figure S16.1.** PDB 6JP5. Top 10 interface-scoring (`interface_delta_X`) poses from RosettaLigand, 100,000 total poses, all atom ligand interface.

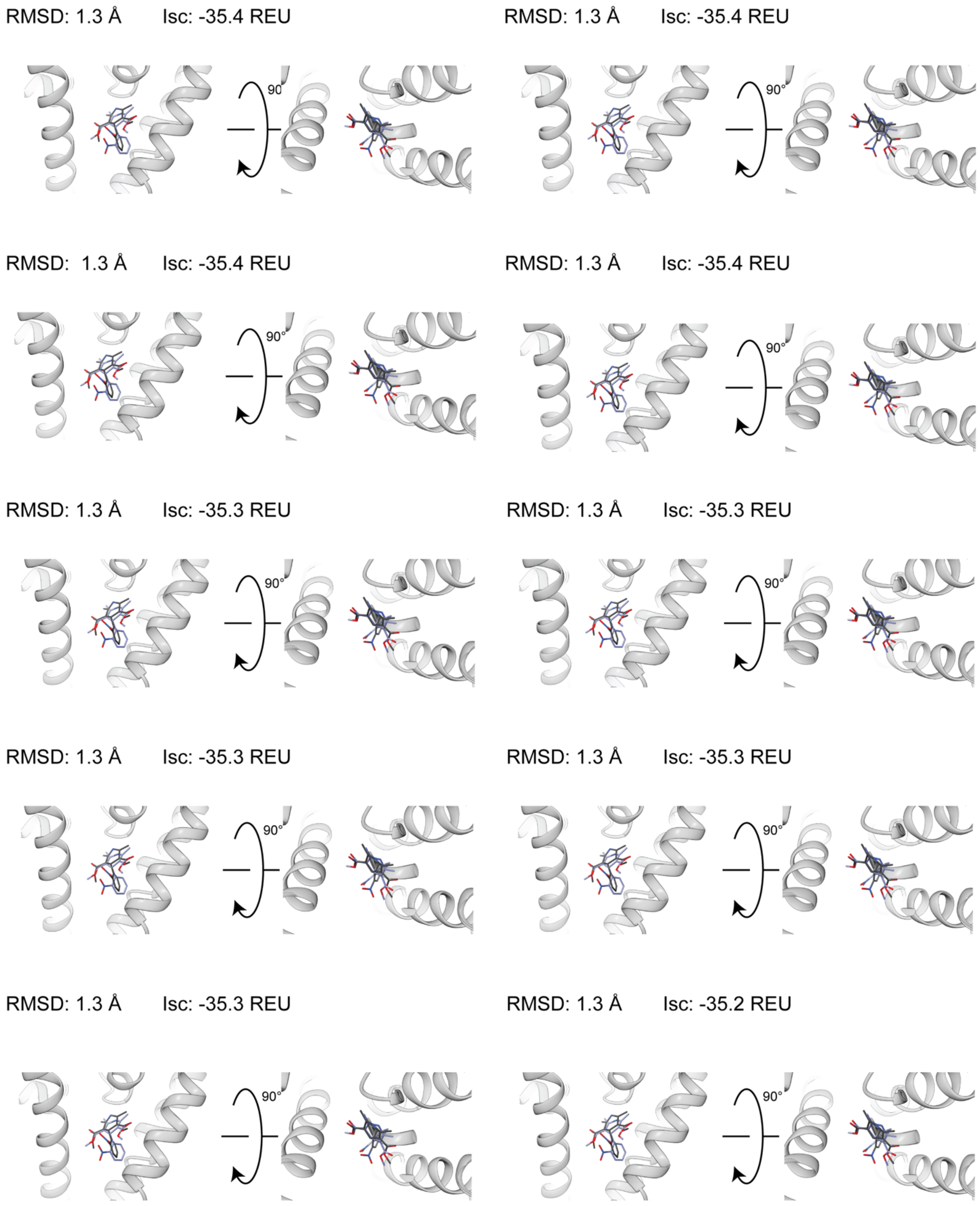


**Figure S16.2.** PDB 6JP5. Top 10 interface-scoring (`LigInterface`) poses from GALigandDock, 100,000 total poses, padding 7 Å.

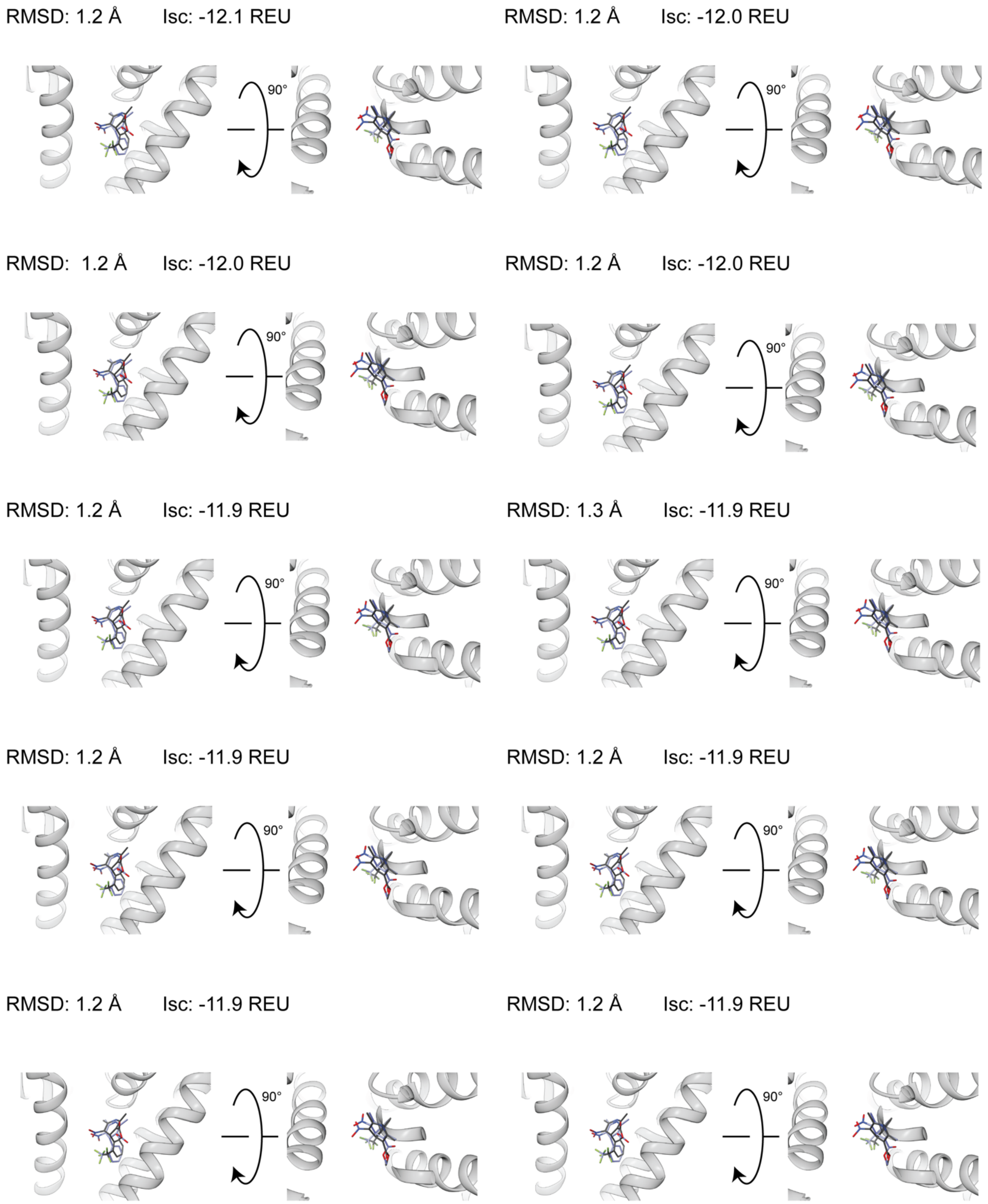


**Figure S17.1.** PDB 6JP8. Top 10 interface-scoring (`interface_delta_X`) poses from RosettaLigand, 100,000 total poses, all atom ligand interface.

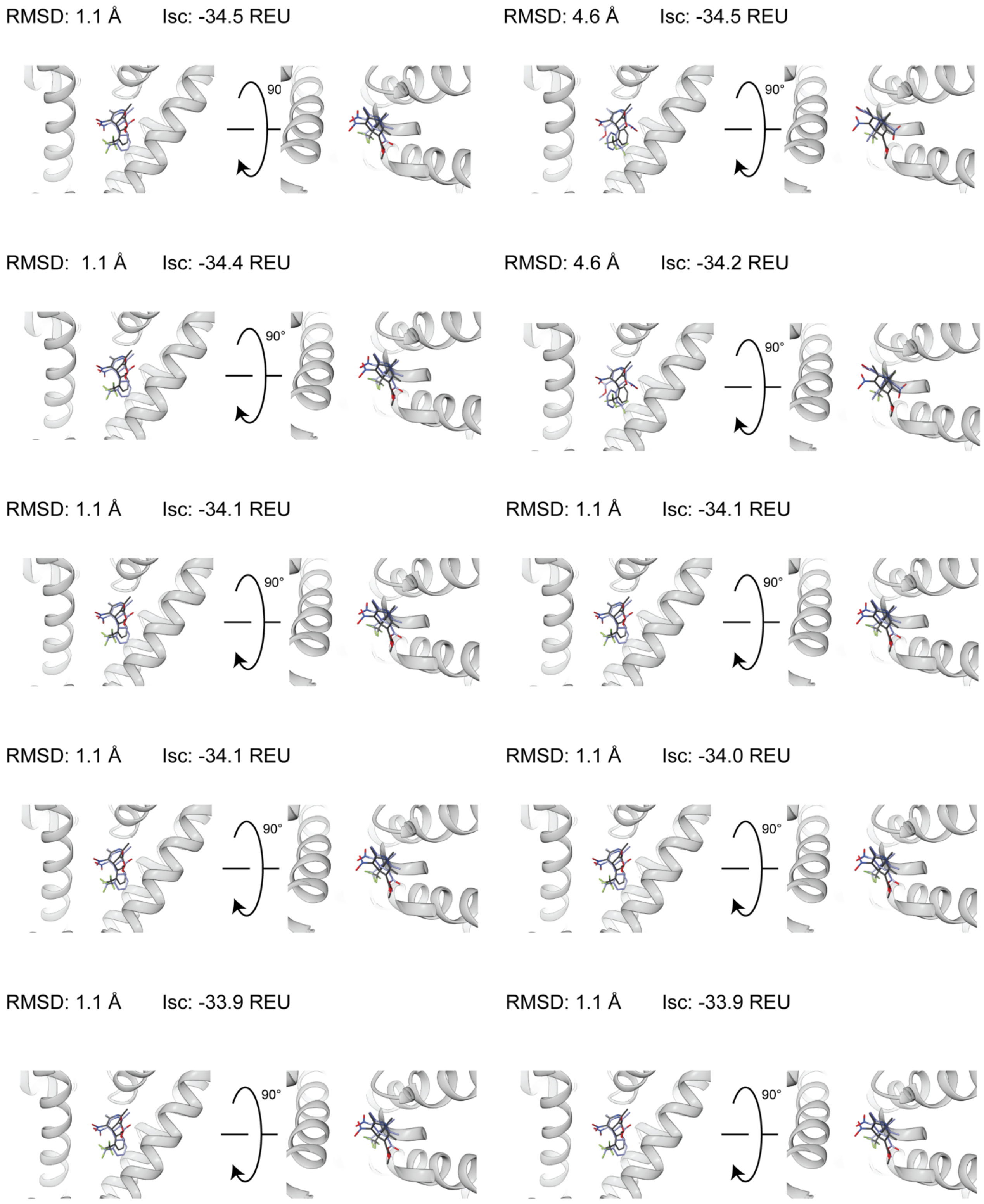


**Figure S17.2.** PDB 6JP8. Top 10 interface-scoring (`LigInterface`) poses from GALigandDock, 100,000 total poses, padding 7 Å.

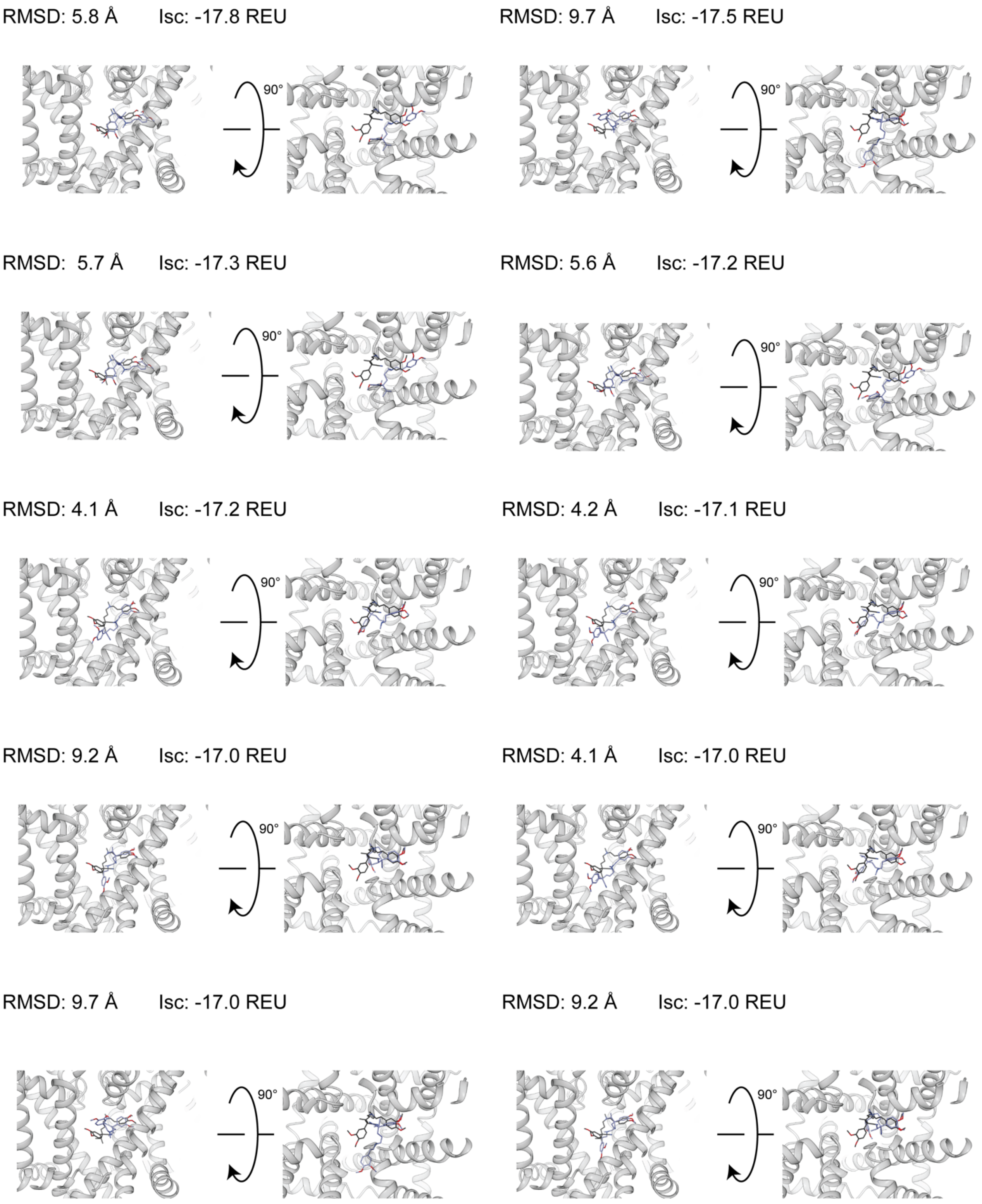


**Figure S18.1.** PDB 6JPA-1. Top 10 interface-scoring (`interface_delta_X`) poses from RosettaLigand, 100,000 total poses, all atom ligand interface.

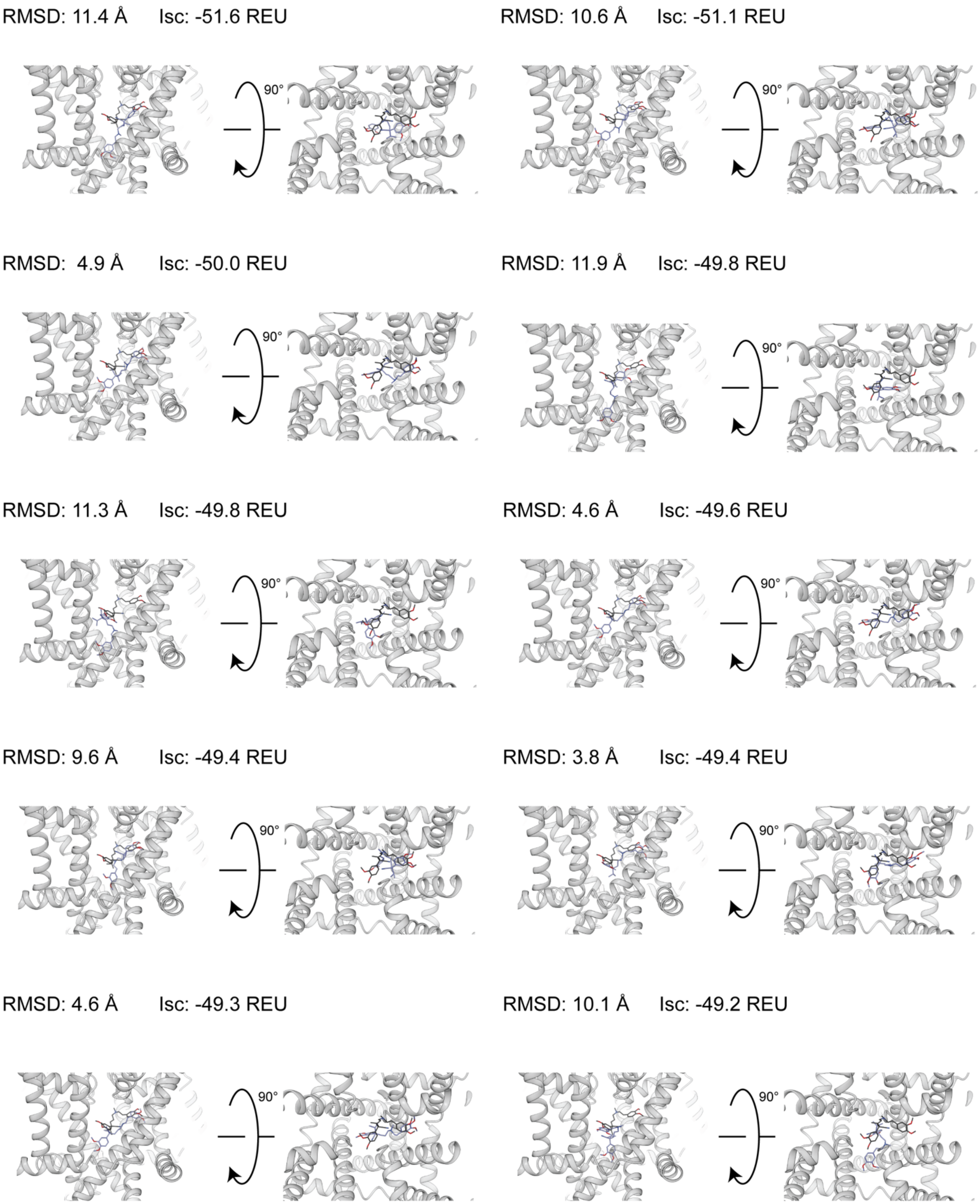


**Figure S18.2.** PDB 6JPA-1. Top 10 interface-scoring (`LigInterface`) poses from GALigandDock, 100,000 total poses, padding 7 Å.

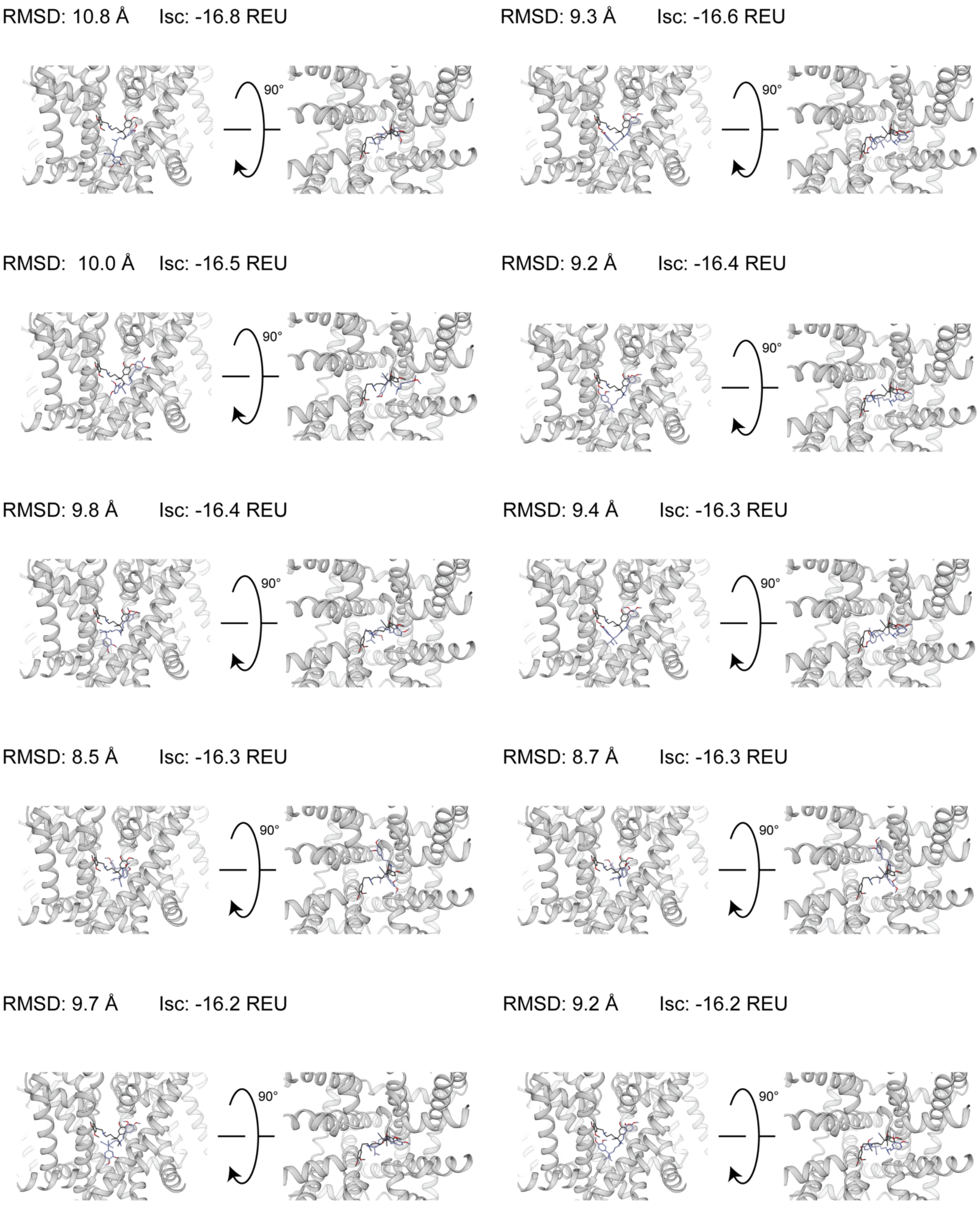


**Figure S19.1.** PDB 6JPA-2. Top 10 interface-scoring (`interface_delta_X`) poses from RosettaLigand, 100,000 total poses, all atom ligand interface.

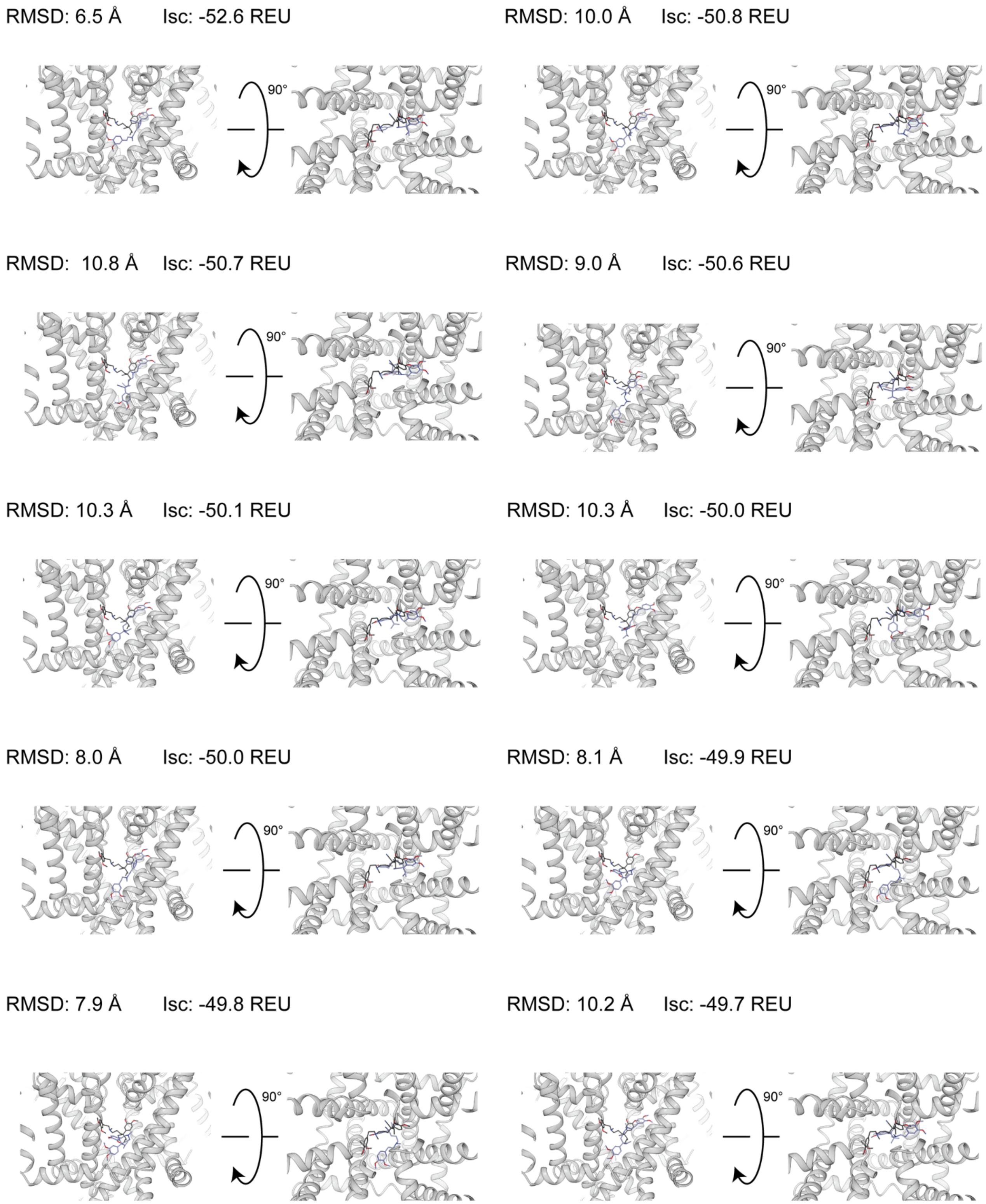


**Figure S19.2.** PDB 6JPA-2. Top 10 interface-scoring (`LigInterface`) poses from GALigandDock, 100,000 total poses, padding 7 Å.

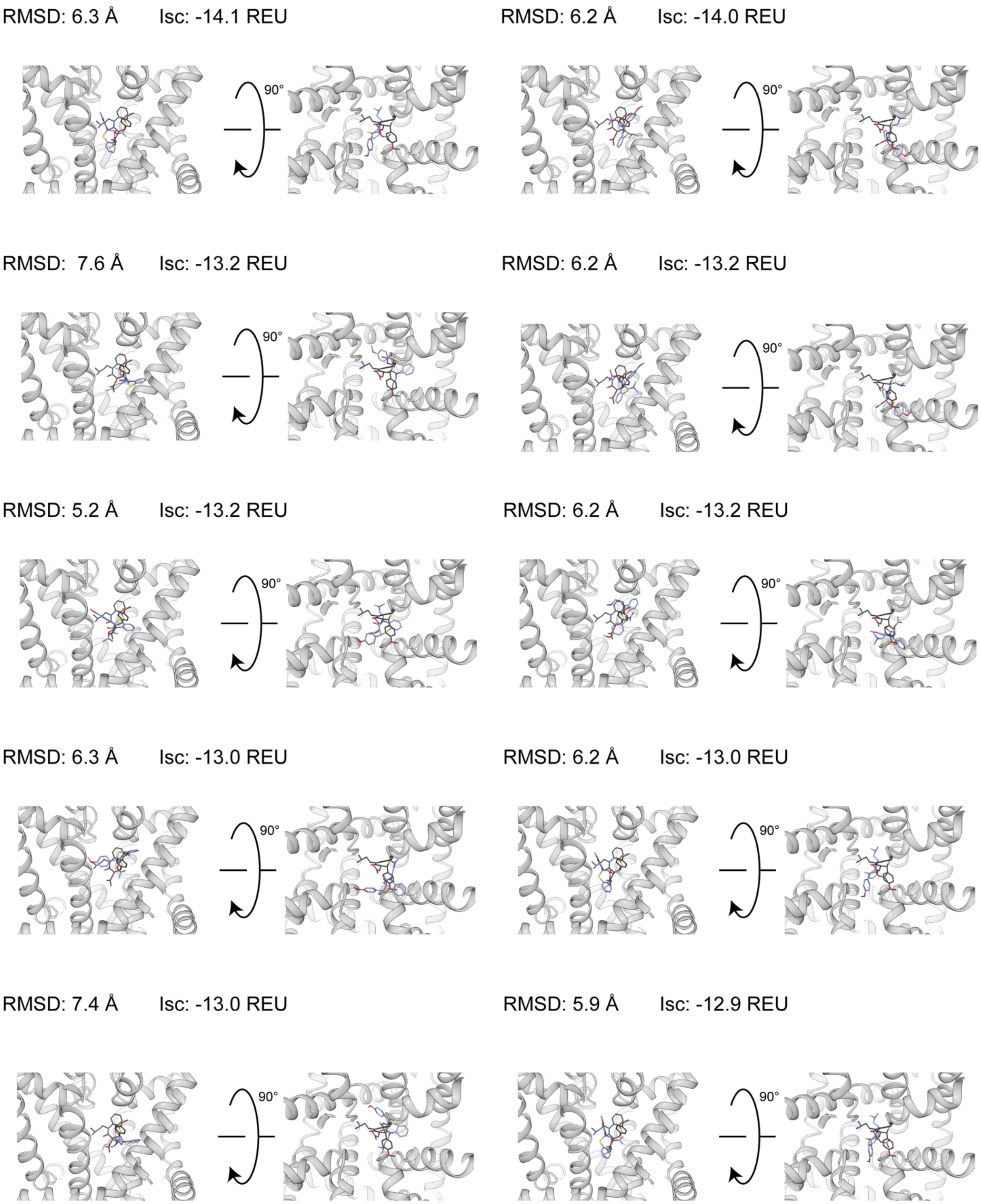


**Figure S20.1.** PDB 6JPB. Top 10 interface-scoring (`interface_delta_X`) poses from RosettaLigand, 100,000 total poses, all atom ligand interface.

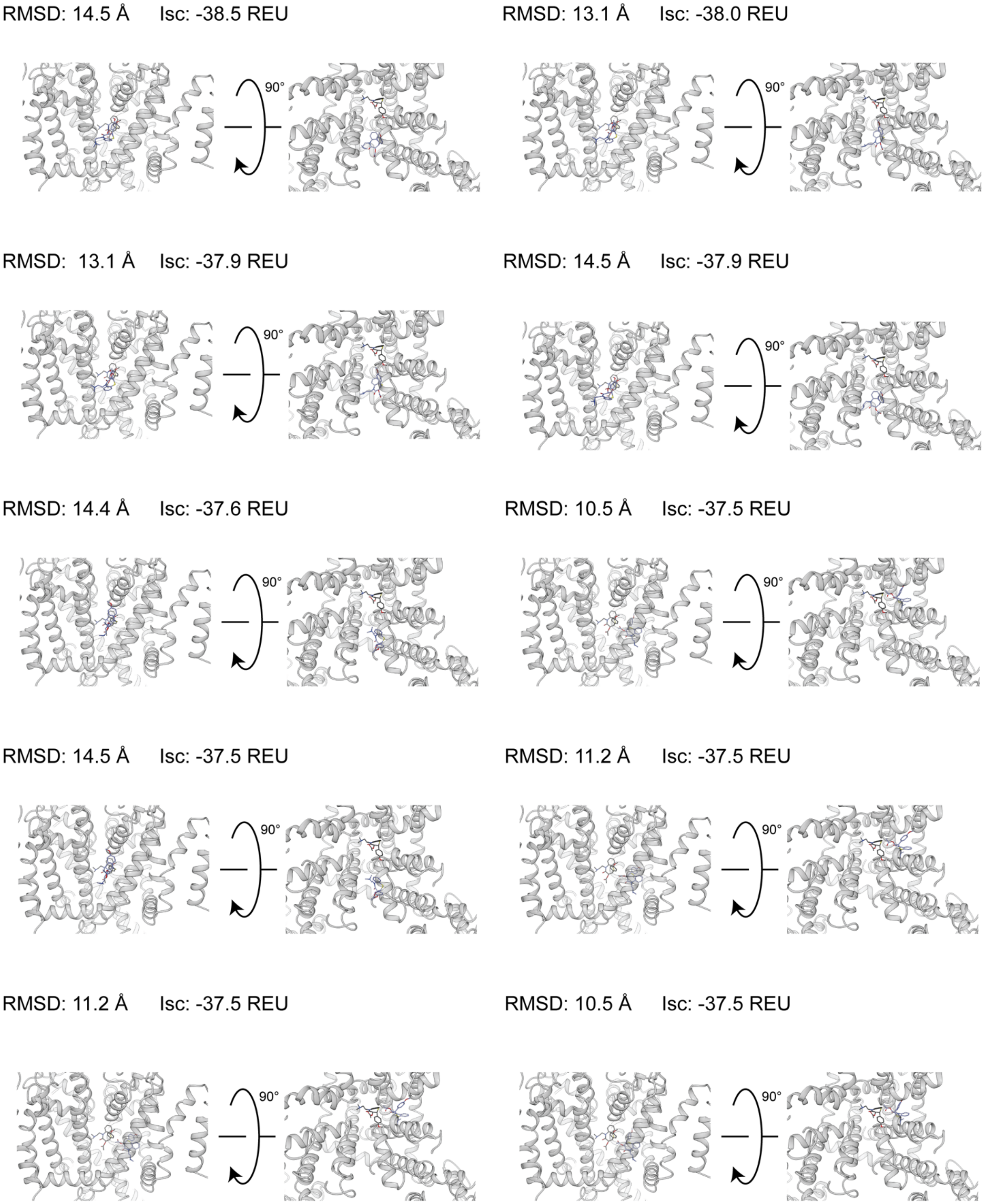


**Figure S20.2.** PDB 6JPB. Top 10 interface-scoring (`LigInterface`) poses from GALigandDock, 100,000 total poses, padding 7 Å.

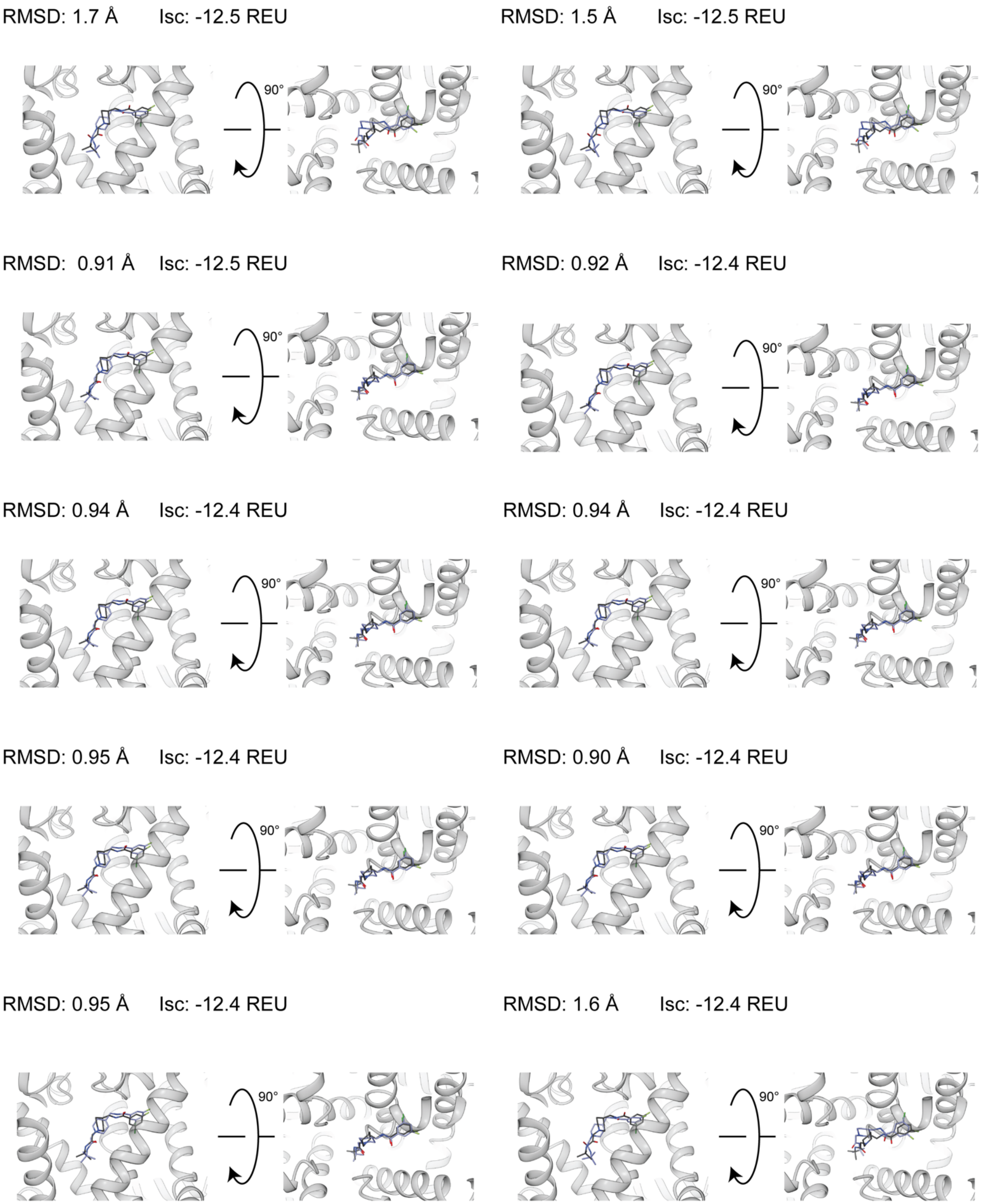


**Figure S21.1.** PDB 6KZP. Top 10 interface-scoring (`interface_delta_X`) poses from RosettaLigand, 100,000 total poses, all atom ligand interface.

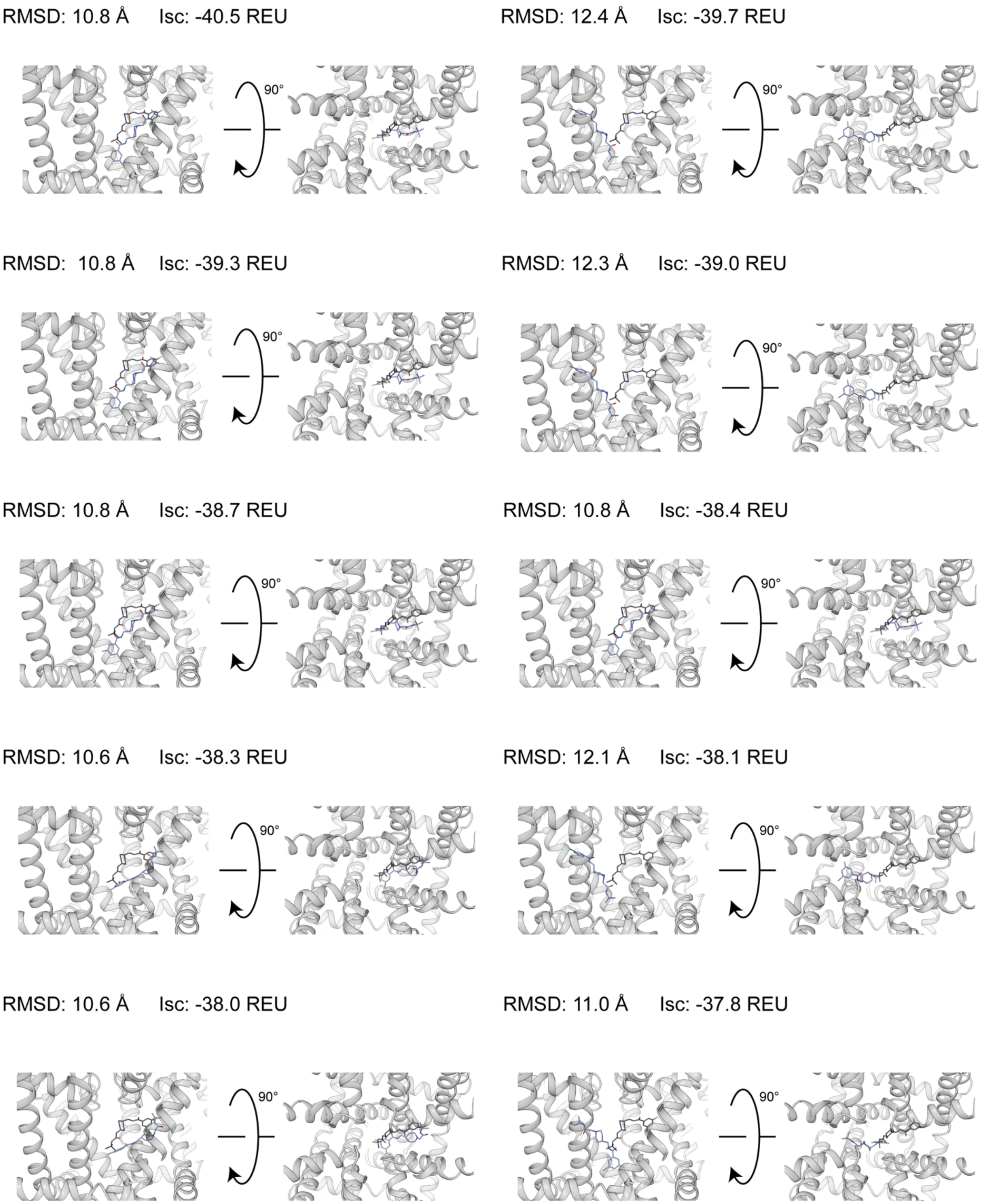


**Figure S21.2.** PDB 6KZP. Top 10 interface-scoring (`LigInterface`) poses from GALigandDock, 100,000 total poses, padding 7 Å.

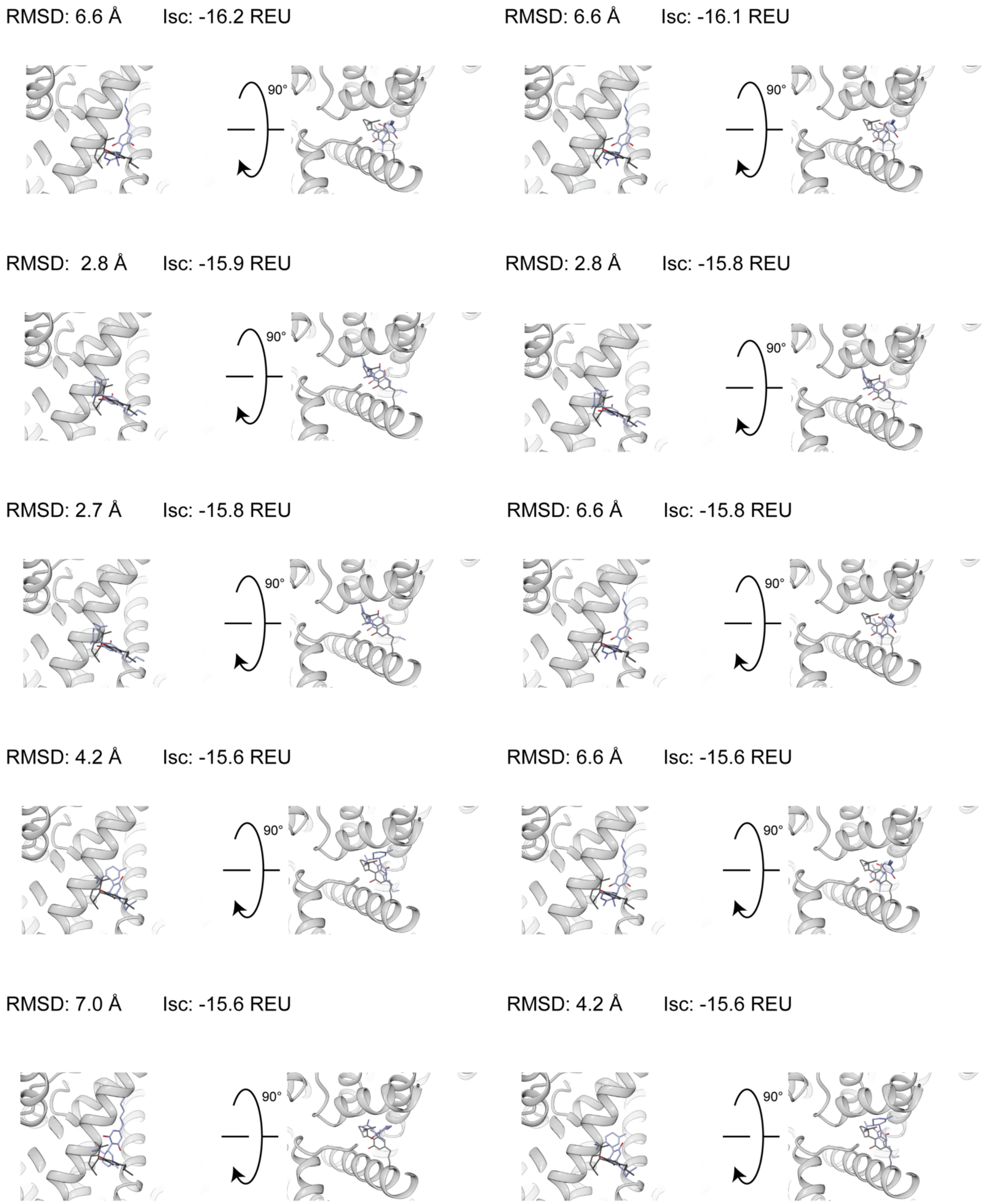


**Figure S22.1.** PDB 6U88. Top 10 interface-scoring (`interface_delta_X`) poses from RosettaLigand, 100,000 total poses, all atom ligand interface.

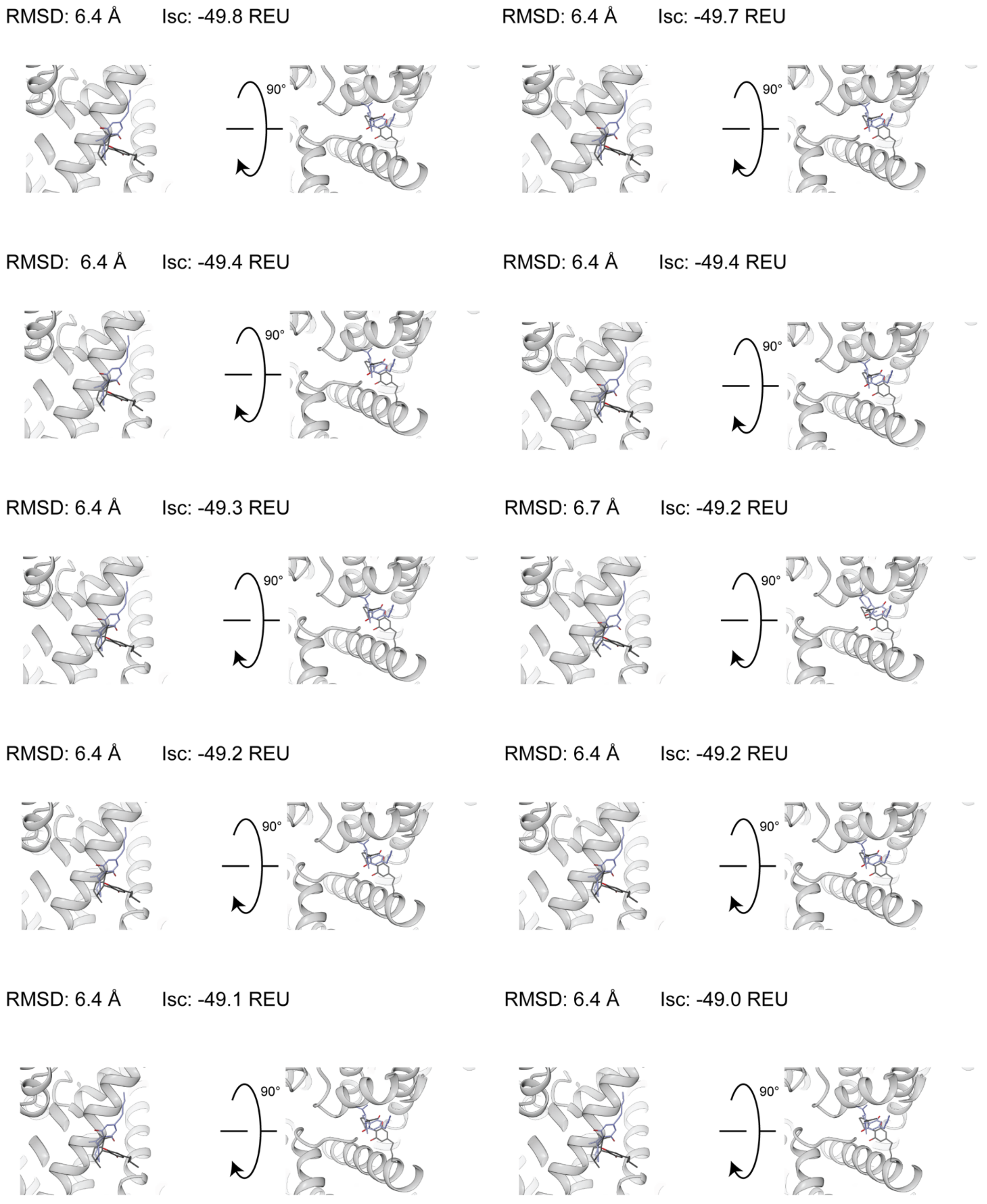


**Figure S22.2.** PDB 6U88. Top 10 interface-scoring (`LigInterface`) poses from GALigandDock, 100,000 total poses, padding 7 Å.

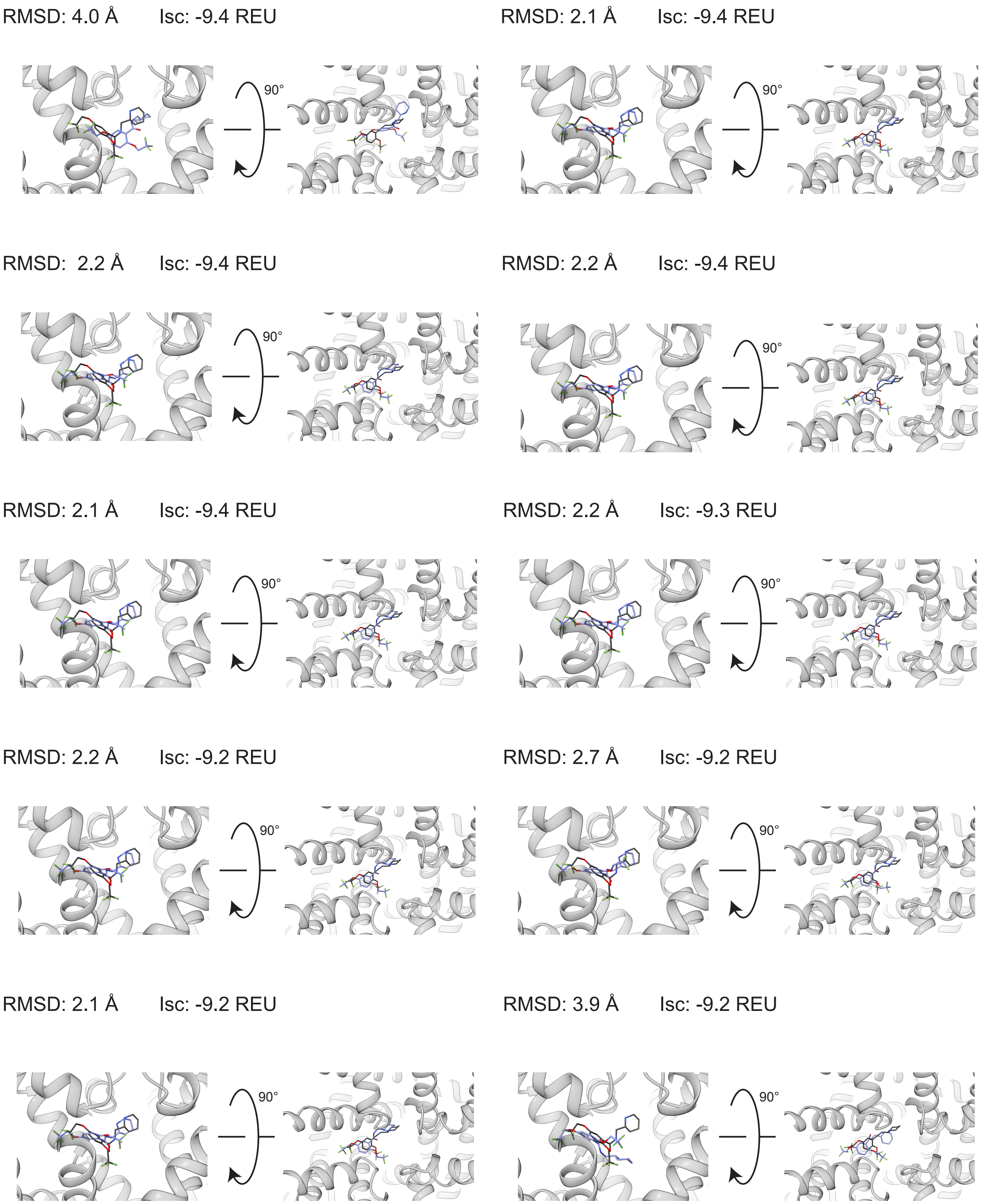


**Figure S23.1.** PDB 6UZ0. Top 10 interface-scoring (`interface_delta_X`) poses from RosettaLigand, 100,000 total poses, all atom ligand interface.

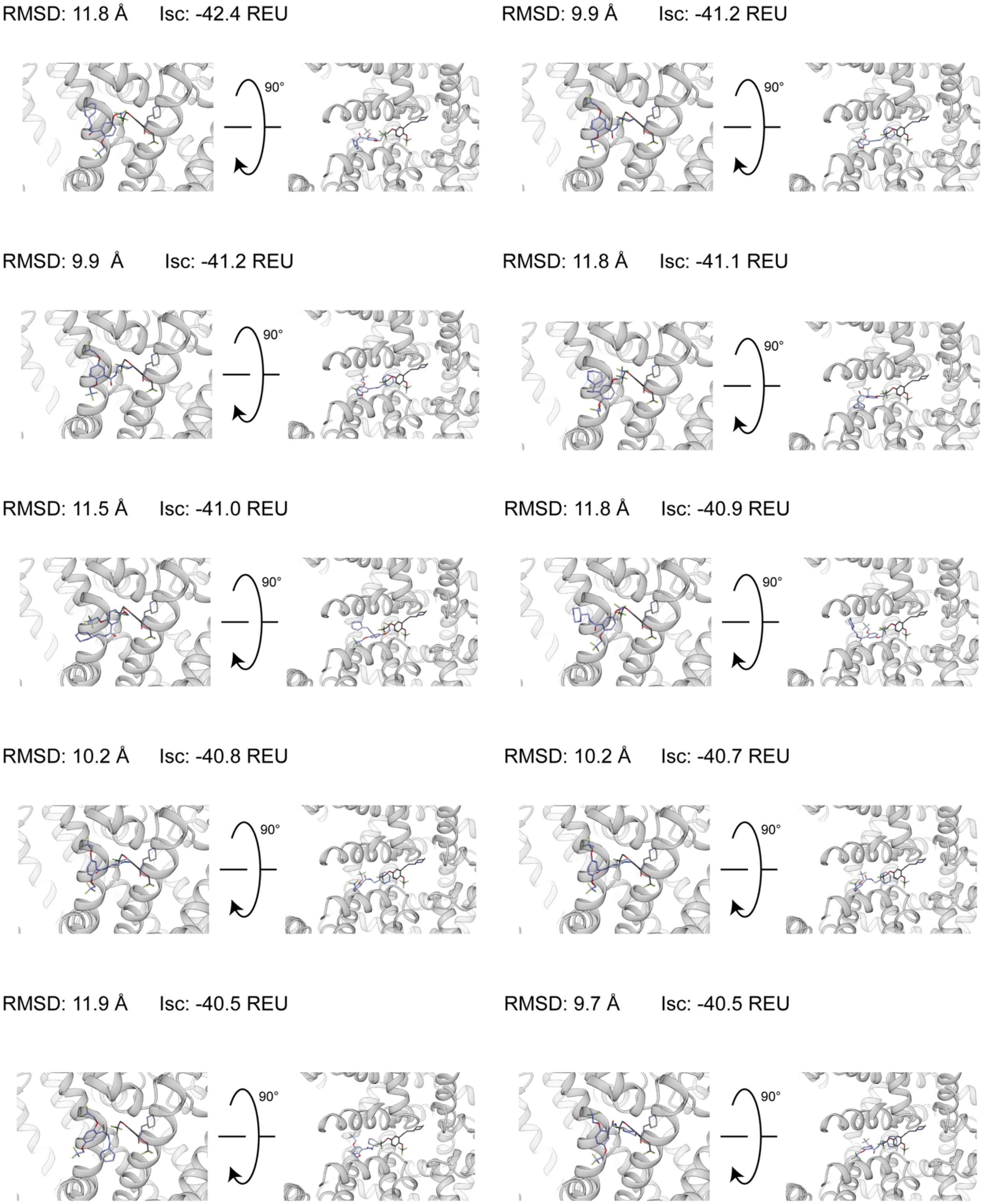


**Figure S23.2.** PDB 6UZ0. Top 10 interface-scoring (`LigInterface`) poses from GALigandDock, 100,000 total poses, padding 7 Å.

**Table S1.1.** Summary of minimum RMSD (Å) pose from each RosettaLigand docking set.

| | All ligand atom cutoff | | Neighbor atom cutoff | |
|---|---|---|---|---|
| **PDB ID** | **20,000 poses** | **100,000 poses** | **20,000 poses** | **100,000 poses** |
| **5EK0** | 0.92 | 0.91 | 0.92 | 0.88 |
| **6J8G** | 0.71 | 0.70 | 0.70 | 0.66 |
| **6J8I** | 0.55 | 0.54 | 0.58 | 0.58 |
| **6JP5** | 0.96 | 0.77 | 0.94 | 0.85 |
| **6JP8** | 0.92 | 0.93 | 1.1 | 0.89 |
| **6JPA-1** | 1.4 | 1.4 | 1.5 | 1.3 |
| **6JPA-2** | 2.3 | 2.2 | 2.6 | 2.2 |
| **6JPB** | 2.3 | 2.5 | 2.5 | 1.9 |
| **6KZP** | 0.71 | 0.73 | 0.73 | 0.72 |
| **6U88** | 1.0 | 1.0 | 1.2 | 1.1 |
| **6UZ0** | 1.2 | 1.2 | 1.2 | 1.2 |
| **Average** | 1.2 ± 0.6 | 1.2 ± 0.6 | 1.3 ± 0.7 | 1.1 ± 0.5 |
| **Median** | 1.0 | 0.93 | 1.1 | 0.89 |

**Table S1.2.** Summary of minimum RMSD (Å) pose from each GALigandDock docking set.

| | Padding 2 Å | | Padding 4 Å | | Padding 7 Å | |
|---|---|---|---|---|---|---|
| **PDB ID** | **20,000 poses** | **100,000 poses** | **20,000 poses** | **100,000 poses** | **20,000 poses** | **100,000 poses** |
| **5EK0** | 0.51 | 0.56 | 0.56 | 0.57 | 0.71 | 0.65 |
| **6J8G** | 0.77 | 0.75 | 1.1 | 0.65 | 0.8 | 0.76 |
| **6J8I** | 0.96 | 0.94 | 0.96 | 0.83 | 0.81 | 0.81 |
| **6JP5** | 1.8 | 1.8 | 1.3 | 1.2 | 1.3 | 1.3 |
| **6JP8** | 1.2 | 1.2 | 1.0 | 1.0 | 1.1 | 0.97 |
| **6JPA-1** | 1.9 | 2.0 | 2.3 | 2.0 | 2.7 | 2.0 |
| **6JPA-2** | 2.7 | 2.4 | 2.4 | 2.4 | 2.3 | 2.1 |
| **6JPB** | 1.7 | 1.3 | 1.7 | 1.7 | 1.1 | 1.1 |
| **6KZP** | 0.83 | 0.83 | 0.84 | 0.83 | 0.81 | 0.84 |
| **6U88** | 1.3 | 1.4 | 1.4 | 1.4 | 0.99 | 0.90 |
| **6UZ0** | 1.5 | 1.3 | 1.5 | 1.4 | 1.2 | 0.94 |
| **Average** | 1.4 ± 0.6 | 1.3 ± 0.6 | 1.4 ± 0.6 | 1.3 ± 0.6 | 1.3 ± 0.6 | 1.1 ± 0.5 |
| **Median** | 1.3 | 1.3 | 1.3 | 1.2 | 1.1 | 0.94 |

**Table S2.1.** RMSD (Å) of closest pose to native structural coordinates and lowest interface energy (`interface_delta_X`) pose of the top 10 % of poses by total energy (`total_score`) for RosettaLigand, using all ligand atoms for interface scoring cutoff. †

| | All ligand atom cutoff, 20,000 poses | | | | All ligand atom cutoff, 100,000 poses | | | |
|---|---|---|---|---|---|---|---|---|
| | Closest pose to native | | Lowest Interface Energy | | Closest pose to native | | Lowest Interface Energy | |
| **PDB ID** | **RMSD (Å)** | **Interface energy (REU)** | **RMSD (Å)** | **Interface energy (REU)** | **RMSD (Å)** | **Interface energy (REU)** | **RMSD (Å)** | **Interface energy (REU)** |
| **5EK0** | 0.92 | -13.5 | 2.8 | -17.4 | 0.91 | -11.1 | 2.7 | -17.4 |
| **6J8G** | 0.71 | -6.0 | 4.2 | -14.8 | 0.7 | -7.6 | 4.4 | -14.4 |
| **6J8I** | 0.55 | -2.7 | 4.1 | -7.2 | 0.54 | -4.5 | 4.1 | -7.6 |
| **6JP5** | 0.96 | -10.4 | 4.4 | -13.2 | 0.77 | -9.9 | 1.7 | -13.7 |
| **6JP8** | 0.92 | -9.5 | 1.2 | -12.0 | 0.93 | -9.7 | 1.2 | -12.1 |
| **6JPA-1** | 1.4 | -11.8 | 9.7 | -17.5 | 1.4 | -13.1 | 5.8 | -17.8 |
| **6JPA-2** | 2.3 | -7.5 | 9.5 | -16.5 | 2.2 | -10.0 | 10.8 | -16.8 |
| **6JPB** | 2.3 | -8.1 | 6.0 | -12.9 | 2.5 | -2.2 | 6.3 | -14.1 |
| **6KZP** | 0.71 | -8.9 | 0.91 | -12.5 | 0.73 | -10.2 | 1.7 | -12.5 |
| **6U88** | 1.0 | -9.6 | 6.0 | -15.8 | 1.0 | -6.0 | 6.6 | -16.2 |
| **6UZ0** | 1.2 | -7.7 | 3.4 | -9.5 | 1.2 | -4.5 | 4.0 | -9.4 |
| **Average** | 1.2 ± 0.6 | -- | 4.7 ± 2.9 | -- | 1.2 ± 0.6 | -- | 4.5 ± 2.8 | -- |
| **Median** | 1.0 | -- | 4.2 | -- | 0.93 | -- | 4.1 | -- |

† Interface energy statistics were not computed since they are not standardized like RMSD.

**Table S2.2.** RMSD (Å) of closest pose to native structural coordinates and lowest interface energy (`interface_delta_X`) pose of the top 10 % of poses by total energy (`total_score`) from RosettaLigand with ligand neighbor atom for interface scoring cutoff. †

| | Neighbor atom cutoff, 20,000 poses | | | | Neighbor atom cutoff, 100,000 poses | | | |
|---|---|---|---|---|---|---|---|---|
| | Closest pose to native | | Lowest Interface Energy | | Closest pose to native | | Lowest Interface Energy | |
| **PDB ID** | **RMSD (Å)** | **Interface energy (REU)** | **RMSD (Å)** | **Interface energy (REU)** | **RMSD (Å)** | **Interface energy (REU)** | **RMSD (Å)** | **Interface energy (REU)** |
| **5EK0** | 0.92 | -7.8 | 2.6 | -16.7 | 0.88 | -7.7 | 2.5 | -17.9 |
| **6J8G** | 0.70 | -7.3 | 4.2 | -15.2 | 0.66 | -5.9 | 4.2 | -15.4 |
| **6J8I** | 0.58 | -4.7 | 4.2 | -7.5 | 0.58 | -4.3 | 4.1 | -7.9 |
| **6JP5** | 0.94 | -10.3 | 1.8 | -13.5 | 0.85 | -7.6 | 1.5 | -14.0 |
| **6JP8** | 1.1 | -10.1 | 1.2 | -12.2 | 0.89 | -9.9 | 1.3 | -12.5 |
| **6JPA-1** | 1.5 | -11.5 | 7.7 | -16.6 | 1.3 | -12.5 | 5.2 | -16.8 |
| **6JPA-2** | 2.6 | -5.0 | 6.6 | -16.6 | 2.2 | -9.2 | 9.5 | -16.1 |
| **6JPB** | 2.5 | -6.1 | 6.2 | -14.5 | 1.9 | -7.8 | 6.0 | -13.9 |
| **6KZP** | 0.73 | -9.6 | 1.0 | -12.5 | 0.72 | -8.5 | 6.5 | -13.1 |
| **6U88** | 1.2 | -9.5 | 6.1 | -14.7 | 1.1 | -9.2 | 6.6 | -16.4 |
| **6UZ0** | 1.2 | -7.7 | 4.0 | -8.9 | 1.2 | -7.8 | 9.1 | -10.2 |
| **Average** | 1.3 ± 0.7 | -- | 4.1 ± 2.3 | -- | 1.1 ± 0.5 | -- | 5.1 ± 2.8 | -- |
| **Median** | 1.1 | -- | 4.2 | -- | 0.89 | -- | 5.2 | -- |

† Interface energy statistics were not computed since they are not standardized like RMSD.

**Table S2.3.** RMSD (Å) of closest pose to native structural coordinates and lowest interface energy (`LigInterface`) pose from GALigandDock, using the top 10 % of poses by total energy (`total_score`) and 2 Å padding. †

| | 2 Å padding, 20,000 poses | | | | 2 Å padding, 100,000 poses | | | |
|---|---|---|---|---|---|---|---|---|
| | Closest pose to native | | Lowest Interface Energy | | Closest pose to native | | Lowest Interface Energy | |
| **PDB ID** | **RMSD (Å)** | **Interface energy (REU)** | **RMSD (Å)** | **Interface energy (REU)** | **RMSD (Å)** | **Interface energy (REU)** | **RMSD (Å)** | **Interface energy (REU)** |
| **5EK0** | 0.51 | -36.0 | 0.72 | -38.5 | 0.56 | -36.4 | 1.3 | -38.8 |
| **6J8G** | 0.77 | -18.7 | 3.8 | -29.5 | 0.75 | -18.9 | 3.8 | -31.4 |
| **6J8I** | 0.96 | -12.2 | 6.2 | -30.0 | 0.94 | -12.3 | 6.2 | -29.9 |
| **6JP5** | 1.8 | -20.6 | 4.7 | -30.6 | 1.8 | -20.5 | 5.3 | -31.5 |
| **6JP8** | 1.2 | -11.7 | 6.1 | -25.3 | 1.2 | -13.4 | 6.1 | -25.3 |
| **6JPA-1** | 1.9 | -37.8 | 5.6 | -47.6 | 2.0 | -38.3 | 5.4 | -48.8 |
| **6JPA-2** | 2.7 | -35.6 | 11.8 | -48.2 | 2.4 | -30.9 | 11.7 | -48.2 |
| **6JPB** | 1.7 | -20.8 | 4.7 | -30.7 | 1.3 | -18.2 | 4.6 | -30.8 |
| **6KZP** | 0.83 | -32.4 | 10.3 | -32.6 | 0.83 | -33.4 | 0.86 | -33.4 |
| **6U88** | 1.3 | -37.6 | 6.4 | -45.4 | 1.4 | -32.4 | 6.4 | -45.5 |
| **6UZ0** | 1.5 | -31.6 | 4.9 | -37.2 | 1.3 | -28.0 | 7.1 | -37.3 |
| **Average** | 1.4 ± 0.6 | -- | 5.9 ± 3.0 | -- | 1.3 ± 0.6 | -- | 5.3 ± 2.9 | -- |
| **Median** | 1.3 | -- | 5.6 | -- | 1.3 | -- | 5.4 | -- |

† Interface energy statistics were not computed since they are not standardized like RMSD.

**Table S2.4.** RMSD (Å) of closest pose to native structural coordinates and lowest interface energy (`LigInterface`) pose from GALigandDock, using the top 10 % of poses by total energy (`total_score`) and 4 Å padding. †

| | 4 Å padding, 20,000 poses | | | | 4 Å padding, 100,000 poses | | | |
|---|---|---|---|---|---|---|---|---|
| | Closest pose to native | | Lowest Interface Energy | | Closest pose to native | | Lowest Interface Energy | |
| **PDB ID** | **RMSD (Å)** | **Interface energy (REU)** | **RMSD (Å)** | **Interface energy (REU)** | **RMSD (Å)** | **Interface energy (REU)** | **RMSD (Å)** | **Interface energy (REU)** |
| **5EK0** | 0.56 | -37.1 | 0.74 | -38.7 | 0.57 | -37.0 | 0.66 | -38.7 |
| **6J8G** | 1.1 | -20.6 | 3.8 | -30.6 | 0.65 | -21.4 | 3.8 | -33.2 |
| **6J8I** | 0.96 | -18.1 | 3.9 | -33.9 | 0.83 | -17.4 | 2.2 | -34.9 |
| **6JP5** | 1.3 | -32.3 | 1.3 | -35.2 | 1.2 | -29.7 | 1.3 | -35.1 |
| **6JP8** | 1.0 | -30.4 | 1.1 | -32.4 | 1.0 | -30.5 | 1.1 | -32.4 |
| **6JPA-1** | 2.3 | -39.3 | 6.7 | -48.6 | 2.0 | -38.7 | 10.8 | -49.2 |
| **6JPA-2** | 2.4 | -32.7 | 9.8 | -47.7 | 2.4 | -32.7 | 11.1 | -50.4 |
| **6JPB** | 1.7 | -18.4 | 5.7 | -35.4 | 1.7 | -20.2 | 5.7 | -35.7 |
| **6KZP** | 0.84 | -31.1 | 10.2 | -34.1 | 0.83 | -30.8 | 10.8 | -38.6 |
| **6U88** | 1.4 | -41.4 | 6.3 | -47.6 | 1.4 | -36.0 | 6.3 | -48.5 |
| **6UZ0** | 1.5 | -30.6 | 10.0 | -38.4 | 1.4 | -29.2 | 11.3 | -38.6 |
| **Average** | 1.4 ± 0.6 | -- | 5.4 ± 3.6 | -- | 1.3 ± 0.6 | -- | 5.9 ± 4.4 | -- |
| **Median** | 1.3 | -- | 5.7 | -- | 1.2 | -- | 5.7 | -- |

† Interface energy statistics were not computed since they are not standardized like RMSD.

**Table S2.5.** RMSD (Å) of closest pose to native structural coordinates and lowest interface energy (`LigInterface`) pose from GALigandDock, using the top 10 % of poses by total energy (`total_score`) and 7 Å padding. †

| PDB ID | 7 Å padding, 20,000 poses | | | | 7 Å padding, 100,000 poses | | | |
|---|---|---|---|---|---|---|---|---|
| | Closest pose to native | | Lowest Interface Energy | | Closest pose to native | | Lowest Interface Energy | |
| | RMSD (Å) | Interface energy (REU) | RMSD (Å) | Interface energy (REU) | RMSD (Å) | Interface energy (REU) | RMSD (Å) | Interface energy (REU) |
| **5EK0** | 0.71 | -36.1 | 0.74 | -39.0 | 0.65 | -38.7 | 0.83 | -40.4 |
| **6J8G** | 0.80 | -21.9 | 4.0 | -32.5 | 0.76 | -23.8 | 4.0 | -32.4 |
| **6J8I** | 0.81 | -19.1 | 3.9 | -34.2 | 0.81 | -18.9 | 2.2 | -37.5 |
| **6JP5** | 1.3 | -34.4 | 1.3 | -34.4 | 1.3 | -35.4 | 1.3 | -35.4 |
| **6JP8** | 1.1 | -30.7 | 1.1 | -33.9 | 0.97 | -30.9 | 1.1 | -34.5 |
| **6JPA-1** | 2.7 | -47.6 | 3.6 | -49.1 | 2.0 | -39.0 | 11.4 | -51.6 |
| **6JPA-2** | 2.3 | -33.4 | 9.0 | -51.0 | 2.1 | -31.9 | 6.5 | -52.6 |
| **6JPB** | 1.1 | -20.1 | 14.6 | -38.0 | 1.1 | -20.4 | 14.5 | -38.5 |
| **6KZP** | 0.81 | -31.2 | 10.7 | -39.2 | 0.84 | -32.1 | 10.8 | -40.5 |
| **6U88** | 0.99 | -42.5 | 6.4 | -49.8 | 0.9 | -41.1 | 6.4 | -49.8 |
| **6UZ0** | 1.2 | -30.6 | 11.8 | -41.9 | 0.94 | -31.3 | 11.8 | -42.4 |
| **Average** | 1.3 ± 0.6 | -- | 6.1 ± 4.8 | -- | 1.1 ± 0.5 | -- | 6.4 ± 5.0 | -- |
| **Median** | 1.1 | -- | 4.0 | -- | 0.94 | -- | 6.4 | -- |

† Interface energy statistics were not computed since they are not standardized like RMSD.

**Table S3.** Pearson's correlation coefficient (r), confidence interval, and associated two-tailed p-values for covariates compared. Null hypothesis: there is no correlation between covariate 1 and covariate 2. n=10, α=0.05. Statistically significant values denoted with asterisk (*).

| Covariate 1 | Covariate 2 | Pearson's r | Confidence Interval | p-value |
|---|---|---|---|---|
| Total ligand rotatable bonds | Total ligand heavy atoms | 0.73 | [0.19, 0.93] | 0.016* |
| Total ligand rotatable bonds | Ligand molecular weight (Da) | 0.66 | [0.06, 0.91] | 0.036* |
| Total ligand rotatable bonds | Structure resolution (Å) | -0.26 | [-0.76, 0.45] | 0.47 |
| Total ligand rotatable bonds | Log10 RosettaLigand conformers | 0.74 | [0.21, 0.93] | 0.014* |
| Total ligand heavy atoms | Ligand molecular weight (Da) | 0.99 | [0.96, 1.0] | $3.9 \cdot 10^{-8}$* |
| Total ligand heavy atoms | Structure resolution (Å) | 0.12 | [-0.55, 0.7] | 0.74 |
| Total ligand heavy atoms | Log10 RosettaLigand conformers | 0.67 | [0.07, 0.91] | 0.035* |
| Ligand molecular weight (Da) | Structure resolution (Å) | 0.21 | [-0.48, 0.74] | 0.56 |
| Ligand molecular weight (Da) | Log10 RosettaLigand conformers | 0.68 | [0.10, 0.92] | 0.029* |
| Structure resolution (Å) | Log10 RosettaLigand conformers | 0.29 | [-0.41, 0.78] | 0.41 |

**Table S4.1.** $P_{Near}$ of RMSD vs `interface_delta_X` for RosettaLigand, all poses, all ligand atom mode.

| | $k_BT$=1.0 | | | | | | $k_BT$=0.62 | | | | | |
|---|---|---|---|---|---|---|---|---|---|---|---|---|
| | 20,000 poses | | | 100,000 poses | | | 20,000 poses | | | 100,000 poses | | |
| **PDB ID** | **λ=1.0** | **λ=1.5** | **λ=2.0** | **λ=1.0** | **λ=1.5** | **λ=2.0** | **λ=1.0** | **λ=1.5** | **λ=2.0** | **λ=1.0** | **λ=1.5** | **λ=2.0** |
| **5EK0** | $1.8•10^{-2}$ | $8.2•10^{-2}$ | 0.19 | $2.5•10^{-2}$ | $8.9•10^{-2}$ | 0.18 | $1.0•10^{-2}$ | $7.2•10^{-2}$ | 0.20 | $2.6•10^{-2}$ | 0.10 | 0.22 |
| **6J8G** | $4.2•10^{-3}$ | $1.0•10^{-2}$ | $2.5•10^{-2}$ | $2.8•10^{-3}$ | $9.0•10^{-3}$ | $2.4•10^{-2}$ | $1.6•10^{-3}$ | $3.2•10^{-3}$ | $1.4•10^{-2}$ | $5.2•10^{-4}$ | $1.8•10^{-3}$ | $1.2•10^{-2}$ |
| **6J8I** | $5.0•10^{-2}$ | $7.9•10^{-2}$ | 0.11 | $4.9•10^{-2}$ | $7.7•10^{-2}$ | 0.11 | $5.3•10^{-2}$ | $8.4•10^{-2}$ | 0.12 | $5.0•10^{-2}$ | $8.0•10^{-2}$ | 0.12 |
| **6JP5** | $3.3•10^{-2}$ | $9.4•10^{-2}$ | 0.16 | $3.3•10^{-2}$ | $9.4•10^{-2}$ | 0.16 | $5.0•10^{-2}$ | 0.14 | 0.21 | $5.1•10^{-2}$ | 0.14 | 0.21 |
| **6JP8** | 0.14 | 0.37 | 0.51 | 0.14 | 0.36 | 0.51 | 0.19 | 0.45 | 0.62 | 0.19 | 0.45 | 0.62 |
| **6JPA-1** | $6.3•10^{-5}$ | $8.0•10^{-4}$ | $6.5•10^{-3}$ | $8.7•10^{-5}$ | $9.1•10^{-4}$ | $6.6•10^{-3}$ | $1.3•10^{-5}$ | $3.9•10^{-4}$ | $5.2•10^{-3}$ | $1.9•10^{-5}$ | $4.0•10^{-4}$ | $4.7•10^{-3}$ |
| **6JPA-2** | $3.2•10^{-7}$ | $5.5•10^{-5}$ | $6.3•10^{-4}$ | $3.8•10^{-7}$ | $6.8•10^{-5}$ | $7.0•10^{-4}$ | $1.8•10^{-7}$ | $3.6•10^{-5}$ | $4.5•10^{-4}$ | $2.9•10^{-7}$ | $5.7•10^{-5}$ | $5.0•10^{-4}$ |
| **6JPB** | $1.9•10^{-7}$ | $4.4•10^{-5}$ | $1.1•10^{-3}$ | $8.9•10^{-8}$ | $4.4•10^{-5}$ | $1.1•10^{-3}$ | $5.1•10^{-8}$ | $2.1•10^{-5}$ | $7.7•10^{-4}$ | $3.1•10^{-8}$ | $1.9•10^{-5}$ | $6.8•10^{-4}$ |
| **6KZP** | 0.10 | 0.23 | 0.34 | $9.8•10^{-2}$ | 0.23 | 0.34 | 0.18 | 0.36 | 0.50 | 0.17 | 0.36 | 0.50 |
| **6U88** | $5.0•10^{-3}$ | $2.9•10^{-2}$ | $7.9•10^{-2}$ | $5.3•10^{-3}$ | $3.0•10^{-2}$ | $8.1•10^{-2}$ | $2.5•10^{-3}$ | $2.2•10^{-2}$ | $7.0•10^{-2}$ | $3.9•10^{-3}$ | $2.9•10^{-2}$ | $8.3•10^{-2}$ |
| **6UZ0** | $7.4•10^{-3}$ | $5.3•10^{-2}$ | 0.14 | $7.4•10^{-3}$ | $5.3•10^{-2}$ | 0.14 | $8.1•10^{-3}$ | $5.8•10^{-2}$ | 0.15 | $8.0•10^{-3}$ | $5.6•10^{-2}$ | 0.14 |

**Table S4.2.** $P_{Near}$ of RMSD vs `interface_delta_X` for RosettaLigand, 10% lowest poses by `total_score`, all ligand atom mode.

| | $k_BT$=1.0 | | | | | | $k_BT$=0.62 | | | | | |
|---|---|---|---|---|---|---|---|---|---|---|---|---|
| | **20,000 poses** | | | **100,000 poses** | | | **20,000 poses** | | | **100,000 poses** | | |
| **PDB ID** | **λ=1.0** | **λ=1.5** | **λ=2.0** | **λ=1.0** | **λ=1.5** | **λ=2.0** | **λ=1.0** | **λ=1.5** | **λ=2.0** | **λ=1.0** | **λ=1.5** | **λ=2.0** |
| **5EK0** | $2.1\cdot10^{-2}$ | $9.5\cdot10^{-2}$ | 0.21 | $4.1\cdot10^{-2}$ | 0.13 | 0.24 | $1.0\cdot10^{-2}$ | $7.3\cdot10^{-2}$ | 0.20 | $4.0\cdot10^{-2}$ | 0.13 | 0.26 |
| **6J8G** | $1.4\cdot10^{-4}$ | $1.3\cdot10^{-3}$ | $1.3\cdot10^{-2}$ | $3.7\cdot10^{-4}$ | $2.1\cdot10^{-3}$ | $1.4\cdot10^{-2}$ | $1.3\cdot10^{-5}$ | $4.0\cdot10^{-4}$ | $1.0\cdot10^{-2}$ | $9.1\cdot10^{-5}$ | $6.8\cdot10^{-4}$ | $9.3\cdot10^{-3}$ |
| **6J8I** | $3.0\cdot10^{-2}$ | $4.7\cdot10^{-2}$ | $7.1\cdot10^{-2}$ | $3.2\cdot10^{-2}$ | $5.1\cdot10^{-2}$ | $7.8\cdot10^{-2}$ | $2.9\cdot10^{-2}$ | $4.7\cdot10^{-2}$ | $7.5\cdot10^{-2}$ | $2.7\cdot10^{-2}$ | $4.4\cdot10^{-2}$ | $7.8\cdot10^{-2}$ |
| **6JP5** | $6.4\cdot10^{-2}$ | 0.17 | 0.25 | $6.2\cdot10^{-2}$ | 0.16 | 0.24 | $7.5\cdot10^{-2}$ | 0.19 | 0.28 | $7.6\cdot10^{-2}$ | 0.19 | 0.28 |
| **6JP8** | 0.18 | 0.45 | 0.62 | 0.18 | 0.45 | 0.62 | 0.20 | 0.48 | 0.66 | 0.20 | 0.48 | 0.66 |
| **6JPA-1** | $2.3\cdot10^{-5}$ | $5.6\cdot10^{-4}$ | $6.2\cdot10^{-3}$ | $5.9\cdot10^{-5}$ | $6.5\cdot10^{-4}$ | $5.9\cdot10^{-3}$ | $3.1\cdot10^{-6}$ | $3.1\cdot10^{-4}$ | $5.0\cdot10^{-3}$ | $1.0\cdot10^{-5}$ | $2.9\cdot10^{-4}$ | $4.3\cdot10^{-3}$ |
| **6JPA-2** | $5.2\cdot10^{-9}$ | $1.3\cdot10^{-5}$ | $2.6\cdot10^{-4}$ | $6.5\cdot10^{-9}$ | $9.7\cdot10^{-6}$ | $2.1\cdot10^{-4}$ | $3.2\cdot10^{-9}$ | $9.6\cdot10^{-6}$ | $2.0\cdot10^{-4}$ | $2.0\cdot10^{-9}$ | $5.7\cdot10^{-6}$ | $1.2\cdot10^{-4}$ |
| **6JPB** | $3.7\cdot10^{-8}$ | $3.2\cdot10^{-5}$ | $9.2\cdot10^{-4}$ | $5.3\cdot10^{-8}$ | $3.3\cdot10^{-5}$ | $9.6\cdot10^{-4}$ | $1.2\cdot10^{-8}$ | $1.5\cdot10^{-5}$ | $6.6\cdot10^{-4}$ | $1.6\cdot10^{-8}$ | $1.4\cdot10^{-5}$ | $6.4\cdot10^{-4}$ |
| **6KZP** | 0.19 | 0.40 | 0.55 | 0.18 | 0.39 | 0.54 | 0.25 | 0.48 | 0.63 | 0.24 | 0.47 | 0.62 |
| **6U88** | $3.0\cdot10^{-4}$ | $9.4\cdot10^{-3}$ | $4.2\cdot10^{-2}$ | $4.8\cdot10^{-4}$ | $8.9\cdot10^{-3}$ | $3.9\cdot10^{-2}$ | $2.7\cdot10^{-4}$ | $1.0\cdot10^{-2}$ | $4.6\cdot10^{-2}$ | $4.1\cdot10^{-4}$ | $9.8\cdot10^{-3}$ | $4.3\cdot10^{-2}$ |
| **6UZ0** | $6.9\cdot10^{-3}$ | $4.9\cdot10^{-2}$ | 0.13 | $4.5\cdot10^{-3}$ | $4.1\cdot10^{-2}$ | 0.11 | $7.4\cdot10^{-3}$ | $5.3\cdot10^{-2}$ | 0.14 | $4.9\cdot10^{-3}$ | $4.5\cdot10^{-2}$ | 0.12 |

**Table S4.3.** $P_{Near}$ of RMSD vs `interface_delta_X` for RosettaLigand, all poses, neighbor ligand atom mode.

| | $k_BT$=1.0 | | | | | | $k_BT$=0.62 | | | | | |
|---|---|---|---|---|---|---|---|---|---|---|---|---|
| | 20,000 poses | | | 100,000 poses | | | 20,000 poses | | | 100,000 poses | | |
| **PDB ID** | **λ=1.0** | **λ=1.5** | **λ=2.0** | **λ=1.0** | **λ=1.5** | **λ=2.0** | **λ=1.0** | **λ=1.5** | **λ=2.0** | **λ=1.0** | **λ=1.5** | **λ=2.0** |
| **5EK0** | $9.2\cdot10^{-3}$ | $6.0\cdot10^{-2}$ | 0.16 | $1.7\cdot10^{-2}$ | $7.5\cdot10^{-2}$ | 0.17 | $5.5\cdot10^{-3}$ | $6.2\cdot10^{-2}$ | 0.19 | $1.0\cdot10^{-2}$ | $6.9\cdot10^{-2}$ | 0.19 |
| **6J8G** | $5.6\cdot10^{-3}$ | $1.5\cdot10^{-2}$ | $3.3\cdot10^{-2}$ | $4.0\cdot10^{-3}$ | $1.1\cdot10^{-2}$ | $2.7\cdot10^{-2}$ | $1.1\cdot10^{-3}$ | $3.1\cdot10^{-3}$ | $1.5\cdot10^{-2}$ | $7.4\cdot10^{-4}$ | $2.0\cdot10^{-3}$ | $1.2\cdot10^{-2}$ |
| **6J8I** | $6.1\cdot10^{-2}$ | $9.4\cdot10^{-2}$ | 0.12 | $5.9\cdot10^{-2}$ | $9.0\cdot10^{-2}$ | 0.12 | $6.8\cdot10^{-2}$ | 0.10 | 0.14 | $5.9\cdot10^{-2}$ | $9.2\cdot10^{-2}$ | 0.13 |
| **6JP5** | $2.5\cdot10^{-2}$ | $7.6\cdot10^{-2}$ | 0.13 | $2.6\cdot10^{-2}$ | $7.7\cdot10^{-2}$ | 0.13 | $3.6\cdot10^{-2}$ | 0.11 | 0.18 | $4.1\cdot10^{-2}$ | 0.11 | 0.18 |
| **6JP8** | 0.14 | 0.36 | 0.51 | 0.15 | 0.37 | 0.51 | 0.19 | 0.46 | 0.62 | 0.19 | 0.46 | 0.63 |
| **6JPA-1** | $2.2\cdot10^{-4}$ | $1.4\cdot10^{-3}$ | $6.7\cdot10^{-3}$ | $1.6\cdot10^{-4}$ | $1.1\cdot10^{-3}$ | $5.8\cdot10^{-3}$ | $8.7\cdot10^{-5}$ | $6.8\cdot10^{-4}$ | $4.2\cdot10^{-3}$ | $5.3\cdot10^{-5}$ | $4.8\cdot10^{-4}$ | $3.5\cdot10^{-3}$ |
| **6JPA-2** | $5.8\cdot10^{-8}$ | $2.2\cdot10^{-5}$ | $3.6\cdot10^{-4}$ | $2.1\cdot10^{-7}$ | $2.8\cdot10^{-5}$ | $3.8\cdot10^{-4}$ | $1.3\cdot10^{-8}$ | $9.6\cdot10^{-6}$ | $2.0\cdot10^{-4}$ | $3.7\cdot10^{-8}$ | $8.5\cdot10^{-6}$ | $1.7\cdot10^{-4}$ |
| **6JPB** | $1.0\cdot10^{-7}$ | $3.8\cdot10^{-5}$ | $1.0\cdot10^{-3}$ | $2.8\cdot10^{-7}$ | $4.7\cdot10^{-5}$ | $1.1\cdot10^{-3}$ | $2.6\cdot10^{-8}$ | $1.5\cdot10^{-5}$ | $5.8\cdot10^{-4}$ | $7.7\cdot10^{-8}$ | $1.9\cdot10^{-5}$ | $6.3\cdot10^{-4}$ |
| **6KZP** | $6.2\cdot10^{-2}$ | 0.17 | 0.28 | $5.7\cdot10^{-2}$ | 0.16 | 0.27 | 0.11 | 0.26 | 0.40 | $9.4\cdot10^{-2}$ | 0.24 | 0.37 |
| **6U88** | $1.2\cdot10^{-3}$ | $1.9\cdot10^{-2}$ | $6.5\cdot10^{-2}$ | $1.4\cdot10^{-3}$ | $2.0\cdot10^{-2}$ | $6.5\cdot10^{-2}$ | $1.7\cdot10^{-3}$ | $2.7\cdot10^{-2}$ | $7.9\cdot10^{-2}$ | $2.0\cdot10^{-3}$ | $2.5\cdot10^{-2}$ | $7.0\cdot10^{-2}$ |
| **6UZ0** | $7.4\cdot10^{-3}$ | $5.0\cdot10^{-2}$ | 0.13 | $6.9\cdot10^{-3}$ | $4.8\cdot10^{-2}$ | 0.13 | $8.0\cdot10^{-3}$ | $5.3\cdot10^{-2}$ | 0.13 | $7.2\cdot10^{-3}$ | $4.9\cdot10^{-2}$ | 0.13 |

**Table S4.4.** $P_{Near}$ of RMSD vs `interface_delta_X` for RosettaLigand, 10% lowest poses by `total_score`, neighbor atom mode.

| | $k_BT$=1.0 | | | | | | $k_BT$=0.62 | | | | | |
|---|---|---|---|---|---|---|---|---|---|---|---|---|
| | 20,000 poses | | | 100,000 poses | | | 20,000 poses | | | 100,000 poses | | |
| **PDB ID** | **λ=1.0** | **λ=1.5** | **λ=2.0** | **λ=1.0** | **λ=1.5** | **λ=2.0** | **λ=1.0** | **λ=1.5** | **λ=2.0** | **λ=1.0** | **λ=1.5** | **λ=2.0** |
| **5EK0** | $1.5\bullet10^{-2}$ | $7.8\bullet10^{-2}$ | 0.18 | $3.1\bullet10^{-2}$ | 0.11 | 0.22 | $7.4\bullet10^{-3}$ | $6.8\bullet10^{-2}$ | 0.20 | $1.6\bullet10^{-2}$ | $8.2\bullet10^{-2}$ | 0.21 |
| **6J8G** | $1.4\bullet10^{-3}$ | $4.1\bullet10^{-3}$ | $1.6\bullet10^{-2}$ | $5.5\bullet10^{-4}$ | $2.2\bullet10^{-3}$ | $1.3\bullet10^{-2}$ | $4.0\bullet10^{-4}$ | $1.3\bullet10^{-3}$ | $1.2\bullet10^{-2}$ | $9.1\bullet10^{-5}$ | $5.8\bullet10^{-4}$ | $1.0\bullet10^{-2}$ |
| **6J8I** | $1.1\bullet10^{-2}$ | $2.0\bullet10^{-2}$ | $3.8\bullet10^{-2}$ | $5.5\bullet10^{-3}$ | $1.1\bullet10^{-2}$ | $2.7\bullet10^{-2}$ | $1.2\bullet10^{-2}$ | $2.3\bullet10^{-2}$ | $5.0\bullet10^{-2}$ | $4.5\bullet10^{-3}$ | $1.2\bullet10^{-2}$ | $3.8\bullet10^{-2}$ |
| **6JP5** | $4.5\bullet10^{-2}$ | 0.13 | 0.21 | $4.8\bullet10^{-2}$ | 0.13 | 0.20 | $6.3\bullet10^{-2}$ | 0.18 | 0.28 | $7.0\bullet10^{-2}$ | 0.18 | 0.27 |
| **6JP8** | 0.18 | 0.45 | 0.62 | 0.18 | 0.45 | 0.62 | 0.20 | 0.49 | 0.66 | 0.20 | 0.48 | 0.66 |
| **6JPA-1** | $7.3\bullet10^{-6}$ | $3.8\bullet10^{-4}$ | $5.1\bullet10^{-3}$ | $7.6\bullet10^{-6}$ | $3.3\bullet10^{-4}$ | $4.5\bullet10^{-3}$ | $1.5\bullet10^{-6}$ | $1.9\bullet10^{-4}$ | $3.2\bullet10^{-3}$ | $1.4\bullet10^{-6}$ | $1.7\bullet10^{-4}$ | $2.8\bullet10^{-3}$ |
| **6JPA-2** | $4.4\bullet10^{-9}$ | $9.9\bullet10^{-6}$ | $2.5\bullet10^{-4}$ | $2.9\bullet10^{-8}$ | $1.3\bullet10^{-5}$ | $2.6\bullet10^{-4}$ | $1.6\bullet10^{-9}$ | $6.8\bullet10^{-6}$ | $1.7\bullet10^{-4}$ | $4.8\bullet10^{-9}$ | $5.1\bullet10^{-6}$ | $1.3\bullet10^{-4}$ |
| **6JPB** | $1.6\bullet10^{-8}$ | $1.9\bullet10^{-5}$ | $6.4\bullet10^{-4}$ | $8.8\bullet10^{-8}$ | $2.6\bullet10^{-5}$ | $6.9\bullet10^{-4}$ | $2.7\bullet10^{-9}$ | $5.8\bullet10^{-6}$ | $3.0\bullet10^{-4}$ | $2.9\bullet10^{-8}$ | $1.1\bullet10^{-5}$ | $4.1\bullet10^{-4}$ |
| **6KZP** | $7.1\bullet10^{-2}$ | 0.19 | 0.31 | $7.9\bullet10^{-2}$ | 0.20 | 0.31 | 0.11 | 0.27 | 0.41 | 0.12 | 0.26 | 0.38 |
| **6U88** | $6.4\bullet10^{-5}$ | $4.7\bullet10^{-3}$ | $2.6\bullet10^{-2}$ | $4.4\bullet10^{-5}$ | $3.3\bullet10^{-3}$ | $2.0\bullet10^{-2}$ | $3.6\bullet10^{-5}$ | $3.1\bullet10^{-3}$ | $1.9\bullet10^{-2}$ | $1.2\bullet10^{-5}$ | $1.2\bullet10^{-3}$ | $8.5\bullet10^{-3}$ |
| **6UZ0** | $1.3\bullet10^{-3}$ | $2.2\bullet10^{-2}$ | $8.6\bullet10^{-2}$ | $1.6\bullet10^{-3}$ | $2.3\bullet10^{-2}$ | $8.7\bullet10^{-2}$ | $1.3\bullet10^{-3}$ | $2.0\bullet10^{-2}$ | $7.7\bullet10^{-2}$ | $1.9\bullet10^{-3}$ | $2.1\bullet10^{-2}$ | $7.4\bullet10^{-2}$ |

**Table S4.5.** $P_{Near}$ of RMSD vs `LigInterface` for GALigandDock, padding 2 Å. Full population of poses.

| | $k_BT$=1.0 | | | | | | $k_BT$=0.62 | | | | | |
|---|---|---|---|---|---|---|---|---|---|---|---|---|
| | 20,000 poses | | | 100,000 poses | | | 20,000 poses | | | 100,000 poses | | |
| **PDB ID** | **λ=1.0** | **λ=1.5** | **λ=2.0** | **λ=1.0** | **λ=1.5** | **λ=2.0** | **λ=1.0** | **λ=1.5** | **λ=2.0** | **λ=1.0** | **λ=1.5** | **λ=2.0** |
| **5EK0** | 0.46 | 0.63 | 0.71 | 0.42 | 0.60 | 0.68 | 0.53 | 0.71 | 0.78 | 0.46 | 0.63 | 0.72 |
| **6J8G** | $9.1\bullet10^{-5}$ | $8.4\bullet10^{-4}$ | $1.1\bullet10^{-2}$ | $8.2\bullet10^{-5}$ | $1.0\bullet10^{-3}$ | $1.2\bullet10^{-2}$ | $1.8\bullet10^{-6}$ | $6.5\bullet10^{-4}$ | $1.4\bullet10^{-2}$ | $3.3\bullet10^{-6}$ | $8.9\bullet10^{-4}$ | $1.5\bullet10^{-2}$ |
| **6J8I** | $4.6\bullet10^{-5}$ | $2.3\bullet10^{-4}$ | $1.3\bullet10^{-3}$ | $6.7\bullet10^{-5}$ | $2.9\bullet10^{-4}$ | $1.3\bullet10^{-3}$ | $1.3\bullet10^{-7}$ | $1.3\bullet10^{-6}$ | $1.2\bullet10^{-4}$ | $4.8\bullet10^{-7}$ | $2.9\bullet10^{-6}$ | $1.7\bullet10^{-4}$ |
| **6JP5** | $3.1\bullet10^{-6}$ | $4.7\bullet10^{-4}$ | $9.9\bullet10^{-3}$ | $3.1\bullet10^{-6}$ | $4.7\bullet10^{-4}$ | $9.9\bullet10^{-3}$ | $5.7\bullet10^{-7}$ | $4.9\bullet10^{-4}$ | $1.1\bullet10^{-2}$ | $3.7\bullet10^{-7}$ | $4.9\bullet10^{-4}$ | $1.1\bullet10^{-2}$ |
| **6JP8** | $1.2\bullet10^{-5}$ | $1.9\bullet10^{-4}$ | $2.4\bullet10^{-3}$ | $1.7\bullet10^{-5}$ | $2.0\bullet10^{-4}$ | $2.4\bullet10^{-3}$ | $4.3\bullet10^{-7}$ | $4.7\bullet10^{-5}$ | $1.6\bullet10^{-3}$ | $1.6\bullet10^{-6}$ | $4.9\bullet10^{-5}$ | $1.5\bullet10^{-3}$ |
| **6JPA-1** | $5.0\bullet10^{-7}$ | $1.5\bullet10^{-4}$ | $3.9\bullet10^{-3}$ | $5.3\bullet10^{-7}$ | $1.4\bullet10^{-4}$ | $3.1\bullet10^{-3}$ | $3.0\bullet10^{-8}$ | $8.1\bullet10^{-5}$ | $2.9\bullet10^{-3}$ | $1.9\bullet10^{-8}$ | $2.3\bullet10^{-5}$ | $1.2\bullet10^{-3}$ |
| **6JPA-2** | $6.3\bullet10^{-10}$ | $5.2\bullet10^{-7}$ | $1.6\bullet10^{-5}$ | $6.6\bullet10^{-10}$ | $9.7\bullet10^{-7}$ | $2.8\bullet10^{-5}$ | $2.3\bullet10^{-12}$ | $7.2\bullet10^{-9}$ | $7.5\bullet10^{-7}$ | $5.6\bullet10^{-12}$ | $2.4\bullet10^{-8}$ | $1.5\bullet10^{-6}$ |
| **6JPB** | $8.7\bullet10^{-7}$ | $6.4\bullet10^{-5}$ | $2.6\bullet10^{-3}$ | $9.5\bullet10^{-7}$ | $6.2\bullet10^{-5}$ | $2.7\bullet10^{-3}$ | $9.1\bullet10^{-9}$ | $4.1\bullet10^{-5}$ | $3.2\bullet10^{-3}$ | $8.9\bullet10^{-9}$ | $4.3\bullet10^{-5}$ | $3.3\bullet10^{-3}$ |
| **6KZP** | $2.7\bullet10^{-2}$ | $8.3\bullet10^{-2}$ | 0.15 | $3.1\bullet10^{-2}$ | $9.3\bullet10^{-2}$ | 0.16 | $2.4\bullet10^{-2}$ | $8.1\bullet10^{-2}$ | 0.15 | $3.1\bullet10^{-2}$ | $8.7\bullet10^{-2}$ | 0.15 |
| **6U88** | $7.6\bullet10^{-4}$ | $1.1\bullet10^{-2}$ | $4.4\bullet10^{-2}$ | $1.1\bullet10^{-3}$ | $1.2\bullet10^{-2}$ | $4.5\bullet10^{-2}$ | $1.2\bullet10^{-4}$ | $4.8\bullet10^{-3}$ | $2.5\bullet10^{-2}$ | $9.5\bullet10^{-4}$ | $8.9\bullet10^{-3}$ | $3.2\bullet10^{-2}$ |
| **6UZ0** | $2.0\bullet10^{-5}$ | $3.3\bullet10^{-4}$ | $2.1\bullet10^{-3}$ | $2.5\bullet10^{-5}$ | $3.3\bullet10^{-4}$ | $1.9\bullet10^{-3}$ | $1.8\bullet10^{-6}$ | $7.1\bullet10^{-5}$ | $1.6\bullet10^{-3}$ | $2.1\bullet10^{-6}$ | $5.1\bullet10^{-5}$ | $1.3\bullet10^{-3}$ |

**Table S4.6.** $P_{Near}$ of RMSD vs `LigInterface` for GALigandDock, padding 2 Å. Lowest 10% of poses by `total_score`.

| | $k_BT$=1.0 | | | | | | $k_BT$=0.62 | | | | | |
|---|---|---|---|---|---|---|---|---|---|---|---|---|
| | 20,000 poses | | | 100,000 poses | | | 20,000 poses | | | 100,000 poses | | |
| **PDB ID** | λ=1.0 | λ=1.5 | λ=2.0 | λ=1.0 | λ=1.5 | λ=2.0 | λ=1.0 | λ=1.5 | λ=2.0 | λ=1.0 | λ=1.5 | λ=2.0 |
| **5EK0** | 0.53 | 0.73 | 0.83 | 0.51 | 0.71 | 0.81 | 0.58 | 0.78 | 0.87 | 0.55 | 0.75 | 0.85 |
| **6J8G** | $7.2\bullet10^{-8}$ | $4.0\bullet10^{-4}$ | $9.9\bullet10^{-3}$ | $9.2\bullet10^{-8}$ | $4.1\bullet10^{-4}$ | $9.9\bullet10^{-3}$ | $1.2\bullet10^{-7}$ | $6.1\bullet10^{-4}$ | $1.3\bullet10^{-2}$ | $1.8\bullet10^{-7}$ | $6.9\bullet10^{-4}$ | $1.4\bullet10^{-2}$ |
| **6J8I** | $6.0\bullet10^{-7}$ | $5.7\bullet10^{-5}$ | $8.5\bullet10^{-4}$ | $8.5\bullet10^{-7}$ | $5.4\bullet10^{-5}$ | $8.0\bullet10^{-4}$ | $3.9\bullet10^{-9}$ | $8.1\bullet10^{-7}$ | $1.1\bullet10^{-4}$ | $7.8\bullet10^{-9}$ | $1.3\bullet10^{-6}$ | $1.7\bullet10^{-4}$ |
| **6JP5** | $1.9\bullet10^{-6}$ | $1.4\bullet10^{-4}$ | $4.2\bullet10^{-3}$ | $8.7\bullet10^{-7}$ | $9.4\bullet10^{-5}$ | $3.9\bullet10^{-3}$ | $7.0\bullet10^{-7}$ | $1.1\bullet10^{-4}$ | $4.7\bullet10^{-3}$ | $3.0\bullet10^{-7}$ | $8.8\bullet10^{-5}$ | $4.4\bullet10^{-3}$ |
| **6JP8** | $1.3\bullet10^{-9}$ | $3.4\bullet10^{-5}$ | $1.7\bullet10^{-3}$ | $1.2\bullet10^{-9}$ | $3.1\bullet10^{-5}$ | $1.6\bullet10^{-3}$ | $1.3\bullet10^{-9}$ | $3.1\bullet10^{-5}$ | $1.5\bullet10^{-3}$ | $1.2\bullet10^{-9}$ | $2.9\bullet10^{-5}$ | $1.4\bullet10^{-3}$ |
| **6JPA-1** | $2.8\bullet10^{-7}$ | $1.4\bullet10^{-4}$ | $3.9\bullet10^{-3}$ | $4.2\bullet10^{-7}$ | $1.2\bullet10^{-4}$ | $2.9\bullet10^{-3}$ | $2.5\bullet10^{-8}$ | $8.1\bullet10^{-5}$ | $2.9\bullet10^{-3}$ | $1.7\bullet10^{-8}$ | $2.1\bullet10^{-5}$ | $1.2\bullet10^{-3}$ |
| **6JPA-2** | $1.6\bullet10^{-11}$ | $1.5\bullet10^{-7}$ | $9.1\bullet10^{-6}$ | $1.7\bullet10^{-10}$ | $5.9\bullet10^{-7}$ | $1.9\bullet10^{-5}$ | $2.5\bullet10^{-13}$ | $4.2\bullet10^{-9}$ | $7.0\bullet10^{-7}$ | $2.8\bullet10^{-12}$ | $1.9\bullet10^{-8}$ | $1.4\bullet10^{-6}$ |
| **6JPB** | $1.6\bullet10^{-8}$ | $3.9\bullet10^{-5}$ | $2.8\bullet10^{-3}$ | $1.1\bullet10^{-8}$ | $4.1\bullet10^{-5}$ | $2.9\bullet10^{-3}$ | $4.0\bullet10^{-10}$ | $4.2\bullet10^{-5}$ | $3.3\bullet10^{-3}$ | $3.7\bullet10^{-10}$ | $4.5\bullet10^{-5}$ | $3.5\bullet10^{-3}$ |
| **6KZP** | $4.8\bullet10^{-2}$ | 0.14 | 0.25 | $5.7\bullet10^{-2}$ | 0.17 | 0.28 | $4.5\bullet10^{-2}$ | 0.15 | 0.27 | $7.9\bullet10^{-2}$ | 0.22 | 0.36 |
| **6U88** | $4.1\bullet10^{-4}$ | $9.3\bullet10^{-3}$ | $4.3\bullet10^{-2}$ | $4.8\bullet10^{-4}$ | $9.2\bullet10^{-3}$ | $4.2\bullet10^{-2}$ | $7.8\bullet10^{-5}$ | $4.6\bullet10^{-3}$ | $2.4\bullet10^{-2}$ | $9.8\bullet10^{-5}$ | $4.7\bullet10^{-3}$ | $2.5\bullet10^{-2}$ |
| **6UZ0** | $9.6\bullet10^{-7}$ | $1.6\bullet10^{-4}$ | $2.1\bullet10^{-3}$ | $2.9\bullet10^{-6}$ | $1.1\bullet10^{-4}$ | $1.7\bullet10^{-3}$ | $2.5\bullet10^{-7}$ | $6.6\bullet10^{-5}$ | $2.0\bullet10^{-3}$ | $2.9\bullet10^{-7}$ | $3.5\bullet10^{-5}$ | $1.6\bullet10^{-3}$ |

**Table S4.7.** $P_{Near}$ of RMSD vs `LigInterface` for GALigandDock, padding 4 Å. Full population of poses.

| | $k_BT$=1.0 | | | | | | $k_BT$=0.62 | | | | | |
|---|---|---|---|---|---|---|---|---|---|---|---|---|
| | 20,000 poses | | | 100,000 poses | | | 20,000 poses | | | 100,000 poses | | |
| **PDB ID** | **λ=1.0** | **λ=1.5** | **λ=2.0** | **λ=1.0** | **λ=1.5** | **λ=2.0** | **λ=1.0** | **λ=1.5** | **λ=2.0** | **λ=1.0** | **λ=1.5** | **λ=2.0** |
| **5EK0** | 0.53 | 0.72 | 0.81 | 0.45 | 0.64 | 0.73 | 0.59 | 0.78 | 0.87 | 0.56 | 0.75 | 0.83 |
| **6J8G** | $5.5\bullet10^{-5}$ | $9.4\bullet10^{-4}$ | $1.3\bullet10^{-2}$ | $1.9\bullet10^{-4}$ | $1.5\bullet10^{-3}$ | $1.7\bullet10^{-2}$ | $2.3\bullet10^{-6}$ | $1.0\bullet10^{-3}$ | $1.8\bullet10^{-2}$ | $5.7\bullet10^{-6}$ | $1.5\bullet10^{-3}$ | $2.5\bullet10^{-2}$ |
| **6J8I** | $5.2\bullet10^{-3}$ | $6.3\bullet10^{-2}$ | 0.16 | $4.1\bullet10^{-3}$ | $5.8\bullet10^{-2}$ | 0.15 | $3.2\bullet10^{-3}$ | $4.5\bullet10^{-2}$ | 0.12 | $3.3\bullet10^{-3}$ | $5.1\bullet10^{-2}$ | 0.14 |
| **6JP5** | $1.7\bullet10^{-2}$ | $5.2\bullet10^{-2}$ | $9.1\bullet10^{-2}$ | $1.7\bullet10^{-2}$ | $5.4\bullet10^{-2}$ | $9.5\bullet10^{-2}$ | $4.7\bullet10^{-2}$ | 0.13 | 0.20 | $5.1\bullet10^{-2}$ | 0.15 | 0.22 |
| **6JP8** | 0.15 | 0.30 | 0.39 | 0.14 | 0.29 | 0.38 | 0.24 | 0.48 | 0.62 | 0.23 | 0.47 | 0.61 |
| **6JPA-1** | $2.1\bullet10^{-7}$ | $1.4\bullet10^{-4}$ | $3.4\bullet10^{-3}$ | $7.9\bullet10^{-7}$ | $1.4\bullet10^{-4}$ | $3.0\bullet10^{-3}$ | $1.1\bullet10^{-8}$ | $5.6\bullet10^{-5}$ | $2.8\bullet10^{-3}$ | $3.1\bullet10^{-8}$ | $3.3\bullet10^{-5}$ | $1.3\bullet10^{-3}$ |
| **6JPA-2** | $1.2\bullet10^{-9}$ | $1.0\bullet10^{-6}$ | $2.6\bullet10^{-5}$ | $7.2\bullet10^{-10}$ | $7.0\bullet10^{-7}$ | $1.7\bullet10^{-5}$ | $8.0\bullet10^{-12}$ | $3.7\bullet10^{-8}$ | $1.7\bullet10^{-6}$ | $5.0\bullet10^{-12}$ | $7.3\bullet10^{-9}$ | $4.1\bullet10^{-7}$ |
| **6JPB** | $1.6\bullet10^{-7}$ | $1.2\bullet10^{-5}$ | $7.7\bullet10^{-4}$ | $1.3\bullet10^{-7}$ | $1.3\bullet10^{-5}$ | $8.0\bullet10^{-4}$ | $1.6\bullet10^{-10}$ | $1.6\bullet10^{-6}$ | $3.8\bullet10^{-4}$ | $1.1\bullet10^{-10}$ | $1.6\bullet10^{-6}$ | $3.8\bullet10^{-4}$ |
| **6KZP** | $1.4\bullet10^{-2}$ | $4.2\bullet10^{-2}$ | $7.4\bullet10^{-2}$ | $4.8\bullet10^{-3}$ | $1.4\bullet10^{-2}$ | $2.3\bullet10^{-2}$ | $8.0\bullet10^{-3}$ | $2.2\bullet10^{-2}$ | $4.0\bullet10^{-2}$ | $2.6\bullet10^{-4}$ | $6.2\bullet10^{-4}$ | $1.2\bullet10^{-3}$ |
| **6U88** | $4.8\bullet10^{-4}$ | $9.2\bullet10^{-3}$ | $4.2\bullet10^{-2}$ | $3.7\bullet10^{-4}$ | $6.6\bullet10^{-3}$ | $3.0\bullet10^{-2}$ | $7.8\bullet10^{-5}$ | $4.7\bullet10^{-3}$ | $2.5\bullet10^{-2}$ | $2.9\bullet10^{-5}$ | $1.5\bullet10^{-3}$ | $8.0\bullet10^{-3}$ |
| **6UZ0** | $1.8\bullet10^{-5}$ | $3.6\bullet10^{-4}$ | $1.6\bullet10^{-3}$ | $1.7\bullet10^{-5}$ | $3.3\bullet10^{-4}$ | $1.5\bullet10^{-3}$ | $1.5\bullet10^{-6}$ | $3.7\bullet10^{-5}$ | $6.4\bullet10^{-4}$ | $9.1\bullet10^{-7}$ | $2.3\bullet10^{-5}$ | $4.4\bullet10^{-4}$ |

**Table S4.8.** $P_{Near}$ of RMSD vs `LigInterface` for GALigandDock, padding 4 Å. Lowest 10% of poses by `total_score`.

| | $k_BT$=1.0 | | | | | | $k_BT$=0.62 | | | | | |
|---|---|---|---|---|---|---|---|---|---|---|---|---|
| | **20,000 poses** | | | **100,000 poses** | | | **20,000 poses** | | | **100,000 poses** | | |
| **PDB ID** | **λ=1.0** | **λ=1.5** | **λ=2.0** | **λ=1.0** | **λ=1.5** | **λ=2.0** | **λ=1.0** | **λ=1.5** | **λ=2.0** | **λ=1.0** | **λ=1.5** | **λ=2.0** |
| **5EK0** | 0.55 | 0.75 | 0.84 | 0.48 | 0.67 | 0.77 | 0.59 | 0.79 | 0.88 | 0.57 | 0.76 | 0.85 |
| **6J8G** | $2.2\cdot10^{-7}$ | $6.4\cdot10^{-4}$ | $1.2\cdot10^{-2}$ | $3.5\cdot10^{-7}$ | $9.1\cdot10^{-4}$ | $1.7\cdot10^{-2}$ | $3.1\cdot10^{-7}$ | $1.0\cdot10^{-3}$ | $1.8\cdot10^{-2}$ | $4.7\cdot10^{-7}$ | $1.5\cdot10^{-3}$ | $2.5\cdot10^{-2}$ |
| **6J8I** | $4.3\cdot10^{-3}$ | $5.9\cdot10^{-2}$ | 0.15 | $3.9\cdot10^{-3}$ | $5.7\cdot10^{-2}$ | 0.15 | $3.1\cdot10^{-3}$ | $4.4\cdot10^{-2}$ | 0.12 | $3.3\cdot10^{-3}$ | $5.0\cdot10^{-2}$ | 0.14 |
| **6JP5** | $2.3\cdot10^{-2}$ | $6.8\cdot10^{-2}$ | 0.11 | $2.3\cdot10^{-2}$ | $6.6\cdot10^{-2}$ | 0.11 | $5.4\cdot10^{-2}$ | 0.15 | 0.22 | $5.8\cdot10^{-2}$ | 0.16 | 0.23 |
| **6JP8** | 0.21 | 0.41 | 0.53 | 0.20 | 0.40 | 0.52 | 0.26 | 0.52 | 0.67 | 0.26 | 0.52 | 0.66 |
| **6JPA-1** | $2.1\cdot10^{-7}$ | $1.5\cdot10^{-4}$ | $3.5\cdot10^{-3}$ | $6.1\cdot10^{-7}$ | $1.4\cdot10^{-4}$ | $3.1\cdot10^{-3}$ | $1.6\cdot10^{-8}$ | $5.5\cdot10^{-5}$ | $1.8\cdot10^{-3}$ | $2.6\cdot10^{-8}$ | $3.4\cdot10^{-5}$ | $1.4\cdot10^{-3}$ |
| **6JPA-2** | $1.4\cdot10^{-10}$ | $7.3\cdot10^{-7}$ | $2.0\cdot10^{-5}$ | $1.7\cdot10^{-10}$ | $4.3\cdot10^{-7}$ | $1.3\cdot10^{-5}$ | $5.2\cdot10^{-12}$ | $3.3\cdot10^{-8}$ | $1.5\cdot10^{-6}$ | $1.0\cdot10^{-12}$ | $5.2\cdot10^{-9}$ | $3.8\cdot10^{-7}$ |
| **6JPB** | $8.9\cdot10^{-11}$ | $9.6\cdot10^{-6}$ | $9.2\cdot10^{-4}$ | $1.1\cdot10^{-10}$ | $9.8\cdot10^{-6}$ | $9.4\cdot10^{-4}$ | $4.8\cdot10^{-12}$ | $1.6\cdot10^{-6}$ | $3.9\cdot10^{-4}$ | $4.1\cdot10^{-12}$ | $1.6\cdot10^{-6}$ | $3.9\cdot10^{-4}$ |
| **6KZP** | $2.9\cdot10^{-2}$ | $8.6\cdot10^{-2}$ | 0.15 | $6.4\cdot10^{-3}$ | $1.8\cdot10^{-2}$ | $3.0\cdot10^{-2}$ | $2.3\cdot10^{-2}$ | $6.3\cdot10^{-2}$ | 0.10 | $2.9\cdot10^{-4}$ | $6.6\cdot10^{-4}$ | $1.0\cdot10^{-3}$ |
| **6U88** | $3.7\cdot10^{-4}$ | $1.2\cdot10^{-2}$ | $5.7\cdot10^{-2}$ | $2.4\cdot10^{-4}$ | $8.4\cdot10^{-3}$ | $4.2\cdot10^{-2}$ | $7.8\cdot10^{-5}$ | $6.1\cdot10^{-3}$ | $3.2\cdot10^{-2}$ | $2.9\cdot10^{-5}$ | $2.4\cdot10^{-3}$ | $1.3\cdot10^{-2}$ |
| **6UZ0** | $1.8\cdot10^{-6}$ | $9.8\cdot10^{-5}$ | $1.4\cdot10^{-3}$ | $2.8\cdot10^{-7}$ | $5.7\cdot10^{-5}$ | $1.1\cdot10^{-3}$ | $1.9\cdot10^{-7}$ | $2.3\cdot10^{-5}$ | $1.2\cdot10^{-3}$ | $3.5\cdot10^{-8}$ | $1.5\cdot10^{-5}$ | $9.4\cdot10^{-4}$ |

**Table S4.9.** $P_{Near}$ of RMSD vs `LigInterface` for GALigandDock, padding 7 Å. Full population of poses.

| | $k_BT$=1.0 | | | | | | $k_BT$=0.62 | | | | | |
|---|---|---|---|---|---|---|---|---|---|---|---|---|
| | 20,000 poses | | | 100,000 poses | | | 20,000 poses | | | 100,000 poses | | |
| **PDB ID** | **λ=1.0** | **λ=1.5** | **λ=2.0** | **λ=1.0** | **λ=1.5** | **λ=2.0** | **λ=1.0** | **λ=1.5** | **λ=2.0** | **λ=1.0** | **λ=1.5** | **λ=2.0** |
| **5EK0** | 0.45 | 0.68 | 0.80 | 0.46 | 0.69 | 0.81 | 0.47 | 0.70 | 0.81 | 0.50 | 0.72 | 0.83 |
| **6J8G** | $4.0\cdot10^{-5}$ | $7.1\cdot10^{-4}$ | $1.2\cdot10^{-2}$ | $7.1\cdot10^{-5}$ | $1.2\cdot10^{-3}$ | $1.3\cdot10^{-2}$ | $8.2\cdot10^{-7}$ | $7.3\cdot10^{-4}$ | $1.5\cdot10^{-2}$ | $2.1\cdot10^{-5}$ | $1.1\cdot10^{-3}$ | $1.6\cdot10^{-2}$ |
| **6J8I** | $3.9\cdot10^{-3}$ | $6.5\cdot10^{-2}$ | 0.19 | $5.1\cdot10^{-3}$ | $8.1\cdot10^{-2}$ | 0.22 | $3.5\cdot10^{-3}$ | $7.5\cdot10^{-2}$ | 0.22 | $6.3\cdot10^{-3}$ | $1.0\cdot10^{-1}$ | 0.27 |
| **6JP5** | $6.1\cdot10^{-3}$ | $2.3\cdot10^{-2}$ | $4.7\cdot10^{-2}$ | $1.9\cdot10^{-2}$ | $5.9\cdot10^{-2}$ | 0.10 | $1.3\cdot10^{-2}$ | $4.2\cdot10^{-2}$ | $7.2\cdot10^{-2}$ | $5.8\cdot10^{-2}$ | 0.16 | 0.24 |
| **6JP8** | 0.17 | 0.33 | 0.44 | 0.18 | 0.36 | 0.47 | 0.23 | 0.46 | 0.60 | 0.23 | 0.47 | 0.60 |
| **6JPA-1** | $1.1\cdot10^{-5}$ | $1.0\cdot10^{-3}$ | $9.0\cdot10^{-3}$ | $1.3\cdot10^{-6}$ | $4.2\cdot10^{-4}$ | $5.5\cdot10^{-3}$ | $9.4\cdot10^{-6}$ | $1.0\cdot10^{-3}$ | $9.3\cdot10^{-3}$ | $2.4\cdot10^{-7}$ | $1.6\cdot10^{-4}$ | $2.4\cdot10^{-3}$ |
| **6JPA-2** | $4.5\cdot10^{-10}$ | $4.1\cdot10^{-7}$ | $1.4\cdot10^{-5}$ | $2.9\cdot10^{-10}$ | $3.1\cdot10^{-7}$ | $1.7\cdot10^{-5}$ | $2.0\cdot10^{-12}$ | $7.9\cdot10^{-9}$ | $4.0\cdot10^{-6}$ | $3.8\cdot10^{-13}$ | $8.5\cdot10^{-9}$ | $1.8\cdot10^{-5}$ |
| **6JPB** | $1.8\cdot10^{-8}$ | $8.7\cdot10^{-7}$ | $4.1\cdot10^{-5}$ | $2.1\cdot10^{-8}$ | $8.2\cdot10^{-7}$ | $4.2\cdot10^{-5}$ | $5.4\cdot10^{-12}$ | $2.4\cdot10^{-8}$ | $2.2\cdot10^{-6}$ | $4.6\cdot10^{-12}$ | $2.0\cdot10^{-8}$ | $2.8\cdot10^{-6}$ |
| **6KZP** | $1.4\cdot10^{-3}$ | $3.8\cdot10^{-3}$ | $6.6\cdot10^{-3}$ | $1.2\cdot10^{-3}$ | $3.0\cdot10^{-3}$ | $4.8\cdot10^{-3}$ | $2.9\cdot10^{-5}$ | $1.2\cdot10^{-4}$ | $2.9\cdot10^{-4}$ | $2.5\cdot10^{-5}$ | $6.7\cdot10^{-5}$ | $1.2\cdot10^{-4}$ |
| **6U88** | $3.1\cdot10^{-3}$ | $1.1\cdot10^{-2}$ | $2.9\cdot10^{-2}$ | $3.9\cdot10^{-3}$ | $1.3\cdot10^{-2}$ | $3.2\cdot10^{-2}$ | $2.2\cdot10^{-4}$ | $1.1\cdot10^{-3}$ | $4.1\cdot10^{-3}$ | $3.7\cdot10^{-4}$ | $1.5\cdot10^{-3}$ | $5.0\cdot10^{-3}$ |
| **6UZ0** | $3.9\cdot10^{-6}$ | $3.8\cdot10^{-5}$ | $1.6\cdot10^{-4}$ | $4.6\cdot10^{-6}$ | $3.8\cdot10^{-5}$ | $1.6\cdot10^{-4}$ | $1.7\cdot10^{-8}$ | $3.1\cdot10^{-7}$ | $6.7\cdot10^{-6}$ | $2.2\cdot10^{-8}$ | $2.9\cdot10^{-7}$ | $5.9\cdot10^{-6}$ |

**Table S4.10.** $P_{Near}$ of RMSD vs `LigInterface` for GALigandDock, padding 7 Å. Lowest 10% of poses by `total_score`.

| | $k_BT$=1.0 | | | | | | $k_BT$=0.62 | | | | | |
|---|---|---|---|---|---|---|---|---|---|---|---|---|
| | 20,000 poses | | | 100,000 poses | | | 20,000 poses | | | 100,000 poses | | |
| **PDB ID** | **λ=1.0** | **λ=1.5** | **λ=2.0** | **λ=1.0** | **λ=1.5** | **λ=2.0** | **λ=1.0** | **λ=1.5** | **λ=2.0** | **λ=1.0** | **λ=1.5** | **λ=2.0** |
| **5EK0** | 0.45 | 0.68 | 0.80 | 0.46 | 0.69 | 0.81 | 0.47 | 0.70 | 0.81 | 0.50 | 0.72 | 0.83 |
| **6J8G** | $4.9\cdot10^{-5}$ | $9.1\cdot10^{-4}$ | $1.6\cdot10^{-2}$ | $3.5\cdot10^{-5}$ | $1.2\cdot10^{-3}$ | $1.4\cdot10^{-2}$ | $1.0\cdot10^{-6}$ | $9.1\cdot10^{-4}$ | $1.8\cdot10^{-2}$ | $2.1\cdot10^{-5}$ | $1.2\cdot10^{-3}$ | $1.6\cdot10^{-2}$ |
| **6J8I** | $3.0\cdot10^{-3}$ | $3.5\cdot10^{-2}$ | $1.0\cdot10^{-1}$ | $5.7\cdot10^{-3}$ | $8.3\cdot10^{-2}$ | 0.22 | $1.8\cdot10^{-3}$ | $2.6\cdot10^{-2}$ | $7.9\cdot10^{-2}$ | $7.1\cdot10^{-3}$ | 0.10 | 0.27 |
| **6JP5** | $5.9\cdot10^{-3}$ | $2.5\cdot10^{-2}$ | $5.3\cdot10^{-2}$ | $3.0\cdot10^{-2}$ | $8.8\cdot10^{-2}$ | 0.14 | $1.0\cdot10^{-2}$ | $3.5\cdot10^{-2}$ | $6.3\cdot10^{-2}$ | $7.1\cdot10^{-2}$ | 0.19 | 0.29 |
| **6JP8** | 0.19 | 0.38 | 0.50 | 0.20 | 0.40 | 0.52 | 0.23 | 0.47 | 0.61 | 0.24 | 0.47 | 0.61 |
| **6JPA-1** | $1.4\cdot10^{-5}$ | $1.3\cdot10^{-3}$ | $1.2\cdot10^{-2}$ | $1.5\cdot10^{-6}$ | $4.8\cdot10^{-4}$ | $6.2\cdot10^{-3}$ | $1.8\cdot10^{-5}$ | $1.9\cdot10^{-3}$ | $1.8\cdot10^{-2}$ | $2.9\cdot10^{-7}$ | $1.9\cdot10^{-4}$ | $2.8\cdot10^{-3}$ |
| **6JPA-2** | $1.7\cdot10^{-10}$ | $2.1\cdot10^{-7}$ | $9.5\cdot10^{-6}$ | $6.0\cdot10^{-11}$ | $2.1\cdot10^{-7}$ | $1.5\cdot10^{-5}$ | $1.3\cdot10^{-12}$ | $5.2\cdot10^{-9}$ | $4.0\cdot10^{-6}$ | $1.8\cdot10^{-13}$ | $8.3\cdot10^{-9}$ | $1.9\cdot10^{-5}$ |
| **6JPB** | $1.9\cdot10^{-10}$ | $9.5\cdot10^{-7}$ | $6.4\cdot10^{-5}$ | $4.9\cdot10^{-11}$ | $9.9\cdot10^{-7}$ | $8.0\cdot10^{-5}$ | $1.5\cdot10^{-12}$ | $3.7\cdot10^{-8}$ | $3.3\cdot10^{-6}$ | $4.6\cdot10^{-13}$ | $5.9\cdot10^{-8}$ | $7.8\cdot10^{-6}$ |
| **6KZP** | $2.0\cdot10^{-3}$ | $5.4\cdot10^{-3}$ | $9.1\cdot10^{-3}$ | $1.4\cdot10^{-3}$ | $3.5\cdot10^{-3}$ | $5.6\cdot10^{-3}$ | $3.3\cdot10^{-5}$ | $1.4\cdot10^{-4}$ | $3.2\cdot10^{-4}$ | $2.4\cdot10^{-5}$ | $6.6\cdot10^{-5}$ | $1.1\cdot10^{-4}$ |
| **6U88** | $2.4\cdot10^{-4}$ | $4.7\cdot10^{-3}$ | $2.3\cdot10^{-2}$ | $1.7\cdot10^{-4}$ | $4.4\cdot10^{-3}$ | $2.3\cdot10^{-2}$ | $9.9\cdot10^{-6}$ | $6.2\cdot10^{-4}$ | $3.4\cdot10^{-3}$ | $8.4\cdot10^{-6}$ | $6.8\cdot10^{-4}$ | $3.8\cdot10^{-3}$ |
| **6UZ0** | $5.8\cdot10^{-6}$ | $7.5\cdot10^{-5}$ | $4.0\cdot10^{-4}$ | $7.2\cdot10^{-6}$ | $8.2\cdot10^{-5}$ | $4.2\cdot10^{-4}$ | $3.9\cdot10^{-8}$ | $9.2\cdot10^{-7}$ | $2.1\cdot10^{-5}$ | $7.4\cdot10^{-8}$ | $1.3\cdot10^{-6}$ | $2.9\cdot10^{-5}$ |

**Table S5.** $P_{Near}$ central pore cavity cases subclassified by position. Values reflect the primary text for RosettaLigand (all ligand atom mode with 100,000 output poses) and GALigandDock (padding=7 Å with 100,000 output poses), $k_BT$=0.62, λ=2.0.

| Sub-classification | PDB ID | Rotatable bonds | $P_{Near}$ | | RMSD (Å) | |
|---|---|---|---|---|---|---|
| | | | RosettaLigand | GALigandDock | RosettaLigand | GALigandDock |
| Fenestration | 6JP5 | 5 | 0.28 | 0.29 | 0.77 | 1.3 |
| | 6JP8 | 3 | 0.66 | 0.61 | 0.93 | 0.97 |
| | 6U88 | 6 | $4.3 \cdot 10^{-2}$ | $3.8 \cdot 10^{-3}$ | 1.0 | 0.90 |
| Pore and fenestration | 6JPA, position 1 | 13 | $4.3 \cdot 10^{-3}$ | $2.8 \cdot 10^{-3}$ | 1.4 | 2.0 |
| | 6KZP | 6 | 0.62 | $1.1 \cdot 10^{-4}$ | 0.73 | 0.84 |
| | 6UZ0 | 7 | 0.12 | $2.9 \cdot 10^{-5}$ | 1.2 | 0.94 |
| Pore | 6JPA, position 2 | 13 | $1.2 \cdot 10^{-4}$ | $1.9 \cdot 10^{-5}$ | 2.2 | 2.1 |
| | 6JPB | 7 | $6.4 \cdot 10^{-4}$ | $7.8 \cdot 10^{-6}$ | 2.5 | 1.1 |

**Appendix S1.** Example Antechamber input for AM1BCC ligand minimization

```
antechamber -i $ligand_input -fi mol2 -o $ligand_output -fo mol2 -c bcc -nc $net_charge -at sybyl

```

where `$ligand_input` is the .mol2 ligand input, `$ligand_output` is the .mol2 ligand output name, and `$net_charge` is the net charge of the ligand

**Appendix S2.** Example Bash script generating ligand conformers with OpenEye Omega and ligand parameters generation for RosettaLigand

```
#!/bin/bash
# New to OpenEye? Install the openeye toolkit with Conda. (Read the OpenEye toolkit README)
# Using Python2.7? Look for pip install packages at: https://anaconda.org/OpenEye
rosetta_src={YOUR_FOLDER_PATH}/rosetta_bin_linux_2021.07.61567_bundle/main/source
openeye_bin={YOUR_FOLDER_PATH}/openeye/bin

if [ $# -lt 1 ]; then
    echo "USAGE: generate-ligand-conformers.sh <ligand name only (input in .mol2 format)>"
    exit
fi
drug=$1
dir=$(echo ${PWD})
set -v

# Set up directory structure
mkdir -p ligand
# Make ligands
pushd ligand
```

```
mkdir -p {fa,cen}/{conf1,confs,kins,withxtal}

omega="${openeye_bin}/omega2 -includeInput -commentEnergy"
$omega -in $dir/$drug.mol2 -out $drug.omega.mol2 -prefix _$drug
python ${rosetta_src}/src/apps/public/ligand_docking/assign_charges.py
< $drug.omega.mol2 > $drug.am1bcc.mol2
python ${rosetta_src}/scripts/python/public/molfile_to_params.py -c -nX00
-p$drug -k$drug.kin $drug.am1bcc.mol2
cat ${drug}_????.fa.pdb | gzip -c > fa/withxtal/${drug}_confs.fa.pdb.gz &&
( [ -f ${drug}_0002.fa.pdb ] || cp ${drug}_0001.fa.pdb ${drug}_0002.fa.pdb )
mv ${drug}_0001.fa.pdb fa/conf1/
cat ${drug}_????.fa.pdb | gzip -c > fa/${drug}_confs.fa.pdb.gz
mv ${drug}_????.fa.pdb fa/confs/
echo "PDB_ROTAMERS ${drug}_confs.fa.pdb" >> $drug.fa.params
cp $drug.fa.params fa/withxtal/
mv $drug.fa.params fa/
mv $drug.fa.kin fa/kins/
cat ${drug}_????.cen.pdb | gzip -c > cen/withxtal/${drug}_confs.cen.pdb.gz &&
 ( [ -f ${drug}_0002.cen.pdb ] || cp ${drug}_0001.cen.pdb ${drug}_0002.cen.pdb )
mv ${drug}_0001.cen.pdb cen/conf1/
cat ${drug}_????.cen.pdb | gzip -c > cen/${drug}_confs.cen.pdb.gz
mv ${drug}_????.cen.pdb cen/confs/
echo 'PDB_ROTAMERS $drug_confs.cen.pdb' >> $drug.cen.params
```

```
cp $drug.cen.params cen/withxtal/
mv $drug.cen.params cen/
mv $drug.cen.kin cen/kins/
popd

```

where `rosetta_src` is the file path to Rosetta software, `openeye_bin` is the file path to the OpenEye directory, and `drug` is the ligand name only (input in .mol2 format).

**Appendix S3**. Example input for small molecule crystal structure prediction

```
#!/bin/bash
## Formatted USAGE conditions and variables, reformatted to reflect example script from
Park H. et al 2021. https://dx.doi.org/10.1021/acs.jctc.0c01184

if [ $# -lt 2 ]; then
      echo "USAGE: ./bjh-GADock-dockrigid.sh <Ligand.pdb> <File handle for ligand
params>”
      exit
fi

## For De novo prediction, sgin=random_achiral or random_chiral & random=1. For native
prediction, sgin=input & random=0

rosetta_execute={ROSETTA_PATH}/main/source/bin/rosetta_scripts.default.linuxgccrelease
rosetta_database={ROSETTA_PATH}/main/database
sgin=random_achiral
random=1
```

```
nstruct=1

## DO NOT EDIT BELOW THIS LINE ##
PDB=${1}
file_handle=${2}
genpot=${PWD}/genpot_condensed.wts
$rosetta_execute \
    -in:path:database $rosetta_database \
    -parser:protocol ${PWD}/crystdock.xml \
    -parser:script_vars sg=$sgin random=$random \
    -s ${PWD}/${PDB} \
    -nstruct $nstruct \
    -gen_potential \
    -in:file:fullatom \
    -crystal_refine \
    -overwrite \
    -fa_max_dis 10.0 \
    -elec_max_dis 10.0 \
    -extra_res_fa ./seed_files/${file_handle}.params \
    -out:file:silent ${i}_GADock-sc_${file_handle}_.silent \
    -out:file:silent_struct_type binary \
    -out:file:scorefile GADock-sc_${file_handle}.sc \

```

**Appendix S4**. Example Python script for determining maximum ligand atom-atom distance for a set of conformers.

```
import warnings
import math
import pandas
import os
```

```python
warnings.simplefilter(action='ignore', category=FutureWarning)

# EDIT ME
base_path = 'example/path/'
seed_files_path = base_path + 'folder_containing_params/'
# END EDITS

# identify the params file
params_file = [f for f in os.listdir(seed_files_path) if f.endswith('.params')][0]
# in the params file, identify the file located on the same line as PDB_ROTAMERS
with open(seed_files_path + params_file, 'r') as f:
    for line in f:
        if 'PDB_ROTAMERS' in line:
            pdb_rotamers_file = line.split('PDB_ROTAMERS')[1].strip()
            print(pdb_rotamers_file)

# create coordinates_df dataframe
coordinates_df = pandas.DataFrame(columns=['x', 'y', 'z'])

# initialize the max_distance variable
max_distance = 0

# read the pdb_rotamers file line by line
with open(seed_files_path + pdb_rotamers_file, 'r') as f:
    for line in f:
        # if the line starts with 'HETATM', store the x, y, and z coordinates
        if line.startswith('HETATM'):
            x = float(line[30:38].strip())
            y = float(line[38:46].strip())
            z = float(line[46:54].strip())
            coordinates_df = coordinates_df.append({'x': x, 'y': y, 'z': z},
```

```
ignore_index=True)
        if line.startswith('TER'):
            # for each coordinate, compute the distance to all other coordinates in dataframe, store the largest distance
            for index, row in coordinates_df.iterrows():
                for index2, row2 in coordinates_df.iterrows():
                    if index != index2:
                        distance = math.sqrt((row['x'] - row2['x']) ** 2 + (row['y'] - row2['y']) ** 2 + (row['z'] - row2['z']) ** 2)
                        if distance > max_distance:
                            max_distance = distance
            # reset the coordinates_df for the next conformer
            coordinates_df = pandas.DataFrame(columns=['x', 'y', 'z'])

print('The max distance is ' + str(max_distance))
exit()

```

**Appendix S5**. Example Bash script for RosettaLigand input.

```
#!/bin/bash

if [ $# -lt 3 ]; then
      echo "USAGE: ./RosettaLigand.sh <NativePDB.pdb> <Test Protein Ligand Complex > <File handle for params>"
      exit
fi

rosetta_execute={PATH_TO_ROSETTA}/main/source/bin/rosetta_scripts.static.linuxgccrelease
rosetta_database={PATH_TO_ROSETTA}}/main/database

```

```
## DO NOT EDIT BELOW THIS LINE ##
nativepdb=${1}
Testpdb=${2}
file_handle=${3}

$rosetta_execute \
      -in:path:database $rosetta_database \
      -s ${Testpdb} \
      -parser:protocol dock.xml \
      -parser:script_vars nativepdb=${nativepdb} \
      -extra_res_fa ${file_handle}.params \
      -ex1 \
      -ex2 \
      -ex1aro \
      -extrachi_cutoff 1 \
      -no_optH false \
      -flip_HNQ true \
      -ignore_ligand_chi true \
      -nstruct 10 \
      -overwrite \
      -out:file:silent_struct_type binary \
      -out:file:scorefile _Rosetta_ligand_${file_handle}.sc \
      -out:file:silent _Rosetta_ligand_${file_handle}.silent \
      -mistakes:restore_pre_talaris_2013_behavior \
      -score:analytic_etable_evaluation true

```

**Appendix S6**. Example XML for RosettaLigand

```
<ROSETTASCRIPTS>
  <SCOREFXNS>
    <ScoreFunction name="ligand_soft_rep" weights="ligand_soft_rep">
      <Reweight scoretype="fa_elec" weight="0.42"/>
      <Reweight scoretype="hbond_bb_sc" weight="1.3"/>
      <Reweight scoretype="hbond_sc" weight="1.3"/>
      <Reweight scoretype="rama" weight="0.2"/>
    </ScoreFunction>
    <ScoreFunction name="hard_rep" weights="ligand">
      <Reweight scoretype="fa_intra_rep" weight="0.004"/>
      <Reweight scoretype="fa_elec" weight="0.42"/>
      <Reweight scoretype="hbond_bb_sc" weight="1.3"/>
      <Reweight scoretype="hbond_sc" weight="1.3"/>
      <Reweight scoretype="rama" weight="0.2"/>
    </ScoreFunction>
  </SCOREFXNS>

  <LIGAND_AREAS>
    <LigandArea name="inhibitor_dock_sc" chain="X" cutoff="6.0" add_nbr_radius="false" all_atom_mode="true" minimize_ligand="10"/>
    <LigandArea name="inhibitor_final_sc" chain="X" cutoff="6.0" add_nbr_radius="false" all_atom_mode="true"/>
    <LigandArea name="inhibitor_final_bb" chain="X" cutoff="7.0" add_nbr_radius="false" all_atom_mode="true" Calpha_restraints="0.3"/>
  </LIGAND_AREAS>

  <INTERFACE_BUILDERS>
    <InterfaceBuilder name="side_chain_for_docking" ligand_areas="inhibitor_dock_sc"/>
    <InterfaceBuilder name="side_chain_for_final" ligand_areas="inhibitor_final_sc"/>
```

```
      <InterfaceBuilder name="backbone" ligand_areas="inhibitor_final_bb" extension_window="3"/>
    </INTERFACE_BUILDERS>

    <MOVEMAP_BUILDERS>
      <MoveMapBuilder name="docking" sc_interface="side_chain_for_docking" minimize_water="false"/>
      <MoveMapBuilder name="final" sc_interface="side_chain_for_final" bb_interface="backbone" minimize_water="false"/>
    </MOVEMAP_BUILDERS>

    <SCORINGGRIDS ligand_chain="X" width="34">
      <ClassicGrid grid_name="ClassicGrid" weight="1.0"/>
    </SCORINGGRIDS>

    <MOVERS>
      <Transform name="transform" chain="X" box_size="7.0" move_distance="0.2" angle="20" cycles="500" repeats="1" temperature="5"/>
      <HighResDocker name="high_res_docker" cycles="6" repack_every_Nth="3" scorefxn="ligand_soft_rep" movemap_builder="docking"/>
      <FinalMinimizer name="final" scorefxn="hard_rep" movemap_builder="final"/>
      <InterfaceScoreCalculator name="add_scores" chains="X" scorefxn="hard_rep" native="%%nativepdb%%"/>

      <MultiplePoseMover name="add_filter">
        <ROSETTASCRIPTS>
        <SCOREFXNS>
          <ScoreFunction name="hard_rep" weights="ligand">
            <Reweight scoretype="fa_intra_rep" weight="0.004"/>
            <Reweight scoretype="fa_elec" weight="0.42"/>
            <Reweight scoretype="hbond_bb_sc" weight="1.3"/>
```

```
            <Reweight scoretype="hbond_sc" weight="1.3"/>
            <Reweight scoretype="rama" weight="0.2"/>
          </ScoreFunction>
        </SCOREFXNS>

        <FILTERS>
          <LigInterfaceEnergy name="LigInterface"  scorefxn="hard_rep" confidence="0.0" />
          <DSasa name="DSASA" lower_threshold="0.0" upper_threshold="1.0" confidence="0.0" />
        </FILTERS>

        <PROTOCOLS>
          <Add filter_name="DSASA" />
          <Add filter_name="LigInterface" />
        </PROTOCOLS>
       </ROSETTASCRIPTS>
     </MultiplePoseMover>
   </MOVERS>

   <PROTOCOLS>
     <Add mover_name="transform"/>
     <Add mover_name="high_res_docker"/>
     <Add mover_name="final"/>
     <Add mover_name="add_scores"/>
     <Add mover_name="add_filter"/>
   </PROTOCOLS>

</ROSETTASCRIPTS>
```

**Appendix S7**. Example input to convert RosettaLigand silent file to pdb file.

```
#!/bin/bash

{ROSETTA_PATH}/main/source/bin/extract_pdbs.default.macosclangrelease \
    -database {ROSETTA_PATH}/main/database \
    -in:file:silent_struct_type binary \
    -ignore_unrecognized_res 1 \
    -in:file:fullatom \
    -missing_density_to_jump true \
    -extra_res {PARAMS_FILE} \
    -in:file:silent {SILENT_FILE} \
    -tags {POSE_TAG} \
    -mistakes:restore_pre_talaris_2013_behavior \
    -score:analytic_etable_evaluation true

```

**Appendix S8**. Example input for GALigandDock

```
#!/bin/bash

## Formatted to reflect example script from Park H. et al 2021.
https://dx.doi.org/10.1021/acs.jctc.0c01184

if [ $# -lt 3 ]; then
```

```
	echo "USAGE: GADock.sh <NativePDB.pdb> <Test Protein Ligand Complex > <File handle for params>"
	exit
fi

mkdir scorefile
mkdir results_silent
rosetta_execute={ROSETTA_PATH}/main/source/bin/rosetta_scripts.static.linuxgccrelease
rosetta_database={ROSETTA_PATH}/main/database

## DO NOT EDIT BELOW THIS LINE ##
nativepdb=${1}
TestComplex=${2}
file_handle=${3}

$rosetta_execute \
    -in:path:database $rosetta_database \
    -s ./seed_files/${TestComplex} \
    -parser:protocol ./seed_files/GA-Dock-dockflex_blind.xml \
    -parser:script_vars initial_pool=./seed_files/${TestComplex} nativepdb=./seed_files/${nativepdb} \
    -extra_res_fa ./seed_files/${file_handle}.params \
    -gen_potential \
    -overwrite \
    -crystal_refine \
    -scale_rb 10.0 \
    -score::hb_don_strength hbdon_GENERIC_SC:1.45 \
    -score::hb_acc_strength hbacc_GENERIC_SP2SC:1.19 \
    -score::hb_acc_strength hbacc_GENERIC_SP3SC:1.19 \
    -score::hb_acc_strength hbacc_GENERIC_RINGSC:1.19 \
    -no_autogen_cart_improper \
```

```
     -out:file:silent_struct_type binary \
     -out:file:scorefile ./scorefile/GADock-sc_${file_handle}.sc \
     -out:file:silent ./results_silent/GADock-sc_${file_handle}_.silent \
     -mute core basic protocols.relax
```

**Appendix S9**. Example XML for GALigandDock

```
<ROSETTASCRIPTS>
      <SCOREFXNS>
        <ScoreFunction name="relaxscore" weights="beta_genpot_cart"/>
        <ScoreFunction name="dockscore" weights="beta_genpot">
              <Reweight scoretype="fa_rep" weight="0.2"/>
              <Reweight scoretype="coordinate_constraint" weight="0.1"/>
        </ScoreFunction>
      </SCOREFXNS>
      <RESIDUE_SELECTORS>
      </RESIDUE_SELECTORS>
      <TASKOPERATIONS>
      </TASKOPERATIONS>
      <MOVE_MAP_FACTORIES>
      </MOVE_MAP_FACTORIES>
      <SIMPLE_METRICS>
      </SIMPLE_METRICS>
      <FILTERS>
      </FILTERS>
      <MOVERS>
        <GALigandDock name="dock" runmode="dockflex" scorefxn="dockscore"
scorefxn_relax="relaxscore" grid_step="0.25" padding="7.0" hashsize="8.0" subhash="3"
```

```
final_exact_minimize="sc" random_oversample="10" rotprob="0.9" rotEcut="100" sidechains="aniso" initial_pool="%%initial_pool%%" nativepdb="%%nativepdb%%" >
        </GALigandDock>
        <MultiplePoseMover name="add_filter">
          <ROSETTASCRIPTS>
            <SCOREFXNS>
              <ScoreFunction name="relaxscore" weights="beta_genpot_cart"/>
            </SCOREFXNS>
            <FILTERS>
              <LigInterfaceEnergy name="LigInterface"  scorefxn="relaxscore" confidence="0.0" />
              <DSasa name="DSASA" lower_threshold="0.0" upper_threshold="1.0" confidence="0.0" />
            </FILTERS>
            <PROTOCOLS>
              <Add filter_name="DSASA" />
              <Add filter_name="LigInterface" />
            </PROTOCOLS>
          </ROSETTASCRIPTS>
        </MultiplePoseMover>
      </MOVERS>
      <PROTOCOLS>
          <Add mover_name="dock" />
          <Add mover_name="add_filter" />
      </PROTOCOLS>
      <OUTPUT scorefxn="relaxscore" />
</ROSETTASCRIPTS>
```

**Appendix S10**. Example input to convert GALigandDock silent file to pdb file.

```
#!/bin/bash

{ROSETTA_PATH}/main/source/bin/extract_pdbs.default.macosclangrelease \
    -database {ROSETTA_PATH}/main/database \
    -in:file:silent_struct_type binary \
    -ignore_unrecognized_res 1 \
    -in:file:fullatom \
    -missing_density_to_jump true \
    -extra_res {PARAMS_FILE} \
    -in:file:silent {SILENT_FILE} \
    -tags {POSE_TAG} \
    -beta \
```
